# A global map of the Zika virus phosphoproteome reveals host-driven regulation of viral budding and cytopathogenicity

**DOI:** 10.1101/2021.11.24.469793

**Authors:** Inessa Manuelyan, Anna M. Schmoker, Boyd L. Yount, Philip Eisenhauer, Judith I. Keller, Clarissa Gold, Dante Terino, Christopher M. Ziegler, Jeff Alexander, Heather Driscoll, Edward Hutchinson, David Bhella, Christopher D. Syme, Douglas G. Widman, Mark T. Heise, Ralph S. Baric, Bryan A. Ballif, Jason W. Botten

**Author notes:** Correspondence (J.W.B.); (J.W.B.). Current address: Dartmouth Cancer Center, Geisel School of Medicine, Dartmouth College, 1 Medical Center Drive, Lebanon, NH, USA. Current address: Medical Microbiology and Immunology Dept, University of Alberta, Edmonton, AB, Canada. Current address: Imanis Life Sciences, Rochester, MN, USA. Current address: Seqirus, Inc. 50 Hampshire Street, Cambridge, MA, USA.

## Abstract

Flaviviruses are enveloped, positive-strand RNA viruses that cause millions of infections in the human population annually. Although Zika virus (ZIKV) had been detected in humans as early as the 1950s, its reemergence in South America in 2015 resulted in a global health crisis. While flaviviruses encode 10 proteins that can be post-translationally modified by host enzymes, little is known regarding post-translational modifications (PTMs) of the flavivirus proteome. We used mass spectrometry to comprehensively identify host-driven PTMs on the ZIKV proteome. This approach allowed us to identify 43 PTMs across 8 ZIKV proteins, including several that are highly conserved within the *Flavivirus* genus. Notably, we found two phosphosites on the ZIKV envelope protein that are functionally important for viral propagation. Both appear to regulate viral budding, while one also impacts ZIKV cytopathogenicity. Additionally, we discovered host kinases that interact with ZIKV proteins and determined that Bosutinib—an FDA-approved tyrosine kinase inhibitor that targets some of these host kinases—impairs ZIKV growth, in part by blocking phosphorylation of a tyrosine residue on the envelope protein. Thus, we have defined a high-resolution map of host-driven PTMs on ZIKV proteins as well as cellular interacting kinases, uncovered novel mechanisms of host driven-regulation of ZIKV budding and cytopathogenicity, and identified an FDA-approved inhibitor of ZIKV growth.

## Introduction

ZIKV is a mosquito-borne, enveloped, positive-strand RNA virus that encodes three structural proteins (capsid, C; membrane, prM; and envelope glycoprotein, E) and seven nonstructural (NS) proteins (NS1, NS2A, NS2B, NS3, NS4A, NS4B, and NS5; Fig 1A). ZIKV was first discovered in 1947 and confirmed as a human pathogen in 1952 (1). ZIKV belongs to the *Flavivirus* genus, which has a large disease burden on the human population because of notable members like dengue (DENV), West Nile (WNV), and yellow fever viruses (YFV). Though considered a benign disease for most of its known history, ZIKV emerged in South America in 2015 and resulted in a global health crisis that lasted for more than a year and caused close to 1 million suspected and confirmed infections, but likely infected over 130 million people by the end of 2018 (2, 3). Most ZIKV infections are asymptomatic, while approximately 20% of those infected experience mild disease. In rarer cases, detrimental neurological complications arise in the form of Guillain-Barré syndrome as well as fetal microcephaly (4, 5). Infants afflicted with microcephaly because of ZIKV infection suffer developmental and health consequences into childhood often with no countermeasures available; those born with severe microcephaly require constant care.

**Figure 1.**
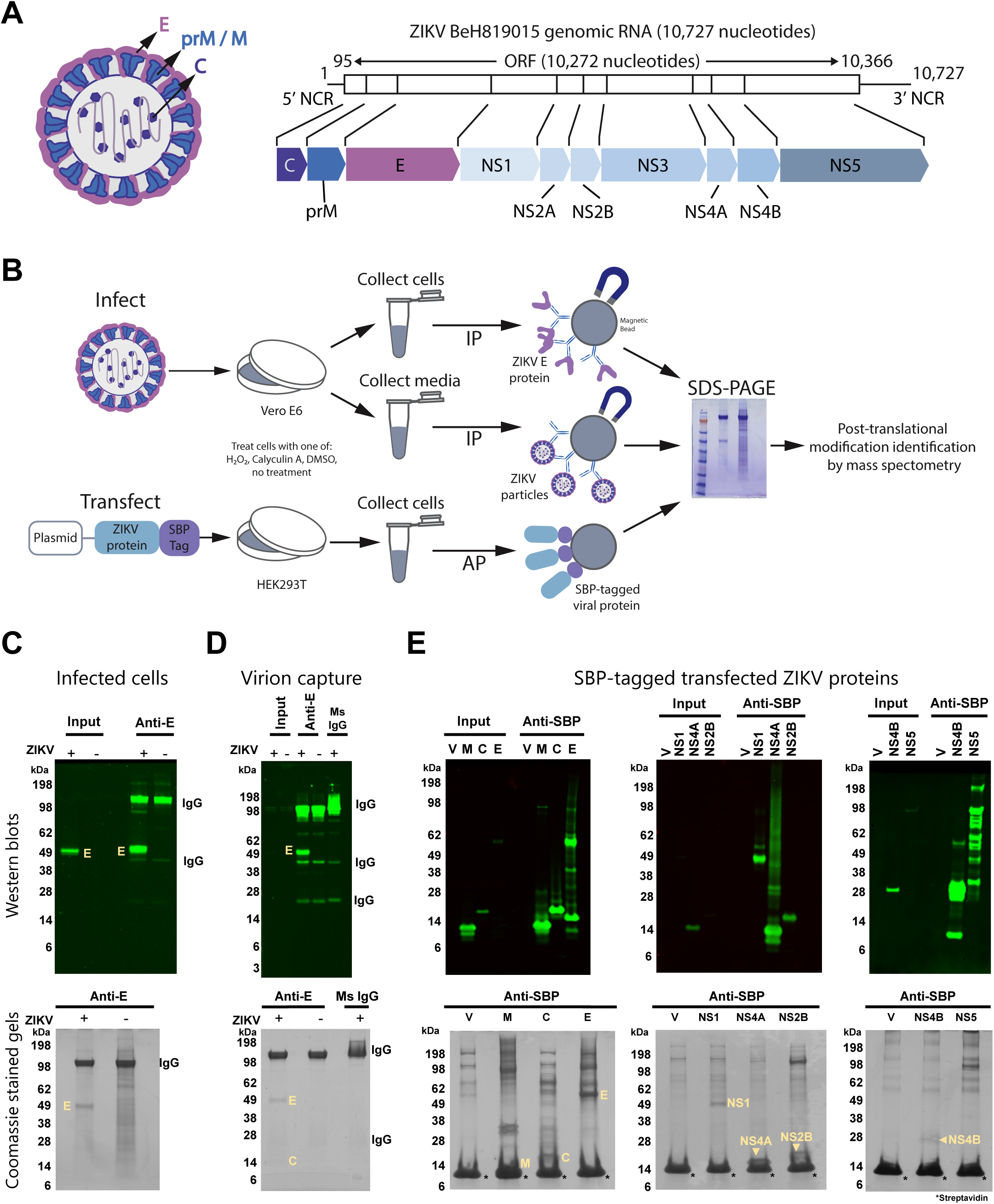
Discovery of post-translational modifications on the ZIKV proteome. (A) Depictions of i) a ZIKV particle with its structural proteins indicated and ii) the organization of the ZIKV positive-strand RNA genome and its encoded ORFeome. (B) Overview of the workflow for discovery of post-translational modifications on ZIKV proteins in the settings of viral infection or plasmid-based expression of ZIKV proteins. IP, immunoprecipitation; AP, affinity purification. (C, D, and E) Western blots and Coomassie stained SDS-page gels of affinity purified ZIKV E from ZIKV infected cells (C), authentic ZIKV particles (D), or the indicated SBP-tagged ZIKV proteins (E). For (C and D), Vero E6 cells were infected with ZIKV strain BeH819015 at an MOI of 0.1 or mock infected. Cells (C) and media (D) were collected 72 hours and 96 hours later, respectively. In (C), infected cells were lysed in Triton-X buffer and intracellular ZIKV E was purified using the anti-E antibody 4G2 and magnetic protein G beads. As a control, we performed immunoprecipitation of uninfected (mock) cellular lysates with the 4G2 antibody. In (D), viral particles were immunoprecipitated from clarified supernatants using the 4G2 antibody and magnetic protein G beads. Control conditions included (i) immunoprecipitation of virus-containing supernatants with a non-specific mouse IgG antibody and (ii) immunoprecipitation of an uninfected (mock) supernatant with the 4G2 antibody. For (E), to purify ZIKV proteins expressed from plasmid, we engineered plasmids to encode the indicated ZIKV ORFs with a C-terminal streptavidin-binding protein (SBP) tag. HEK293T cells were transfected with a plasmid encoding the indicated SBP-tagged ZIKV protein or, as a control, an empty vector (V). Two days after transfection, the cells were lysed in Triton-X buffer and ZIKV proteins were affinity purified using streptavidin-coated magnetic beads (labeled as Anti-SBP in panels). In panels (C-E), for both antibody-and streptavidin-based purifications, captured protein mixtures were orthogonally separated using SDS-PAGE, followed by in-gel processing for subsequent mass spectrometry analysis to discover post-translational modifications or interacting host proteins. The molecular weight in kDa, viral proteins (yellow text and arrows), and immunoglobulin bands are labeled. Note that the asterisks in (E) indicate monomeric streptavidin that was eluted off the streptavidin beads when boiled.

Surveys of post-translational modifications (PTMs) on viral proteins have previously been used to understand how viruses affect cells and vice versa. Viral proteins are multifunctional and PTMs are an important host-driven process that can regulate viral protein functionality. Discovery of PTMs can potentially elucidate novel viral protein functions. For example, in human immunodeficiency virus and lymphocytic choriomeningitis virus (LCMV), phosphorylation of viral proteins has been shown to be important in key processes like viral particle budding and regulation of budding of different classes of viral particles (6–9). There have been a small number of flavivirus phosphorylation sites discovered which demonstrate that phosphorylation of viral proteins can influence the host’s innate immune system or evade it, affect functionality of a viral protein, impact viral protein-protein interactions, or regulate the replication of viral RNA (10–14). These important discoveries emerged from studies of NS5 proteins of several species of flaviviruses. However, there remains an expansive terrain of other flavivirus proteins. Importantly, identification of novel PTMs and their functional relevance is anticipated to reveal unexplored avenues for pharmacological therapeutics in these diseases. To address this gap in knowledge, herein we performed a large-scale mass spectrometry analysis of the ZIKV proteome to broadly map the PTM landscape of ZIKV. Additionally, we mapped host kinases that interact with the ZIKV proteome and tested whether FDA-approved inhibitors of interacting kinases can be used to inhibit ZIKV growth. Finally, we functionally interrogated a subset of the identified PTMs to determine their importance for ZIKV propagation and, where relevant, mapped the stage of the viral life cycle impacted.

## Results

### Discovery of post-translational modifications (PTMs) on the Zika virus proteome

A primary goal of this study was to identify host-driven PTMs that occur on ZIKV proteins produced by mammalian cells, either in the setting of cell-free virions or within cells. Where possible, we wished to examine PTMs in the setting of authentic ZIKV infection. At the time of this study, the only suitable antibody available for purification of a ZIKV protein, expressed in the context of an infected cell, was 4G2, which recognizes the ZIKV envelope (E) protein (15). This antibody was used to purify intracellular ZIKV E and its interacting host partners from infected cells (Figs 1B and 1C) or E-decorated, cell-free virions released from these same infected cells (Figs 1B and 1D). Note that immunopurified ZIKV particles contained the structural proteins E, prM, and C (Fig 1D and Supplementary Information 1). We also isolated cell-free virions from infected mosquito cells via banding on a potassium tartrate density gradient (Fig S1A). To circumvent the lack of antibodies suitable for immunoprecipitation of the remaining ZIKV proteins from cells, we used plasmids to express streptavidin-binding protein (SBP)-tagged ZIKV proteins. This allowed for efficient capture of SBP-tagged viral proteins via magnetic streptavidin beads (Figs 1B, 1E, S2, S3, and S4). To increase the likelihood of detecting phosphorylation events, transfected cells were treated with either H_2_O_2_, Calyculin A, or DMSO to inhibit tyrosine phosphatases, serine/threonine phosphatases, or to act as a control, respectively (Figs 1B, 1E, S2, S3, and S4). Samples derived from immunoprecipitation and affinity purification were subjected to SDS-PAGE to orthogonally separate protein components. Gel lanes were cut into multiple sections, and each piece was subjected to in-gel trypsin digestion. Meanwhile, ultracentrifuged purified virus from mosquito cells (Fig S1B) was digested with trypsin using filter-assisted sample preparation. All resulting tryptic peptides were subjected to liquid chromatography tandem mass spectrometry for identification of PTMs or interacting host proteins as described in the Methods and previously (9, 16). A complete list of peptides from viral proteins including those containing PTMs such as phosphorylation and ubiquitination can be found in Supplementary Information 2. Tandem mass spectra for discovered phosphorylation and ubiquitination sites are available in Supplementary Information 3 (spectra for ZIKV E T351 from virions is available in Figs S1C, S1D).

We identified 18 phosphorylation sites and 5 ubiquitination sites on ZIKV E (Table 1, Table S1, Figs 2A, and 2B). Two of these phosphorylation sites, Y61 and T351, were discovered in the context of infection (Y61 from infected Vero E6 cells and T351 from mosquito cell-derived ZIKV particles) (Fig S1D). T351 was also found on ZIKV E expressed from plasmid in HEK293T cells. The remaining PTMs were identified from plasmid-expressed viral proteins.

**Figure 2.**
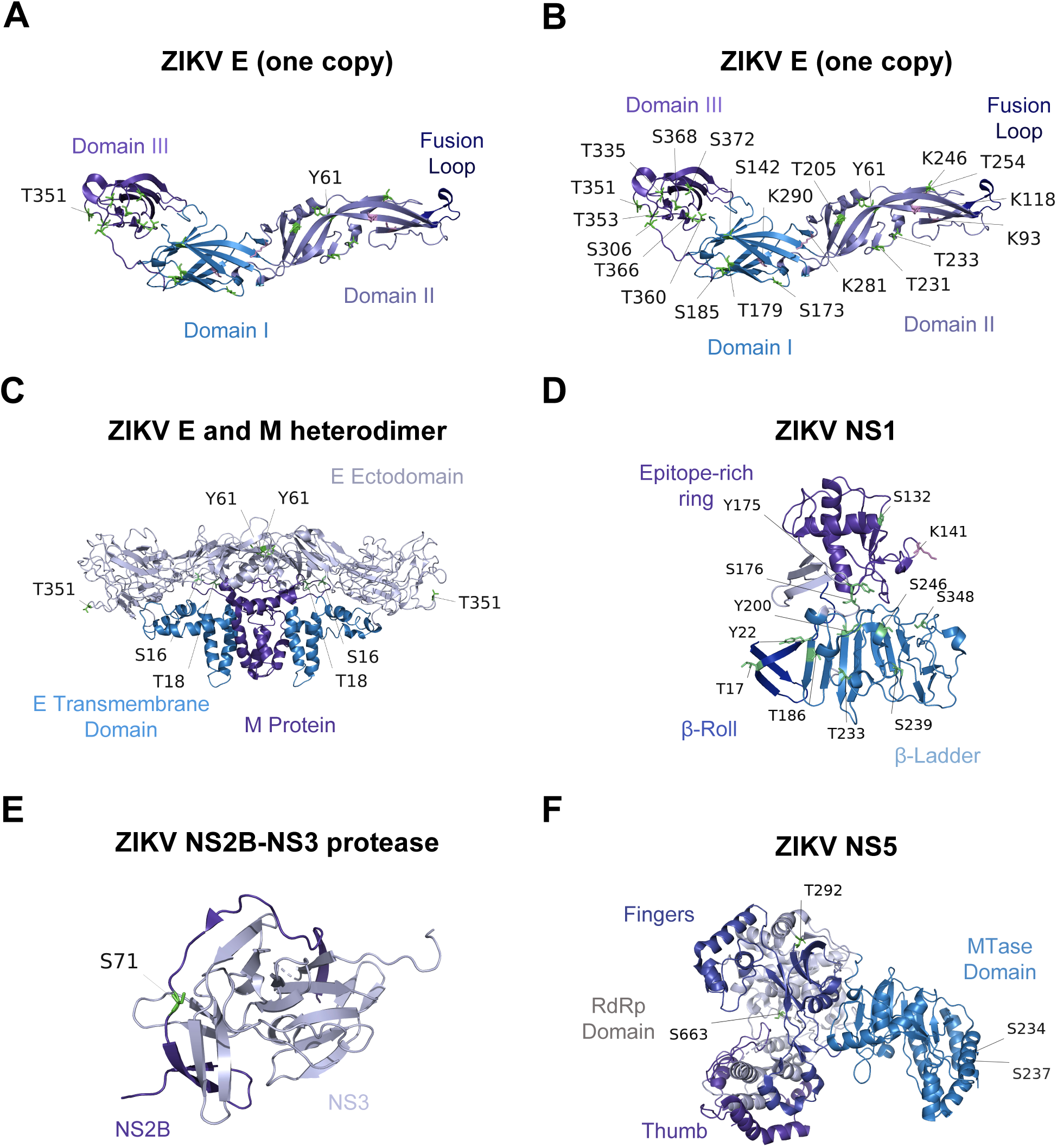
Location of post-translational modification sites on ZIKV proteins. Phosphosites and ubiquitination sites are marked in green and light purple, respectively. For each ZIKV protein, individual domains or specific features are color coded with the text label displayed in the matching color in each instance. (A and B) Location of the Y61 and T351 phosphosites on ZIKV E (A) or location of all post-translational modifications on ZIKV E (B). In both (A and B), PDB ID: 5JHM was used and one copy of E is shown. (C) ZIKV E - M heterodimer (PDB ID: 5IZ7). Y61 and T351 are labeled on the E ectodomain. ZIKV M phosphosites S16 and T18 are labeled on the M protein. (D) PTM locations on ZIKV NS1 (PDB ID: 5GS6). (E) ZIKV NS2B – NS3 protease complex shown in close conformation (PDB ID: 5GPI) with labeled phosphosites on ZIKV NS2B. (F) ZIKV NS5 (PDB ID: 5TMH) phosphosites. RdRp, RNA-dependent RNA polymerase; MTase, methyltransferase domain.

**Table 1.**
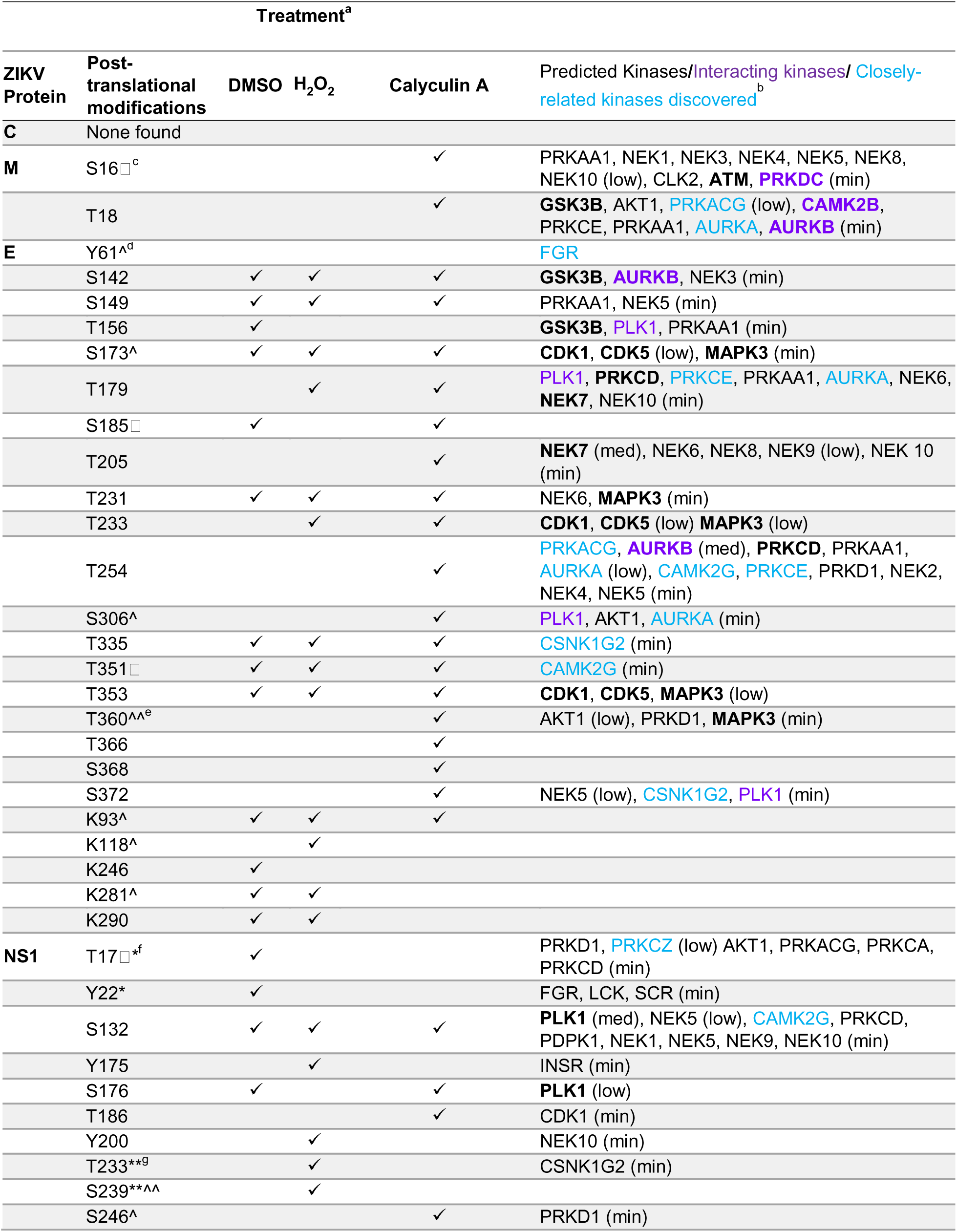
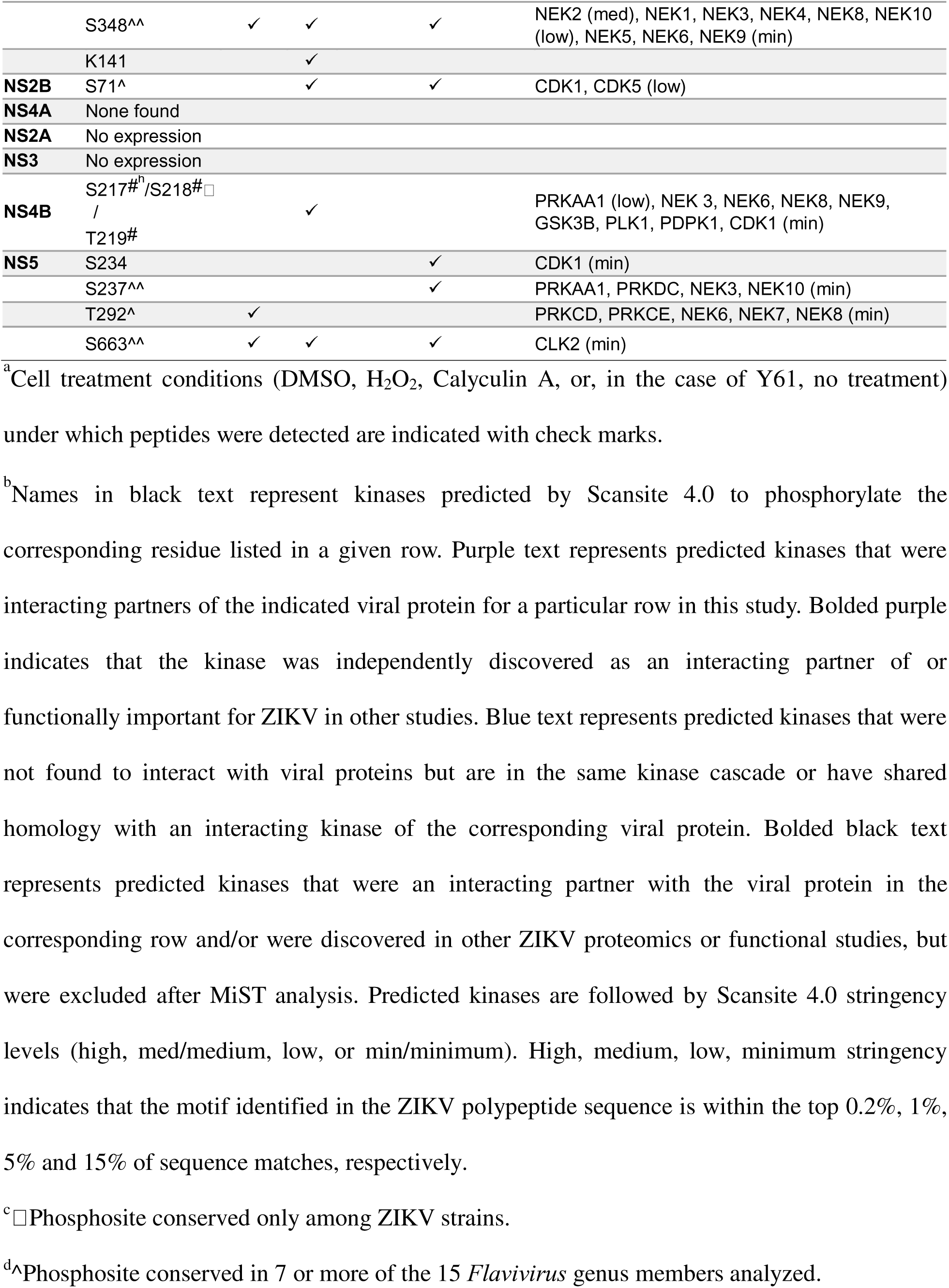
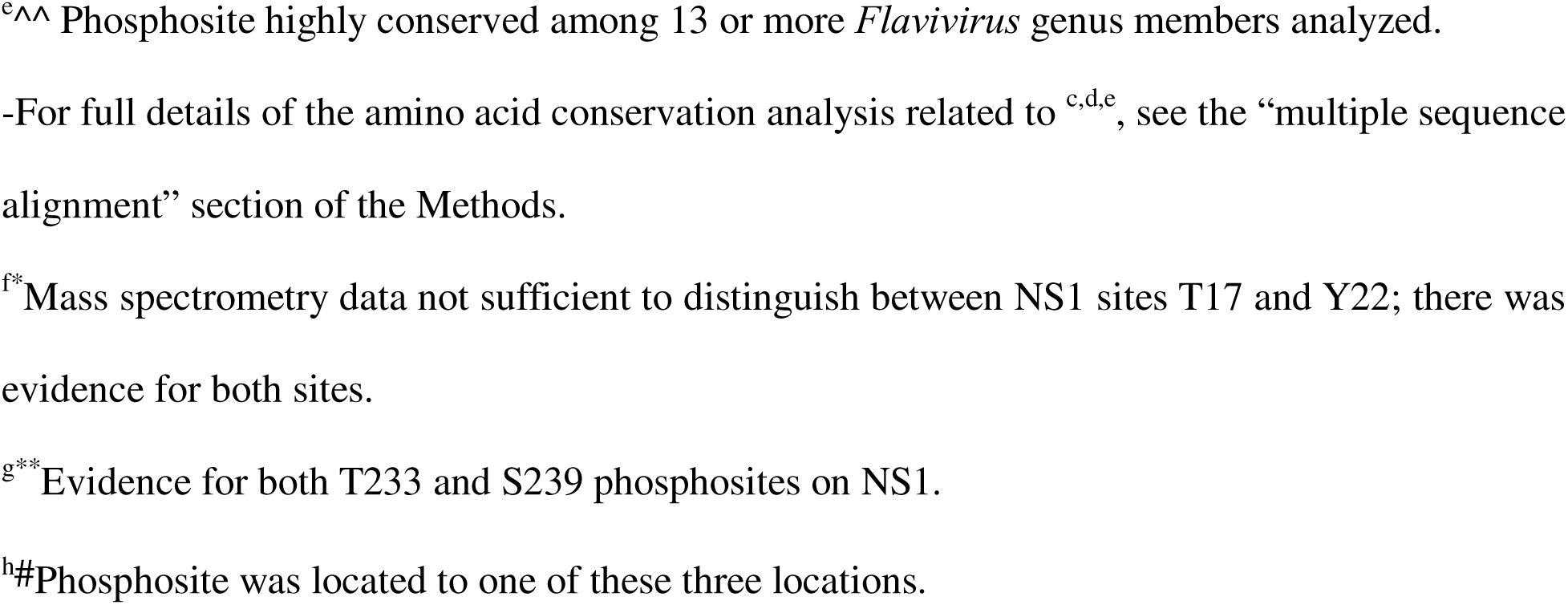
Phosphorylation and ubiquitination sites on the ZIKV proteome.

The locations of Y61, T351, and other PTMs found on ZIKV E are shown in Figs 2A and 2B. Y61 is located in Domain II of E, which is the region responsible for dimerization (17). T351 is located in Domain III, which is often a target of neutralizing antibodies (17). PTMs on E were also mapped using electrostatically labeled models (Fig S5). Figure S5A shows one “face” of the ZIKV E dimer as largely positively charged while Figure S5B shows the other largely negatively charged face. In contrast, the border of the E dimer is mostly neutral (Fig S5C). The majority of ZIKV E phosphosites were found on the neutral border of the envelope protein (Fig S5C). The two phosphorylation sites on M, S16 and T18, are positioned in close proximity to one another at a junction of where E and M associate (Fig 2C). Interactions between the M S16 and T18 phosphorylated residues with E are hard to resolve, however, because both sites are in flexible regions of M and are interacting with flexible regions of E (Fig S6A).

We identified 11 phosphorylation sites and one ubiquitination site on NS1 (Table 1, Table S1). Location mapping of PTMs on NS1 revealed that approximately half of the modified amino acid residues were in the beta-roll or beta ladder domains of NS1 (Fig 2D) important for NS1 intracellular dimerization (18) and host protein interaction (19), respectively. We were unable to successfully express and isolate adequate quantities of NS2A or NS3 to analyze for PTMs (Fig S7). Mass spectrometry revealed a phosphorylated serine at position 71 on the NS2B protein (Table 1, Table S1), a small protein known to associate with NS3 to form the ZIKV protease complex (Fig 2E). There were no PTMs found on NS4A despite an abundance of NS4A peptides observed by mass spectrometry. One phosphorylation site was detected on NS4B, though the precise location could not be determined with certainty. Our mass spectrometry data suggest its location to be at either S217, S218, or T219 (Table 1, Table S1). There are currently no models of ZIKV NS4B, thus we were unable to depict these sites within the protein structure. Lastly, we discovered four phosphorylation sites on NS5 (Table 1, Table S1). Of these, two were located in the RNA-dependent RNA polymerase region of NS5 and two in the methyl-transferase region (Fig 2F).

To determine how conserved the discovered ZIKV protein PTMs might be with other flaviviruses, we constructed a multiple sequence alignment (MSA) that included viruses spanning the *Flavivirus* genus and chose flaviviruses known to cause human disease (Supplementary Information 4; virus list also noted in the Methods section). In total, we compared 15 different flaviviruses representing tick-or mosquito-borne viruses, viruses with no known vectors, and those considered emerging pathogens. Amino acid residues were defined as conserved if they were found in at least seven of the 15 flaviviruses in our panel. Of the 43 discovered PTMs, 14 met this degree of conservation across the *Flavivirus* genus and five were conserved across nearly all flavivirus subfamilies, a result that suggests they may be fundamentally important for many species of flaviviruses (Table 1). Notably, four sites were found only in ZIKV isolates (Table 1).

### Tyrosine 61 and threonine 351 on the ZIKV E protein are critical for viral growth

We next sought to determine whether the PTMs identified were important for viral propagation. We focused on two ZIKV E phosphosites, Y61 and T351, because they were found in both infection-derived and plasmid-expressed contexts. Y61 was discovered from infected cells while T351 was found in virions and on plasmid-expressed ZIKV E. Further, these sites represented residues that are either highly conserved in flaviviruses (Y61) or unique to ZIKV (T351) (Table 1; Supplementary Information 4). To evaluate their importance in viral propagation, we employed a ZIKV reverse genetics system (20) to generate recombinant viruses encoding phosphoablative or phosphomimetic mutations at these sites (Fig 3A). We then used these recombinant viruses in multi-step growth curves to determine how each change impacted viral fitness. Throughout this manuscript the terms “clone” or “infectious clone” are used as general shorthand for recombinant virus generated from a ZIKV infectious cDNA clone using the reverse genetics system. The terms, “mutant”, “mutant virus”, or “mutant clone” are used as shorthand to refer to recombinant viruses carrying mutations at phosphorylation sites of interest.

**Figure 3.**
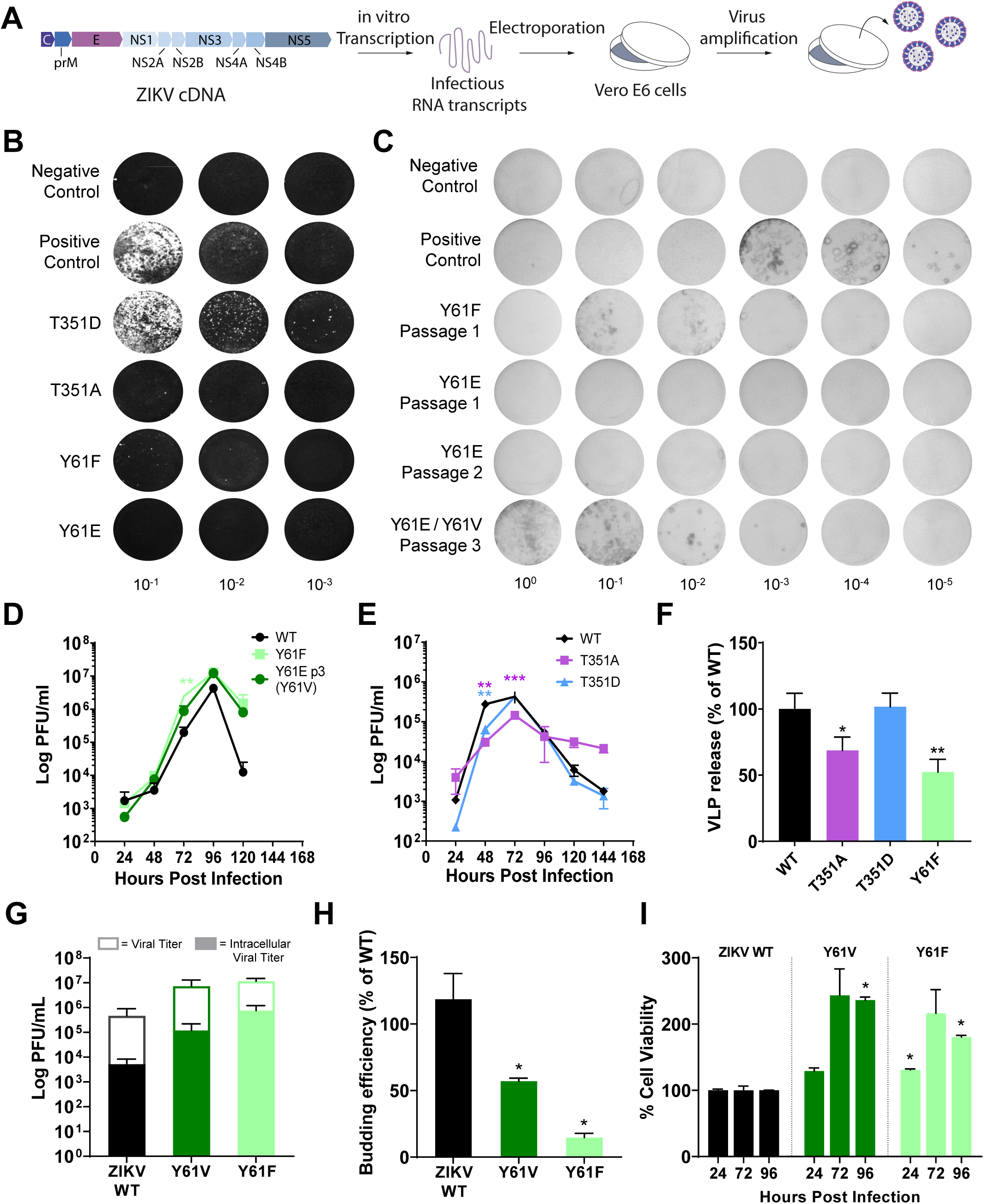
Tyrosine 61 and threonine 351 on the ZIKV E protein are critical for viral propagation at the stage of viral budding. (A-C) Overview of the reverse genetics approach used to recover infectious ZIKV clones bearing mutations at Y61 or T351 in the E protein (A). For each recovery attempt, full-length infectious RNAs were transcribed from a cDNA template of the viral genome. The RNAs were electroporated into Vero E6 cells and 3 days later supernatants from these cultures were screened for the presence of infectious ZIKV particles by either viral plaque assay (B) or focus assay (C). In (C), the rZIKV E Y61E clone was passaged 2 additional times due to the absence of detectable plaques or foci after passage 1. The Y61E passage 3 material showed infectious virus and this rLCMV had mutated from a glutamic acid to a valine at position 61. (D and E) The growth kinetics of the indicated rZIKV E Y61 mutants (D) or T351 mutants (E) was determined by infecting Vero E6 cells at an MOI of 0.001 (D) or A549 cells at an MOI of 0.01 (E) with these viruses and measuring the infectious particles released by plaque assay at 24, 48, 72, 96, 120, or 144 hours after infection. Data represent the mean±SEM from three (D) or two (E) independent biological replicates. For statistical analysis, data were analyzed via mixed-effect analysis (two-way ANOVA) with Holm-Sidak’s test for multiple comparisons in Prism 8.0. (F) The budding activity of WT, Y61-or T351-mutant E proteins was measured by a virus-like particle (VLP) assay. VLPs were generated by transfection of HEK293T cells with plasmid that encodes ZIKV prM and E (either WT E or the indicated E mutants). The quantity of ZIKV E released into the supernatant vs retained in cells was measured by quantitative western blot and budding efficiency was determined as described in the Methods. Data represent the mean±SEM from three (Y61 mutants) or four (T351 mutants) independent biological replicates. Statistical significance was determined in Prism 8.0 software using a one-way ANOVA followed by Fisher’s LSD test. (G) The intracellular and extracellular titers of WT, Y61V, and Y61F were measured by plaque assay. Vero E6 cells were infected with the indicated virus at an MOI of 0.001 for 96 hours. Supernatants were collected, infected cells were lysed with distilled (Milli-Q) water to release intracellular virions, as described in Methods, and titers were determined by plaque assay. Data represent the mean±SEM from two independent biological replicates. (H) Budding efficiency of authentic virions was calculated by normalizing the ratio of mutant extracellular to intracellular infectious titers (shown in panel G) to the ratio of the WT extracellular to intracellular infectious titers, as described in the Methods. Statistical significance was determined in Prism 8.0 software using a one-way ANOVA with Holm-Sidak’s test for multiple comparisons. (I) Cell viability after infection was measured by MTT assays. Vero E6 cells were infected with ZIKV WT, Y61V, or Y61F virus at an MOI of 0.001 and cell viability was determined by MTT assay at 24, 72, and 96 hours post infection, and normalized to the ZIKV WT condition. Data represent the mean±SEM from two biological replicates. Statistical significance was determined in Prism 8.0 software via two-way ANOVA with Holm-Sidak’s test for multiple comparisons. For the indicated statistical tests in (D-F): *, P<0.05; **; P<0.01; ***; P<0.001.

ZIKV E Y61 was mutated to a negatively-charged glutamic acid (Y61E) to reflect a permanent state of phosphorylation (i.e. a phosphomimetic) and to a phenylalanine (Y61F) to represent a non-phosphorylatable state. We recovered the non-phosphorylatable Y61F mutant virus but, interestingly, the phosphomimetic Y61E mutant appeared to be nonviable as assessed by plaque assay (Fig 3B). Recovery of a Y61E mutant clone was attempted three times without success, despite successful recovery of a wild-type clone each time. To account for the possibility that the Y61E mutant virus could be propagating but unable to form plaques (and therefore undetectable by plaque assay), we screened for the presence of infectious virus by focus assay (Fig 3C). In parallel, to account for the possibility that the virus was replicating but not with sufficient kinetics to be detectable by focus or plaque assay after the initial recovery passage, we blind-passaged Y61E three times. This approach led to the successful isolation of a Y61E recombinant virus capable of forming foci beginning at passage 3 (Fig 3C). However, sequencing of this infectious clone at passage 3 revealed that the glutamic acid (Y61E) had mutated to a valine (Y61V), suggesting that there is selection pressure to avoid a permanent negative charge at position 61.

To address whether phosphorylation of Y61 influences viral fitness, we performed a multi-step growth curve comparing the phosphomutant clones Y61F and Y61V to a wild-type clone. Interestingly, both the Y61F and Y61V infectious clones grew with accelerated kinetics and produced higher quantities of infectious virus compared to the wild-type clone (Fig 3D; Y61F p = 0.006).

ZIKV E T351 was mutated to an aspartic acid (T351D) to mimic a state of phosphorylation or an alanine (T351A) to act as a non-phosphorylatable mutant. Infectious clones of each mutant were successfully recovered (Fig 3B) and multi-step growth curves were performed. There was no difference between the growth of the phosphomimetic T351D mutant and the wild-type clone (Fig 3E; p = 0.964). Notably, the growth of the T351A non-phosphorylatable mutant was impaired compared to WT virus or the T351D mutant (Fig 3E; p = 0.0009). After 72 hours, the titers for the T351D and the wild-type infectious clones decreased more rapidly compared to T351A. This could be because the wild-type and T351D clones caused more cytopathic effect at this time point and therefore cells could not support further rounds of infection.

In summary, both the Y61 and T351 sites appear important for viral propagation. Our results suggest that blocking phosphorylation at Y61 enhances viral growth while blocking phosphorylation at T351A impairs it.

### Y61 and T351 on ZIKV E are required for efficient Zika virus budding

To determine the life cycle stage impacted by ZIKV E Y61 or T351, we next tested how a phosphomimetic or non-phosphorylatable mutation at each site would impact viral budding. To do so, we used a virus-like particle (VLP) assay that features a single plasmid that expresses ZIKV prM and E. Once expressed, these two proteins form ZIKV VLPs that bud from cells and can be detected in supernatant (21). The plasmid was mutated at ZIKV E Y61 to Y61E or Y61F and at T351 to T351A or T351D. The Western blots used to quantify ZIKV E expression and calculate VLP budding efficiency are shown in Fig S8. Compared to the wild-type control, VLP budding efficiency was decreased when T351 was mutated to alanine (T351A) but not aspartic acid (T351D) (Fig 3F; p = 0.04). Interestingly, we were unable to express ZIKV E protein when Y61 was mutated to Y61E (Fig S8) despite performing two bacterial transformations and evaluating several clones from each transformation. This result, combined with our inability to isolate genetically stable Y61E infectious clones (Fig 3C), suggests that a constitutive negative charge or phosphorylation at Y61 is not favorable either for protein expression or viral growth. Mutation of Y61 to F also reduced budding efficiency (Fig 3F, p = 0.006), despite the improved growth kinetics of this mutant (Fig 3D).

We next measured the budding efficiency of the Y61F and Y61V infectious clones by comparing the ratios of extracellular to intracellular infectious virions of the mutants to WT virus. Notably, the intracellular infectious titers of both non-phosphorylatable mutants were higher than the WT virus (Fig 3G). Yet, consistent with the VLP assay (Fig 3F), budding efficiency was decreased in both Y61V and Y61F mutant clones (Fig 3H, p = 0.031 and 0.015, respectively).

Taken together, our findings suggest that both the Y61 and T351 phosphosites are required for efficient ZIKV budding.

### Phosphorylation status of ZIKV E Y61 impacts ZIKV cytopathogenicity

Titers of the non-phosphorylatable mutants Y61V and Y61F remain elevated at late time points (Fig 3D), leading us to assess whether phosphorylation at Y61 impacts cell viability. Using MTT assays, we measured viability of Vero E6 cells infected with ZIKV WT, Y61V, or Y61F at 24, 72, and 96 hours post infection. Compared to the WT virus, cells infected with the Y61 non-phosphorylatable mutants had increased viability at several time points, particularly at 96 hours post infection (Fig 3I). We verified that the viruses were adhering to the same growth kinetic patterns seen in Fig 3D by simultaneously measuring viral titers (Fig S9). The highest cell viability, observed at 72 and 96 hours post infection, matched the time points where ZIKV Y61V and Y61F titers surpass WT virus (Fig S9). In summary, these results suggest that blocking host-driven phosphorylation of Y61 protects cells from ZIKV cytopathogenicity.

### Host protein kinases interact with the Zika virus proteome

Given the multiple phosphosites discovered on various ZIKV proteins, we next wished to determine the host kinases that associate with ZIKV proteins. The immunoprecipitation and affinity purification performed in Figs 1C, 1D, 1E, S2, S3, and S4 not only captured ZIKV proteins, but also their host cellular protein partners. Thus, we were able to identify host kinases that interacted with ZIKV C, M, E, NS1, NS2B, NS4A, and NS4B. In total, we found 115 cellular kinases that interacted with ZIKV proteins (Table S2). We subjected this dataset to analysis using MiST, an affinity purification mass spectrometry computational tool that scores protein-protein interactions to remove interactions that could be false positives (22). MiST analysis restricted the interactome to 40 kinases (Table S3, Fig S10) that were above a MiST threshold of 0.62. This threshold was chosen based on how inclusive it was of kinases that were independently discovered in other ZIKV proteomic or functional screens (23–28) but still stringent enough to remove low-scoring interactions. Most kinases were unique to this study, but 14 were independently discovered in other ZIKV proteomic or functional screens (Table S3, Column F). Notably, when comparing the original list of 115 kinases to these studies (23–28), an additional 18 kinases overlap with our study (Table S2, Column F). We have included the complete list of 115 kinases in Table S2 in the event that MiST analysis excluded additional, not-yet-tested, but functionally relevant interactions. Figure S10 depicts the MiST analyzed kinases and highlights their involvement in biological processes such as regulation of protein translation, neuron differentiation, cell cycle arrest, and other cellular processes which are relevant to ZIKV infections.

We next used Scansite 4.0 to predict kinases that might phosphorylate ZIKV proteins at the sites of phosphorylation identified in this study (Table 1, Tables S2 and S3, Column H).

Intriguingly, a subset of these kinases were interacting partners of the ZIKV proteins they were predicted to phosphorylate and/or were discovered in other ZIKV proteomic or functional screens (Table 1, purple text if unique to our dataset, or bolded purple text if discovered in others ZIKV studies). These were AURKB, CAM2KB, PLK1, and PRKDC (serine/threonine-protein kinase Aurora B, calcium/calmodulin-dependent protein kinase type II subunit beta, serine/threonine-protein kinase 13, and DNA-dependent protein kinase catalytic subunit, respectively). This list is more extensive when not accounting for MiST scoring (Table 1, bolded black text). Other kinases predicted by Scansite 4.0, although not found by our mass spectrometry analysis to be interacting partners, either have shared homology with or are found in the same signaling cascades as interacting kinases (Table 1, highlighted in blue). To incorporate the information above, we constructed Figure 4A which is representative of kinase interactions with MiST scores above 0.8 and highlights the broad range of kinases detected from various kinase groups or families/subfamilies, emphasizing those which were discovered in other ZIKV proteomic (bolded text), functional screens (bolded and asterisked text), and predicted by Scansite 4.0 to phosphorylate a ZIKV protein (italicized text). Figure 4A also includes inhibitors known to target the listed kinases, some of which are FDA-approved (bolded text in “Inhibitor” column). In summary, we have identified ZIKV-interacting host kinases, some of which were predicted to phosphorylate ZIKV proteins at sites discovered in this study, and a number of which that were previously confirmed as ZIKV interactors or important for ZIKV propagation (Table 1, Fig. 4A, Table S2, Table S3).

**Figure 4.**
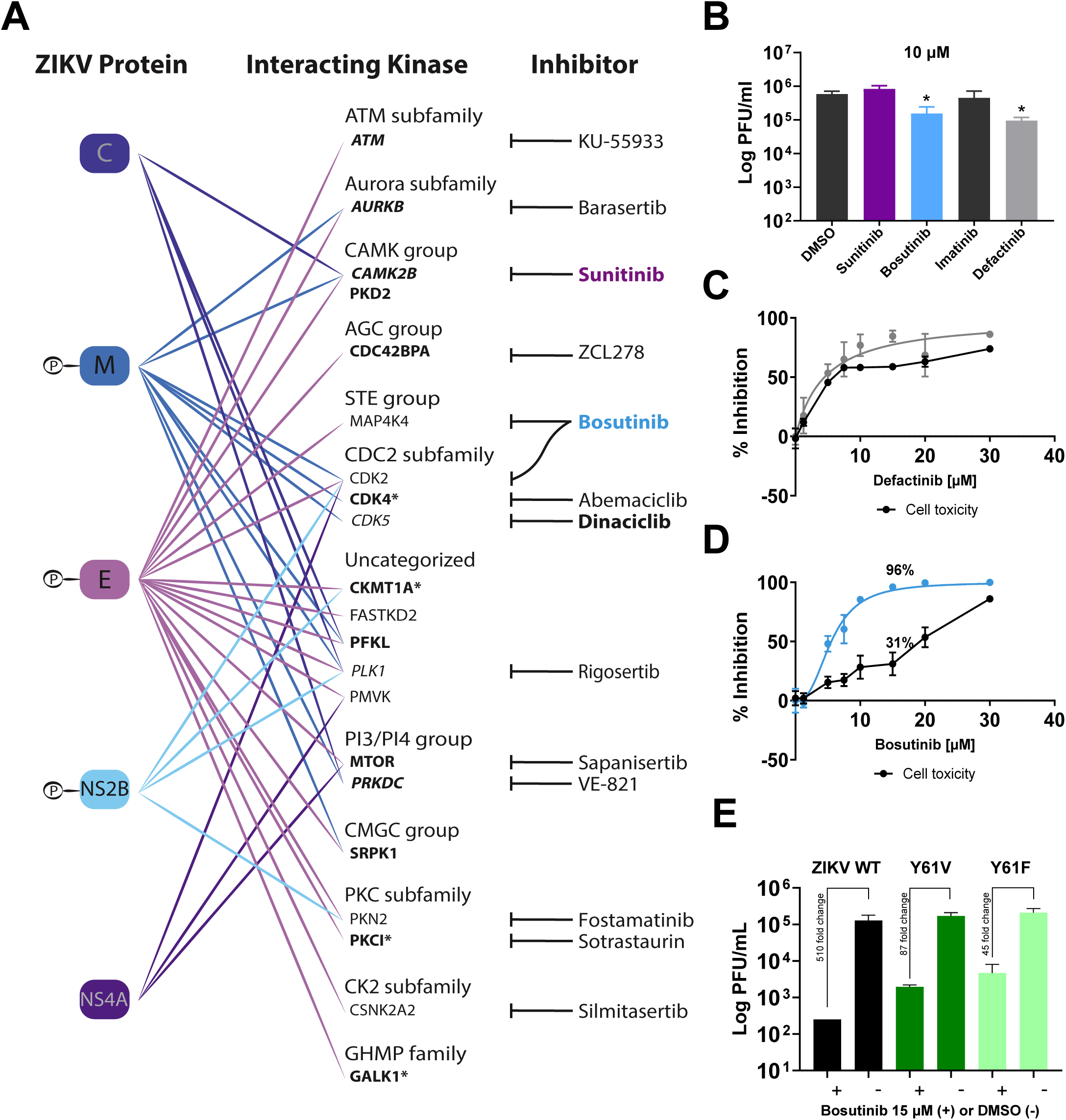
Host kinase interactions the ZIKV proteome and effects of kinase inhibitors on ZIKV growth. (A) Host kinases that interact with ZIKV proteins representative of MiST-scored interactions with a chosen threshold of >0.8. In the “Interacting Kinase” column, kinases are organized into respective groups or families. Bolded text indicates kinases that were also found to interact with ZIKV proteins in other studies. Bolded text with an asterisk were found to be functionally important in other studies. Kinases predicted by Scansite 4.0 to phosphorylate ZIKV proteins at the discovered phosphosites are italicized. To capture the range of the different kinases found, we prioritized displaying kinases discovered in published ZIKV interactomes or functional studies, then those that had the highest MiST scores. Drugs known to inhibit specific kinases are listed in the “Inhibitor” column; drugs with FDA approval are bolded. Note the full list of interacting kinases are listed in Table S2. (B) Efficacy of kinase inhibitors in restricting ZIKV growth. A549 cells were infected with ZIKV at a MOI of 0.1. One hour later, the inoculum was removed and cells were treated with 10 µM of the indicated drug or, as a control, DMSO (vehicle). Two days later infectious virus was quantified from supernatants via plaque assay. Data represent mean±SEM from three independent biological replicates. Statistical significance was determined in Prism 8.0 software via one-way ANOVA with Holm-Sidak’s test for multiple comparisons. *, P<0.05 (C and D) IC_50_ determination for Defactinib and Bosutinib inhibition of ZIKV growth. A549 cells were infected with ZIKV at a MOI of 0.1 and 1 hour later, the inoculum was removed and cells were treated with the indicated doses of Bosutinib or Defactinib. Two days later infectious virus was measured in the supernatants via plaque assay. In parallel, the cytotoxicity of each drug at these same doses was determined via MTT assay as described in the Methods. Data represent mean±SEM from three independent biological replicates. The resulting IC_50_ and selective index (SI) values are listed for each drug. Calculations of both values are described in Methods. (E) Bosutinib inhibits growth of non-phosphorylatable Y61 mutant viruses, but to a lesser degree than WT virus. A549 cells were infected with ZIKV WT, Y61V, or Y61F mutant viruses at an MOI of 0.01. One hour post infection, inoculum was removed and replaced with cell media containing either Bosutinib [15 µM] or DMSO. Supernatants were collected after 48 hours, and titers were measured via plaque assay. Data represent the mean±SEM from two biological replicates. Statistical analysis of the differences in fold changes between groups was determined in Prism 8.0 software via one-way ANOVA with Holm-Sidak’s test for multiple comparisons and showed no statistical difference.

### Inhibition of host cell kinases by Bosutinib impairs Zika virus growth and its mechanism of action is partially associated with blocking Y61 phosphorylation on ZIKV E

Considering that there are no FDA approved treatments for ZIKV, we next tested whether selected FDA-approved kinase inhibitors might be effective in restricting ZIKV growth. At 10 µM, the tyrosine kinase inhibitors Bosutinib and Defactinib both reduced ZIKV titers (Fig 4B; p = 0.03, p = 0.02, respectively) while Sunitinib and Imatinib did not (Fig 4B; p = 0.17, p = 0.35, respectively). To determine the IC_50_ of each drug, we constructed concentration-response curves. To account for drug cytotoxicity, we performed simultaneous MTT assays on cells that had been treated with each kinase inhibitor at the equivalent concentrations. Although Defactinib inhibition of ZIKV infectious virus production reached levels as high as 86%, at the same concentration of drug, cell toxicity measured 73% (Fig 4C). At Defactinib’s IC_50_ of 4.5 µM, cell toxicity was approximately 46%. The fitted curve of inhibition and cell toxicity closely followed each other, suggesting that most of the viral inhibition may have been due to nonviable cells.

After determining the CC_50_ of Defactinib to be 7.53 µM, we calculated the selectivity index for Defactinib to be 1.7. The IC_50_ of Bosutinib was 5.5 µM. In this case, at the same doses of drug, cell toxicity was approximately 17%. At higher concentrations, Bosutinib completely inhibited viral growth while cell toxicity remained at approximately 30% (Fig 4D). Given Bosutinib’s IC_50_ and CC_50_ (17.43 µM), we calculated the selectivity index to be 3.14. Because Bosutinib is a tyrosine kinase inhibitor, we next wanted to determine if ZIKV impairment by the drug was associated with blocking phosphorylation of Y61, the only phosphotyrosine discovered on the ZIKV E protein. We thus tested whether Bosutinib would impair the Y61V and Y61F mutant clones’ growth despite their inability to be phosphorylated. We found that 15 µM Bosutinib, a concentration chosen based on its full inhibition of WT virus (Fig 4D), indeed reduced Y61V and Y61F virus titers by 87-and 45-fold, respectively (Fig 4E). However, the titer reduction was not as robust compared to the 510-fold change in the WT virus. Thus, Bosutinib appears to inhibit ZIKV partially by its inhibitory effect on a host kinase responsible for Y61 phosphorylation, but also through other avenues such as blocking phosphorylation on a different ZIKV tyrosine (i.e., NS1 Y22, Y175, Y200, or one that was not detected in this study) and/or by effecting host proteins that lead to cellular events which are unfavorable for ZIKV growth. In summary, we and others have shown that ZIKV proteins interact with host kinases and some of these host kinases are predicted to phosphorylate ZIKV proteins at the discovered phosphosites in this study (Tables 1, S2, S3). Finally, we have shown that the mechanism by which Bosutinib reduces ZIKV growth (Fig 4D) may be due, in part, to blocking phosphorylation of Y61.

Overall, our results suggest that host cell kinases may be tractable antiviral targets for ZIKV.

## Discussion

Little is known regarding the diversity of host-driven PTMs that occur on flavivirus proteins or how such modifications regulate the functionality of these proteins. We addressed this deficiency by providing the first comprehensive map of phosphorylation and ubiquitination sites found on a flavivirus proteome. A major finding was that selected phosphorylation sites on the ZIKV E protein appear critical for ZIKV propagation, likely at the stage of viral assembly and release.

Moreover, we discovered that blocking phosphorylation of a tyrosine on ZIKV E reduced ZIKV cytopathogenicity, leading to a paradoxical increase in infectious virus, despite a defect in viral budding. We also identified a network of host kinases that interact with ZIKV proteins, some of which were predicted to target the identified phosphosites. Finally, we discovered that Bosutinib, an FDA approved tyrosine kinase inhibitor, likely impairs ZIKV growth partly by blocking phosphorylation of a newly discovered ZIKV E phosphotyrosine. Our data suggest that inhibition of host kinases may be an effective approach to restrict ZIKV growth.

The use of mass spectrometry allowed us to discover 43 sites of phosphorylation and ubiquitination on ZIKV proteins. Due to the lack of antibodies suitable for immunoprecipitation, many of these PTMs were discovered on plasmid-expressed viral proteins in non-infectious settings, introducing a possible limitation to this study. Our subsequent experiments focused on the ZIKV E phosphosite Y61 because it was found in the context of infection and T351 because it was discovered in virions as well as on plasmid expressed ZIKV E. Furthermore, recently, two ubiquitination sites on ZIKV E were shown to be required for viral entry and replication (29).

Notably, one of these ubiquitination sites, K281, was detected on plasmid expressed ZIKV E in our study (Table 1, Table S1) and adds further evidence that phosphosites discovered on the plasmid-expressed viral proteins merit investigation into their functional relevance.

Flavivirus budding occurs at the endoplasmic reticulum and is largely driven by prM and E. This pair of structural proteins together induce curvature in the ER membrane leading to the formation of spherical budded virions that contain the encapsidated viral genome (30). We show that the ZIKV envelope glycoprotein (ZIKV E) has at least two phosphosites, Y61 and T351, which are involved in the budding stage (Fig 3F, 3H). Thus, our data suggests that phosphorylation of E may play a regulatory role in ZIKV budding. To our knowledge, phosphorylation of a flavivirus protein has not been implicated in the budding process.

Moreover, considering the conservation of Y61 in other pathogenic flaviviruses (Table 1, Supplementary Information 4), this PTM could be fundamentally important for flavivirus budding.

Phosphosite Y61 is found in domain II of ZIKV E, the domain responsible for ZIKV E dimerization (17). The inability of the phosphomimetic Y61E infectious clone to efficiently grow over the first two passages following reverse genetics rescue coupled with its conversion at this residue from a glutamic acid to valine by passage 3 (Fig 3C, 3D) suggest that the negatively charged amino acid is not favorable. Indeed, there may be pressure at position 61 to be either a non-charged or hydrophobic amino acid. A zoomed-in view of the ZIKV model reveals that Y61 is near an asparagine (N207) and glutamic acid (E262) (Fig S6B). In the absence of phosphorylation, it is possible that the N207 and E262 residues are interacting. A large phosphate group on Y61 might repel the negative charge on E262 and perhaps disrupt this interaction, providing a possible explanation for why a non-charged state is optimal at this position. This preference is further supported by the fact that we readily rescued the non-phosphorylatable mutant, Y61F, which has a hydrophobic, non-charged phenylalanine at position 61.

The hydrophobic/non-phosphorylatable amino acid changes (Y to F or V) at position 61 resulted in ZIKV infectious clones that were less cytopathic (Fig 3I) and outperformed wild type ZIKV in reaching peak titers (Fig 3D, 3G), yet resulted in a budding defect (Fig 3F, 3H). We interpret these finding in several ways. First, phosphorylation of ZIKV E at Y61 is either not favorable to the virus or is beneficial at very specific time point(s) in the ZIKV life cycle, supported by the observation of the budding defect. If the latter scenario is at work, our growth curve data may not be sufficient to resolve at which other life cycle stages Y61 phosphorylation may be beneficial to ZIKV because the infectious clones employed contained static mutations at this position. The negatively-charged phosphosite may be important in aiding a specific purpose, such as encouraging a conformational change for a period and then returning to an unphosphorylated state. When mutated to Y61E, ZIKV E no longer expressed from a plasmid (Fig S8), suggesting that the ZIKV envelope protein containing a negatively-charged glutamic acid at this position is too unstable for expression or may be quickly marked for degradation.

Interestingly, the Y61 site is highly conserved among flaviviruses. A multiple sequence alignment (Supplementary Information 4) shows that flaviviruses such as WNV, YFV, Japanese encephalitis virus (JEV), and Saint Louis encephalitis virus (SLEV) also contain a tyrosine at this position, suggesting that maintaining control over the charge of this tyrosine likely benefits these flaviviruses. DENV contains an isoleucine at this position and more distantly related flaviviruses contain a leucine, both of which share properties with tyrosine and phenylalanine in that they are also non-charged, hydrophobic amino acids.

The seemingly paradoxical characteristics of decreased budding (Fig 3F, 3H) but enhanced growth of the Y61E mutants (Fig 3D) are resolved when considering the significantly dampened cytopathogenicity of Y61 mutants (Fig 3I). With increased cell viability, mutants may replicate to higher levels, evidenced by the observed increased intracellular titers of Y61 mutants compared to WT (Fig 3G), overcoming their budding defects, and ultimately outperforming the WT virus (summarized in Fig 5).

**Figure 5.**
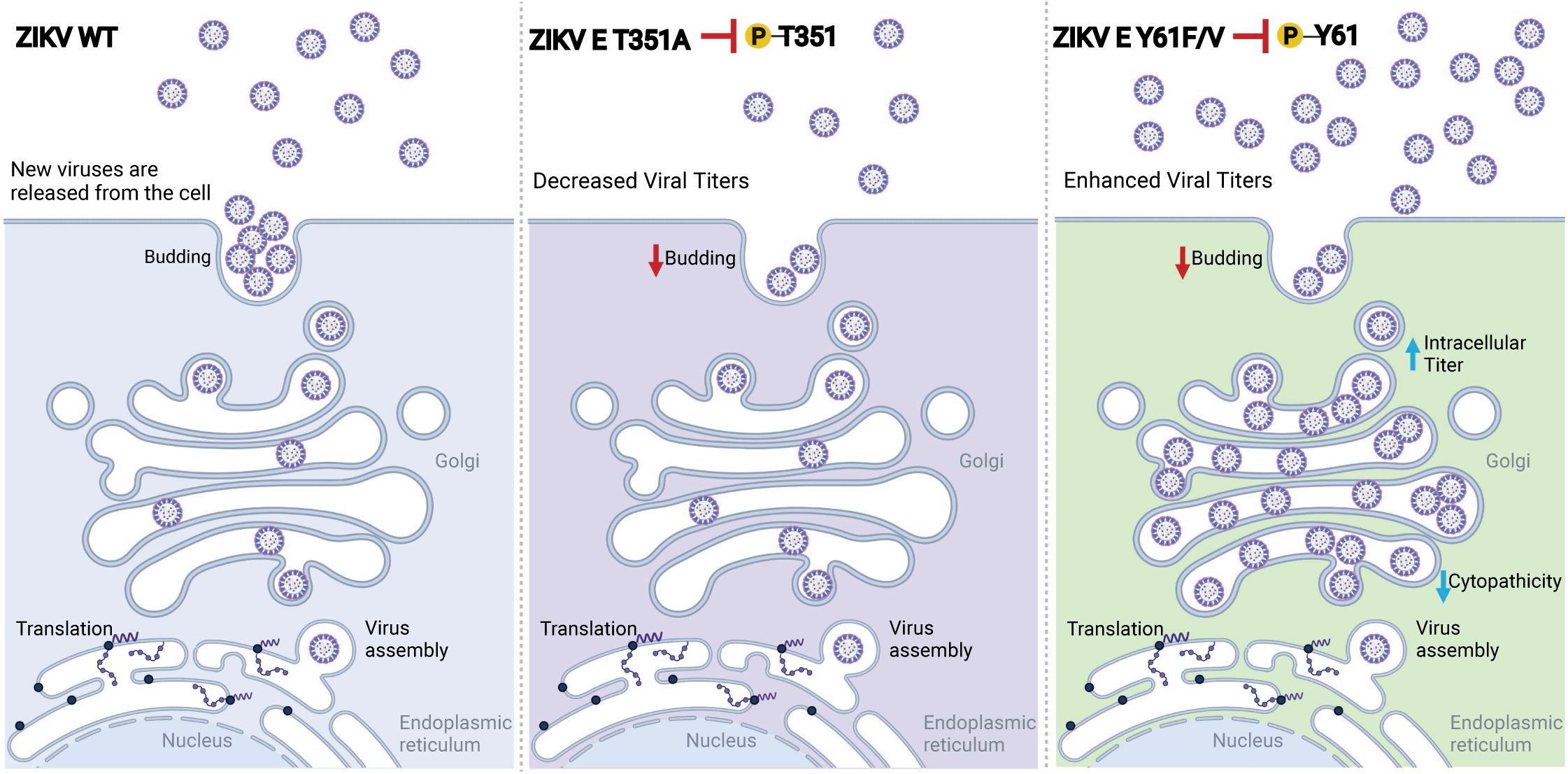
Graphical summary of the effects of blocking phosphorylation of ZIKV E T351 and ZIKV E Y61 on ZIKV growth. A graphical interpretation of results from Fig 3D – I and Fig S9 experiments. Blocking phosphorylation of ZIKV E T351 reduces budding efficiency, resulting in a viral titer decrease. Blocking phosphorylation of ZIKV E Y61 reduces viral budding efficiency, increases intracellular viral titers, and reduces ZIKV cytopathogenicity. Presumably, the decrease in virus-induced cytopathogenicity supersedes the negative impact on virus budding, resulting in an overall higher capacity for viral propagation. Created with BioRender.com.

Phosphorylation of ZIKV E at position T351 was observed both in purified virions and in cells. T351 sits within Domain III of ZIKV E, which contains motifs for cellular receptor binding and is often the target of neutralizing antibodies (17). A negative charge at this position, either from a phosphorylated threonine or an aspartic acid substitution, was optimal for growth (Fig 3E). Similarly, a negatively charged aspartic acid in the budding assay resulted in the maintenance of virus-like particle budding. Conversely, mutation to an alanine, the nonphosphorylatable mutant, resulted in a strong reduction in virus-like particle budding (Fig 3F), suggesting that host-driven phosphorylation of T351 promotes ZIKV budding. Intriguingly, phosphorylation of T351 was discovered both on ZIKV E isolated from human cells as well on virions derived from mosquito cells, suggesting that not only is phosphorylation of T351 important in the host, but that it may also play a role in the virus vector. Future studies could focus on confirming the functional importance of T351 and other discovered phosphosites, such as Y61, in mosquito cells, as well as evaluating whether phosphomutant viruses from mosquito cells impact entry or infectivity in human cells. Because there may be differences in human and arthropod kinomes, it is possible that the ZIKV phosphoproteome in vector cells would diverge from our dataset in mammalian cells. Additionally, it may be that optimal viral fitness requires different patterns of regulatory phosphorylation in the cells of mammalian and arthropod hosts.

Therefore, it would be informational to map the phosphoproteome of ZIKV in mosquito cells for comparison with this dataset as well as for comparison of the roles of phosphorylation in both cell types.

Previous studies have shown that host-driven phosphorylation of the flavivirus NS5 protein is critical for the flavivirus viral life cycle. For example, phosphorylation of YFV NS5 at a highly conserved site (S56) is required for YFV to evade the cap-dependent innate immune response in the host cell (10). Blocking the phosphorylation of another conserved flavivirus residue, T449 on DENV NS5, results in a nonfunctional protein (31). Other key studies linked phosphorylation of NS5 or Hepatitis C virus NS5A to important functions such as its interaction with NS3 (11, 32), localization during different stages of infection (32), and viral particle assembly and viral replication (12, 33). In some instances, phosphorylation of NS5 was detected but not the specific residues driving the observed phenotype (11, 32). The highly conserved NS5 phosphosites identified in our study may be responsible for driving these important functions (Table 1). Indeed, we have uncovered several highly conserved phosphosites in the ZIKV proteome that may help guide the discovery of new functions for these viral proteins or the mechanisms by which these functions are regulated.

Our mass spectrometry analysis revealed 115 host kinases that interacted with one or more of the 8 ZIKV proteins probed (Table S2). A limitation of this study is that we did not orthogonally verify these interactions. An additional consideration is that kinase expression in the cell lines used may not fully reflect that of primary cells targeted by ZIKV, which could influence both the phosphoproteomic landscape and the observed kinase and viral protein interactions. Therefore, it will be important to validate interactions and functionally interrogate these kinases in future studies. However, despite these potential limitations, 32 of the kinases discovered in this study were identified in other ZIKV proteomics or functional studies, providing independent validation of our findings. We used MiST analysis to refine the list of kinases from 115 to 40 and remove potential false positives (Table S3). This resulted in the exclusion of 18 independently discovered kinases (23–28), introducing the possibility that the analysis might remove functionally relevant targets. Therefore, in addition to the more stringent MiST-selected kinases listed in Table S3 and Fig S10, we have also provided the complete list of 115 interacting kinases in Table S2.

The discovery of host kinases as viral protein interactors raised the possibility that some subset may be responsible for viral protein phosphorylation. Treating cells after infection with Bosutinib, an FDA-approved tyrosine kinase inhibitor, reduced ZIKV growth (Fig 4B, 4D).

Indeed, a recent study by Valencia *et al.* confirmed our finding that Bosutinib reduces ZIKV growth in the setting of BHK-21 cells (34). There are different possibilities for how Bosutinib inhibits ZIKV growth. One is that Bosutinib’s inhibitory effect on ZIKV is exclusively through cellular mechanisms. For example, Bosutinib may be acting on host kinases that ultimately lead to a cellular environment not conducive to optimal ZIKV growth. Another possibility is that Bosutinib reduces titers by inhibiting kinases that act directly on ZIKV phosphosites and cellular processes do not play a role in titer reduction. However, Bosutinib is known to act on many cellular kinases (35), making it unlikely that the reduction of ZIKV titers is solely due to its impact on viral phosphosites. Which leads to the last possibility that Bosutinib may be acting through a combination of cellular mechanisms as well as directly on ZIKV phosphosites. To investigate whether Bosutinib’s mechanism of action is associated with blocking phosphorylation of Y61, the lone high confidence phosphotyrosine discovered on ZIKV E, we treated the Y61V and Y61F mutant clones with Bosutinib. If Bosutinib’s only mechanism for reducing ZIKV titers is by blocking Y61 phosphorylation due to the inhibition of a kinase that acts on this site, and the mutant clones are refractory to this kinase because they are non-phosphorylatable, then we would expect that Bosutinib treatment would not lower mutant titers. Conversely, if Bosutinib reduces ZIKV titers through mechanism(s) completely independent to blocking Y61 phosphorylation, then mutations at this site would be irrelevant and we would expect Bosutinib to reduce mutant titers as effectively as it does ZIKV WT titers. Our results showed that Bosutinib reduced the mutant clones’ titers, but to a lesser degree, than the WT virus (Fig 4E), suggesting that Bosutinib’s titer reduction is, in part, due to blocking phosphorylation at Y61 and also through other cellular mechanisms and/or potentially a different ZIKV phosphosite (i.e. NS1 Y22, Y175, Y200, or one not detected in this study). It is possible that Bosutinib treatment, by blocking Y61 phosphorylation, results in a budding defect, similar to the budding defects of VLPs (Fig 3F) and authentic virions (Fig 3H) of non-phosphorylatable Y61 mutant clones. In the mutant clones, this budding defect was ultimately overcome by higher intracellular (Fig 3G) and extracellular (Fig 3D and 3G) titers stemming from a substantial increase in cell viability (Fig 3I).

Although the kinase predicted by Scansite 4.0 to phosphorylate Y61, FGR, was not detected as a ZIKV interacting partner, the closely related tyrosine protein kinase FYN was (Table 1, Table S2). Furthermore, FYN is inhibited by Bosutinib at nanomolar concentrations (35). Additional studies are needed to identify the exact host kinase that phosphorylates Y61. As discussed above, Bosutinib may be affecting other ZIKV phosphosites and we have found additional kinases that are targeted by the drug. It is also important to consider that the concentration at which kinase inhibitors are administered will influence their specificity. So, although targets of Bosutinib (Fig 4A) were not predicted to directly phosphorylate discovered phosphosites (Table 1), Bosutinib may be inhibiting off-target kinases that phosphorylate ZIKV proteins. Further studies using molecules related to Bosutinib may improve efficacy and specificity for blocking specific kinases. Collectively, these data imply that blocking phosphorylation by host kinases is a potential antiviral strategy.

Many of our discovered kinases have been corroborated as ZIKV interactors or functionally important for ZIKV (Fig 4A, Tables 1, S2, S3) (23–28). Of these, a several (ATM, AURKB, CAMK2B, GSK3B, NEK7, PRKDC) are predicted to phosphorylate ZIKV at the phosphosites discovered in this study, making them intriguing targets for further studies. Several interacting kinases (ATM, ATR, and mTOR) discovered in our study were the targets of early drug screens against ZIKV. Specifically, Cherry *et al*. used inhibitors VE-822 and WAY-600 to robustly block ZIKV infection (36). Intriguingly, VE-822 targets kinases ATM and ATR, which we found to be interacting partners of ZIKV M (ATM, ATR) or ZIKV E (ATM) (note that ATM is predicted to phosphorylate S16 on ZIKV M; Table 1). WAY-600 targets mTOR, which we found to be an interacting partner of ZIKV phosphoproteins M and E as well as NS4A (Table S2). Interestingly, mTOR was the most abundant kinase found to interact with any of the ZIKV proteins (Table S2). Its importance was established early in the ZIKV epidemic when it was discovered that ZIKV proteins can suppress the Akt-mTOR pathway, leading to impairment of neurogenesis and upregulation of autophagy, which is beneficial for ZIKV during replication (37, 38). Another study found an overall downregulation of the AKT/mTOR pathway during ZIKV infection (25). Due to ZIKV M associating with high levels of mTOR (see peptide counts in Table S2), we theorize that ZIKV M could be acting as a sink for mTOR, providing an explanation for why the Akt-mTOR pathway is downregulated and there is less phosphorylation of mTOR targets observed during ZIKV infection (25).

In summary, we have detailed a high-resolution map of host-driven PTMs on ZIKV proteins, uncovered a novel mechanism by which ZIKV budding may be regulated, discovered that phosphorylation may influence ZIKV cytopathogenicity, and identified an FDA-approved drug that could be repurposed for the treatment of ZIKV infection. The dataset described here will be useful to the field for the discovery of novel ZIKV protein functions and the mechanisms by which these functions are regulated. Our work further suggests that targeting host kinases may be an effective antiviral strategy and provides potential targets.

## Materials and Methods

### Cells and viruses

HEK 293T/17 cells (CRL-11268), referred to as HEK293T throughout, and A549 cells (CCL-185) were procured from the American Type Culture Collection (ATCC, Manassas, VA). Vero E6 cells were kindly given by J. L. Whitton (The Scripps Research Institute, La Jolla, CA).

HEK293T cells were maintained in Dulbecco’s modified Eagle medium (DMEM) containing 10% fetal bovine serum (FBS), 1% penicillin-streptomycin, 1% HEPES buffer solution, 1% MEM nonessential amino acid solution, and 1% GlutaMAX, from Thermo Fisher Scientific (Waltham, MA). A549 cells were maintained in DMEM-F12 (Thermo Fisher) containing 10% FBS and 1% penicillin-streptomycin. Vero E6 cells were maintained in DMEM containing 10% FBS, 1% penicillin-streptomycin, and 1% HEPES buffer solution. Cell lines described above were incubated at 37°C with 5% CO_2_. *Aedes albopictus* mosquito cells (C6/36; a kind gift of Prof Alain Kohl, MRC-University of Glasgow Centre for Virus Research) were maintained in Leibovitz’s L15 media (Gibco) containing 10% FBS (Gibco), 10% triptose phosphate broth (Gibco), and 1% pen/strep (Gibco) and incubated at 28°C. ZIKV strain BeH819015, a Brazilian isolate (GenBank: KU365778.1), was generously provided by the Baric Lab (University of North Carolina, Chapel Hill, North Carolina, USA) (20). The Zika virus strain used to infect mosquito cells was ZIKV/H.sapiens/Brazil/PE243/2015 (Recife, Brazil) (39). Working stocks of infectious ZIKV, including recombinant ZIKV clones generated via reverse genetics (see below for details on recovery), were grown on Vero E6 cells and titered by standard plaque assay.

### Plasmids

Several plasmids were used for expression of ZIKV proteins. The nucleotide sequences of ZIKV ORFs used in this study were derived from ZIKV strain H/PF/2013 (NCBI gene identifier number AHZ13508). The Heise lab (University of North Carolina, Chapel Hill, North Carolina, USA) provided their pOME plasmid-based library that permits expression of each ZIKV ORF containing a C-terminal 3XFLAG tag (40). To enable streptavidin-based affinity purification of ZIKV proteins, the ZIKV C, E, M, NS1, NS2B, and NS4A ORFs encoded in the pOME plasmids were subcloned into our modified pCAGGS expression vector (41, 42) such that they would encode a C-terminal streptavidin binding peptide (SBP) (MDEKTTGWRGGHVVEGLAGELEQLRARLEHHPQGQREP) in place of the 3XFLAG tag.

Each ZIKV ORF was fused to the SBP tag through an 18 base pair linker. The nucleotide sequence of the ZIKV genes was amplified by PCR from the pOME vectors with forward primers that contained a 5’ overhang containing a Gateway AttB1 site and a Kozak sequence and reverse primers containing an overhang with the linker sequence (all primers used are listed in Supplementary Information 5). The SBP tag was amplified from a previously generated in-house plasmid via PCR using the forward primer SBP (5’-GCAGCTGGAGGTGGAGGTATGGACGAAAAAACCACCGGT-3’), which has a 5’ overhang containing the linker sequence, and the reverse primer SBP (5’-ACCACTTTGTACAAGAAAGCTGGGTCTTACGGTTCACGCTGACCCTGCGG-3’), which contains a 3’ overhang with a stop codon preceding an AttB2 sequence. The two PCR products (each respective amplified ZIKV gene and the amplified SBP tag) were fused by PCR using the appropriate ZIKV gene forward primer and the SBP reverse primer. The resulting cassette was subcloned into the modified pCAGGS vector using the Gateway cloning system (Invitrogen) following the manufacturer’s instructions. pOME plasmids encoding ZIKV NS2A, NS3, NS4B, or NS5, each with the same C-terminal linker and SBP tag as described directly above, were generated by GenScript Biotech Corporation (Piscataway, NJ, USA) but were maintained in the original pOME vector. Plasmids for the ZIKV reverse genetics system were kindly provide by the Baric Lab as described in Widman *et al.* (20). The plasmid from the Baric system which contained the ZIKV E gene was modified by GenScript to include the Y61F, Y61E, T351A, and T351D mutations. The plasmid used for the virus-like-particle budding assay, which encodes the ZIKV prM and E genes, is described here (21) and was generously provided by Emergent BioSolutions, Inc. This plasmid was also modified by GenScript to include the include the Y61F, Y61E, T351A, and T351D mutations. All plasmid sequences were verified by DNA sequencing.

### Identification of post-translational modifications of ZIKV proteins by mass spectrometry

To identify phosphorylation sites on ZIKV proteins via mass spectrometry, three general approaches were taken. In the first approach, to purify ZIKV E, Vero E6 cells were infected with ZIKV at an MOI of 0.1 or mock infected and 72 hours later cells were collected and lysed in Triton buffer consisting of 1% Triton X-100, 0.5% NP40, 140mM NaCl, and 25mM Tris-HCl containing a protease inhibitor cocktail (04693159001, Roche Applied Science, Indianapolis, IN) as well as PhosStop phosphatase inhibitor cocktail (04906837001, Roche Applied Science). The resulting protein lysate was incubated with anti-ZIKV E 4G2 antibody at 4°C, overnight. The next day, magnetic Protein G Dynabeads (ThermoFisher Scientific, Waltham, MA) were added to the lysate-antibody mixture and incubated at 4°C, rotating for at least 4 hours (but not more than 8 hr). Beads were then magnetically isolated, gently washed 3 times in Triton buffer, magnetically isolated for a final time and then boiled in Laemmli buffer (62.5 mM Tris-HCl, 10% glycerol, 2% sodium dodecyl sulfate and 0.01% bromophenol blue (B392, Fisher Scientific, Pittsburgh, PA) for 5 minutes to release antibody, ZIKV E protein, and associated host or viral protein partners. Similarly, ZIKV virions were immunopurified from infected Vero E6 supernatants 96 hours post-infection using Dynabeads coated with anti-ZIKV E 4G2 antibody or an irrelevant mouse IgG then lysed using 10X Triton buffer containing the same protease and phosphatase inhibitor cocktails described directly above. The mock infected lysates and supernatants served as controls during mass spectrometry analysis.

To purify ZIKV proteins expressed from plasmid, our second approach was to transfect HEK293T cells with modified pCAGGS plasmid expressing ZIKV C, M, E, NS1, NS2B, NS4A, NS4B, or NS5 proteins with a C-terminal SBP tag in each case, or an empty plasmid labeled as V for “vector” in Figures 1E, S2, S3, S4 to serve as the control condition during mass spectrometry analysis. 5×10^5^ cells per well in 6-well plates were transfected with 2 μg of plasmid using 8 μg of polyethylenimine (23966, Polysciences, Inc., Warrington, PA), which was reconstituted at 1 mg/mL. Two days later, transfected cells were treated either with DMSO (D2650, Sigma-Aldrich. Saint Louis, MO), H_2_O_2_, (Thermo Fisher) or Calyculin A (Cell Signaling Technology, Danver, MA) for 20 minutes. Cells were collected and lysed with Triton buffer (same formulation described above) and then subjected to affinity purification using magnetic MyOne Streptavidin T1 beads (65601, Thermo Fisher Scientific) according to the manufacturer’s instructions. Briefly, magnetic beads were washed in buffer several times, added to protein lysates from transfected cells, and then rotated overnight at 4°C. The next day, the beads were magnetically isolated and washed several times with lysis buffer containing protease and phosphatase inhibitors and then boiled in Laemmli buffer for 5 minutes to release SBP-tagged ZIKV proteins from the streptavidin beads.

Infected or mock-infected cell lysates (control condition), transfected lysates from cells expressing ZIKV proteins or empty plasmid/empty vector (control condition), and virion protein lysates from infected or mock infected supernatants (control condition) were separated on 4-20% Tris-Glycine polyacrylamide gels (EC60255, Invitrogen). To visualize protein bands for excising and mass spectrometry processing, gels were stained with Coomassie (40% methanol, 20% acetic acid, and 0.1% Brilliant Blue R (B7920, Sigma-Aldrich)). Gels were destained with a solution of 30% methanol and 10% acetic acid and imaged using a Canon Canoscan 8800F scanner. Sample lanes were cut into a series of gel pieces containing prominent protein bands or band groups. After excision, gel pieces were further cut into 1 mm cubes and processed.

Chemicals used for processing were purchased from Thermo Fisher Scientific. Gel pieces were rinsed with HPLC-grade water and then incubated with destain solution (50 mM ammonium bicarbonate and 50% acetonitrile) for 30 minutes at 37°C. Destain was removed and gel pieces were dehydrated by incubating twice with 100% acetonitrile for 5 minutes. The gel pieces were reduced with 25 mM dithiothreitol in 50 mM ammonium bicarbonate for 30 minutes at 55°C. After cooling, gel pieces were dehydrated with 100% acetonitrile for 5 minutes and then alkylated with 10 mM iodoacetamide in 50 mM ammonium biocarbonate for 45 minutes at room temperature, while protected from light. Gel pieces were washed twice in destain solution for 5 minutes, dehydrated with 100% acetonitrile, then rehydrated with water for 10 minutes. Gel pieces were further dehydrated with two 5-minute incubations in 100% acetonitrile. All liquid was removed and gel pieces were left to incubate at room temperature to evaporate trace acetonitrile. Gel pieces were rehydrated with a solution of 12.5 ng/μL sequencing-grade, modified trypsin (V5111, Promega) in 50 mM ammonium bicarbonate on ice for 30 minutes, before digesting overnight at 37°C. Peptides were extracted with a solution of 2.5% formic acid in 50% acetonitrile while spinning in a microcentrifuge at 13,000 RPM for 10 minutes. The supernatant was removed and saved while the gel pieces were subjected to further extraction and rinsing with 100% acetonitrile. The second extraction was combined with the initial extraction. All solvent was removed from the extracts using a vacuum centrifuge at 37°C. The peptides were resuspended in 2.5% formic acid, 2.5% acetonitrile prior to mass spectrometry analysis.

Dried peptides were resuspended in Solvent A (2.5% MeCN, 0.15% formic acid (FA)) and separated using the Easy n-LC 1200 across 15-cm columns packed in-house with 2.7 µm C18 packing material prior to analysis on the Q Exactive Plus mass spectrometer fitted with a Nanospray Flex ion source (2.2 kV) and supplied with Thermo Xcalibur 4.0 software. Peptides were eluted using a 0 - 50% gradient of Solvent B (80% MeCN, 0.15% FA) over 60 min, followed by 10 minutes at 95% Solvent B. The precursor scan (scan range = 360–1700 *m/z*, resolution = 7.0 x 10^4^, maximum IT = 100 ms) was followed by HCD fragmentation spectra in a Top-10 approach (resolution = 3.5 x 10^4^, AGC = 5.0 x 10^4^, maximum IT = 50 ms, isolation window = 61.6 *m/z*, normalized collision energy = 26%, dynamic exclusion = 30 s). Raw spectra were searched against a forward and reverse database of tagged ZIKV proteins with common contaminants added using SEQUEST with a precursor mass tolerance of ± 5 PPM and a fragment ion tolerance of ± 0.006 Da. The following differential modifications were permitted: phosphorylation of serine, threonine and tyrosine (±79.9663 Da), ubiquitylation of lysine (±114.0429 Da), oxidation of methionine (±15.9949 Da), carboxyamidomethylation of cysteine (±57.0215 Da), and acrylamidation of cysteine (±71.0371 Da). No enzyme was indicated in the original search to expand the database, and the resulting hits were filtered for tryptic peptides.

Spectra of phosphopeptide hits were manually assessed and compared to their unphosphorylated counterparts to confirm hits and determine phosphorylation site localization. Any phosphopeptide hits with poor fragmentation or with ambiguous phosphorylation site localization were removed from the dataset. To identify bound human interactors, raw data were searched against a forward and reverse human database containing common contaminants via SEQUEST. Trypsin was specified as the cleavage agent with two missed cleavages permitted.

Mass tolerances and allowed modifications were as stated above. Resulting hits were filtered by cross correlation score (>= 2.0 for z=2, >= 2.2 for z=3, >=2.4 for z=4, >=2.6 for z=5) and delta correlation score (>=0.15), resulting in a FDR of <1%. Host proteins were considered ZIKV interactors if they were identified by two or more unique peptides in experimental conditions and were not present in the controls. Proteins were also considered ZIKV interactors in cases where proteins were >5-fold enriched in experimental over control conditions. We used the MiST computational tool at https://modbase.compbio.ucsf.edu/mist in the PCA (principal component analysis) training mode to score the viral protein – host protein interactions identified by mass spectrometry. After scoring, a threshold of 0.62 was chosen to generate Table S3 and Fig S10.

Fig 4A included kinases with scores greater than 0.8.

In our third approach, which led to the identification of ZIKV E T351 from virions, C6/36 cells were infected with ZIKV at an MOI of 0.1 and then incubated at 28°C. Four days later, the supernatant was harvested and clarified by centrifugation at 5000 RPM. The clarified supernatant was concentrated using 100,000 MW cut-off Centricon concentrators (Millipore). The concentrate was underlayed with a 5 mL sucrose cushion (24% w/v in 1x NTE buffer; NaCl-Tris-EDTA) and ultracentrifuged at 105,000 *g* in a Surespin 630 Rotor (Thermo Scientific) for 1.5 hours. The pellet-associated virus was resuspended in NTE, then overlaid onto a 10-35% (w/v) / density gradient of potassium tartrate in 30% glycerol in NTE, prepared using a Gradient Master (Biocomp Instruments). Virions were banded by ultracentrifugation at 175,000 *g* overnight and collected by pipetting from the top of the gradient. Next, the virus-containing fraction was removed from the gradient, placed in 100,000 MW cut-off dialysis units, and dialyzed several times using 1× NTE to remove the potassium tartrate and glycerol. After dialyzing overnight, virus was concentrated using 100,000 MW cut-off Centricon concentrators (Millipore).

Concentrated ZIKV virions were inactivated at 70°C in 4M final urea in TEAB and then prepared for liquid chromatography and tandem mass spectrometry (LC-MS/MS) as described (43). Briefly, the sample was reduced, alkylated, and then digested with trypsin and LysC using filter-assisted sample preparation (FASP) (44). Peptides were separated by reversed-phase chromatography using a 2h gradient on an Ultimate 3000 RSLCnano HPLC system (Dionex). This was run in direct injection mode and coupled to a Q Exactive mass spectrometer (Thermo), running in ‘Top 10′ data-dependent acquisition mode with fragmentation by higher-energy collisional dissociation (HCD). Charge state + 1 ions were rejected and fragmentation and dynamic exclusion with 40 s was enabled. Mass spectra were annotated in MaxQuant version 1.6.3.4 (45) with reference proteomes from Zika virus isolate ZIKV/H.sapiens/Brazil/PE243/2015 (GenBank KX197192.1) and *Aedes aegypti* (UniProt UP000008820, accessed on 2018-10-26) and the MaxQuant common contaminants list. Standard settings were used, with the enzyme set as trypsin/P and fixed modifications set as carbamidomethyl (C). Variable modifications were oxidation (M), acetyl (Protein N-ter), phospho (STY) and, to account for artefactual modifications when inactivating in urea, carbamyl (KRC). Modified spectra were considered where they could be identified in the absence of artefactual carbamylation. Data from this third approach can be found at https://researchdata.gla.ac.uk/1222/.

### Multiple sequence alignment

A multiple sequence alignment (MSA) of flavivirus FASTA-formatted sequences was generated using the MUSCLE (Multiple Sequence Comparison by Log-Expectation) algorithm (46) as implemented in the NIAID Virus Pathogen Database and Analysis Resource (ViPR) through the web site at http://www.viprbrc.org/ (47). The Uclust parameter was set as a MUSCLE pre-processor to improve both speed and quality of alignment. The alignment visualization with PTM annotations was created in Jalview 2.11.1.4 (48). The following flaviviruses were aligned: ZIKV H/PF/2013 (AHZ13508), ZIKV Uganda (ABI54475), ZIKV SPH2015 (ALU33341), ZIKV Paraiba (ANH10698), ZIKV USA UT (AOO19564), ZIKV BeH819015 (AMA12085), DENV 1 NP059433), DENV 2 (NP056776), DENV 3 (YP001621843), DENV 4 (NP073286), JEV (Japanese encephalitis virus; NP059434), SLEV (Saint Louis Encephalitis Virus; YP001008348), USUV (Usutu virus; AAS59401), WNV 1(YP001527877), WNV 2 (NP041724), AHFV (Alkhurma hemorrhagic fever virus; AAL08421), KFDV (Kyasanur forest disease virus, AAQ91607), OHFV (Omsk hemorrhagic fever virus, AAQ91606), TBEV (tick-borne encephalitis virus; NP043135), ENTV (Entebbe bat virus; AAV34153), MMLV (Montana myotis leukoencephalitis virus, CAC82713), LAMV (Lammi virus, ACR56717), YFV (NP041726), and PCV (Palm Creek virus, AGG76014).

### SDS-PAGE and western blotting

Protein lysates were diluted in Laemmli buffer, boiled for 5 minutes, then separated on NuPAGE 4–12% Bis-Tris gels with MES buffer. Protein was transferred to nitrocellulose membranes using iBlot gel transfer stacks (IB301001 or IB301002, Invitrogen) and the Invitrogen iBlot Device according to manufacturer instructions. Membranes were blocked with 5% milk in PBS for 1 hour and incubated overnight with primary antibody diluted in PBS which contained 5% milk and 0.2% Tween 20 (BP337, Fisher Scientific). The following day the membrane was rinsed several times with TBST (tris-buffered saline with 0.1% Tween 20) then incubated for 1 hour with secondary antibody diluted in PBS containing 5% milk, 0.2% Tween 20, and 0.02% SDS. After secondary antibody incubation, the membrane was washed several times in TBST then washed in PBS before imaging on a LI-COR Odyssey CLx system.

Primary antibodies were used for western blotting at the following concentrations: mouse anti-ZIKV E (4G2) (1:10,000) (generously provided by Baric Lab), mouse anti-streptavidin binding peptide (MAB10764, Millipore, Billerica, MA) (1:10,000), rabbit anti-actin (A2066, Sigma-Aldrich) (1:5,000), and mouse anti-actin (A5441, Sigma-Aldrich) (1:5,000). For quantitative western blotting, the following secondary antibodies from LI-COR were used at a 1:20,000 dilution: IRDye 800CW goat anti-mouse (926–32210) and IRDye 680LT goat anti-rabbit (926–68021).

### Generation of mutant ZIKV clones

We used the Baric lab reverse genetics system to generate phosphomimetic and non-phosphorylatable mutants as described in Widman *et al.* (20). ZIKV E was mutated at Y61 to Y61E and Y61F and separately ZIKV E T351 was mutated to T351D and T351A. The Baric reverse genetics system is composed of 4 plasmids, Fragments A through D, that encode the ZIKV proteome. Fragment A was modified by GenScript from thymine to guanine at nucleotide position 1599 and thymine to adenine at position 1601 to generate the Y61E phosphomimetic while nucleotide 1600 was modified from adenine to thymine for the Y61F non-phosphorylatable mutant. Nucleotides 2470 and 2471 in Fragment A were modified from adenine and cytosine to guanine and adenine, respectively, for the T351D phosphomimetic while position 2470 was modified from adenine to guanine for the T351A non-phosphorylatable mutant. Infectious recombinant ZIKV clones were generated as described in Widman *et al.* (20). Four plasmids containing Fragments A-D, respectively, which encode for the ZIKV genome, were digested with BamHI and Bsu36I for Fragment A, Bsu36I and BstXI for Fragment B, BstXI and SfiI for Fragment C, and SfiI, SmaI, AlwNI for Fragment D. Restriction enzymes were all acquired from New England Biolabs (Ipswich, MA). The four digested fragments were ligated with T4 DNA ligase purchased from New England Biolabs. Ligated DNA was precipitated with chloroform then *in vitro* transcribed using the mMESSAGE mMACHINE™ T7 ULTRA Transcription Kit from ThermoFisher Scientific. The transcribed product was then electroporated into Vero E6 cells using a Gene Pulser Xcell (Bio-Rad Laboratories, Hercules, CA) and incubated at 37°C for 4 days. Cell media was collected after 4 days and passaged onto a new monolayer of Vero E6 cells. Cell media collected from this passage was described as “passage 1/ p.1”. Clarified cell media was titered by plaque assay using Vero E6 cells.

### Virus growth curves

For the Y61F and Y61E mutants, six-well plates were seeded with 2 x 10^5^ Vero E6 cells per well and infected the following day at an MOI of 0.001. For the T351A and T351D mutants, 12-well plates were seeded with 2 x 10^4^ A549 cells and infected the following day at an MOI of 0.01.

Cell media was collected at indicated time points from separate wells. Cell media was clarified by centrifugation at 1200 RPM for 5 minutes, stored at-80°C, and then titered by plaque assay (see plaque and focus assays below for details).

### Plaque and focus assays

Plaque assays were used to measure infectious virus titers in several experiments as follows. Six-well plates were seeded with 2.5 x 10^5^ Vero E6 cells per well and the following day inoculated with 10-fold serial dilutions of virus. Plates were rocked every 15 minutes for 1 hour to allow for viral absorption at 37°C, the cells were overlaid with a solution of 0.7% agarose (20–102, Apex Industrial Chemicals, Aberdeen, United Kingdom) in Vero E6 growth media. Wells with agarose plugs were fixed 3 days later with a solution of 2.5% formaldehyde (1635-4L, Sigma-Aldrich) in 3x PBS. Following removal of the agarose, the fixed cell monolayers were stained with 0.1% crystal violet (C581-100, Fisher Scientific) and 2.1% ethanol in water.

Focus assays were used to assess the presence of the Y61E infectious clones after several viral passages. 24-well plates were seeded with 2 x 10^4^ Vero E6 cells per well and the following day were inoculated with 10-fold serial dilutions of infectious viral clones. Plates were incubated for 1 hour at 37°C with rocking every 15 minutes. After absorption, cells were overlaid with 0.01% methylcellulose in Vero E6 media (see Cells and Viruses for media detail) and were left in a 37°C incubator for 3 days. After 3 days the overlay was removed and cell monolayers were fixed with 4% formaldehyde (28906, Thermo Fisher Scientific) at room temperature, washed, then permeabalized with 0.5% Triton X-100 in PBS. Cell monolayers were then blocked with 5% milk in PBS and thereafter incubated with 4G2 antibody (1:10,000) diluted in 5% milk in PBS for 1 hour at 37°C. Cells were washed with PBS and then incubated with goat anti-mouse secondary antibody at a 1:2000 (5450-0011, LGC Clinical Diagnostics, Inc., Milford, MA) dilution in 5% milk in PBS for 1 hour at 37°C. Cells were washed again and then incubated with TrueBlue Peroxidase Substrate (5510-0030, LGC Clinical Diagnostics, Inc.) until the formation of blue foci and reaction was stopped by removing the substrate.

### Virus-like particle (VLP) release assay

To determine the efficiency of VLP formation and release when ZIKV E contains mutations at Y61 or T351, 5 x 10^5^ HEK293T cells in 6-well plates were transfected with 2 µg of the modified or unmodified (corresponding to either WT ZIKV E or mutant ZIKV E) Emergent BioSolutions, Inc. plasmids (see Plasmids for details) using 8 µg of PEI per well. Twenty-four hours after transfection, fresh cell media was added to wells. 48 hours after transfection, cells and cell media were collected. Cell media was clarified by centrifugation for 3 minutes at 1500 RPM and then VLPs were lysed with 10X Triton buffer for a final dilution of 1X lysis buffer. Cells were collected with cold PBS, centrifuged for 3 minutes at 1500 RPM, and then lysed with Triton buffer. Cells and cell media were subjected to quantitative western blotting for the detection of ZIKV E by way of 4G2 antibody (concentration of 1:10,000). To calculate efficiency of VLP release, each ZIKV E protein value found in cells was first normalized to actin values in cells (rabbit anti-actin antibody at 1:5000). VLP release efficiency was then calculated as the quotient of the E protein quantity in VLPs divided by the quantity of E in cells [(E_mutant_VLP / E_mutant_cells) / (E_WT_ VLP/ E_WT_ cells)*100].

### Intracellular Infectious Virus Titer Assay and Budding Efficiency Calculation

To determine if there were differences in the intracellular titers of Y61 non-phosphorylatable mutant viruses compared to WT, deionized water was used to release intracellular virions from infected cells. To establish if there were differences in the budding efficiencies of mutants compared to WT virus, intracellular titers were compared to their corresponding extracellular titers and then normalized to WT virus. The following methodology was adapted from Kong *et al.* (49). First, 1.4 x 10^6^ Vero E6 cells were infected with ZIKV WT, Y61V, or Y61F virus at an MOI of 0.001. After 96 hours, supernatants were clarified by centrifugation at 1500 RPM for 3 minutes to remove floating cells. Infected cells were centrifuged at 1500 RPM for 3 minutes then washed with PBS, 3 times in tandem, to remove any extracellular virions present in the supernatant. Washed cells were lysed with nuclease free deionized water for 15 minutes at room temperature. Then 2X DMEM was added to the lysate to normalize salts. Next, the lysate was centrifuged to separate cell debris from the intracellular virions in the supernatant. Finally, supernatants and lysates from infected cells were assayed for extracellular and intracellular titers, respectively, by plaque assay. Budding efficiency was calculated using the following formula: [(Mutant _extracellular titer_ / Mutant_intracellular titer_) / (WT virus_extracellular titer_ / WTvirus_intracellular titer_)*100].

### Cell viability assay

To determine if Y61 mutant viruses impact cell viability, we used an MTT (3-(4,5-dimethylthiazol-2-yl)-2,5-diphenyltetrazolium bromide)) assay (M6494, Thermo Fisher Scientific). First, 1.2 x 10^5^ Vero E6 cells were infected with ZIKV WT, Y61V, or Y61F virus. At 24, 48, and 96 hours post infection, supernatants were collected, clarified by centrifugation at 1500 RPM for 3 minutes, stored at-80°C, then titered by plaque assay. At every collection timepoint, cells were treated with 1.2 mM MTT in PBS for 4 hours at 37°C. After incubation, MTT was removed and DMSO was added to each well, incubated for 10 minutes at 37°C, and absorbance was read at 540 nm. Cell viability was determined by normalizing the absorbance readings of mutant-infected wells to the WT-infected control wells.

### Kinase inhibition assays

To determine if treating cells at the time of infection with various kinase inhibitors would impact viral growth, viral challenge assays were performed. First, to screen a panel of inhibitors at a fixed concentration, 2.5 x 10^5^ A549 cells were seeded in 6-well plates. The next day cells were infected at an MOI of 0.1 with ZIKV. After an hour of absorption at 37°C, the inoculum was removed and cells were treated with 10 µM of Sunitinib (B1045, APExBIO Technology LLC, Houston, TX) Bosutinib (PZ0192,Sigma-Aldrich), Defactinib (B4800, APExBIO), Imatinib (9084S, Cell Signaling Technology), or DMSO vehicle control in HEK293T media. The infection proceeded for 48 hours then supernatants were collected, clarified by centrifugation, stored at-80°C, and then titered by plaque assay. The percent inhibition was determined by normalizing titers of drug-treated wells to the DMSO control well titers and multiplying by 100. Second, to determine the IC_50_ of Bosutinib or Defactinib, a dose response curve was set up with drugs at concentrations of 30, 20, 15, 10, 7.5, 5, and 1.25 µM as well as DMSO. 2.8 x 10^4^ A549 cells were seeded into 48 well plates. One day later, cells were infected at an MOI of 0.1. One hour after absorption, the inoculum was removed and cells were treated with A549 media containing drug at the listed concentrations. After 48 hours, supernatants were collected, clarified by centrifugation, stored at-80°C, and then titered by plaque assay. To determine the cell toxicity of each drug, an MTT assay was conducted in parallel to the infection and kinase inhibitor treatment according to manufacturer instructions. A549 cells were set up in 48 well plates and treated with the listed concentrations of Bosutinib. After 48 hours, cell media was removed and replaced with 1.2 mM MTT in PBS for 4 hours at 37°C. After incubation, MTT was removed and DMSO was added to each well, incubated for 10 minutes at 37°C, and absorbance was read at 540 nm. Cell viability was determined by first normalizing the absorbance readings of drug-treated cells to the DMSO control wells. Cell toxicity was determined by subtracting the value of cell viability from 100. To determine the IC_50_ (half-maximal inhibitory concentration) of Bosutinib and Defactinib, we used [inhibitor] versus normalized response curve fitting in Prism 8.0. To determine CC_50_ (cytotoxic concentration causing death to 50% of cells), the same Prism 8.0 curve fitting was applied to our MTT assay data (inhibitor versus cell toxicity). The selectivity index (SI) of each drug was calculated by dividing the CC_50_ by the IC_50_ of each drug.

To determine if Bosutinib impacts growth of Y61 mutants, viral challenge assays were performed. First, 3 x 10^5^ A549 cells were infected with ZIKV WT, Y61V, or Y61F virus at an MOI of 0.01. After an hour of absorption at 37°C, the inoculum was removed and replaced with A549 media containing 15 µM Bosutinib or DMSO. After 48 hours, supernatants were collected, clarified by centrifugation, stored at-80°C, then titered by plaque assay.

## Acknowledgments

We thank the NIH for the following grant support: T32 HL076122 (I.M. and C.M.Z.), R21AI135265 (J.W.B.), P20RR021905 and P30GM118228 (Immunobiology and Infectious Disease COBRE awards) (J.W.B.), AI107731 (R.S.B.), and the National Institute of General Medical Sciences of the NIH (P20GM103449; Vermont Biomedical Research Network Bioinformatics – H.D.; Vermont Biomedical Research Network Proteomics – C.G.). We thank the United Kingdom Research and Innovation Medical Research Council for the following support: ZK/16-012 (D.B) and MR/N008618/1 (E.H.). We also thank the University of Vermont Office of Fellowships, Opportunities and Undergraduate Research for their support of D.T. Funding agencies had no role in study design, collection and analysis of data, manuscript preparation or the decision to publish. We are grateful to Emergent BioSolutions, Inc. and Dr.

Lindsay Whitton for kindly providing critical reagents described in the Materials and Methods, Svenja Hester (Advanced Proteomics Facility, University of Oxford) for performing and advising on mass spectrometry, Glasgow Polyomics for computing resources and Dr. Emily Bruce, the UVM Immunobiology group, and the Vermont Lung Center for technical comments and insightful discussions.

---

**Figure S1.**
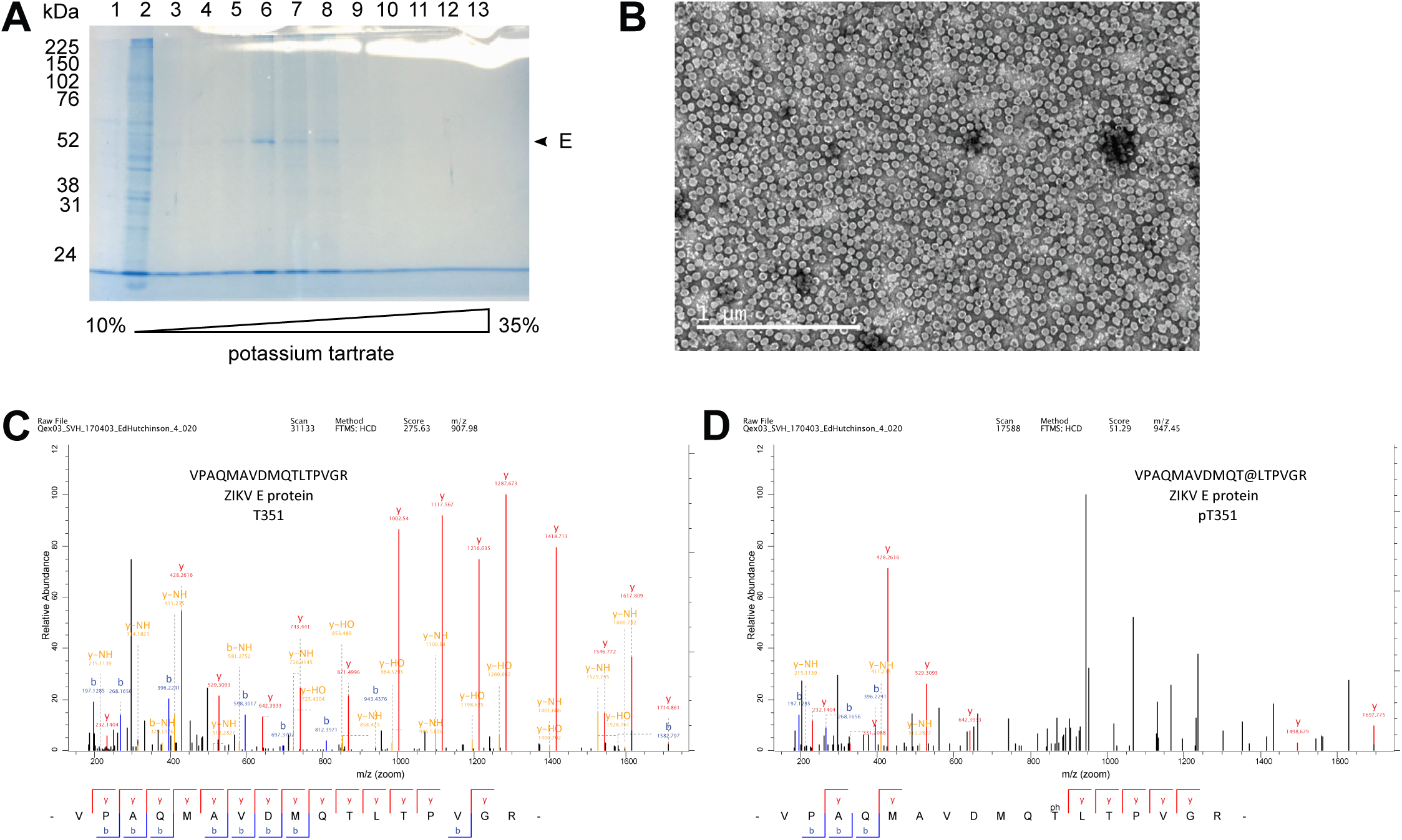
Identification of ZIKV E phosphosite T351 in virions from mosquito cells. C6/36 mosquito cells were inoculated with ZIKV at an MOI of 0.1. Four days later, cell-free ZIKV particles were collected, concentrated, and purified by banding on a potassium tartrate density gradient. (A) Aliquots of gradient fractions were separated by SDS-PAGE and stained with Coomassie Brilliant Blue; the position of ZIKV E is shown. Virus-containing fractions then were dialyzed and concentrated. (B) Negative stain transmission electron micrograph showing concentrated ZIKV virions prepared using this process (from an infection at MOI of 1.0, harvested at 72 h post-infection). Concentrated ZIKV particles were digested with trypsin and LysC and analyzed using mass spectrometry. Annotated mass spectra are shown for ZIKV E T351 in (C) unmodified site and (D) phosphorylated forms; ph/@ = phosphorylation.

**Figure S2.**
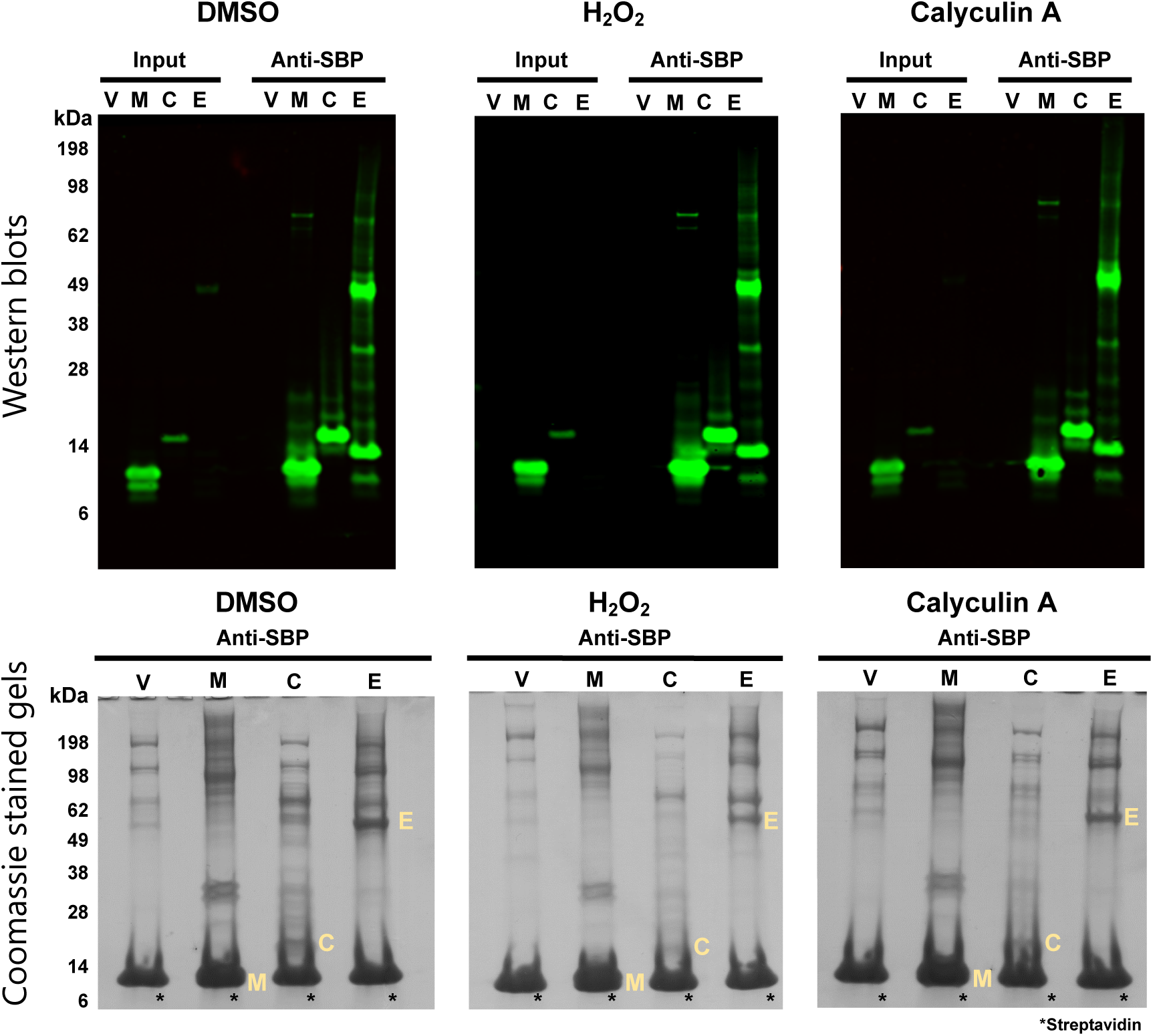
Purification of plasmid-expressed ZIKV M, C, or E in the setting of phosphatase inhibition for mass spectrometry analysis. Western blots and Coomassie stained SDS-page gels of purified SBP-tagged ZIKV proteins. HEK293T cells were transfected with a plasmid encoding the indicated SBP-tagged ZIKV protein or, as a control, an empty vector (V). Two days after transfection, cells were treated with DMSO, H_2_O_2_, or Calyculin A for 30 minutes and then lysed in Triton-X buffer. SBP-tagged ZIKV proteins were affinity purified using streptavidin-coated magnetic beads and the captured protein mixtures were orthogonally separated using SDS-PAGE, followed by in-gel processing for subsequent mass spectrometry analysis to discover post-translational modifications or interacting host proteins. The molecular weight in kDa, viral proteins (yellow text and arrows), and immunoglobulin bands are labeled. Note that the asterisks in indicate monomeric streptavidin that was eluted off of the streptavidin beads when boiled.

**Figure S3.**
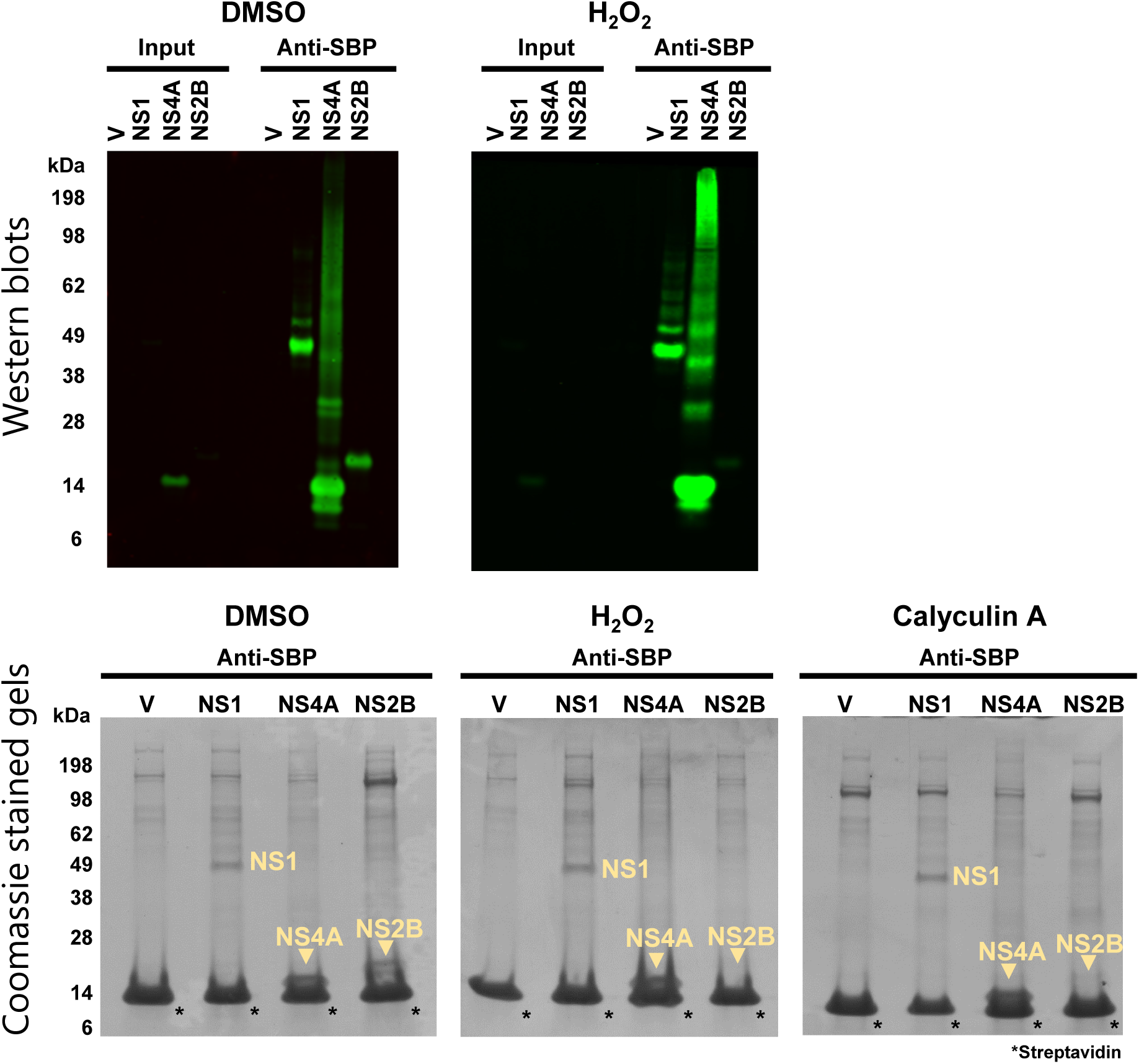
Purification of plasmid-expressed ZIKV NS1, NS4A, and NS2B in the setting of phosphatase inhibition for mass spectrometry analysis. Western blots and Coomassie stained SDS-page gels of purified SBP-tagged ZIKV proteins. HEK293T cells were transfected with a plasmid encoding the indicated SBP-tagged ZIKV protein or, as a control, an empty vector (V). Two days after transfection, cells were treated with DMSO, H_2_O_2_, or Calyculin A for 30 minutes and then lysed in Triton-X buffer. SBP-tagged ZIKV proteins were affinity purified using streptavidin-coated magnetic beads and the captured protein mixtures were orthogonally separated using SDS-PAGE, followed by in-gel processing for subsequent mass spectrometry analysis to discover post-translational modifications or interacting host proteins. The molecular weight in kDa, viral proteins (yellow text and arrows), and immunoglobulin bands are labeled. Note that the asterisks in indicate monomeric streptavidin that was eluted off streptavidin beads when boiled.

**Figure S4.**
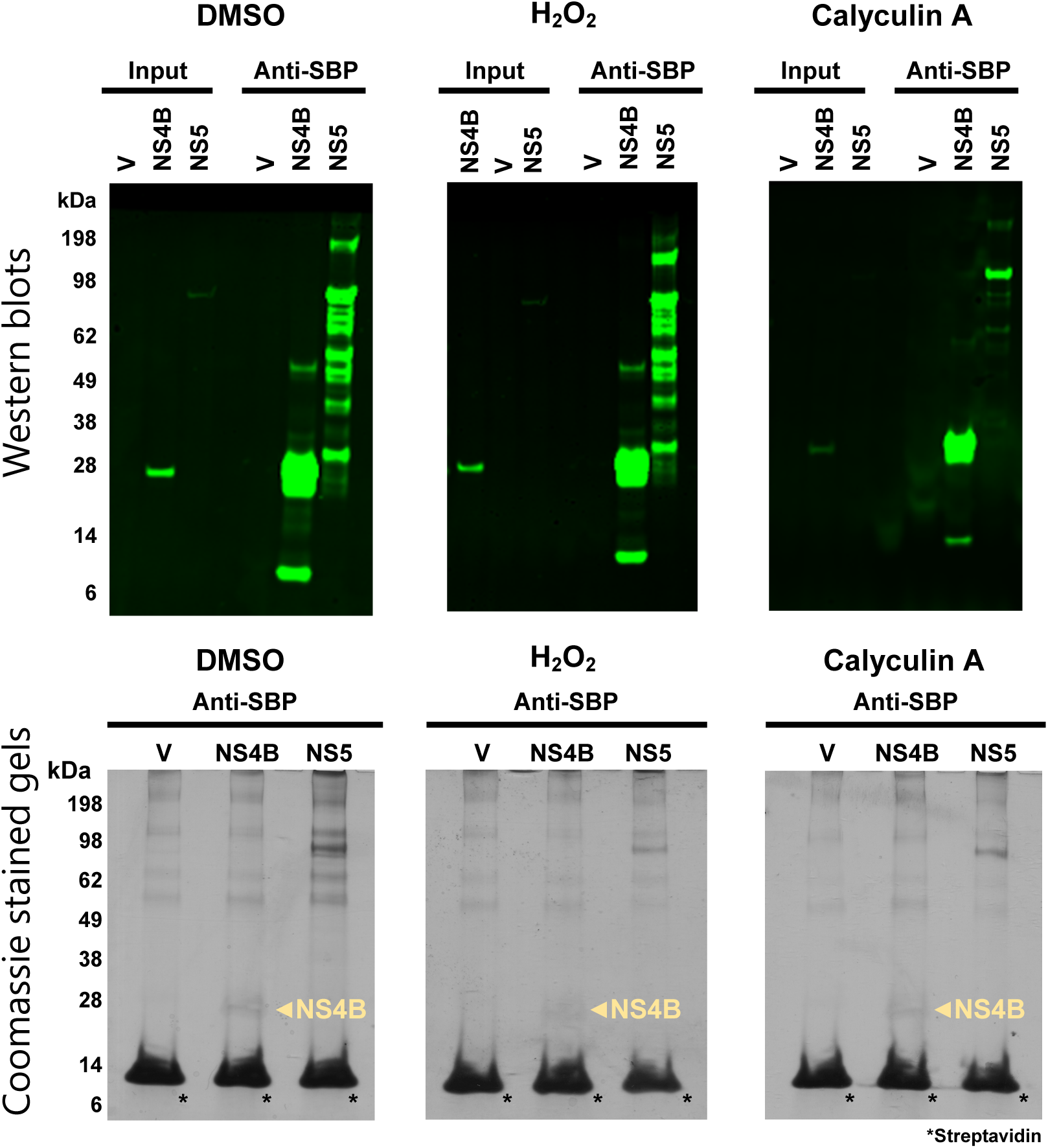
Purification of plasmid expressed ZIKV NS4B and NS5 in the setting of phosphatase inhibition for mass spectrometry analysis. Western blots and Coomassie stained SDS-page gels of purified SBP-tagged ZIKV proteins. HEK293T cells were transfected with a plasmid encoding the indicated SBP-tagged ZIKV protein or, as a control, an empty vector (V). Two days after transfection, cells were treated with DMSO, H_2_O_2_, or Calyculin A for 30 minutes and then lysed in Triton-X buffer. SBP-tagged ZIKV proteins were affinity purified using streptavidin-coated magnetic beads and the captured protein mixtures were orthogonally separated using SDS-PAGE, followed by in-gel processing for subsequent mass spectrometry analysis to discover post-translational modifications or interacting host proteins. The molecular weight in kDa, viral proteins (yellow text and arrows), and immunoglobulin bands are labeled. Note that the asterisks in indicate monomeric streptavidin that was eluted off of the streptavidin beads when boiled.

**Figure S5.**
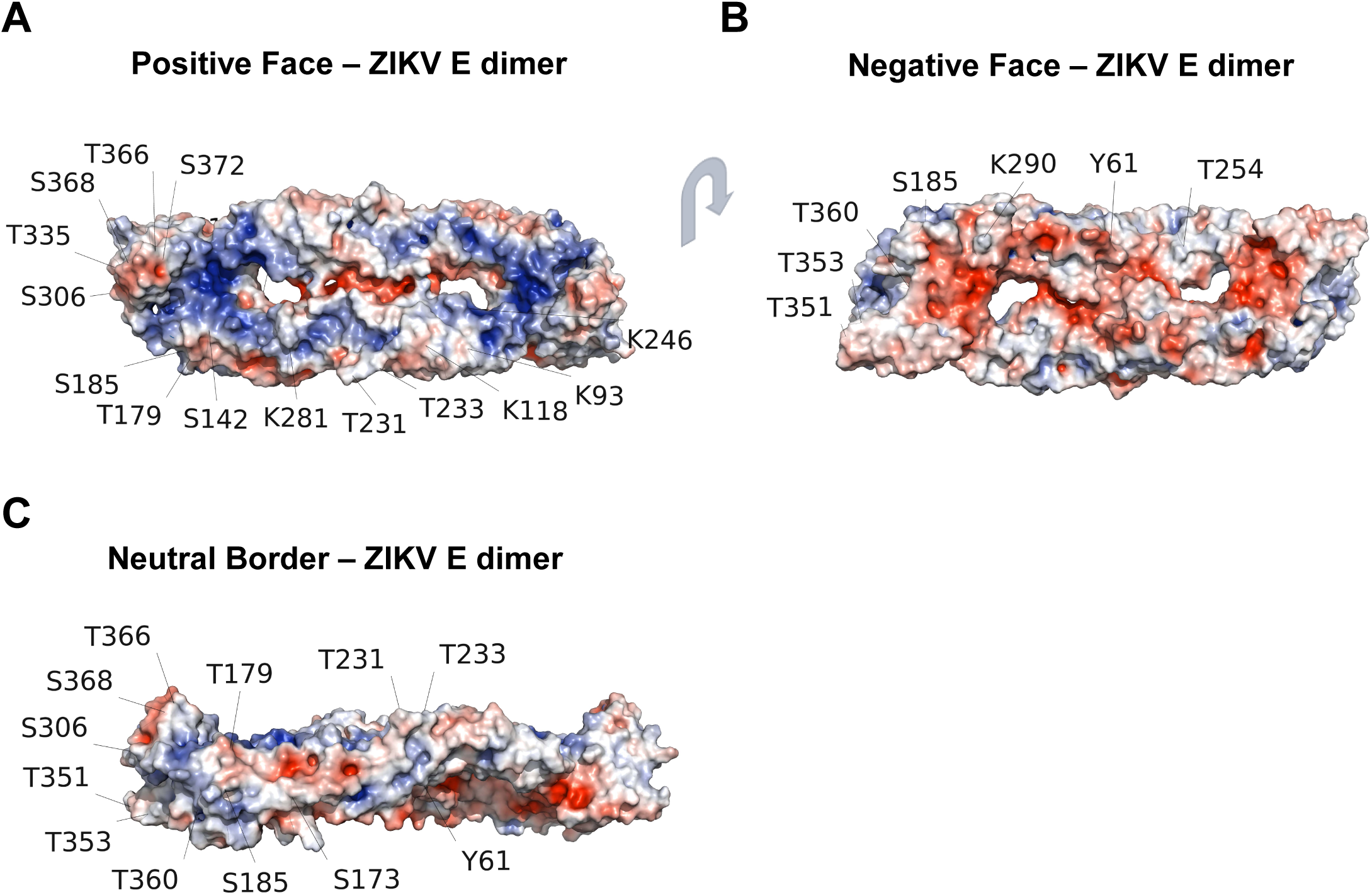
PTMs labeled on an electrostatic model of ZIKV E dimer. (PDB ID: 5JHM). Blue indicates positive charges while red indicates negative charges. (A) Electrostatic view of ZIKV E dimer. (B) rotated to the negatively charged “face”. (C) “Side-view” positioning showing the neutral border of electrostatically labeled ZIKV E dimer along with location of post-translational modification.

**Figure S6.**
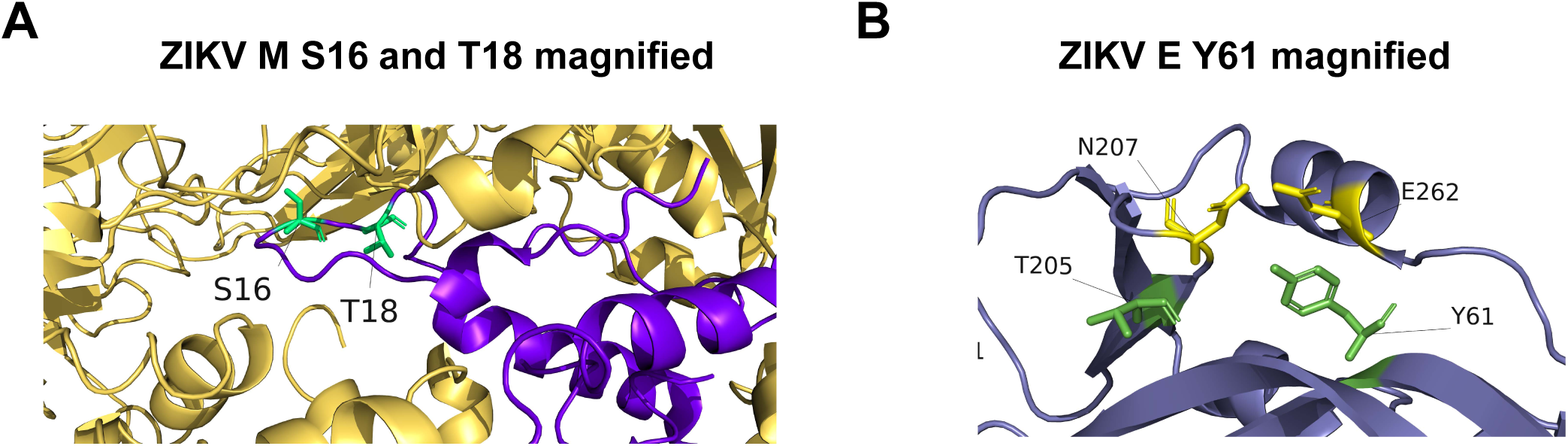
ZIKV E Y61 interactions with neighboring residues and a view of the junction of ZIKV E and. **M.** (A) Zoomed in view of the junction of ZIKV E (yellow) and M heterodimer (dark purple) with M phosphosites labeled in green. (B) Zoomed in view of Y61 on ZIKV E. Y61 is labeled in green while residues in close proximity, N207 and E262, are labeled in yellow.

**Figure S7.**
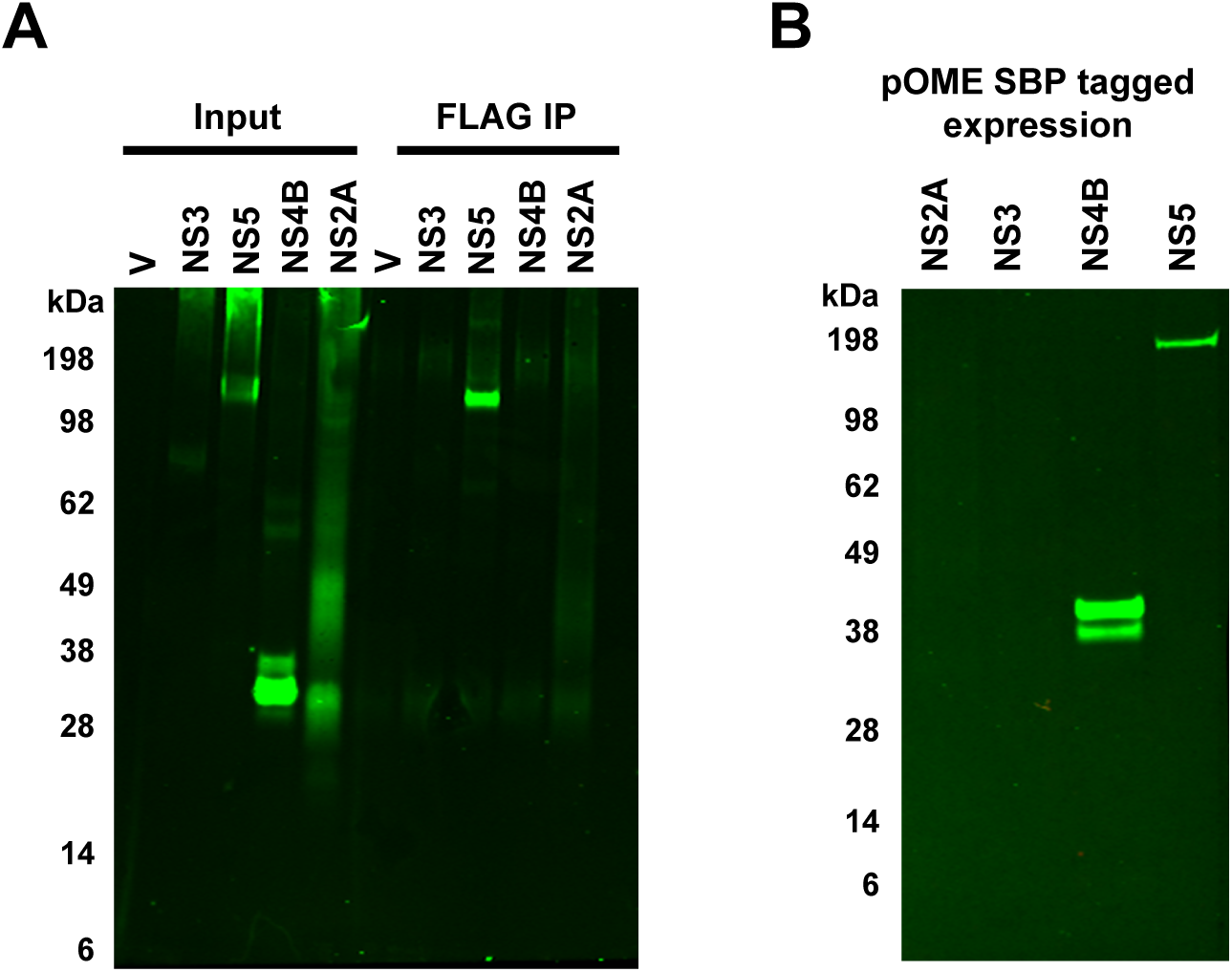
NS2A and NS3 did not express at sufficiently high levels for mass spectrometry analysis. Attempts were made to express and purify NS2A and NS3 via plasmid transfection of mammalian cells. HEK293T cells were transfected with plasmids that encoded ZIKV proteins with either a C-terminal 3x FLAG-tag (A) or a SBP tag (B). Two days later, cells were lysed and tag-based protein purifications were attempted with an anti-FLAG antibody and magnetic protein G beads (A) or magnetic streptavidin beads (B).

**Figure S8.**
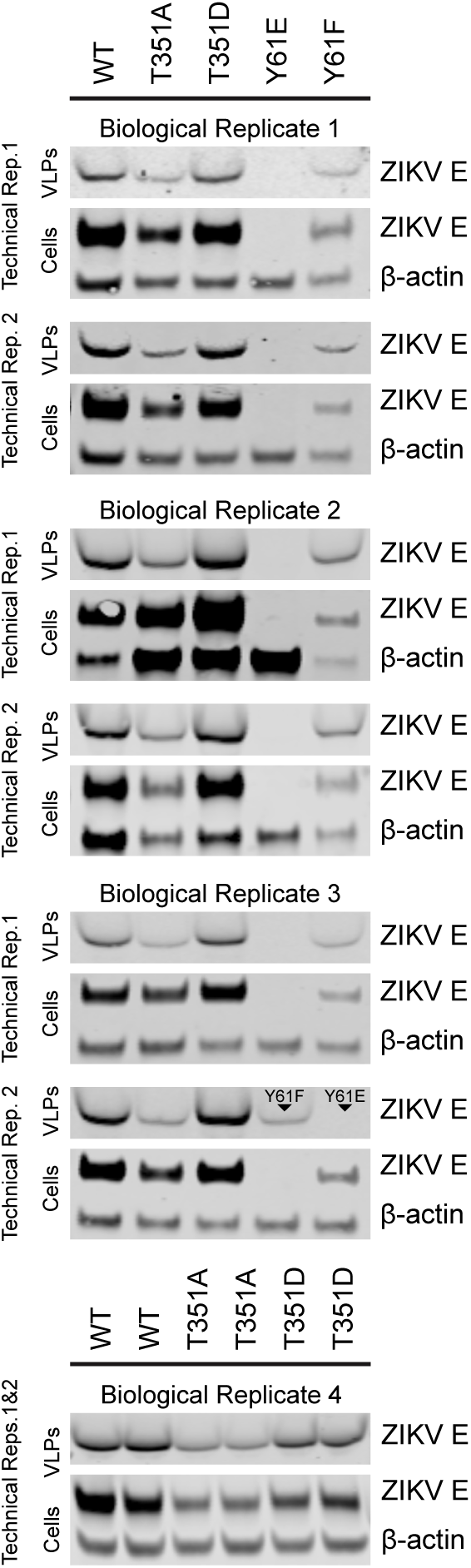
Expression of WT and mutant forms of plasmid-derived ZIKV. **E.** Accompanying Western blots of the VLP budding assay shown in Fig 3F. HEK293T were transfected with a plasmid that encodes ZIKV prM and E (either WT E or the indicated E mutants). Forty-eight hours after transfection, cells and cell media (labeled “VLPs”) were subjected to quantitative western blotting for the detection of ZIKV E by way of 4G2 antibody or β-Actin.

**Figure S9.**
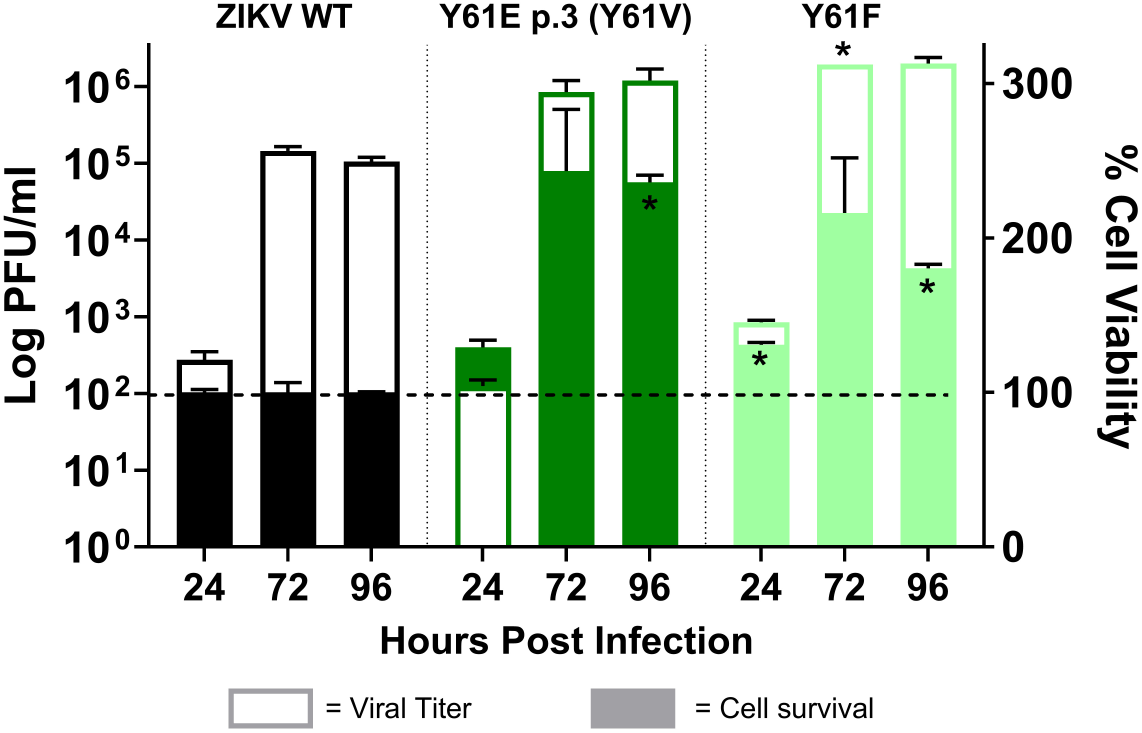
Elevated titers of non-phosphorylatable Y61 mutant viruses correspond with higher cell viability. Left Y-axis: Vero E6 cells were infected with WT, Y61V, or Y61F virus at an MOI of 0.001. Supernatants were collected at 24, 72, and 96 hours post infection and viral titers (open bars) were determined by plaque assay. Right Y-axis: simultaneous to titer determination, cell viability (shaded bars) was measured by MTT assays at 24, 72, and 96 hours post infection. Data represent the mean±SEM from two biological replicates. Statistical significance was determined in Prism 8.0 software via two-way ANOVA with Holm-Sidak’s test for multiple comparisons. *, P<0.05.

**Figure S10.**
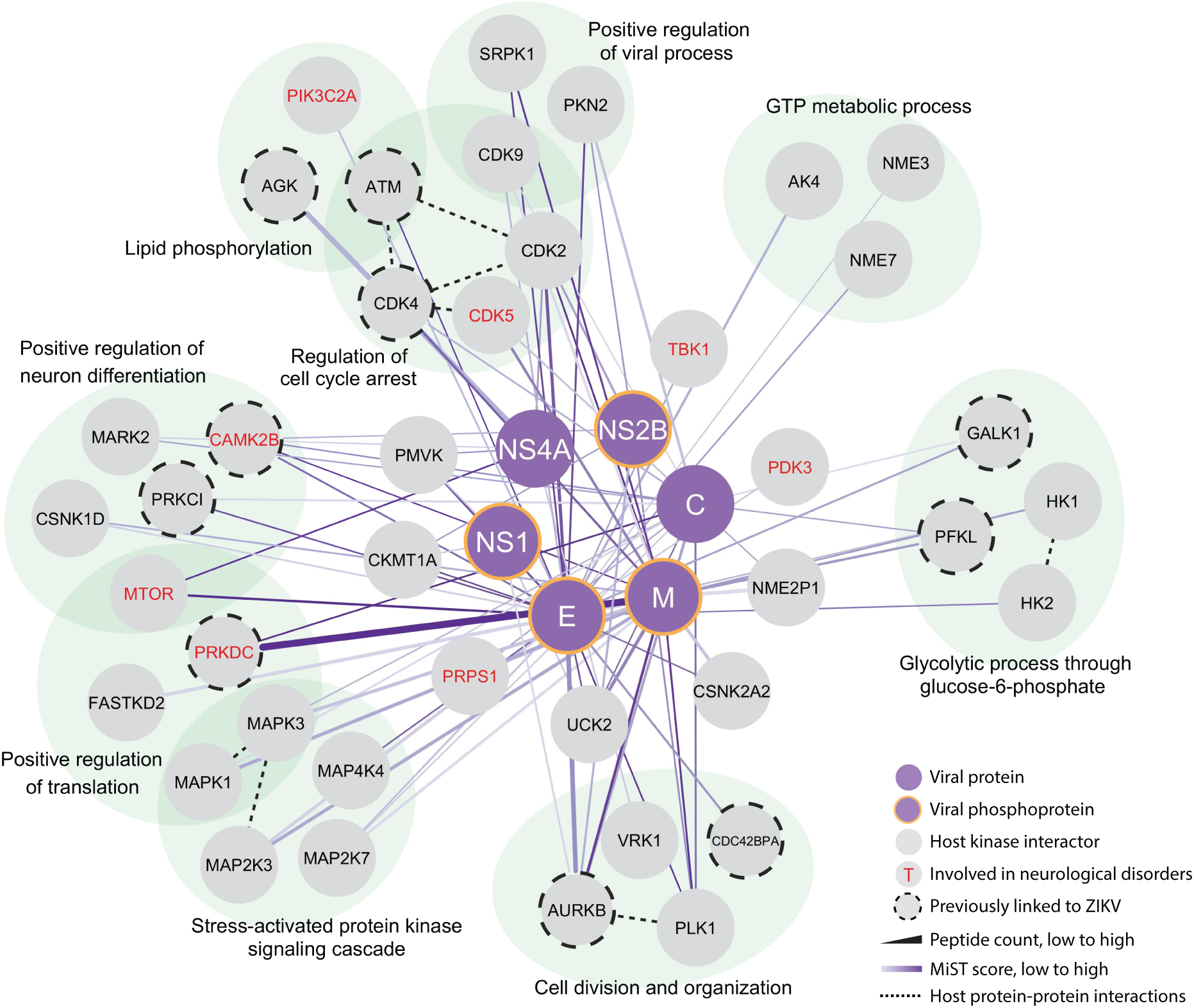
ZIKV-human kinase interaction network. The complete list of 115 interacting kinases found in this study (Table S2) was subjected to MiST scoring with a score threshold of 0.62, resulting in 40 kinases and 85 viral-host interactions. The network depicts interactions between ZIKV proteins (purple circles / with orange border if ZIKV protein was phosphorylated) and cellular kinases (grey circles) discovered during mass spectrometry. Kinases involved in neurological disorders such as microcephaly, intellectual disability, developmental delays, vision and hearing issues, paralysis, and dementia (grey circle with red text), and kinases that were independently found in other ZIKV interactome or functional studies (grey circle with dashed borders) are indicated. Interactions between viral and host proteins are depicted by solid lines with color representing MiST scoring (lowest to highest corresponding to lightest to darkest purple) and width representing peptide counts (lowest to highest corresponding to thinnest to thickest). Dashed lines indicate published interactions or associations between kinases integrated with the STRING application in Cytoscape 3.9.1.

## References

1. Smithburn KC. Neutralizing antibodies against certain recently isolated viruses in the sera of human beings residing in East Africa. J Immunol. 1952;69(2):223–34. Epub 1952/08/01. PubMed PMID: 14946416.

2. Zika cases and congenital syndrome associated with Zika virus reported by countries and territories in the Americas, 2015-2018 cumulative cases. Pan Amerian Health Organization (PAHO), 2018.

3. Moore SM, Oidtman RJ, Soda KJ, Siraj AS, Reiner RC, Jr., Barker CM, Perkins TA. Leveraging multiple data types to estimate the size of the Zika epidemic in the Americas. PLoS Negl Trop Dis. 2020;14(9):e0008640. Epub 2020/09/29. doi: 10.1371/journal.pntd.0008640. PubMed PMID: 32986701; PMCID: PMC7544039.

4. Cao-Lormeau VM, Blake A, Mons S, Lastere S, Roche C, Vanhomwegen J, Dub T, Baudouin L, Teissier A, Larre P, Vial AL, Decam C, Choumet V, Halstead SK, Willison HJ, Musset L, Manuguerra JC, Despres P, Fournier E, Mallet HP, Musso D, Fontanet A, Neil J, Ghawche F. Guillain-Barre Syndrome outbreak associated with Zika virus infection in French Polynesia: a case-control study. Lancet. 2016;387(10027):1531–9. Epub 2016/03/08. doi: 10.1016/S0140-6736(16)00562-6. PubMed PMID: 26948433; PMCID: PMC5444521.

5. Mlakar J, Korva M, Tul N, Popovic M, Poljsak-Prijatelj M, Mraz J, Kolenc M, Resman Rus K, Vesnaver Vipotnik T, Fabjan Vodusek V, Vizjak A, Pizem J, Petrovec M, Avsic Zupanc T. Zika Virus Associated with Microcephaly. N Engl J Med. 2016;374(10):951–8. Epub 2016/02/11. doi: 10.1056/NEJMoa1600651. PubMed PMID: 26862926.

6. Hemonnot B, Cartier C, Gay B, Rebuffat S, Bardy M, Devaux C, Boyer V, Briant L. The host cell MAP kinase ERK-2 regulates viral assembly and release by phosphorylating the p6gag protein of HIV-1. J Biol Chem. 2004;279(31):32426–34. Epub 2004/05/25. doi: 10.1074/jbc.M313137200. PubMed PMID: 15155723.

7. Ziegler CM, Eisenhauer P, Bruce EA, Weir ME, King BR, Klaus JP, Krementsov DN, Shirley DJ, Ballif BA, Botten J. The Lymphocytic Choriomeningitis Virus Matrix Protein PPXY Late Domain Drives the Production of Defective Interfering Particles. PLoS Pathog. 2016;12(3):e1005501. doi: 10.1371/journal.ppat.1005501. PubMed PMID: 27010636; PMCID: PMC4806877.

8. Ziegler CM, Eisenhauer P, Bruce EA, Beganovic V, King BR, Weir ME, Ballif BA, Botten J. A novel phosphoserine motif in the LCMV matrix protein Z regulates the release of infectious virus and defective interfering particles. J Gen Virol. 2016. doi: 10.1099/jgv.0.000550. PubMed PMID: 27421645.

9. Ziegler CM, Eisenhauer P, Manuelyan I, Weir ME, Bruce EA, Ballif BA, Botten J. Host-Driven Phosphorylation Appears to Regulate the Budding Activity of the Lassa Virus Matrix Protein. Pathogens. 2018;7(4). doi: 10.3390/pathogens7040097. PubMed PMID: 30544850; PMCID: PMC6313517.

10. Bhattacharya D, Hoover S, Falk SP, Weisblum B, Vestling M, Striker R. Phosphorylation of yellow fever virus NS5 alters methyltransferase activity. Virology. 2008;380(2):276–84. doi: 10.1016/j.virol.2008.07.013. PubMed PMID: 18757072; PMCID: PMC2583469.

11. Kapoor M, Zhang L, Ramachandra M, Kusukawa J, Ebner KE, Padmanabhan R. Association between NS3 and NS5 proteins of dengue virus type 2 in the putative RNA replicase is linked to differential phosphorylation of NS5. J Biol Chem. 1995;270(32):19100–6. PubMed PMID: 7642575.

12. Yamauchi S, Takeuchi K, Chihara K, Sun X, Honjoh C, Yoshiki H, Hotta H, Sada K. Hepatitis C Virus Particle Assembly Involves Phosphorylation of NS5A by the c-Abl Tyrosine Kinase. J Biol Chem. 2015;290(36):21857–64. Epub 2015/07/24. doi: 10.1074/jbc.M115.666859. PubMed PMID: 26203192; PMCID: PMC4571941.

13. Quintavalle M, Sambucini S, Summa V, Orsatti L, Talamo F, De Francesco R, Neddermann P. Hepatitis C virus NS5A is a direct substrate of casein kinase I-alpha, a cellular kinase identified by inhibitor affinity chromatography using specific NS5A hyperphosphorylation inhibitors. J Biol Chem. 2007;282(8):5536–44. Epub 2006/12/15. doi: 10.1074/jbc.M610486200. PubMed PMID: 17166835.

14. Tellinghuisen TL, Foss KL, Treadaway J. Regulation of hepatitis C virion production via phosphorylation of the NS5A protein. PLoS Pathog. 2008;4(3):e1000032. doi: 10.1371/journal.ppat.1000032. PubMed PMID: 18369478; PMCID: PMC2265800.

15. Henchal EA, Gentry MK, McCown JM, Brandt WE. Dengue virus-specific and flavivirus group determinants identified with monoclonal antibodies by indirect immunofluorescence. Am J Trop Med Hyg. 1982;31(4):830–6. Epub 1982/07/01. doi: 10.4269/ajtmh.1982.31.830. PubMed PMID: 6285749.

16. King BR, Hershkowitz D, Eisenhauer PL, Weir ME, Ziegler CM, Russo J, Bruce EA, Ballif BA, Botten J. A map of the arenavirus nucleoprotein-host protein interactome reveals that Junin virus selectively impairs the antiviral activity of PKR. J Virol. 2017. doi: 10.1128/JVI.00763-17. PubMed PMID: 28539447.

17. Mukhopadhyay S, Kuhn RJ, Rossmann MG. A structural perspective of the flavivirus life cycle. Nat Rev Microbiol. 2005;3(1):13–22. Epub 2004/12/21. doi: 10.1038/nrmicro1067. PubMed PMID: 15608696.

18. Akey DL, Brown WC, Dutta S, Konwerski J, Jose J, Jurkiw TJ, DelProposto J, Ogata CM, Skiniotis G, Kuhn RJ, Smith JL. Flavivirus NS1 structures reveal surfaces for associations with membranes and the immune system. Science. 2014;343(6173):881-5. Epub 2014/02/08. doi: 10.1126/science.1247749. PubMed PMID: 24505133; PMCID: PMC4263348.

19. Rastogi M, Sharma N, Singh SK. Flavivirus NS1: a multifaceted enigmatic viral protein. Virol J. 2016;13:131. Epub 2016/07/31. doi: 10.1186/s12985-016-0590-7. PubMed PMID: 27473856; PMCID: PMC4966872.

20. Widman DG, Young E, Yount BL, Plante KS, Gallichotte EN, Carbaugh DL, Peck KM, Plante J, Swanstrom J, Heise MT, Lazear HM, Baric RS. A Reverse Genetics Platform That Spans the Zika Virus Family Tree. MBio. 2017;8(2). doi: 10.1128/mBio.02014-16. PubMed PMID: 28270583; PMCID: PMC5340872.

21. Espinosa D, Mendy J, Manayani D, Vang L, Wang C, Richard T, Guenther B, Aruri J, Avanzini J, Garduno F, Farness P, Gurwith M, Smith J, Harris E, Alexander J. Passive Transfer of Immune Sera Induced by a Zika Virus-Like Particle Vaccine Protects AG129 Mice Against Lethal Zika Virus Challenge. EBioMedicine. 2018;27:61–70. Epub 2017/12/23. doi: 10.1016/j.ebiom.2017.12.010. PubMed PMID: 29269041; PMCID: PMC5828544.

22. Jager S, Cimermancic P, Gulbahce N, Johnson JR, McGovern KE, Clarke SC, Shales M, Mercenne G, Pache L, Li K, Hernandez H, Jang GM, Roth SL, Akiva E, Marlett J, Stephens M, D’Orso I, Fernandes J, Fahey M, Mahon C, O’Donoghue AJ, Todorovic A, Morris JH, Maltby DA, Alber T, Cagney G, Bushman FD, Young JA, Chanda SK, Sundquist WI, Kortemme T, Hernandez RD, Craik CS, Burlingame A, Sali A, Frankel AD, Krogan NJ. Global landscape of HIV-human protein complexes. Nature. 2011;481(7381):365-70. Epub 2011/12/23. doi: 10.1038/nature10719. PubMed PMID: 22190034; PMCID: PMC3310911.

23. Savidis G, McDougall WM, Meraner P, Perreira JM, Portmann JM, Trincucci G, John SP, Aker AM, Renzette N, Robbins DR, Guo Z, Green S, Kowalik TF, Brass AL. Identification of Zika Virus and Dengue Virus Dependency Factors using Functional Genomics. Cell Rep. 2016;16(1):232–46. Epub 2016/06/28. doi: 10.1016/j.celrep.2016.06.028. PubMed PMID: 27342126.

24. Coyaud E, Ranadheera C, Cheng D, Goncalves J, Dyakov BJA, Laurent EMN, St-Germain J, Pelletier L, Gingras AC, Brumell JH, Kim PK, Safronetz D, Raught B. Global Interactomics Uncovers Extensive Organellar Targeting by Zika Virus. Mol Cell Proteomics. 2018;17(11):2242–55. Epub 2018/07/25. doi: 10.1074/mcp.TIR118.000800. PubMed PMID: 30037810; PMCID: PMC6210227.

25. Scaturro P, Stukalov A, Haas DA, Cortese M, Draganova K, Plaszczyca A, Bartenschlager R, Gotz M, Pichlmair A. An orthogonal proteomic survey uncovers novel Zika virus host factors. Nature. 2018;561(7722):253-7. Epub 2018/09/05. doi: 10.1038/s41586-018-0484-5. PubMed PMID: 30177828.

26. Shah PS, Link N, Jang GM, Sharp PP, Zhu T, Swaney DL, Johnson JR, Von Dollen J, Ramage HR, Satkamp L, Newton B, Huttenhain R, Petit MJ, Baum T, Everitt A, Laufman O, Tassetto M, Shales M, Stevenson E, Iglesias GN, Shokat L, Tripathi S, Balasubramaniam V, Webb LG, Aguirre S, Willsey AJ, Garcia-Sastre A, Pollard KS, Cherry S, Gamarnik AV, Marazzi I, Taunton J, Fernandez-Sesma A, Bellen HJ, Andino R, Krogan NJ. Comparative Flavivirus-Host Protein Interaction Mapping Reveals Mechanisms of Dengue and Zika Virus Pathogenesis. Cell. 2018;175(7):1931–45 e18. Epub 2018/12/15. doi: 10.1016/j.cell.2018.11.028. PubMed PMID: 30550790; PMCID: PMC6474419.

27. Li Y, Muffat J, Omer Javed A, Keys HR, Lungjangwa T, Bosch I, Khan M, Virgilio MC, Gehrke L, Sabatini DM, Jaenisch R. Genome-wide CRISPR screen for Zika virus resistance in human neural cells. Proc Natl Acad Sci U S A. 2019;116(19):9527–32. Epub 2019/04/26. doi: 10.1073/pnas.1900867116. PubMed PMID: 31019072; PMCID: PMC6510995.

28. Wang S, Zhang Q, Tiwari SK, Lichinchi G, Yau EH, Hui H, Li W, Furnari F, Rana TM. Integrin alphavbeta5 Internalizes Zika Virus during Neural Stem Cells Infection and Provides a Promising Target for Antiviral Therapy. Cell Rep. 2020;30(4):969–83 e4. Epub 2020/01/21. doi: 10.1016/j.celrep.2019.11.020. PubMed PMID: 31956073; PMCID: PMC7293422.

29. Giraldo MI, Xia H, Aguilera-Aguirre L, Hage A, van Tol S, Shan C, Xie X, Sturdevant GL, Robertson SJ, McNally KL, Meade-White K, Azar SR, Rossi SL, Maury W, Woodson M, Ramage H, Johnson JR, Krogan NJ, Morais MC, Best SM, Shi PY, Rajsbaum R. Envelope protein ubiquitination drives entry and pathogenesis of Zika virus. Nature. 2020;585(7825):414-9. Epub 2020/07/10. doi: 10.1038/s41586-020-2457-8. PubMed PMID: 32641828; PMCID: PMC7501154.

30. Tabata K, Arimoto M, Arakawa M, Nara A, Saito K, Omori H, Arai A, Ishikawa T, Konishi E, Suzuki R, Matsuura Y, Morita E. Unique Requirement for ESCRT Factors in Flavivirus Particle Formation on the Endoplasmic Reticulum. Cell Rep. 2016;16(9):2339–47. Epub 2016/08/23. doi: 10.1016/j.celrep.2016.07.068. PubMed PMID: 27545892.

31. Bhattacharya D, Mayuri, Best SM, Perera R, Kuhn RJ, Striker R. Protein kinase G phosphorylates mosquito-borne flavivirus NS5. J Virol. 2009;83(18):9195–205. doi: 10.1128/JVI.00271-09. PubMed PMID: 19587048; PMCID: PMC2738234.

32. Morozova OV, Tsekhanovskaya NA, Maksimova TG, Bachvalova VN, Matveeva VA, Kit Y. Phosphorylation of tick-borne encephalitis virus NS5 protein. Virus Res. 1997;49(1):9–15. PubMed PMID: 9178492.

33. Lemay KL, Treadaway J, Angulo I, Tellinghuisen TL. A hepatitis C virus NS5A phosphorylation site that regulates RNA replication. J Virol. 2013;87(2):1255–60. Epub 2012/11/02. doi: 10.1128/JVI.02154-12. PubMed PMID: 23115292; PMCID: PMC3554050.

34. Valencia HJ, de Aguiar M, Costa MA, Mendonca DC, Reis EV, Arias NEC, Drumond BP, Bonjardim CA. Evaluation of kinase inhibitors as potential therapeutics for flavivirus infections. Arch Virol. 2021;166(5):1433–8. Epub 2021/03/09. doi: 10.1007/s00705-021-05021-1. PubMed PMID: 33683474; PMCID: PMC7938686.

35. Remsing Rix LL, Rix U, Colinge J, Hantschel O, Bennett KL, Stranzl T, Muller A, Baumgartner C, Valent P, Augustin M, Till JH, Superti-Furga G. Global target profile of the kinase inhibitor bosutinib in primary chronic myeloid leukemia cells. Leukemia. 2009;23(3):477–85. Epub 2008/11/29. doi: 10.1038/leu.2008.334. PubMed PMID: 19039322.

36. Rausch K, Hackett BA, Weinbren NL, Reeder SM, Sadovsky Y, Hunter CA, Schultz DC, Coyne CB, Cherry S. Screening Bioactives Reveals Nanchangmycin as a Broad Spectrum Antiviral Active against Zika Virus. Cell Rep. 2017;18(3):804–15. Epub 2017/01/19. doi: 10.1016/j.celrep.2016.12.068. PubMed PMID: 28099856; PMCID: PMC5270376.

37. Liang Q, Luo Z, Zeng J, Chen W, Foo SS, Lee SA, Ge J, Wang S, Goldman SA, Zlokovic BV, Zhao Z, Jung JU. Zika Virus NS4A and NS4B Proteins Deregulate Akt-mTOR Signaling in Human Fetal Neural Stem Cells to Inhibit Neurogenesis and Induce Autophagy. Cell Stem Cell. 2016;19(5):663–71. doi: 10.1016/j.stem.2016.07.019. PubMed PMID: 27524440; PMCID: PMC5144538.

38. Chiramel AI, Best SM. Role of autophagy in Zika virus infection and pathogenesis. Virus Res. 2018;254:34–40. Epub 2017/09/14. doi: 10.1016/j.virusres.2017.09.006. PubMed PMID: 28899653; PMCID: PMC5844781.

39. Donald CL, Brennan B, Cumberworth SL, Rezelj VV, Clark JJ, Cordeiro MT, Freitas de Oliveira Franca R, Pena LJ, Wilkie GS, Da Silva Filipe A, Davis C, Hughes J, Varjak M, Selinger M, Zuvanov L, Owsianka AM, Patel AH, McLauchlan J, Lindenbach BD, Fall G, Sall AA, Biek R, Rehwinkel J, Schnettler E, Kohl A. Full Genome Sequence and sfRNA Interferon Antagonist Activity of Zika Virus from Recife, Brazil. PLoS Negl Trop Dis. 2016;10(10):e0005048. Epub 2016/10/06. doi: 10.1371/journal.pntd.0005048. PubMed PMID: 27706161; PMCID: PMC5051680.

40. Alzhanova D, Corcoran K, Bailey AG, Long K, Taft-Benz S, Graham RL, Broussard GS, Heise M, Neumann G, Halfmann P, Kawaoka Y, Baric RS, Damania B, Dittmer DP. Novel modulators of p53-signaling encoded by unknown genes of emerging viruses. PLoS Pathog. 2021;17(1):e1009033. Epub 2021/01/08. doi: 10.1371/journal.ppat.1009033. PubMed PMID: 33411764; PMCID: PMC7790267 Corcoran was unable to confirm their authorship contributions. On their behalf, the corresponding author has reported their contributions to the best of their knowledge.

41. Klaus JP, Eisenhauer P, Russo J, Mason AB, Do D, King B, Taatjes D, Cornillez-Ty C, Boyson JE, Thali M, Zheng C, Liao L, Yates JR, 3rd, Zhang B, Ballif BA, Botten JW. The intracellular cargo receptor ERGIC-53 is required for the production of infectious arenavirus, coronavirus, and filovirus particles. Cell Host Microbe. 2013;14(5):522–34. Epub 2013/11/19. doi: 10.1016/j.chom.2013.10.010. PubMed PMID: 24237698; PMCID: PMC3999090.

42. Cornillez-Ty CT, Liao L, Yates JR, 3rd, Kuhn P, Buchmeier MJ. Severe acute respiratory syndrome coronavirus nonstructural protein 2 interacts with a host protein complex involved in mitochondrial biogenesis and intracellular signaling. Journal of Virology. 2009;83(19):10314–8. Epub 2009/07/31. doi: JVI.00842-09 [pii] 10.1128/JVI.00842-09. PubMed PMID: 19640993; PMCID: 2748024.

43. Hutchinson EC, Stegmann M. Purification and Proteomics of Influenza Virions. Methods Mol Biol. 2018;1836:89–120. Epub 2018/08/29. doi: 10.1007/978-1-4939-8678-1_5. PubMed PMID: 30151570.

44. Wisniewski JR, Zougman A, Nagaraj N, Mann M. Universal sample preparation method for proteome analysis. Nat Methods. 2009;6(5):359–62. Epub 2009/04/21. doi: 10.1038/nmeth.1322. PubMed PMID: 19377485.

45. Tyanova S, Temu T, Cox J. The MaxQuant computational platform for mass spectrometry-based shotgun proteomics. Nat Protoc. 2016;11(12):2301–19. Epub 2016/11/04. doi: 10.1038/nprot.2016.136. PubMed PMID: 27809316.

46. Edgar RC. MUSCLE: multiple sequence alignment with high accuracy and high throughput. Nucleic Acids Res. 2004;32(5):1792–7. Epub 2004/03/23. doi: 10.1093/nar/gkh340. PubMed PMID: 15034147; PMCID: PMC390337.

47. Pickett BE, Sadat EL, Zhang Y, Noronha JM, Squires RB, Hunt V, Liu M, Kumar S, Zaremba S, Gu Z, Zhou L, Larson CN, Dietrich J, Klem EB, Scheuermann RH. ViPR: an open bioinformatics database and analysis resource for virology research. Nucleic Acids Res. 2012;40(Database issue):D593-8. Epub 2011/10/19. doi: 10.1093/nar/gkr859. PubMed PMID: 22006842; PMCID: PMC3245011.

48. Waterhouse AM, Procter JB, Martin DM, Clamp M, Barton GJ. Jalview Version 2--a multiple sequence alignment editor and analysis workbench. Bioinformatics. 2009;25(9):1189–91. Epub 2009/01/20. doi: 10.1093/bioinformatics/btp033. PubMed PMID: 19151095; PMCID: PMC2672624.

49. Kong BW, Foster LK, Foster DN. A method for the rapid isolation of virus from cultured cells. Biotechniques. 2008;44(1):97–9. Epub 2008/02/08. doi: 10.2144/000112589. PubMed PMID: 18254386.

