## Supplementary Information 3 for "A global map of the Zika virus phosphoproteome reveals host-driven regulation of viral budding and cytopathogenicity"

Supplementary Information **3**: The tandem mass spectra of discovered post translational modifications on ZIKV proteins. Tandem mass spectra were collected in a Q Exactive Plus mass spectrometer. Each spectrum is labeled with the peptide and ZIKV protein where a modification was found, as well as the residue number on the ZIKV protein. Spectra of residues are shown first without post-translational modifications then the same residues labeled with "p" or "ub" to represent phosphorylation or ubiquitination, respectively. Special characters within peptide sequences indicate modifications as follows: @ = phosphorylation, * = oxidation, ~ = ubiquitination, ^ = carbamidomethylation, # = acrylamidation. Accompanying tables show the corresponding calculated and measured b- and y-type ions (colored in blue and red, respectively) indicating identified fragment ion masses.

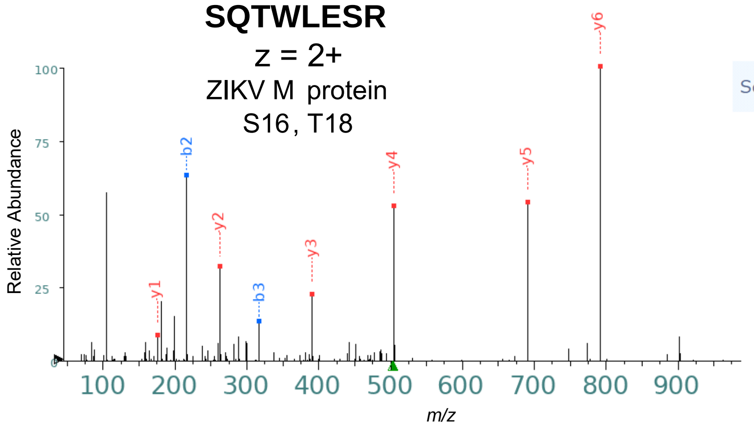


| Seq | # | b: Δ Error | b | y | y: Δ Error | +1 |
|---|---|---|---|---|---|---|
| S | 1 | --- | 88.039 | --- | --- | 8 |
| Q | 2 | -0.762 | 216.098 | 919.463 | --- | 7 |
| T | 3 | -1.321 | 317.146 | 791.405 | -1.191 | 6 |
| W | 4 | --- | 503.225 | 690.357 | -1.086 | 5 |
| L | 5 | --- | 616.309 | 504.278 | -0.947 | 4 |
| E | 6 | --- | 745.352 | 391.194 | -1.173 | 3 |
| S | 7 | --- | 832.384 | 262.151 | -0.971 | 2 |
| R | 8 | --- | --- | 175.119 | -0.407 | 1 |

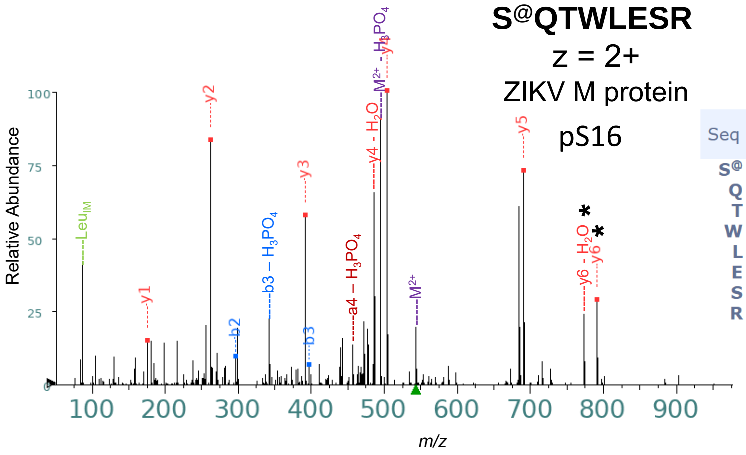


| Seq | # | b: Δ Error | b | y | y: Δ Error | +1 |
|---|---|---|---|---|---|---|
| S@ | 1 | --- | 168.006 | --- | --- | 8 |
| Q | 2 | -1.145 | 296.064 | 919.463 | --- | 7 |
| T | 3 | -2.262 | 397.112 | 791.405 | -0.343 | 6 |
| W | 4 | --- | 583.191 | 690.357 | -0.909 | 5 |
| L | 5 | --- | 696.275 | 504.278 | -0.947 | 4 |
| E | 6 | --- | 825.318 | 391.194 | -1.095 | 3 |
| S | 7 | --- | 912.350 | 262.151 | -1.321 | 2 |
| R | 8 | --- | --- | 175.119 | -0.843 | 1 |

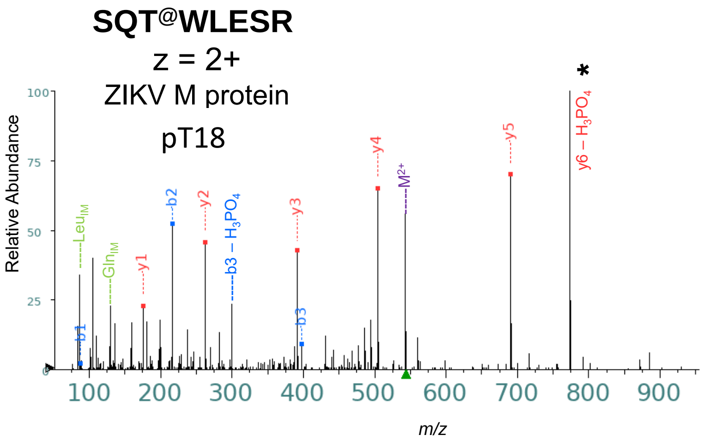


| Seq | # | b: Δ Error | b | y | y: Δ Error | +1 |
|---|---|---|---|---|---|---|
| S | 1 | 4.185 | 88.039 | --- | --- | 8 |
| Q | 2 | -1.186 | 216.098 | 999.430 | --- | 7 |
| T@ | 3 | -1.878 | 397.112 | 871.371 | --- | 6 |
| W | 4 | --- | 583.191 | 690.357 | -1.705 | 5 |
| L | 5 | --- | 696.275 | 504.278 | -1.431 | 4 |
| E | 6 | --- | 825.318 | 391.194 | -1.797 | 3 |
| S | 7 | --- | 912.350 | 262.151 | -1.553 | 2 |
| R | 8 | --- | --- | 175.119 | -1.279 | 1 |

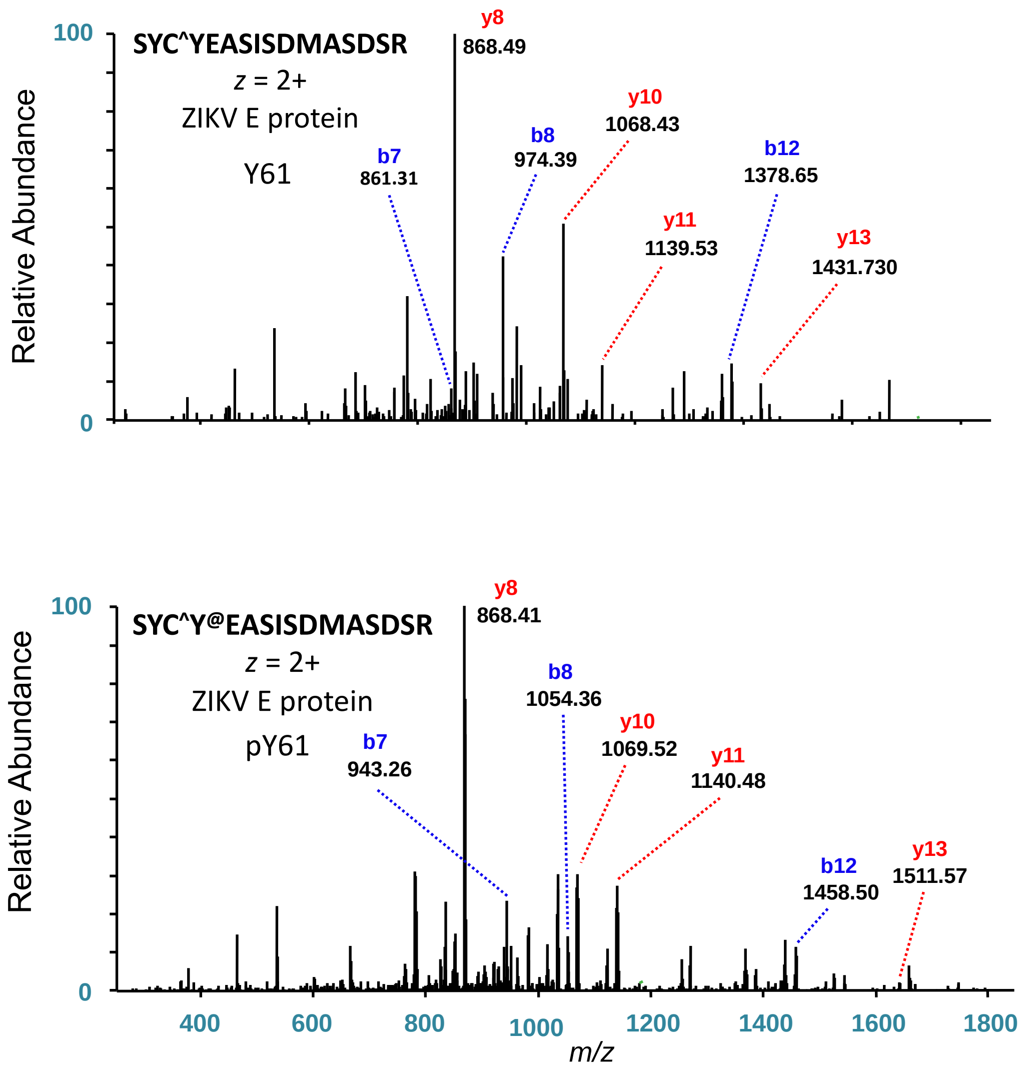

100
SYC^YEASISDMASDSR
z = 2+
ZIKV E protein
Y61
Relative Abundance
0
y8
868.49
b7
861.31
b8
974.39
y10
1068.43
y11
1139.53
b12
1378.65
y13
1431.730
100
SYC^Y@EASISDMASDSR
z = 2+
ZIKV E protein
pY61
Relative Abundance
0
y8
868.41
b7
943.26
b8
1054.36
y10
1069.52
y11
1140.48
b12
1458.50
y13
1511.57
400
600
800
1000
1200
1400
1600
1800
m/z

**IM*LSVHGSQHSGM*IVNDTGHETDENR**

z = 4+

ZIKV E protein

S142

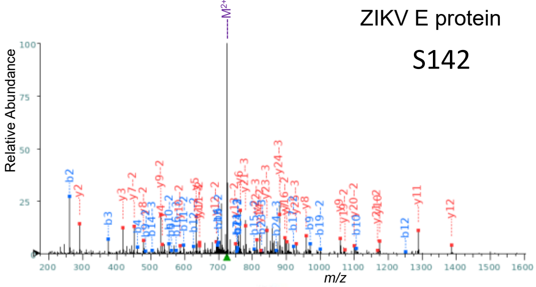


+1

| Seq | # | b: Δ Error | b | y | y: Δ Error | +1 |
|---|---|---|---|---|---|---|
| I | 1 | --- | 114.091 | --- | --- | 26 |
| M* | 2 | -1.635 | 261.127 | 2783.211 | --- | 25 |
| L | 3 | -1.435 | 374.211 | 2636.175 | --- | 24 |
| S | 4 | -0.404 | 461.243 | 2523.091 | --- | 23 |
| V | 5 | -2.881 | 560.311 | 2436.059 | --- | 22 |
| H | 6 | -2.683 | 697.370 | 2336.991 | --- | 21 |
| G | 7 | -2.534 | 754.392 | 2199.932 | --- | 20 |
| S | 8 | --- | 841.424 | 2142.910 | --- | 19 |
| Q | 9 | -0.177 | 969.482 | 2055.878 | --- | 18 |
| H | 10 | -0.111 | 1106.541 | 1927.820 | --- | 17 |
| S | 11 | --- | 1193.573 | 1790.761 | --- | 16 |
| G | 12 | -1.583 | 1250.595 | 1703.729 | --- | 15 |
| M* | 13 | --- | 1397.630 | 1646.707 | --- | 14 |
| I | 14 | --- | 1510.714 | 1499.672 | --- | 13 |
| V | 15 | --- | 1609.783 | 1386.588 | -1.330 | 12 |
| N | 16 | --- | 1723.825 | 1287.520 | 0.411 | 11 |
| D | 17 | --- | 1838.852 | 1173.477 | -0.936 | 10 |
| T | 18 | --- | 1939.900 | 1058.450 | -0.840 | 9 |
| G | 19 | --- | 1996.922 | 957.402 | -0.727 | 8 |
| H | 20 | --- | 2133.980 | 900.381 | -1.542 | 7 |
| E | 21 | --- | 2263.023 | 763.322 | -1.402 | 6 |
| T | 22 | --- | 2364.071 | 634.279 | -1.798 | 5 |
| D | 23 | --- | 2479.098 | 533.231 | -1.548 | 4 |
| E | 24 | --- | 2608.140 | 418.204 | -1.399 | 3 |
| N | 25 | --- | 2722.183 | 289.162 | -1.740 | 2 |
| R | 26 | --- | --- | 175.119 | --- | 1 |

+2

| Seq | # | b: Δ Error | b | y | y: Δ Error | +1 |
|---|---|---|---|---|---|---|
| I | 1 | --- | 57.549 | --- | --- | 26 |
| M* | 2 | --- | 131.067 | 1392.109 | --- | 25 |
| L | 3 | --- | 187.609 | 1318.591 | --- | 24 |
| S | 4 | --- | 231.125 | 1262.049 | --- | 23 |
| V | 5 | --- | 280.659 | 1218.533 | --- | 22 |
| H | 6 | --- | 349.189 | 1168.999 | -0.567 | 21 |
| G | 7 | --- | 377.699 | 1100.470 | 0.208 | 20 |
| S | 8 | --- | 421.215 | 1071.959 | -1.618 | 19 |
| Q | 9 | -2.768 | 485.245 | 1028.443 | --- | 18 |
| H | 10 | -0.068 | 553.774 | 964.414 | --- | 17 |
| S | 11 | -1.430 | 597.290 | 895.884 | -0.243 | 16 |
| G | 12 | -1.836 | 625.801 | 852.368 | --- | 15 |
| M* | 13 | -1.555 | 699.319 | 823.857 | -0.990 | 14 |
| I | 14 | 0.205 | 755.861 | 750.340 | -0.599 | 13 |
| V | 15 | 3.265 | 805.395 | 693.798 | -2.086 | 12 |
| N | 16 | --- | 862.416 | 644.263 | -4.573 | 11 |
| D | 17 | -0.285 | 919.930 | 587.242 | 0.353 | 10 |
| T | 18 | --- | 970.454 | 529.729 | -0.794 | 9 |
| G | 19 | -0.043 | 998.964 | 479.205 | -2.141 | 8 |
| H | 20 | --- | 1067.494 | 450.694 | -1.012 | 7 |
| E | 21 | --- | 1132.015 | 382.164 | --- | 6 |
| T | 22 | --- | 1182.539 | 317.643 | --- | 5 |
| D | 23 | --- | 1240.052 | 267.119 | --- | 4 |
| E | 24 | --- | 1304.574 | 209.606 | --- | 3 |
| N | 25 | --- | 1361.595 | 145.085 | --- | 2 |
| R | 26 | --- | --- | 88.063 | --- | 1 |

+3

| Seq | # | b: Δ Error | b | y | y: Δ Error | +1 |
|---|---|---|---|---|---|---|
| I | 1 | --- | 38.702 | --- | --- | 26 |
| M* | 2 | --- | 87.714 | 928.408 | -0.917 | 25 |
| L | 3 | --- | 125.408 | 879.397 | 0.165 | 24 |
| S | 4 | --- | 154.419 | 841.702 | 0.591 | 23 |
| V | 5 | --- | 187.442 | 812.691 | 1.282 | 22 |
| H | 6 | --- | 233.128 | 779.668 | -0.415 | 21 |
| G | 7 | --- | 252.135 | 733.982 | --- | 20 |
| S | 8 | --- | 281.146 | 714.975 | --- | 19 |
| Q | 9 | --- | 323.832 | 685.964 | --- | 18 |
| H | 10 | --- | 369.519 | 643.278 | 2.449 | 17 |
| S | 11 | --- | 398.529 | 597.592 | --- | 16 |
| G | 12 | --- | 417.536 | 568.581 | --- | 15 |
| M* | 13 | --- | 466.548 | 549.574 | --- | 14 |
| I | 14 | 4.735 | 504.243 | 500.562 | --- | 13 |
| V | 15 | --- | 537.266 | 462.868 | --- | 12 |
| N | 16 | 2.074 | 575.280 | 429.845 | --- | 11 |
| D | 17 | --- | 613.622 | 391.830 | --- | 10 |
| T | 18 | --- | 647.305 | 353.488 | --- | 9 |
| G | 19 | --- | 666.312 | 319.806 | --- | 8 |
| H | 20 | --- | 711.998 | 300.798 | --- | 7 |
| E | 21 | -0.099 | 755.013 | 255.112 | --- | 6 |
| T | 22 | --- | 788.695 | 212.098 | --- | 5 |
| D | 23 | -2.860 | 827.037 | 178.415 | --- | 4 |
| E | 24 | 1.447 | 870.052 | 140.073 | --- | 3 |
| N | 25 | --- | 908.066 | 97.059 | --- | 2 |
| R | 26 | --- | --- | 59.045 | --- | 1 |

**IM*LS@VHGSQHSGM*IVNDTGHETDENR**

z = 4+

ZIKV E

pS142

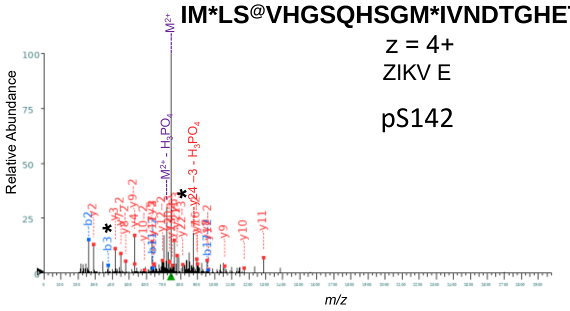


+1

| Seq | # | b: Δ Error | b | y | y: Δ Error | +1 |
|---|---|---|---|---|---|---|
| I | 1 | --- | 114.091 | --- | --- | 26 |
| M* | 2 | -1.051 | 261.127 | 2863.177 | --- | 25 |
| L | 3 | 0.440 | 374.211 | 2716.142 | --- | 24 |
| S@ | 4 | --- | 541.209 | 2603.058 | --- | 23 |
| V | 5 | --- | 640.278 | 2436.059 | --- | 22 |
| H | 6 | --- | 777.336 | 2336.991 | --- | 21 |
| G | 7 | --- | 834.358 | 2199.932 | --- | 20 |
| S | 8 | --- | 921.390 | 2142.910 | --- | 19 |
| Q | 9 | --- | 1049.449 | 2055.878 | --- | 18 |
| H | 10 | --- | 1186.507 | 1927.820 | --- | 17 |
| S | 11 | --- | 1273.539 | 1790.761 | --- | 16 |
| G | 12 | --- | 1330.561 | 1703.729 | --- | 15 |
| M* | 13 | --- | 1477.596 | 1646.707 | --- | 14 |
| I | 14 | --- | 1590.680 | 1499.672 | --- | 13 |
| V | 15 | --- | 1689.749 | 1386.588 | --- | 12 |
| N | 16 | --- | 1803.792 | 1287.520 | -0.821 | 11 |
| D | 17 | --- | 1918.819 | 1173.477 | -0.416 | 10 |
| T | 18 | --- | 2019.866 | 1058.450 | 1.697 | 9 |
| G | 19 | --- | 2076.888 | 957.402 | -0.090 | 8 |
| H | 20 | --- | 2213.947 | 900.381 | 1.509 | 7 |
| E | 21 | --- | 2342.989 | 763.322 | -1.242 | 6 |
| T | 22 | --- | 2444.037 | 634.279 | -1.221 | 5 |
| D | 23 | --- | 2559.064 | 533.231 | 0.856 | 4 |
| E | 24 | --- | 2688.107 | 418.204 | -1.399 | 3 |
| N | 25 | --- | 2802.149 | 289.162 | -0.895 | 2 |
| R | 26 | --- | --- | 175.119 | --- | 1 |

+2

| Seq | # | b: Δ Error | b | y | y: Δ Error | +1 |
|---|---|---|---|---|---|---|
| I | 1 | --- | 57.549 | --- | --- | 26 |
| M* | 2 | --- | 131.067 | 1432.092 | --- | 25 |
| L | 3 | --- | 187.609 | 1358.574 | --- | 24 |
| S@ | 4 | --- | 271.108 | 1302.032 | --- | 23 |
| V | 5 | --- | 320.642 | 1218.533 | --- | 22 |
| H | 6 | --- | 389.172 | 1168.999 | --- | 21 |
| G | 7 | --- | 417.683 | 1100.470 | --- | 20 |
| S | 8 | --- | 461.199 | 1071.959 | --- | 19 |
| Q | 9 | --- | 525.228 | 1028.443 | --- | 18 |
| H | 10 | --- | 593.757 | 964.414 | -1.783 | 17 |
| S | 11 | 1.995 | 637.273 | 895.884 | -1.674 | 16 |
| G | 12 | --- | 665.784 | 852.368 | --- | 15 |
| M* | 13 | --- | 739.302 | 823.857 | --- | 14 |
| I | 14 | --- | 795.844 | 750.340 | 3.793 | 13 |
| V | 15 | --- | 845.378 | 693.798 | 1.345 | 12 |
| N | 16 | --- | 902.400 | 644.263 | -2.204 | 11 |
| D | 17 | 2.450 | 959.913 | 587.242 | -2.246 | 10 |
| T | 18 | --- | 1010.437 | 529.729 | -0.679 | 9 |
| G | 19 | --- | 1038.948 | 479.205 | -0.103 | 8 |
| H | 20 | --- | 1107.477 | 450.694 | -0.268 | 7 |
| E | 21 | --- | 1171.998 | 382.164 | --- | 6 |
| T | 22 | --- | 1222.522 | 317.643 | --- | 5 |
| D | 23 | --- | 1280.036 | 267.119 | --- | 4 |
| E | 24 | --- | 1344.557 | 209.606 | --- | 3 |
| N | 25 | --- | 1401.578 | 145.085 | --- | 2 |
| R | 26 | --- | --- | 88.063 | --- | 1 |

+3

| Seq | # | b: Δ Error | b | y | y: Δ Error | +1 |
|---|---|---|---|---|---|---|
| I | 1 | --- | 38.702 | --- | --- | 26 |
| M* | 2 | --- | 87.714 | 955.064 | --- | 25 |
| L | 3 | --- | 125.408 | 906.052 | --- | 24 |
| S@ | 4 | --- | 181.075 | 868.357 | --- | 23 |
| V | 5 | --- | 214.097 | 812.691 | -0.521 | 22 |
| H | 6 | --- | 259.784 | 779.668 | 0.289 | 21 |
| G | 7 | --- | 278.791 | 733.982 | 0.258 | 20 |
| S | 8 | --- | 307.802 | 714.975 | --- | 19 |
| Q | 9 | --- | 350.488 | 685.964 | --- | 18 |
| H | 10 | --- | 396.174 | 643.278 | 0.931 | 17 |
| S | 11 | --- | 425.185 | 597.592 | --- | 16 |
| G | 12 | --- | 444.192 | 568.581 | --- | 15 |
| M* | 13 | --- | 493.204 | 549.574 | --- | 14 |
| I | 14 | --- | 530.898 | 500.562 | --- | 13 |
| V | 15 | --- | 563.921 | 462.868 | --- | 12 |
| N | 16 | --- | 601.935 | 429.845 | --- | 11 |
| D | 17 | --- | 640.278 | 391.830 | --- | 10 |
| T | 18 | --- | 673.960 | 353.488 | --- | 9 |
| G | 19 | --- | 692.967 | 319.806 | --- | 8 |
| H | 20 | --- | 738.654 | 300.798 | --- | 7 |
| E | 21 | --- | 781.668 | 255.112 | --- | 6 |
| T | 22 | --- | 815.351 | 212.098 | --- | 5 |
| D | 23 | --- | 853.693 | 178.415 | --- | 4 |
| E | 24 | --- | 896.707 | 140.073 | --- | 3 |
| N | 25 | --- | 934.721 | 97.059 | --- | 2 |
| R | 26 | --- | --- | 59.045 | --- | 1 |

**IM*LSVHGSQHSGM*IVNDTGHETDENR**

*z* = 4+

ZIKV E protein

S149, T156

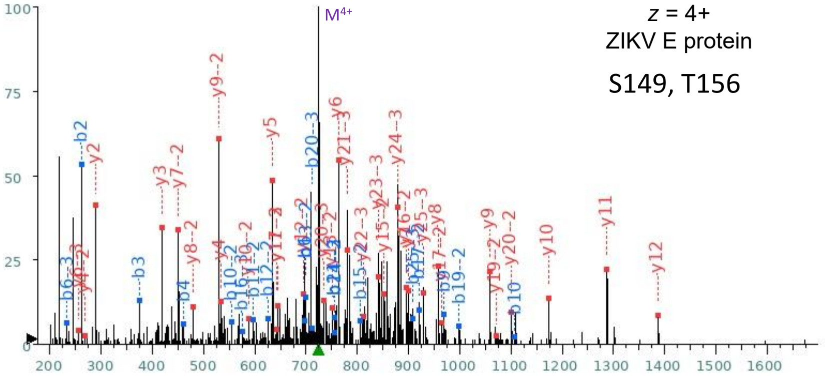


+1

| Seq | # | b: Δ Error | b | y | y: Δ Error | +1 |
|---|---|---|---|---|---|---|
| I | 1 | --- | 114.091 | --- | --- | 26 |
| M* | 2 | -1.635 | 261.127 | 2783.211 | --- | 25 |
| L | 3 | -1.761 | 374.211 | 2636.175 | --- | 24 |
| S | 4 | -0.404 | 461.243 | 2523.091 | --- | 23 |
| V | 5 | --- | 560.311 | 2436.059 | --- | 22 |
| H | 6 | -2.421 | 697.370 | 2336.991 | --- | 21 |
| G | 7 | --- | 754.392 | 2199.932 | --- | 20 |
| S | 8 | --- | 841.424 | 2142.910 | --- | 19 |
| Q | 9 | -4.961 | 969.482 | 2055.878 | --- | 18 |
| H | 10 | 1.433 | 1106.541 | 1927.820 | --- | 17 |
| S | 11 | --- | 1193.573 | 1790.761 | --- | 16 |
| G | 12 | --- | 1250.595 | 1703.729 | --- | 15 |
| M* | 13 | --- | 1397.630 | 1646.707 | --- | 14 |
| I | 14 | --- | 1510.714 | 1499.672 | --- | 13 |
| V | 15 | --- | 1609.783 | 1386.588 | -5.380 | 12 |
| N | 16 | --- | 1723.825 | 1287.520 | -1.675 | 11 |
| D | 17 | --- | 1838.852 | 1173.477 | -1.352 | 10 |
| T | 18 | --- | 1939.900 | 1058.450 | 0.659 | 9 |
| G | 19 | --- | 1996.922 | 957.402 | -2.257 | 8 |
| H | 20 | --- | 2133.980 | 900.381 | -2.965 | 7 |
| E | 21 | --- | 2263.023 | 763.322 | -1.722 | 6 |
| T | 22 | --- | 2364.071 | 634.279 | -1.702 | 5 |
| D | 23 | --- | 2479.098 | 533.231 | -2.120 | 4 |
| E | 24 | --- | 2608.140 | 418.204 | -2.056 | 3 |
| N | 25 | --- | 2722.183 | 289.162 | -1.634 | 2 |
| R | 26 | --- | --- | 175.119 | --- | 1 |

+2

| Seq | # | b: Δ Error | b | y | y: Δ Error | +1 |
|---|---|---|---|---|---|---|
| I | 1 | --- | 57.549 | --- | --- | 26 |
| M* | 2 | --- | 131.067 | 1392.109 | --- | 25 |
| L | 3 | --- | 187.609 | 1318.591 | --- | 24 |
| S | 4 | --- | 231.125 | 1262.049 | --- | 23 |
| V | 5 | --- | 280.659 | 1218.533 | --- | 22 |
| H | 6 | --- | 349.189 | 1168.999 | --- | 21 |
| G | 7 | --- | 377.699 | 1100.470 | -2.232 | 20 |
| S | 8 | --- | 421.215 | 1071.959 | -0.024 | 19 |
| Q | 9 | --- | 485.245 | 1028.443 | --- | 18 |
| H | 10 | 1.034 | 553.774 | 964.414 | -3.998 | 17 |
| S | 11 | -2.554 | 597.290 | 895.884 | -3.581 | 16 |
| G | 12 | -1.933 | 625.801 | 852.368 | 2.779 | 15 |
| M* | 13 | 0.540 | 699.319 | 823.857 | --- | 14 |
| I | 14 | -1.330 | 755.861 | 750.340 | -0.518 | 13 |
| V | 15 | -1.130 | 805.395 | 693.798 | -0.151 | 12 |
| N | 16 | --- | 862.416 | 644.263 | -2.868 | 11 |
| D | 17 | -2.607 | 919.930 | 587.242 | -4.844 | 10 |
| T | 18 | --- | 970.454 | 529.729 | -1.024 | 9 |
| G | 19 | -4.564 | 998.964 | 479.205 | -2.650 | 8 |
| H | 20 | --- | 1067.494 | 450.694 | -1.690 | 7 |
| E | 21 | --- | 1132.015 | 382.164 | --- | 6 |
| T | 22 | --- | 1182.539 | 317.643 | --- | 5 |
| D | 23 | --- | 1240.052 | 267.119 | -40.529 | 4 |
| E | 24 | --- | 1304.574 | 209.606 | --- | 3 |
| N | 25 | --- | 1361.595 | 145.085 | --- | 2 |
| R | 26 | --- | --- | 88.063 | --- | 1 |

+3

| Seq | # | b: Δ Error | b | y | y: Δ Error | +1 |
|---|---|---|---|---|---|---|
| I | 1 | --- | 38.702 | --- | --- | 26 |
| M* | 2 | --- | 87.714 | 928.408 | 0.990 | 25 |
| L | 3 | --- | 125.408 | 879.397 | -2.542 | 24 |
| S | 4 | --- | 154.419 | 841.702 | -3.325 | 23 |
| V | 5 | --- | 187.442 | 812.691 | -1.272 | 22 |
| H | 6 | 13.280 | 233.128 | 779.668 | -1.276 | 21 |
| G | 7 | --- | 252.135 | 733.982 | -6.977 | 20 |
| S | 8 | --- | 281.146 | 714.975 | --- | 19 |
| Q | 9 | --- | 323.832 | 685.964 | --- | 18 |
| H | 10 | --- | 369.519 | 643.278 | -3.149 | 17 |
| S | 11 | --- | 398.529 | 597.592 | --- | 16 |
| G | 12 | --- | 417.536 | 568.581 | --- | 15 |
| M* | 13 | --- | 466.548 | 549.574 | --- | 14 |
| I | 14 | --- | 504.243 | 500.562 | --- | 13 |
| V | 15 | --- | 537.266 | 462.868 | --- | 12 |
| N | 16 | 0.483 | 575.280 | 429.845 | --- | 11 |
| D | 17 | --- | 613.622 | 391.830 | --- | 10 |
| T | 18 | --- | 647.305 | 353.488 | --- | 9 |
| G | 19 | --- | 666.312 | 319.806 | --- | 8 |
| H | 20 | -3.996 | 711.998 | 300.798 | --- | 7 |
| E | 21 | -0.827 | 755.013 | 255.112 | -16.465 | 6 |
| T | 22 | --- | 788.695 | 212.098 | --- | 5 |
| D | 23 | --- | 827.037 | 178.415 | --- | 4 |
| E | 24 | --- | 870.052 | 140.073 | --- | 3 |
| N | 25 | 7.608 | 908.066 | 97.059 | --- | 2 |
| R | 26 | --- | --- | 59.045 | --- | 1 |

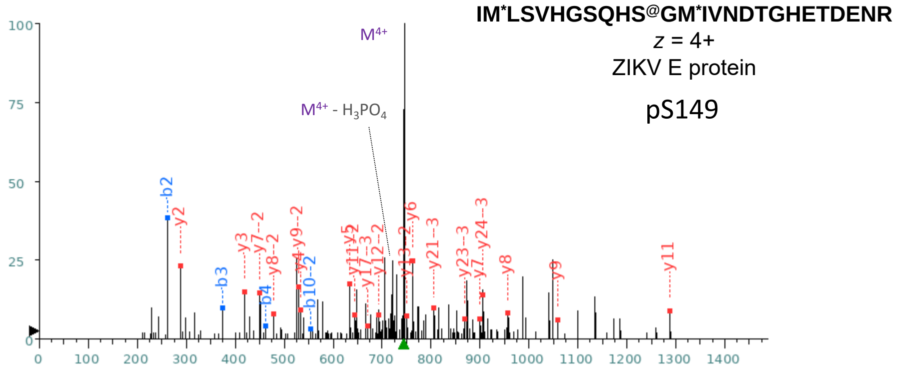


**+1**

| Seq | # | b: Δ Error | b | y | y: Δ Error | +1 |
|---|---|---|---|---|---|---|
| I | 1 | --- | 114.091 | --- | --- | 26 |
| M* | 2 | 0.352 | **261.127** | 2863.177 | --- | 25 |
| L | 3 | 1.908 | **374.211** | 2716.142 | --- | 24 |
| S | 4 | 0.258 | **461.243** | 2603.058 | --- | 23 |
| V | 5 | --- | 560.311 | 2516.026 | --- | 22 |
| H | 6 | --- | 697.370 | 2416.957 | --- | 21 |
| G | 7 | --- | 754.392 | 2279.898 | --- | 20 |
| S | 8 | --- | 841.424 | 2222.877 | --- | 19 |
| Q | 9 | --- | 969.482 | 2135.845 | --- | 18 |
| H | 10 | --- | 1106.541 | 2007.786 | --- | 17 |
| S@ | 11 | --- | 1273.539 | 1870.727 | --- | 16 |
| G | 12 | --- | 1330.561 | 1703.729 | --- | 15 |
| M* | 13 | --- | 1477.596 | 1646.707 | --- | 14 |
| I | 14 | --- | 1590.680 | 1499.672 | --- | 13 |
| V | 15 | --- | 1689.749 | 1386.588 | --- | 12 |
| N | 16 | --- | 1803.792 | **1287.520** | 1.644 | 11 |
| D | 17 | --- | 1918.819 | 1173.477 | --- | 10 |
| T | 18 | --- | 2019.866 | **1058.450** | 5.503 | 9 |
| G | 19 | --- | 2076.888 | **957.402** | -2.576 | 8 |
| H | 20 | --- | 2213.947 | **900.381** | 1.509 | 7 |
| E | 21 | --- | 2342.989 | **763.322** | 0.038 | 6 |
| T | 22 | --- | 2444.037 | **634.279** | 1.859 | 5 |
| D | 23 | --- | 2559.064 | **533.231** | 2.230 | 4 |
| E | 24 | --- | 2688.107 | **418.204** | 1.228 | 3 |
| N | 25 | --- | 2802.149 | **289.162** | -1.634 | 2 |
| R | 26 | --- | --- | 175.119 | --- | 1 |

**+2**

| Seq | # | b: Δ Error | b | y | y: Δ Error | +1 |
|---|---|---|---|---|---|---|
| I | 1 | --- | 57.549 | --- | --- | 26 |
| M* | 2 | --- | 131.067 | 1432.092 | --- | 25 |
| L | 3 | --- | 187.609 | 1358.574 | --- | 24 |
| S | 4 | --- | 231.125 | 1302.032 | --- | 23 |
| V | 5 | --- | 280.659 | 1258.516 | --- | 22 |
| H | 6 | --- | 349.189 | 1208.982 | --- | 21 |
| G | 7 | --- | 377.699 | 1140.453 | --- | 20 |
| S | 8 | --- | 421.215 | 1111.942 | --- | 19 |
| Q | 9 | --- | 485.245 | 1068.426 | --- | 18 |
| H | 10 | 0.263 | **553.774** | 1004.397 | --- | 17 |
| S@ | 11 | --- | 637.273 | 935.867 | --- | 16 |
| G | 12 | --- | 665.784 | 852.368 | --- | 15 |
| M* | 13 | --- | 739.302 | 823.857 | --- | 14 |
| I | 14 | --- | 795.844 | **750.340** | 6.478 | 13 |
| V | 15 | --- | 845.378 | **693.798** | 2.576 | 12 |
| N | 16 | --- | 902.400 | **644.263** | -0.215 | 11 |
| D | 17 | --- | 959.913 | 587.242 | --- | 10 |
| T | 18 | --- | 1010.437 | **529.729** | -0.679 | 9 |
| G | 19 | --- | 1038.948 | **479.205** | 0.598 | 8 |
| H | 20 | --- | 1107.477 | **450.694** | -0.606 | 7 |
| E | 21 | --- | 1171.998 | 382.164 | --- | 6 |
| T | 22 | --- | 1222.522 | 317.643 | --- | 5 |
| D | 23 | --- | 1280.036 | 267.119 | --- | 4 |
| E | 24 | --- | 1344.557 | 209.606 | --- | 3 |
| N | 25 | --- | 1401.578 | 145.085 | --- | 2 |
| R | 26 | --- | --- | 88.063 | --- | 1 |

**+3**

| Seq | # | b: Δ Error | b | y | y: Δ Error | +1 |
|---|---|---|---|---|---|---|
| I | 1 | --- | 38.702 | --- | --- | 26 |
| M* | 2 | --- | 87.714 | 955.064 | --- | 25 |
| L | 3 | --- | 125.408 | **906.052** | -2.048 | 24 |
| S | 4 | --- | 154.419 | **868.357** | 4.032 | 23 |
| V | 5 | --- | 187.442 | 839.347 | --- | 22 |
| H | 6 | --- | 233.128 | **806.324** | 0.599 | 21 |
| G | 7 | --- | 252.135 | 760.638 | --- | 20 |
| S | 8 | --- | 281.146 | 741.630 | --- | 19 |
| Q | 9 | --- | 323.832 | 712.620 | --- | 18 |
| H | 10 | --- | 369.519 | **669.934** | 5.834 | 17 |
| S@ | 11 | --- | 425.185 | 624.247 | --- | 16 |
| G | 12 | --- | 444.192 | 568.581 | --- | 15 |
| M* | 13 | --- | 493.204 | 549.574 | --- | 14 |
| I | 14 | --- | 530.898 | 500.562 | --- | 13 |
| V | 15 | --- | 563.921 | 462.868 | --- | 12 |
| N | 16 | --- | 601.935 | 429.845 | --- | 11 |
| D | 17 | --- | 640.278 | 391.830 | --- | 10 |
| T | 18 | --- | 673.960 | 353.488 | --- | 9 |
| G | 19 | --- | 692.967 | 319.806 | --- | 8 |
| H | 20 | --- | 738.654 | 300.798 | --- | 7 |
| E | 21 | --- | 781.668 | 255.112 | --- | 6 |
| T | 22 | --- | 815.351 | 212.098 | --- | 5 |
| D | 23 | --- | 853.693 | 178.415 | --- | 4 |
| E | 24 | --- | 896.707 | 140.073 | --- | 3 |
| N | 25 | --- | 934.721 | 97.059 | --- | 2 |
| R | 26 | --- | --- | 59.045 | --- | 1 |

**IM*LSVHGSQHSGM*IVNDT@GHETDENR**

*z* = 4+

ZIKV E protein

pT156

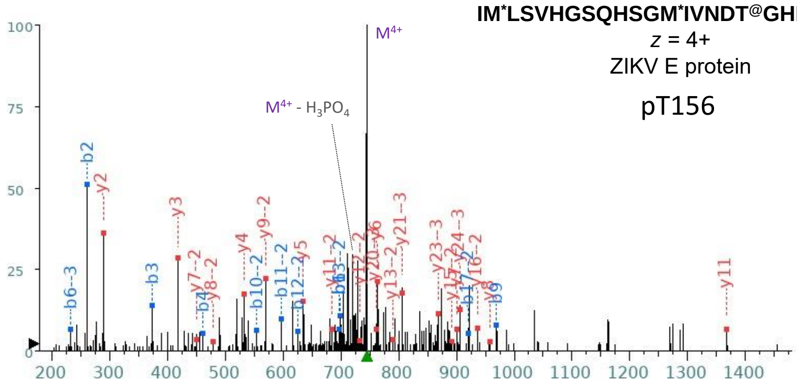


+1

| Seq | # | b: Δ Error | b | y | y: Δ Error | +1 |
|---|---|---|---|---|---|---|
| I | 1 | --- | 114.091 | --- | --- | 26 |
| M* | 2 | -0.349 | 261.127 | 2863.177 | --- | 25 |
| L | 3 | -0.049 | 374.211 | 2716.142 | --- | 24 |
| S | 4 | 0.721 | 461.243 | 2603.058 | --- | 23 |
| V | 5 | --- | 560.311 | 2516.026 | --- | 22 |
| H | 6 | 1.605 | 697.370 | 2416.957 | --- | 21 |
| G | 7 | --- | 754.392 | 2279.898 | --- | 20 |
| S | 8 | --- | 841.424 | 2222.877 | --- | 19 |
| Q | 9 | -0.366 | 969.482 | 2135.845 | --- | 18 |
| H | 10 | 2.205 | 1106.541 | 2007.786 | --- | 17 |
| S | 11 | --- | 1193.573 | 1870.727 | --- | 16 |
| G | 12 | --- | 1250.595 | 1783.695 | --- | 15 |
| M* | 13 | --- | 1397.630 | 1726.674 | --- | 14 |
| I | 14 | --- | 1510.714 | 1579.638 | --- | 13 |
| V | 15 | --- | 1609.783 | 1466.554 | --- | 12 |
| N | 16 | --- | 1723.825 | 1367.486 | -2.664 | 11 |
| D | 17 | --- | 1838.852 | 1253.443 | 5.923 | 10 |
| T@ | 18 | --- | 2019.866 | 1138.416 | -0.693 | 9 |
| G | 19 | --- | 2076.888 | 957.402 | 7.306 | 8 |
| H | 20 | --- | 2213.947 | 900.381 | 2.119 | 7 |
| E | 21 | --- | 2342.989 | 763.322 | 0.117 | 6 |
| T | 22 | --- | 2444.037 | 634.279 | -1.702 | 5 |
| D | 23 | --- | 2559.064 | 533.231 | -0.518 | 4 |
| E | 24 | --- | 2688.107 | 418.204 | -0.305 | 3 |
| N | 25 | --- | 2802.149 | 289.162 | -0.684 | 2 |
| R | 26 | --- | --- | 175.119 | --- | 1 |

+2

| Seq | # | b: Δ Error | b | y | y: Δ Error | +1 |
|---|---|---|---|---|---|---|
| I | 1 | --- | 57.549 | --- | --- | 26 |
| M* | 2 | --- | 131.067 | 1432.092 | --- | 25 |
| L | 3 | --- | 187.609 | 1358.574 | --- | 24 |
| S | 4 | --- | 231.125 | 1302.032 | --- | 23 |
| V | 5 | --- | 280.659 | 1258.516 | --- | 22 |
| H | 6 | --- | 349.189 | 1208.982 | --- | 21 |
| G | 7 | --- | 377.699 | 1140.453 | -1.842 | 20 |
| S | 8 | --- | 421.215 | 1111.942 | --- | 19 |
| Q | 9 | --- | 485.245 | 1068.426 | --- | 18 |
| H | 10 | 2.357 | 553.774 | 1004.397 | --- | 17 |
| S | 11 | 0.410 | 597.290 | 935.867 | 1.582 | 16 |
| G | 12 | 1.578 | 625.801 | 892.351 | 4.694 | 15 |
| M* | 13 | 4.904 | 699.319 | 863.841 | --- | 14 |
| I | 14 | -1.814 | 755.861 | 790.323 | -5.216 | 13 |
| V | 15 | --- | 805.395 | 733.781 | -2.487 | 12 |
| N | 16 | --- | 862.416 | 684.247 | 1.031 | 11 |
| D | 17 | 4.890 | 919.930 | 627.225 | --- | 10 |
| T@ | 18 | --- | 1010.437 | 569.712 | 0.743 | 9 |
| G | 19 | --- | 1038.948 | 479.205 | 0.152 | 8 |
| H | 20 | --- | 1107.477 | 450.694 | 1.561 | 7 |
| E | 21 | --- | 1171.998 | 382.164 | --- | 6 |
| T | 22 | --- | 1222.522 | 317.643 | --- | 5 |
| D | 23 | --- | 1280.036 | 267.119 | -40.986 | 4 |
| E | 24 | --- | 1344.557 | 209.606 | --- | 3 |
| N | 25 | --- | 1401.578 | 145.085 | --- | 2 |
| R | 26 | --- | --- | 88.063 | --- | 1 |

+3

+3

| Seq | # | b: Δ Error | b | y | y: Δ Error | +1 |
|---|---|---|---|---|---|---|
| I | 1 | --- | 38.702 | --- | --- | 26 |
| M* | 2 | --- | 87.714 | 955.064 | 1.231 | 25 |
| L | 3 | --- | 125.408 | 906.052 | -1.307 | 24 |
| S | 4 | --- | 154.419 | 868.357 | -2.435 | 23 |
| V | 5 | --- | 187.442 | 839.347 | 1.838 | 22 |
| H | 6 | 15.374 | 233.128 | 806.324 | 1.734 | 21 |
| G | 7 | --- | 252.135 | 760.638 | 1.790 | 20 |
| S | 8 | --- | 281.146 | 741.630 | --- | 19 |
| Q | 9 | --- | 323.832 | 712.620 | --- | 18 |
| H | 10 | --- | 369.519 | 669.934 | 0.823 | 17 |
| S | 11 | --- | 398.529 | 624.247 | --- | 16 |
| G | 12 | --- | 417.536 | 595.237 | --- | 15 |
| M* | 13 | --- | 466.548 | 576.229 | --- | 14 |
| I | 14 | --- | 504.243 | 527.218 | --- | 13 |
| V | 15 | --- | 537.266 | 489.523 | --- | 12 |
| N | 16 | --- | 575.280 | 456.500 | --- | 11 |
| D | 17 | --- | 613.622 | 418.486 | --- | 10 |
| T@ | 18 | --- | 673.960 | 380.144 | --- | 9 |
| G | 19 | --- | 692.967 | 319.806 | --- | 8 |
| H | 20 | --- | 738.654 | 300.798 | --- | 7 |
| E | 21 | --- | 781.668 | 255.112 | --- | 6 |
| T | 22 | 8.679 | 815.351 | 212.098 | --- | 5 |
| D | 23 | --- | 853.693 | 178.415 | --- | 4 |
| E | 24 | --- | 896.707 | 140.073 | --- | 3 |
| N | 25 | --- | 934.721 | 97.059 | --- | 2 |
| R | 26 | --- | --- | 59.045 | --- | 1 |

**AKVEITPNSPR**

z = 2+

ZIKV E protein

S173

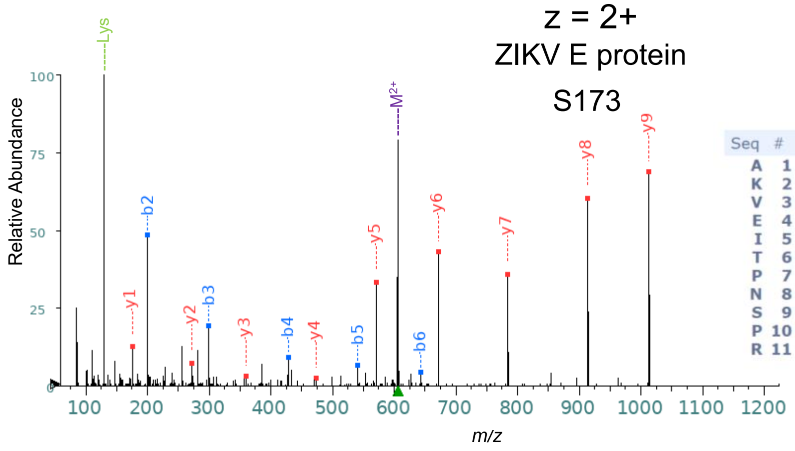


| Seq | # | b: Δ Error | b | y | y: Δ Error | +1 |
|---|---|---|---|---|---|---|
| A | 1 | --- | 72.044 | --- | --- | 11 |
| K | 2 | -1.344 | 200.139 | 1140.637 | --- | 10 |
| V | 3 | -1.744 | 299.208 | 1012.542 | -1.758 | 9 |
| E | 4 | -2.693 | 428.250 | 913.474 | -1.087 | 8 |
| I | 5 | -0.361 | 541.334 | 784.431 | -1.823 | 7 |
| T | 6 | -2.790 | 642.382 | 671.347 | -1.557 | 6 |
| P | 7 | --- | 739.435 | 570.299 | -1.708 | 5 |
| N | 8 | --- | 853.478 | 473.247 | -4.640 | 4 |
| S | 9 | --- | 940.510 | 359.204 | -1.640 | 3 |
| P | 10 | --- | 1037.563 | 272.172 | -2.332 | 2 |
| R | 11 | --- | --- | 175.119 | -1.453 | 1 |

**AKVEITPNS@PR**

z = 2+

ZIKV E protein

pS173

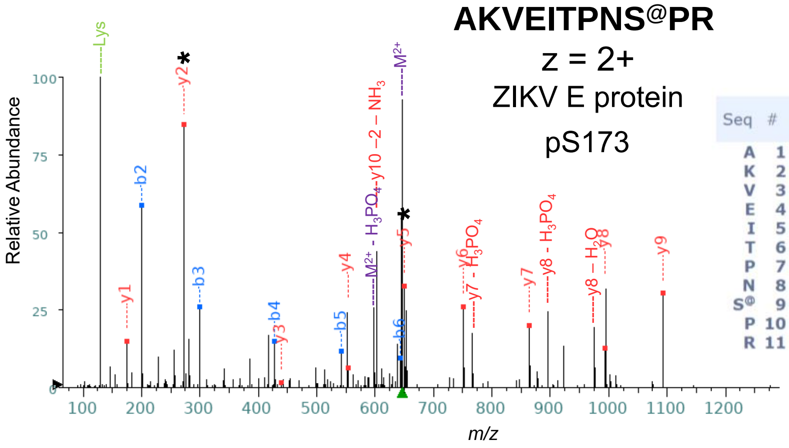


| Seq | # | b: Δ Error | b | y | y: Δ Error | +1 |
|---|---|---|---|---|---|---|
| A | 1 | --- | 72.044 | --- | --- | 11 |
| K | 2 | 1.172 | 200.139 | 1220.603 | --- | 10 |
| V | 3 | 0.398 | 299.208 | 1092.509 | 0.473 | 9 |
| E | 4 | 1.084 | 428.250 | 993.440 | 1.128 | 8 |
| I | 5 | 0.992 | 541.334 | 864.398 | 0.651 | 7 |
| T | 6 | 2.531 | 642.382 | 751.313 | 0.611 | 6 |
| P | 7 | --- | 739.435 | 650.266 | 1.003 | 5 |
| N | 8 | --- | 853.478 | 553.213 | 2.445 | 4 |
| S@ | 9 | --- | 1020.476 | 439.170 | 4.515 | 3 |
| P | 10 | --- | 1117.529 | 272.172 | 0.583 | 2 |
| R | 11 | --- | --- | 175.119 | 1.597 | 1 |

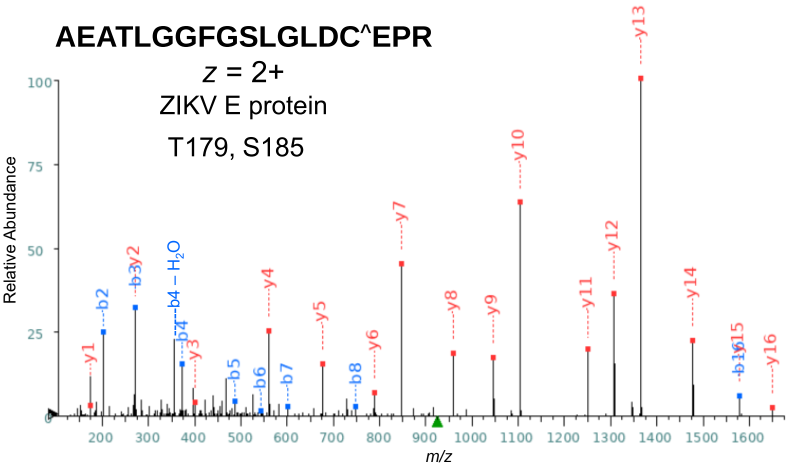


| Seq | # | b: Δ Error | b | y | y: Δ Error | +1 |
|---|---|---|---|---|---|---|
| A | 1 | --- | 72.044 | --- | --- | 18 |
| E | 2 | -0.418 | 201.087 | 1778.838 | --- | 17 |
| A | 3 | -0.830 | 272.124 | 1649.795 | -1.786 | 16 |
| T | 4 | -0.468 | 373.172 | 1578.758 | 0.379 | 15 |
| L | 5 | -1.276 | 486.256 | 1477.710 | 0.040 | 14 |
| G | 6 | -1.835 | 543.277 | 1364.626 | -0.524 | 13 |
| G | 7 | --- | 600.299 | 1307.605 | -0.470 | 12 |
| F | 8 | 1.234 | 747.367 | 1250.583 | -0.703 | 11 |
| G | 9 | --- | 804.389 | 1103.515 | -0.636 | 10 |
| S | 10 | --- | 891.421 | 1046.494 | -0.224 | 9 |
| L | 11 | --- | 1004.505 | 959.461 | -0.387 | 8 |
| G | 12 | --- | 1061.526 | 846.377 | -0.562 | 7 |
| L | 13 | --- | 1174.610 | 789.356 | 0.532 | 6 |
| D | 14 | --- | 1289.637 | 676.272 | 0.016 | 5 |
| C^ | 15 | --- | 1449.668 | 561.245 | -0.695 | 4 |
| E | 16 | --- | 1578.710 | 401.214 | -0.037 | 3 |
| P | 17 | --- | 1675.763 | 272.172 | -0.651 | 2 |
| R | 18 | --- | --- | 175.119 | -1.976 | 1 |

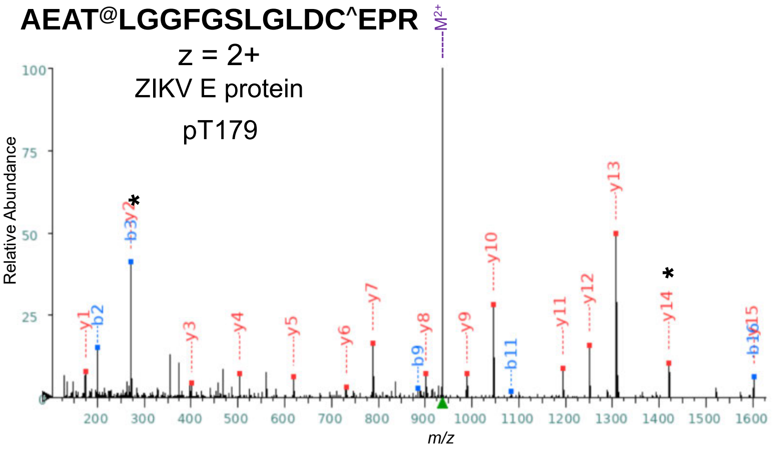


| Seq | # | b: Δ Error | b | y | y: Δ Error | +1 |
|---|---|---|---|---|---|---|
| A | 1 | --- | 72.044 | --- | --- | 18 |
| E | 2 | -1.329 | 201.087 | 1801.783 | --- | 17 |
| A | 3 | -2.400 | 272.124 | 1672.740 | --- | 16 |
| T@ | 4 | --- | 453.138 | 1601.703 | --- | 15 |
| L | 5 | --- | 566.222 | 1420.689 | -1.176 | 14 |
| G | 6 | --- | 623.244 | 1307.605 | -1.216 | 13 |
| G | 7 | --- | 680.265 | 1250.583 | -1.191 | 12 |
| F | 8 | --- | 827.334 | 1193.562 | 0.065 | 11 |
| G | 9 | --- | 884.355 | 1046.494 | -1.391 | 10 |
| S | 10 | --- | 971.387 | 989.472 | -2.540 | 9 |
| L | 11 | --- | 1084.471 | 902.440 | -0.773 | 8 |
| G | 12 | --- | 1141.493 | 789.356 | -1.324 | 7 |
| L | 13 | --- | 1254.577 | 732.335 | 0.378 | 6 |
| D | 14 | --- | 1369.604 | 619.250 | -2.776 | 5 |
| C | 15 | --- | 1472.613 | 504.223 | -2.448 | 4 |
| E | 16 | --- | 1601.655 | 401.214 | -0.646 | 3 |
| P | 17 | --- | 1698.708 | 272.172 | -1.884 | 2 |
| R | 18 | --- | --- | 175.119 | -1.191 | 1 |

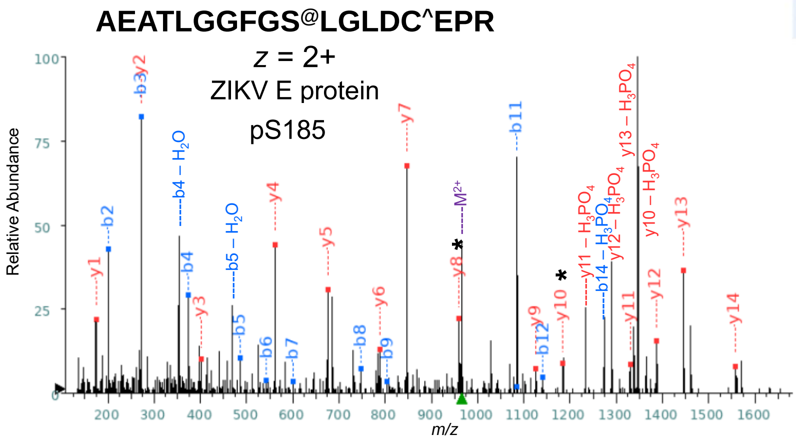


| Seq | # | b: Δ Error | b | y | y: Δ Error | +1 |
|---|---|---|---|---|---|---|
| A | 1 | --- | 72.044 | --- | --- | 18 |
| E | 2 | -1.480 | 201.087 | 1858.804 | --- | 17 |
| A | 3 | -2.176 | 272.124 | 1729.762 | --- | 16 |
| T | 4 | -2.104 | 373.172 | 1658.724 | --- | 15 |
| L | 5 | 0.481 | 486.256 | 1557.677 | 0.102 | 14 |
| G | 6 | 0.637 | 543.277 | 1444.593 | -1.862 | 13 |
| G | 7 | --- | 600.299 | 1387.571 | -1.162 | 12 |
| F | 8 | 1.071 | 747.367 | 1330.550 | --- | 11 |
| G | 9 | --- | 804.389 | 1183.481 | -0.302 | 10 |
| S@ | 10 | --- | 971.387 | 1126.460 | --- | 9 |
| L | 11 | -1.011 | 1084.471 | 959.461 | -2.741 | 8 |
| G | 12 | --- | 1141.493 | 846.377 | -2.364 | 7 |
| L | 13 | --- | 1254.577 | 789.356 | -1.324 | 6 |
| D | 14 | --- | 1369.604 | 676.272 | -1.969 | 5 |
| C^ | 15 | --- | 1529.634 | 561.245 | -1.891 | 4 |
| E | 16 | --- | 1658.677 | 401.214 | -1.178 | 3 |
| P | 17 | --- | 1755.730 | 272.172 | -1.660 | 2 |
| R | 18 | --- | --- | 175.119 | -1.453 | 1 |

**TGLDFSDLYYLTMNNK**

z = 2+

ZIKV E protein

T205

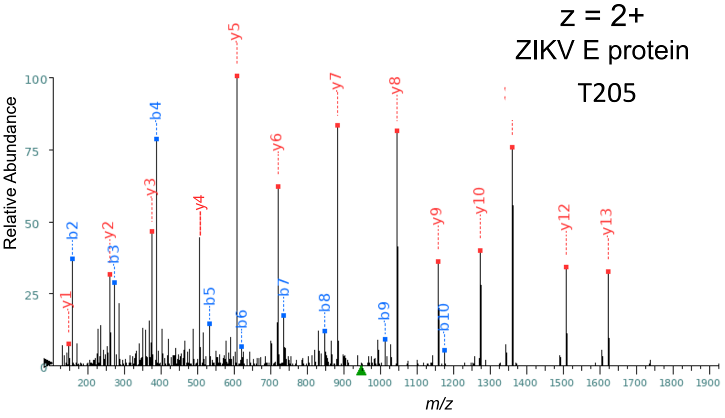


| Seq | # | b: Δ Error | b | y | y: Δ Error | +1 |
| --- | --- | --- | --- | --- | --- | --- |
| T | 1 | --- | 102.055 | --- | --- | 16 |
| G | 2 | -0.685 | 159.076 | 1793.841 | --- | 15 |
| L | 3 | 0.372 | 272.160 | 1736.820 | --- | 14 |
| D | 4 | -0.674 | 387.187 | 1623.736 | -0.585 | 13 |
| F | 5 | 0.381 | 534.256 | 1508.709 | 0.642 | 12 |
| S | 6 | 1.039 | 621.288 | 1361.641 | -0.683 | 11 |
| D | 7 | --- | 736.315 | 1274.609 | 0.168 | 10 |
| L | 8 | --- | 849.399 | 1159.582 | 0.365 | 9 |
| Y | 9 | --- | 1012.462 | 1046.498 | 0.131 | 8 |
| Y | 10 | --- | 1175.526 | 883.434 | 0.402 | 7 |
| L | 11 | --- | 1288.610 | 720.371 | -0.136 | 6 |
| T | 12 | --- | 1389.657 | 607.287 | -0.131 | 5 |
| M | 13 | --- | 1520.698 | 506.239 | --- | 4 |
| N | 14 | 0.993 | 1634.741 | 375.199 | -1.600 | 3 |
| N | 15 | --- | 1748.784 | 261.156 | -0.704 | 2 |
| K | 16 | --- | --- | 147.113 | -0.390 | 1 |

**TGLDFSDLYYLT@MNNK**

z = 2+

ZIKV E protein

pT205

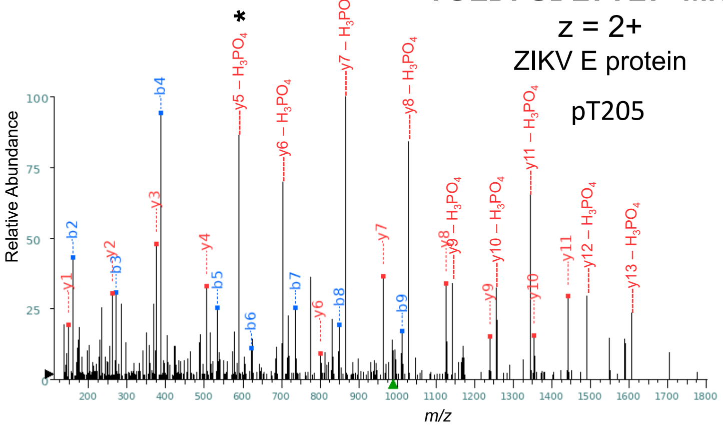


| Seq | # | b: Δ Error | b | y | y: Δ Error | +1 |
| --- | --- | --- | --- | --- | --- | --- |
| T | 1 | --- | 102.055 | --- | --- | 16 |
| G | 2 | -0.397 | 159.076 | 1873.808 | --- | 15 |
| L | 3 | -0.188 | 272.160 | 1816.786 | --- | 14 |
| D | 4 | -0.516 | 387.187 | 1703.702 | --- | 13 |
| F | 5 | 1.523 | 534.256 | 1588.675 | --- | 12 |
| S | 6 | -2.792 | 621.288 | 1441.607 | 1.541 | 11 |
| D | 7 | 1.255 | 736.315 | 1354.575 | -2.201 | 10 |
| L | 8 | 0.779 | 849.399 | 1239.548 | 0.324 | 9 |
| Y | 9 | -1.732 | 1012.462 | 1126.464 | -1.415 | 8 |
| Y | 10 | --- | 1175.526 | 963.401 | -0.414 | 7 |
| L | 11 | --- | 1288.610 | 800.337 | 1.833 | 6 |
| T℗ | 12 | --- | 1469.624 | 687.253 | --- | 5 |
| M | 13 | --- | 1600.664 | 506.239 | 1.913 | 4 |
| N | 14 | --- | 1714.707 | 375.199 | -0.787 | 3 |
| N | 15 | --- | 1828.750 | 261.156 | -0.353 | 2 |
| K | 16 | --- | --- | 147.113 | -0.079 | 1 |

**EWFHDIPLPWHAGADTGTPHWNNK**

z = 3+

ZIKV E protein

T231, T233

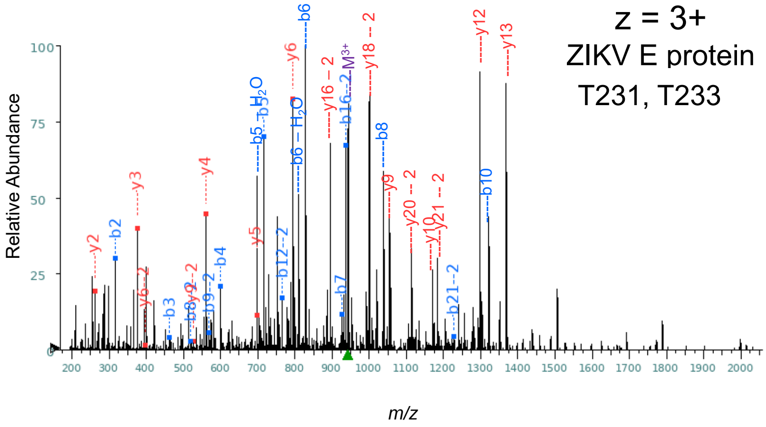


+1

| Seq | # | b: Δ Error | b | y | y: Δ Error | +1 |
|---|---|---|---|---|---|---|
| E | 1 | --- | 130.050 | --- | --- | 24 |
| W | 2 | -5.506 | **316.129** | 2697.275 | --- | 23 |
| F | 3 | -4.930 | **463.198** | 2511.195 | --- | 22 |
| H | 4 | -4.741 | **600.257** | 2364.127 | --- | 21 |
| D | 5 | -4.186 | **715.283** | 2227.068 | --- | 20 |
| I | 6 | --- | 828.368 | 2112.041 | --- | 19 |
| P | 7 | -3.090 | **925.420** | 1998.957 | --- | 18 |
| L | 8 | --- | 1038.504 | 1901.904 | --- | 17 |
| P | 9 | --- | 1135.557 | 1788.820 | --- | 16 |
| W | 10 | --- | 1321.636 | 1691.767 | --- | 15 |
| H | 11 | --- | 1458.695 | 1505.688 | --- | 14 |
| A | 12 | --- | 1529.732 | 1368.629 | --- | 13 |
| G | 13 | --- | 1586.754 | 1297.592 | --- | 12 |
| A | 14 | --- | 1657.791 | 1240.571 | --- | 11 |
| D | 15 | --- | 1772.818 | 1169.533 | --- | 10 |
| T | 16 | --- | 1873.866 | 1054.506 | --- | 9 |
| G | 17 | --- | 1930.887 | 953.459 | --- | 8 |
| T | 18 | --- | 2031.935 | 896.437 | --- | 7 |
| P | 19 | --- | 2128.988 | **795.390** | -3.683 | 6 |
| H | 20 | --- | 2266.046 | **698.337** | -4.065 | 5 |
| W | 21 | --- | 2452.126 | **561.278** | -4.709 | 4 |
| N | 22 | --- | 2566.169 | **375.199** | -5.423 | 3 |
| N | 23 | --- | 2680.212 | **261.156** | -5.495 | 2 |
| K | 24 | --- | --- | 147.113 | --- | 1 |

+2

| Seq | # | b: Δ Error | b | y | y: Δ Error | +1 |
|---|---|---|---|---|---|---|
| E | 1 | --- | 65.529 | --- | --- | 24 |
| W | 2 | --- | 158.568 | 1349.141 | --- | 23 |
| F | 3 | --- | 232.102 | 1256.101 | --- | 22 |
| H | 4 | --- | 300.632 | 1182.567 | --- | 21 |
| D | 5 | --- | 358.145 | 1114.038 | --- | 20 |
| I | 6 | --- | 414.687 | 1056.524 | --- | 19 |
| P | 7 | --- | 463.214 | 999.982 | --- | 18 |
| L | 8 | -4.249 | **519.756** | 951.456 | --- | 17 |
| P | 9 | -4.771 | **568.282** | 894.914 | --- | 16 |
| W | 10 | --- | 661.322 | 846.387 | --- | 15 |
| H | 11 | --- | 729.851 | 753.348 | --- | 14 |
| A | 12 | -1.738 | **765.370** | 684.818 | --- | 13 |
| G | 13 | --- | 793.881 | 649.300 | --- | 12 |
| A | 14 | --- | 829.399 | 620.789 | --- | 11 |
| D | 15 | --- | 886.913 | 585.270 | --- | 10 |
| T | 16 | 0.654 | **937.436** | **527.757** | -2.729 | 9 |
| G | 17 | --- | 965.947 | 477.233 | --- | 8 |
| T | 18 | --- | 1016.471 | 448.722 | --- | 7 |
| P | 19 | --- | 1064.997 | **398.198** | -6.377 | 6 |
| H | 20 | --- | 1133.527 | 349.672 | --- | 5 |
| W | 21 | -1.491 | **1226.567** | 281.143 | --- | 4 |
| N | 22 | --- | 1283.588 | 188.103 | --- | 3 |
| N | 23 | --- | 1340.609 | 131.082 | --- | 2 |
| K | 24 | --- | --- | 74.060 | --- | 1 |

**EWFHDIPLPWHAGADT@GTPHWNNK**

z = 3+

ZIKV E protein

pT231

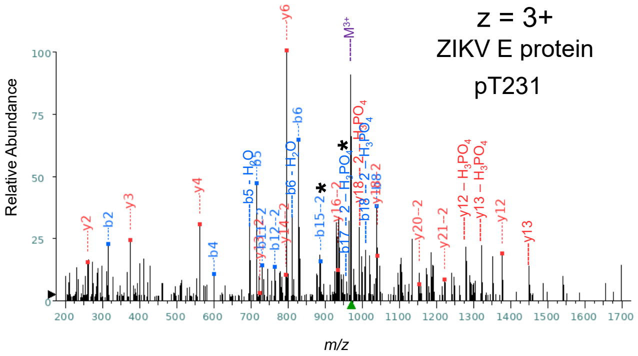


| +1 | | | | | | |
|---|---|---|---|---|---|---|
| Seq | # | b: Δ Error | b | y | y: Δ Error | +1 |
| E | 1 | --- | 130.050 | --- | --- | 24 |
| W | 2 | -1.934 | **316.129** | 2777.241 | --- | 23 |
| F | 3 | --- | 463.198 | 2591.162 | --- | 22 |
| H | 4 | -1.894 | **600.257** | 2444.093 | --- | 21 |
| D | 5 | -2.650 | **715.283** | 2307.034 | --- | 20 |
| I | 6 | -1.942 | **828.368** | 2192.007 | --- | 19 |
| P | 7 | --- | 925.420 | 2078.923 | --- | 18 |
| L | 8 | -0.303 | **1038.504** | 1981.870 | --- | 17 |
| P | 9 | --- | 1135.557 | 1868.786 | --- | 16 |
| W | 10 | --- | 1321.636 | 1771.734 | --- | 15 |
| H | 11 | --- | 1458.695 | 1585.654 | --- | 14 |
| A | 12 | --- | 1529.732 | 1448.595 | --- | 13 |
| G | 13 | --- | 1586.754 | **1377.558** | -0.864 | 12 |
| A | 14 | --- | 1657.791 | 1320.537 | --- | 11 |
| D | 15 | --- | 1772.818 | 1249.500 | --- | 10 |
| T@ | 16 | --- | 1953.832 | 1134.473 | --- | 9 |
| G | 17 | --- | 2010.853 | 953.459 | --- | 8 |
| T | 18 | --- | 2111.901 | 896.437 | --- | 7 |
| P | 19 | --- | 2208.954 | **795.390** | -1.611 | 6 |
| H | 20 | --- | 2346.013 | 698.337 | --- | 5 |
| W | 21 | --- | 2532.092 | **561.278** | -0.576 | 4 |
| N | 22 | --- | 2646.135 | **375.199** | -2.332 | 3 |
| N | 23 | --- | 2760.178 | **261.156** | -1.171 | 2 |
| K | 24 | --- | --- | 147.113 | --- | 1 |

| +2 | | | | | | |
|---|---|---|---|---|---|---|
| Seq | # | b: Δ Error | b | y | y: Δ Error | +1 |
| E | 1 | --- | 65.529 | --- | --- | 24 |
| W | 2 | --- | 158.568 | 1389.124 | --- | 23 |
| F | 3 | --- | 232.102 | 1296.084 | --- | 22 |
| H | 4 | --- | 300.632 | **1222.550** | 2.254 | 21 |
| D | 5 | --- | 358.145 | **1154.021** | 1.257 | 20 |
| I | 6 | --- | 414.687 | 1096.507 | --- | 19 |
| P | 7 | --- | 463.214 | **1039.965** | -2.633 | 18 |
| L | 8 | --- | 519.756 | 991.439 | --- | 17 |
| P | 9 | --- | 568.282 | **934.897** | 0.981 | 16 |
| W | 10 | --- | 661.322 | 886.370 | --- | 15 |
| H | 11 | -0.147 | **729.851** | **793.331** | -0.770 | 14 |
| A | 12 | -1.020 | **765.370** | **724.801** | 1.482 | 13 |
| G | 13 | --- | 793.881 | 689.283 | --- | 12 |
| A | 14 | --- | 829.399 | 660.772 | --- | 11 |
| D | 15 | -1.265 | **886.913** | 625.254 | --- | 10 |
| T@ | 16 | --- | 977.420 | 567.740 | --- | 9 |
| G | 17 | --- | 1005.930 | 477.233 | --- | 8 |
| T | 18 | --- | 1056.454 | 448.722 | --- | 7 |
| P | 19 | --- | 1104.981 | 398.198 | --- | 6 |
| H | 20 | --- | 1173.510 | 349.672 | --- | 5 |
| W | 21 | --- | 1266.550 | 281.143 | --- | 4 |
| N | 22 | --- | 1323.571 | 188.103 | --- | 3 |
| N | 23 | --- | 1380.593 | 131.082 | --- | 2 |
| K | 24 | --- | --- | 74.060 | --- | 1 |

**EWFHDIPLPWHAGADTGT@PHWNNK**

z = 3+

ZIKV E protein

pT233

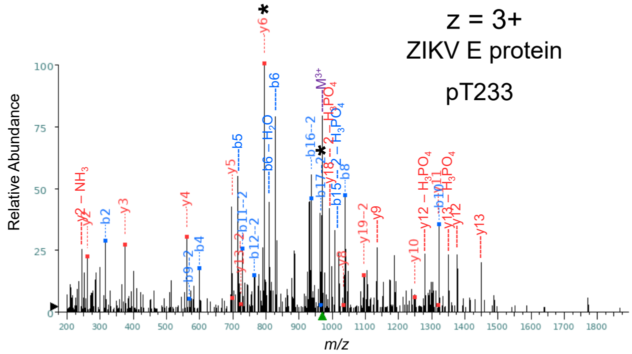


| +1 | | | | | | |
|---|---|---|---|---|---|---|
| Seq | # | b: Δ Error | b | y | y: Δ Error | +1 |
| E | 1 | --- | 130.050 | --- | --- | 24 |
| W | 2 | -4.637 | **316.129** | 2777.241 | --- | 23 |
| F | 3 | --- | 463.198 | 2591.162 | --- | 22 |
| H | 4 | -3.927 | **600.257** | 2444.093 | --- | 21 |
| D | 5 | --- | 715.283 | 2307.034 | --- | 20 |
| I | 6 | --- | 828.368 | 2192.007 | --- | 19 |
| P | 7 | --- | 925.420 | 2078.923 | --- | 18 |
| L | 8 | -1.713 | **1038.504** | 1981.870 | --- | 17 |
| P | 9 | --- | 1135.557 | 1868.786 | --- | 16 |
| W | 10 | -0.235 | **1321.636** | 1771.734 | --- | 15 |
| H | 11 | --- | 1458.695 | 1585.654 | --- | 14 |
| A | 12 | --- | 1529.732 | 1448.595 | --- | 13 |
| G | 13 | --- | 1586.754 | 1377.558 | --- | 12 |
| A | 14 | --- | 1657.791 | **1320.537** | -1.380 | 11 |
| D | 15 | --- | 1772.818 | **1249.500** | 0.695 | 10 |
| T | 16 | --- | 1873.866 | 1134.473 | --- | 9 |
| G | 17 | --- | 1930.887 | **1033.425** | 0.285 | 8 |
| T@ | 18 | --- | 2111.901 | 976.404 | --- | 7 |
| P | 19 | --- | 2208.954 | **795.390** | -3.683 | 6 |
| H | 20 | --- | 2346.013 | **698.337** | 0.218 | 5 |
| W | 21 | --- | 2532.092 | **561.278** | -4.056 | 4 |
| N | 22 | --- | 2646.135 | **375.199** | -4.040 | 3 |
| N | 23 | --- | 2760.178 | **261.156** | -3.976 | 2 |
| K | 24 | --- | --- | 147.113 | --- | 1 |

| +2 | | | | | | |
|---|---|---|---|---|---|---|
| Seq | # | b: Δ Error | b | y | y: Δ Error | +1 |
| E | 1 | --- | 65.529 | --- | --- | 24 |
| W | 2 | --- | 158.568 | 1389.124 | --- | 23 |
| F | 3 | --- | 232.102 | 1296.084 | --- | 22 |
| H | 4 | --- | 300.632 | 1222.550 | --- | 21 |
| D | 5 | --- | 358.145 | 1154.021 | --- | 20 |
| I | 6 | --- | 414.687 | **1096.507** | 1.140 | 19 |
| P | 7 | --- | 463.214 | 1039.965 | --- | 18 |
| L | 8 | --- | 519.756 | 991.439 | --- | 17 |
| P | 9 | 0.599 | **568.282** | 934.897 | --- | 16 |
| W | 10 | --- | 661.322 | 886.370 | --- | 15 |
| H | 11 | -2.656 | **729.851** | 793.331 | --- | 14 |
| A | 12 | -3.812 | **765.370** | **724.801** | -1.633 | 13 |
| G | 13 | --- | 793.881 | 689.283 | --- | 12 |
| A | 14 | --- | 829.399 | 660.772 | --- | 11 |
| D | 15 | --- | 886.913 | 625.254 | --- | 10 |
| T | 16 | 0.263 | **937.436** | 567.740 | --- | 9 |
| G | 17 | 1.593 | **965.947** | 517.216 | --- | 8 |
| T@ | 18 | --- | 1056.454 | 488.705 | --- | 7 |
| P | 19 | --- | 1104.981 | 398.198 | --- | 6 |
| H | 20 | --- | 1173.510 | 349.672 | --- | 5 |
| W | 21 | --- | 1266.550 | 281.143 | --- | 4 |
| N | 22 | --- | 1323.571 | 188.103 | --- | 3 |
| N | 23 | --- | 1380.593 | 131.082 | --- | 2 |
| K | 24 | --- | --- | 74.060 | --- | 1 |

**RQTVVVLGSQEGAVHTALAGALEAEM*DGAK**

z = 3+

ZIKV E protein

T254

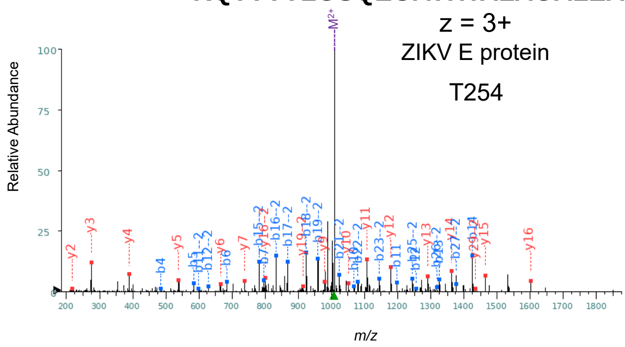

| Seq | # | b: Δ Error | b (+1) | y (+1) | y: Δ Error | +1 |
|---|---|---|---|---|---|---|
| R | 1 | --- | 157.108 | --- | --- | 30 |
| Q | 2 | --- | 285.167 | 2868.441 | --- | 29 |
| T | 3 | --- | 386.215 | 2740.382 | --- | 28 |
| V | 4 | -2.340 | **485.283** | 2639.335 | --- | 27 |
| V | 5 | 2.037 | **584.351** | 2540.266 | --- | 26 |
| V | 6 | 0.501 | **683.420** | 2441.198 | --- | 25 |
| L | 7 | -0.436 | **796.504** | 2342.129 | --- | 24 |
| G | 8 | --- | 853.525 | 2229.045 | --- | 23 |
| S | 9 | --- | 940.557 | 2172.024 | --- | 22 |
| Q | 10 | -2.219 | **1068.616** | 2084.992 | --- | 21 |
| E | 11 | 0.373 | **1197.659** | 1956.933 | --- | 20 |
| G | 12 | -0.601 | **1254.680** | 1827.891 | --- | 19 |
| A | 13 | 1.362 | **1325.717** | 1770.869 | --- | 18 |
| V | 14 | -0.913 | **1424.786** | 1699.832 | --- | 17 |
| H | 15 | --- | 1561.845 | **1600.764** | 1.145 | 16 |
| T | 16 | --- | 1662.892 | **1463.705** | -1.032 | 15 |
| A | 17 | --- | 1733.929 | **1362.657** | 0.377 | 14 |
| L | 18 | --- | 1847.013 | **1291.620** | 2.197 | 13 |
| A | 19 | --- | 1918.050 | **1178.536** | -0.529 | 12 |
| G | 20 | --- | 1975.072 | **1107.499** | 0.103 | 11 |
| A | 21 | --- | 2046.109 | **1050.477** | -1.190 | 10 |
| L | 22 | --- | 2159.193 | **979.440** | -2.206 | 9 |
| E | 23 | --- | 2288.236 | 866.356 | --- | 8 |
| A | 24 | --- | 2359.273 | **737.313** | -0.104 | 7 |
| E | 25 | --- | 2488.315 | **666.276** | -0.017 | 6 |
| M* | 26 | --- | 2635.351 | **537.234** | -0.607 | 5 |
| D | 27 | --- | 2750.378 | **390.198** | -1.542 | 4 |
| G | 28 | --- | 2807.399 | **275.171** | -0.759 | 3 |
| A | 29 | --- | 2878.436 | **218.150** | -1.402 | 2 |
| K | 30 | --- | --- | 147.113 | --- | 1 |

| Seq | # | b: Δ Error | b (+2) | y (+2) | y: Δ Error | +1 |
|---|---|---|---|---|---|---|
| R | 1 | --- | 79.058 | --- | --- | 30 |
| Q | 2 | --- | 143.087 | **1434.724** | 1.262 | 29 |
| T | 3 | --- | 193.611 | 1370.695 | --- | 28 |
| V | 4 | --- | 243.145 | 1320.171 | --- | 27 |
| V | 5 | --- | 292.679 | 1270.637 | --- | 26 |
| V | 6 | --- | 342.214 | 1221.102 | --- | 25 |
| L | 7 | --- | 398.756 | 1171.568 | --- | 24 |
| G | 8 | --- | 427.266 | 1115.026 | --- | 23 |
| S | 9 | --- | 470.782 | 1086.516 | --- | 22 |
| Q | 10 | --- | 534.812 | 1042.999 | --- | 21 |
| E | 11 | -4.781 | **599.333** | 978.970 | --- | 20 |
| G | 12 | 0.410 | **627.844** | **914.449** | 2.009 | 19 |
| A | 13 | --- | 663.362 | 885.938 | --- | 18 |
| V | 14 | --- | 712.896 | 850.420 | --- | 17 |
| H | 15 | -0.068 | **781.426** | **800.885** | -1.264 | 16 |
| T | 16 | -0.327 | **831.950** | 732.356 | --- | 15 |
| A | 17 | 0.880 | **867.468** | 681.832 | --- | 14 |
| L | 18 | -0.207 | **924.010** | 646.314 | --- | 13 |
| A | 19 | 1.897 | **959.529** | 589.772 | --- | 12 |
| G | 20 | --- | 988.040 | 554.253 | --- | 11 |
| A | 21 | -0.360 | **1023.558** | 525.742 | --- | 10 |
| L | 22 | 1.147 | **1080.100** | 490.224 | --- | 9 |
| E | 23 | 0.074 | **1144.621** | 433.682 | --- | 8 |
| A | 24 | --- | 1180.140 | 369.160 | --- | 7 |
| E | 25 | 0.315 | **1244.661** | 333.642 | --- | 6 |
| M* | 26 | 0.576 | **1318.179** | 269.121 | --- | 5 |
| D | 27 | 0.342 | **1375.693** | 195.603 | --- | 4 |
| G | 28 | --- | 1404.203 | 138.089 | --- | 3 |
| A | 29 | --- | 1439.722 | 109.579 | --- | 2 |
| K | 30 | --- | --- | 74.060 | --- | 1 |

**RQT@VVVLGSQEGAVHTALAGALEAEM*DGAK**

z = 3+

ZIKV E protein

pT254

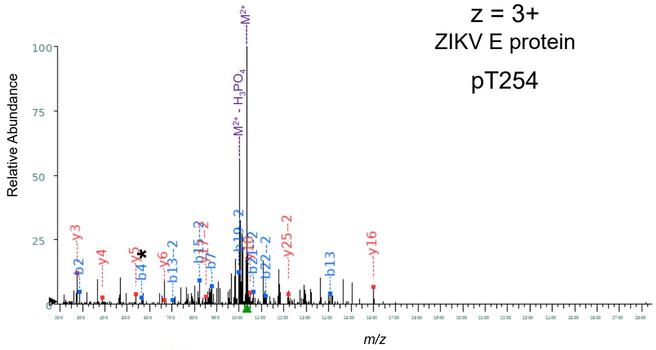

| Seq | # | b: Δ Error | b (+1) | y (+1) | y: Δ Error | +1 |
|---|---|---|---|---|---|---|
| R | 1 | --- | 157.108 | --- | --- | 30 |
| Q | 2 | -3.007 | **285.167** | 2948.407 | --- | 29 |
| T@ | 3 | --- | 466.181 | 2820.349 | --- | 28 |
| V | 4 | -1.724 | **565.249** | 2639.335 | --- | 27 |
| V | 5 | --- | 664.318 | 2540.266 | --- | 26 |
| V | 6 | --- | 763.386 | 2441.198 | --- | 25 |
| L | 7 | -2.789 | **876.470** | 2342.129 | --- | 24 |
| G | 8 | --- | 933.492 | 2229.045 | --- | 23 |
| S | 9 | --- | 1020.524 | 2172.024 | --- | 22 |
| Q | 10 | --- | 1148.582 | 2084.992 | --- | 21 |
| E | 11 | --- | 1277.625 | 1956.933 | --- | 20 |
| G | 12 | --- | 1334.646 | 1827.891 | --- | 19 |
| A | 13 | -1.944 | **1405.684** | 1770.869 | --- | 18 |
| V | 14 | --- | 1504.752 | 1699.832 | --- | 17 |
| H | 15 | --- | 1641.811 | **1600.764** | -1.600 | 16 |
| T | 16 | --- | 1742.859 | 1463.705 | --- | 15 |
| A | 17 | --- | 1813.896 | 1362.657 | --- | 14 |
| L | 18 | --- | 1926.980 | 1291.620 | --- | 13 |
| A | 19 | --- | 1998.017 | 1178.536 | --- | 12 |
| G | 20 | --- | 2055.038 | 1107.499 | --- | 11 |
| A | 21 | --- | 2126.075 | **1050.477** | -1.306 | 10 |
| L | 22 | --- | 2239.159 | 979.440 | --- | 9 |
| E | 23 | --- | 2368.202 | 866.356 | --- | 8 |
| A | 24 | --- | 2439.239 | 737.313 | --- | 7 |
| E | 25 | --- | 2568.282 | **666.276** | -0.108 | 6 |
| M* | 26 | --- | 2715.317 | **537.234** | -1.402 | 5 |
| D | 27 | --- | 2830.344 | **390.198** | -3.732 | 4 |
| G | 28 | --- | 2887.366 | **275.171** | -5.972 | 3 |
| A | 29 | --- | 2958.403 | 218.150 | --- | 2 |
| K | 30 | --- | --- | 147.113 | --- | 1 |

| Seq | # | b: Δ Error | b (+2) | y (+2) | y: Δ Error | +1 |
|---|---|---|---|---|---|---|
| R | 1 | --- | 79.058 | --- | --- | 30 |
| Q | 2 | --- | 143.087 | 1474.707 | --- | 29 |
| T@ | 3 | --- | 233.594 | 1410.678 | --- | 28 |
| V | 4 | --- | 283.128 | 1320.171 | --- | 27 |
| V | 5 | --- | 332.663 | 1270.637 | --- | 26 |
| V | 6 | --- | 382.197 | **1221.102** | 2.349 | 25 |
| L | 7 | --- | 438.739 | 1171.568 | --- | 24 |
| G | 8 | --- | 467.250 | 1115.026 | --- | 23 |
| S | 9 | --- | 510.766 | 1086.516 | --- | 22 |
| Q | 10 | --- | 574.795 | 1042.999 | --- | 21 |
| E | 11 | --- | 639.316 | 978.970 | --- | 20 |
| G | 12 | --- | 667.827 | 914.449 | --- | 19 |
| A | 13 | -1.735 | **703.345** | 885.938 | --- | 18 |
| V | 14 | --- | 752.880 | **850.420** | 0.787 | 17 |
| H | 15 | -3.496 | **821.409** | 800.885 | --- | 16 |
| T | 16 | --- | 871.933 | 732.356 | --- | 15 |
| A | 17 | --- | 907.451 | 681.832 | --- | 14 |
| L | 18 | --- | 963.993 | 646.314 | --- | 13 |
| A | 19 | -1.609 | **999.512** | 589.772 | --- | 12 |
| G | 20 | --- | 1028.023 | 554.253 | --- | 11 |
| A | 21 | -1.447 | **1063.541** | 525.742 | --- | 10 |
| L | 22 | -2.173 | **1120.083** | 490.224 | --- | 9 |
| E | 23 | --- | 1184.605 | 433.682 | --- | 8 |
| A | 24 | --- | 1220.123 | 369.160 | --- | 7 |
| E | 25 | --- | 1284.645 | 333.642 | --- | 6 |
| M* | 26 | --- | 1358.162 | 269.121 | --- | 5 |
| D | 27 | --- | 1415.676 | 195.603 | --- | 4 |
| G | 28 | --- | 1444.186 | 138.089 | --- | 3 |
| A | 29 | --- | 1479.705 | 109.579 | --- | 2 |
| K | 30 | --- | --- | 74.060 | --- | 1 |

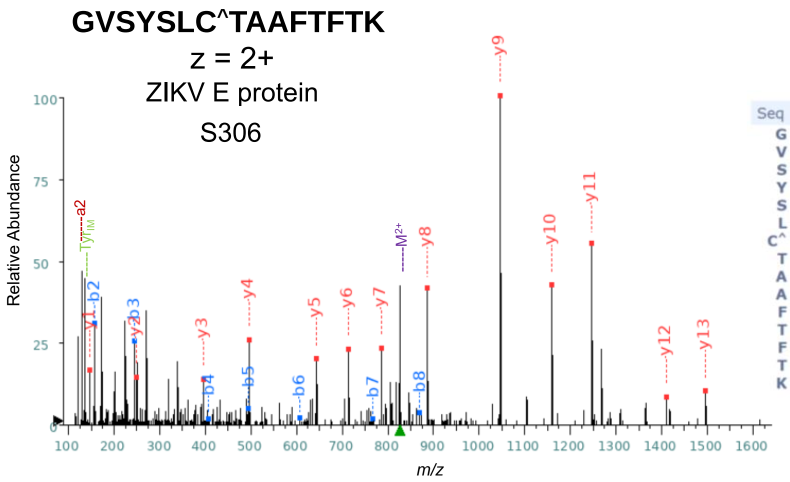


| Seq | # | b: Δ Error | b | y | y: Δ Error | +1 |
|---|---|---|---|---|---|---|
| G | 1 | --- | 58.029 | --- | --- | 15 |
| V | 2 | -0.492 | 157.097 | 1595.777 | --- | 14 |
| S | 3 | -1.192 | 244.129 | 1496.709 | 0.810 | 13 |
| Y | 4 | 4.333 | 407.193 | 1409.677 | --- | 12 |
| S | 5 | 0.822 | 494.225 | 1246.614 | -0.382 | 11 |
| L | 6 | --- | 607.309 | 1159.582 | -0.161 | 10 |
| C^ | 7 | --- | 767.339 | 1046.498 | -0.452 | 9 |
| T | 8 | --- | 868.387 | 886.467 | -0.386 | 8 |
| A | 9 | --- | 939.424 | 785.419 | 0.742 | 7 |
| A | 10 | --- | 1010.461 | 714.382 | -0.545 | 6 |
| F | 11 | --- | 1157.530 | 643.345 | -0.408 | 5 |
| T | 12 | --- | 1258.577 | 496.277 | -0.235 | 4 |
| F | 13 | --- | 1405.646 | 395.229 | -1.196 | 3 |
| T | 14 | --- | 1506.693 | 248.160 | -1.686 | 2 |
| K | 15 | --- | --- | 147.113 | -1.219 | 1 |

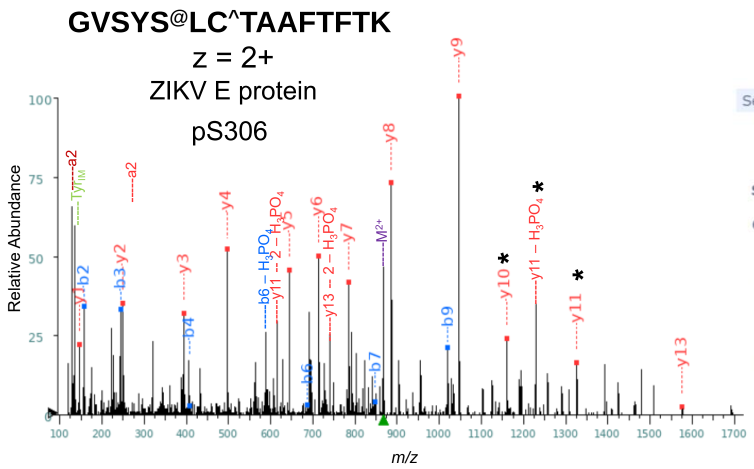


| Seq | # | b: Δ Error | b | y | y: Δ Error | +1 |
|---|---|---|---|---|---|---|
| G | 1 | --- | 58.029 | --- | --- | 15 |
| V | 2 | -0.880 | 157.097 | 1675.744 | --- | 14 |
| S | 3 | -1.317 | 244.129 | 1576.675 | -0.097 | 13 |
| Y | 4 | 0.885 | 407.193 | 1489.643 | --- | 12 |
| S@ | 5 | --- | 574.191 | 1326.580 | -0.560 | 11 |
| L | 6 | --- | 687.275 | 1159.582 | -0.371 | 10 |
| C^ | 7 | --- | 847.306 | 1046.498 | -0.686 | 9 |
| T | 8 | --- | 948.353 | 886.467 | -0.593 | 8 |
| A | 9 | --- | 1019.390 | 785.419 | -1.278 | 7 |
| A | 10 | --- | 1090.427 | 714.382 | -0.459 | 6 |
| F | 11 | --- | 1237.496 | 643.345 | -0.693 | 5 |
| T | 12 | --- | 1338.544 | 496.277 | -1.096 | 4 |
| F | 13 | --- | 1485.612 | 395.229 | -1.274 | 3 |
| T | 14 | --- | 1586.660 | 248.160 | -1.624 | 2 |
| K | 15 | --- | --- | 147.113 | -1.012 | 1 |

**IPAETLHGTVTVEVQYAGTDGPC#KVPAQMAVDM*QTLTPVGR**

*z* = 4+

ZIKV E protein

T335

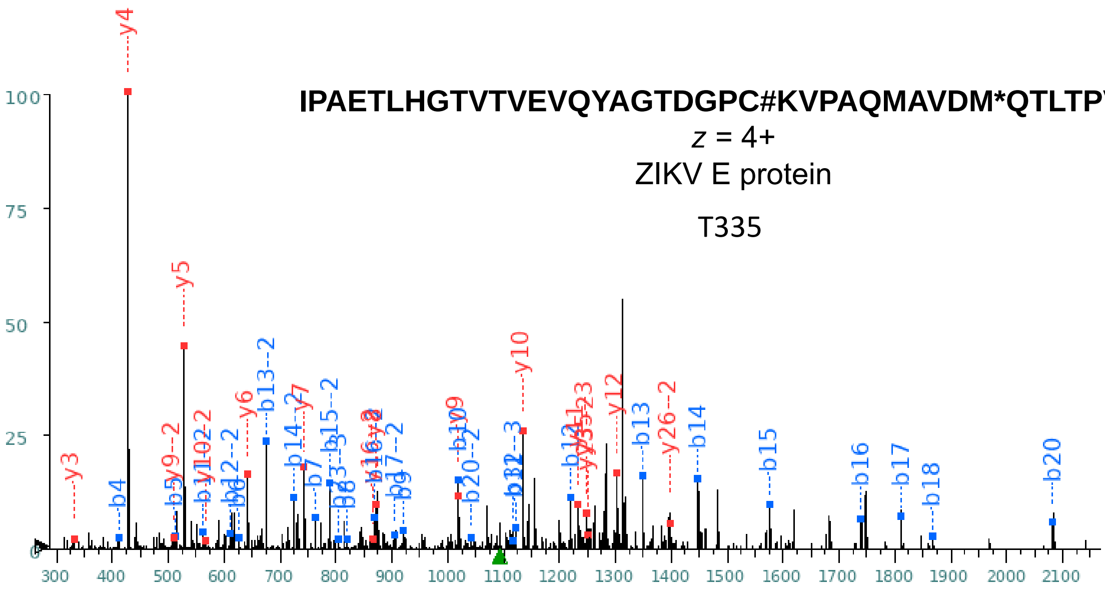


| | | | +1 | | | |
|---|---|---|---|---|---|---|
| Seq | # | b: Δ Error | b | y | y: Δ Error | +1 |
| I | 1 | --- | 114.091 | --- | --- | 41 |
| P | 2 | --- | 211.144 | 4255.083 | --- | 40 |
| A | 3 | --- | 282.181 | 4158.030 | --- | 39 |
| E | 4 | 17.378 | **411.224** | 4086.993 | --- | 38 |
| T | 5 | -2.511 | **512.271** | 3957.951 | --- | 37 |
| L | 6 | -1.404 | **625.356** | 3856.903 | --- | 36 |
| H | 7 | -0.848 | **762.414** | 3743.819 | --- | 35 |
| G | 8 | -2.552 | **819.436** | 3606.760 | --- | 34 |
| T | 9 | 3.951 | **920.484** | 3549.739 | --- | 33 |
| V | 10 | -4.868 | **1019.552** | 3448.691 | --- | 32 |
| T | 11 | -0.081 | **1120.600** | 3349.623 | --- | 31 |
| V | 12 | 5.286 | **1219.668** | 3248.575 | --- | 30 |
| E | 13 | -0.281 | **1348.711** | 3149.506 | --- | 29 |
| V | 14 | 0.206 | **1447.779** | 3020.464 | --- | 28 |
| Q | 15 | 3.143 | **1575.838** | 2921.395 | --- | 27 |
| Y | 16 | 0.757 | **1738.901** | 2793.337 | --- | 26 |
| A | 17 | -0.758 | **1809.938** | 2630.274 | --- | 25 |
| G | 18 | -2.620 | **1866.960** | 2559.236 | --- | 24 |
| T | 19 | --- | 1968.007 | 2502.215 | --- | 23 |
| D | 20 | -4.359 | **2083.034** | 2401.167 | --- | 22 |
| G | 21 | --- | 2140.056 | 2286.140 | --- | 21 |
| P | 22 | --- | 2237.108 | 2229.119 | --- | 20 |
| C# | 23 | --- | 2411.155 | 2132.066 | --- | 19 |
| K | 24 | --- | 2539.250 | 1958.020 | --- | 18 |
| V | 25 | --- | 2638.318 | 1829.925 | --- | 17 |
| P | 26 | --- | 2735.371 | 1730.856 | --- | 16 |
| A | 27 | --- | 2806.408 | 1633.804 | --- | 15 |
| Q | 28 | --- | 2934.467 | 1562.767 | --- | 14 |
| M | 29 | --- | 3065.507 | 1434.708 | --- | 13 |
| A | 30 | --- | 3136.544 | **1303.667** | -9.350 | 12 |
| V | 31 | --- | 3235.613 | **1232.630** | 0.216 | 11 |
| D | 32 | --- | 3350.640 | **1133.562** | 0.283 | 10 |
| M* | 33 | --- | 3497.675 | **1018.535** | -0.498 | 9 |
| Q | 34 | --- | 3625.734 | **871.500** | 5.650 | 8 |
| T | 35 | --- | 3726.781 | **743.441** | -0.295 | 7 |
| L | 36 | --- | 3839.865 | **642.393** | -0.135 | 6 |
| T | 37 | --- | 3940.913 | **529.309** | -0.129 | 5 |
| P | 38 | --- | 4037.966 | **428.262** | -0.421 | 4 |
| V | 39 | --- | 4137.034 | **331.209** | -1.746 | 3 |
| G | 40 | --- | 4194.056 | 232.140 | --- | 2 |
| R | 41 | --- | --- | 175.119 | --- | 1 |

| | | | +2 | | | |
|---|---|---|---|---|---|---|
| Seq | # | b: Δ Error | b | y | y: Δ Error | +1 |
| I | 1 | --- | 57.549 | --- | --- | 41 |
| P | 2 | --- | 106.076 | 2128.045 | --- | 40 |
| A | 3 | --- | 141.594 | 2079.519 | --- | 39 |
| E | 4 | --- | 206.116 | 2044.000 | --- | 38 |
| T | 5 | --- | 256.639 | 1979.479 | --- | 37 |
| L | 6 | --- | 313.181 | 1928.955 | --- | 36 |
| H | 7 | --- | 381.711 | 1872.413 | --- | 35 |
| G | 8 | --- | 410.222 | 1803.884 | --- | 34 |
| T | 9 | --- | 460.745 | 1775.373 | --- | 33 |
| V | 10 | --- | 510.280 | 1724.849 | --- | 32 |
| T | 11 | 1.051 | **560.803** | 1675.315 | --- | 31 |
| V | 12 | 2.721 | **610.338** | 1624.791 | --- | 30 |
| E | 13 | -0.336 | **674.859** | 1575.257 | --- | 29 |
| V | 14 | 1.587 | **724.393** | 1510.736 | --- | 28 |
| Q | 15 | 0.153 | **788.422** | 1461.201 | --- | 27 |
| Y | 16 | -0.408 | **869.954** | **1397.172** | 5.638 | 26 |
| A | 17 | 1.560 | **905.473** | 1315.640 | --- | 25 |
| G | 18 | --- | 933.983 | 1280.122 | --- | 24 |
| T | 19 | --- | 984.507 | **1251.611** | -11.718 | 23 |
| D | 20 | -0.233 | **1042.021** | 1201.087 | --- | 22 |
| G | 21 | --- | 1070.531 | 1143.574 | --- | 21 |
| P | 22 | --- | 1119.058 | 1115.063 | --- | 20 |
| C# | 23 | --- | 1206.081 | 1066.537 | --- | 19 |
| K | 24 | --- | 1270.128 | 979.514 | --- | 18 |
| V | 25 | --- | 1319.663 | 915.466 | --- | 17 |
| P | 26 | --- | 1368.189 | **865.932** | -12.141 | 16 |
| A | 27 | --- | 1403.708 | 817.405 | --- | 15 |
| Q | 28 | --- | 1467.737 | 781.887 | --- | 14 |
| M | 29 | --- | 1533.257 | 717.858 | --- | 13 |
| A | 30 | --- | 1568.776 | 652.337 | --- | 12 |
| V | 31 | --- | 1618.310 | 616.819 | --- | 11 |
| D | 32 | --- | 1675.823 | **567.285** | -1.504 | 10 |
| M* | 33 | --- | 1749.341 | **509.771** | -0.689 | 9 |
| Q | 34 | --- | 1813.370 | 436.253 | --- | 8 |
| T | 35 | --- | 1863.894 | 372.224 | --- | 7 |
| L | 36 | --- | 1920.436 | 321.700 | --- | 6 |
| T | 37 | --- | 1970.960 | 265.158 | --- | 5 |
| P | 38 | --- | 2019.486 | 214.634 | --- | 4 |
| V | 39 | --- | 2069.021 | 166.108 | --- | 3 |
| G | 40 | --- | 2097.531 | 116.574 | --- | 2 |
| R | 41 | --- | --- | 88.063 | --- | 1 |

**IPAETLHGTVTVEVQYAGT@DGPCKVPAQMAVDM*QTLTPVGR**

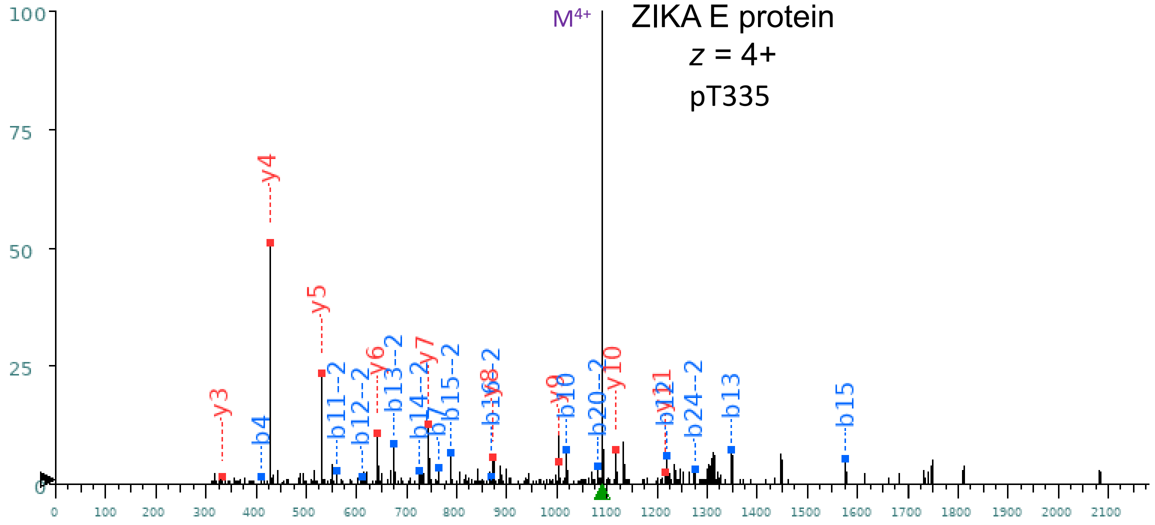


| +1 | | | | | | |
|---|---|---|---|---|---|---|
| Seq | # | b: Δ Error | b | y | y: Δ Error | +1 |
| I | 1 | --- | 114.091 | --- | --- | 41 |
| P | 2 | --- | 211.144 | 4248.017 | --- | 40 |
| A | 3 | --- | 282.181 | 4150.965 | --- | 39 |
| E | 4 | -0.210 | **411.224** | 4079.928 | --- | 38 |
| T | 5 | --- | 512.271 | 3950.885 | --- | 37 |
| L | 6 | --- | 625.356 | 3849.837 | --- | 36 |
| H | 7 | -2.289 | **762.414** | 3736.753 | --- | 35 |
| G | 8 | --- | 819.436 | 3599.694 | --- | 34 |
| T | 9 | --- | 920.484 | 3542.673 | --- | 33 |
| V | 10 | 1.418 | **1019.552** | 3441.625 | --- | 32 |
| T | 11 | --- | 1120.600 | 3342.557 | --- | 31 |
| V | 12 | 3.284 | **1219.668** | 3241.509 | --- | 30 |
| E | 13 | 9.222 | **1348.711** | 3142.441 | --- | 29 |
| V | 14 | --- | 1447.779 | 3013.398 | --- | 28 |
| Q | 15 | 6.939 | **1575.838** | 2914.330 | --- | 27 |
| Y | 16 | --- | 1738.901 | 2786.271 | --- | 26 |
| A | 17 | --- | 1809.938 | 2623.208 | --- | 25 |
| G | 18 | --- | 1866.960 | 2552.171 | --- | 24 |
| T@ | 19 | --- | 2047.974 | 2495.149 | --- | 23 |
| D | 20 | --- | 2163.001 | 2314.135 | --- | 22 |
| G | 21 | --- | 2220.022 | 2199.108 | --- | 21 |
| P | 22 | --- | 2317.075 | 2142.087 | --- | 20 |
| C | 23 | --- | 2420.084 | 2045.034 | --- | 19 |
| K | 24 | --- | 2548.179 | 1942.025 | --- | 18 |
| V | 25 | --- | 2647.247 | 1813.930 | --- | 17 |
| P | 26 | --- | 2744.300 | 1714.861 | --- | 16 |
| A | 27 | --- | 2815.337 | 1617.809 | --- | 15 |
| Q | 28 | --- | 2943.396 | 1546.772 | --- | 14 |
| M | 29 | --- | 3074.436 | 1418.713 | --- | 13 |
| A | 30 | --- | 3145.473 | 1287.673 | --- | 12 |
| V | 31 | --- | 3244.542 | **1216.635** | -8.275 | 11 |
| D | 32 | --- | 3359.569 | **1117.567** | 3.055 | 10 |
| M | 33 | --- | 3490.609 | **1002.540** | 8.790 | 9 |
| Q | 34 | --- | 3618.668 | **871.500** | 8.381 | 8 |
| T | 35 | --- | 3719.715 | **743.441** | 1.840 | 7 |
| L | 36 | --- | 3832.800 | **642.393** | 1.670 | 6 |
| T | 37 | --- | 3933.847 | **529.309** | 1.831 | 5 |
| P | 38 | --- | 4030.900 | **428.262** | 1.503 | 4 |
| V | 39 | --- | 4129.968 | **331.209** | 0.097 | 3 |
| G | 40 | --- | 4186.990 | 232.140 | --- | 2 |
| R | 41 | --- | --- | 175.119 | --- | 1 |

| +2 | | | | | | |
|---|---|---|---|---|---|---|
| Seq | # | b: Δ Error | b | y | y: Δ Error | +1 |
| I | 1 | --- | 57.549 | --- | --- | 41 |
| P | 2 | --- | 106.076 | 2124.512 | --- | 40 |
| A | 3 | --- | 141.594 | 2075.986 | --- | 39 |
| E | 4 | --- | 206.116 | 2040.467 | --- | 38 |
| T | 5 | --- | 256.639 | 1975.946 | --- | 37 |
| L | 6 | --- | 313.181 | 1925.422 | --- | 36 |
| H | 7 | --- | 381.711 | 1868.880 | --- | 35 |
| G | 8 | --- | 410.222 | 1800.351 | --- | 34 |
| T | 9 | --- | 460.745 | 1771.840 | --- | 33 |
| V | 10 | --- | 510.280 | 1721.316 | --- | 32 |
| T | 11 | -0.582 | **560.803** | 1671.782 | --- | 31 |
| V | 12 | -3.379 | **610.338** | 1621.258 | --- | 30 |
| E | 13 | 1.744 | **674.859** | 1571.724 | --- | 29 |
| V | 14 | -1.193 | **724.393** | 1507.203 | --- | 28 |
| Q | 15 | -2.634 | **788.422** | 1457.668 | --- | 27 |
| Y | 16 | -3.004 | **869.954** | 1393.639 | --- | 26 |
| A | 17 | --- | 905.473 | 1312.108 | --- | 25 |
| G | 18 | --- | 933.983 | 1276.589 | --- | 24 |
| T@ | 19 | --- | 1024.490 | 1248.078 | --- | 23 |
| D | 20 | 9.919 | **1082.004** | 1157.571 | --- | 22 |
| G | 21 | --- | 1110.515 | 1100.058 | --- | 21 |
| P | 22 | --- | 1159.041 | 1071.547 | --- | 20 |
| C | 23 | --- | 1210.546 | 1023.021 | --- | 19 |
| K | 24 | 7.115 | **1274.593** | 971.516 | --- | 18 |
| V | 25 | --- | 1324.127 | 907.469 | --- | 17 |
| P | 26 | --- | 1372.654 | 857.934 | --- | 16 |
| A | 27 | --- | 1408.172 | 809.408 | --- | 15 |
| Q | 28 | --- | 1472.202 | 773.889 | --- | 14 |
| M | 29 | --- | 1537.722 | 709.860 | --- | 13 |
| A | 30 | --- | 1573.240 | 644.340 | --- | 12 |
| V | 31 | --- | 1622.775 | 608.821 | --- | 11 |
| D | 32 | --- | 1680.288 | 559.287 | --- | 10 |
| M | 33 | --- | 1745.808 | 501.774 | --- | 9 |
| Q | 34 | --- | 1809.838 | 436.253 | --- | 8 |
| T | 35 | --- | 1860.361 | 372.224 | --- | 7 |
| L | 36 | --- | 1916.903 | 321.700 | --- | 6 |
| T | 37 | --- | 1967.427 | 265.158 | --- | 5 |
| P | 38 | --- | 2015.954 | 214.634 | --- | 4 |
| V | 39 | --- | 2065.488 | 166.108 | --- | 3 |
| G | 40 | --- | 2093.999 | 116.574 | --- | 2 |
| R | 41 | --- | --- | 88.063 | --- | 1 |

#### VPAQMAVDM*QTLTPVGR

z = 2+

ZIKV E protein

T351, T353

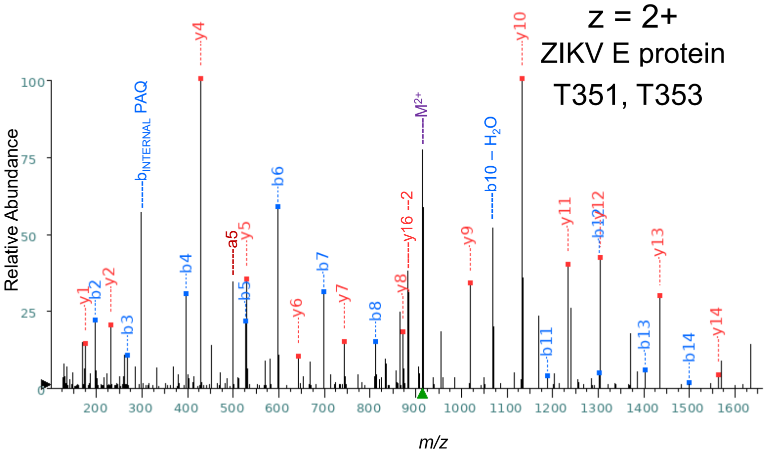


| Seq | # | b: Δ Error | b | y | y: Δ Error | +1 |
|---|---|---|---|---|---|---|
| V | 1 | --- | 100.076 | --- | --- | 17 |
| P | 2 | -0.801 | 197.128 | 1730.856 | --- | 16 |
| A | 3 | -0.150 | 268.166 | 1633.804 | --- | 15 |
| Q | 4 | -0.369 | 396.224 | 1562.767 | -0.263 | 14 |
| M | 5 | 0.036 | 527.265 | 1434.708 | 0.298 | 13 |
| A | 6 | -0.588 | 598.302 | 1303.667 | 1.512 | 12 |
| V | 7 | -1.370 | 697.370 | 1232.630 | 0.018 | 11 |
| D | 8 | 1.496 | 812.397 | 1133.562 | -0.471 | 10 |
| M* | 9 | --- | 959.432 | 1018.535 | 0.102 | 9 |
| Q | 10 | --- | 1087.491 | 871.500 | 0.047 | 8 |
| T | 11 | -1.899 | 1188.539 | 743.441 | 0.937 | 7 |
| L | 12 | 1.769 | 1301.623 | 642.393 | -0.040 | 6 |
| T | 13 | 1.678 | 1402.670 | 529.309 | 0.217 | 5 |
| P | 14 | 1.875 | 1499.723 | 428.262 | -0.849 | 4 |
| V | 15 | --- | 1598.792 | 331.209 | --- | 3 |
| G | 16 | --- | 1655.813 | 232.140 | -0.678 | 2 |
| R | 17 | --- | --- | 175.119 | 0.638 | 1 |

#### VPAQMAVDMQT@LTPVGR

z = 2+

ZIKV E protein

pT351

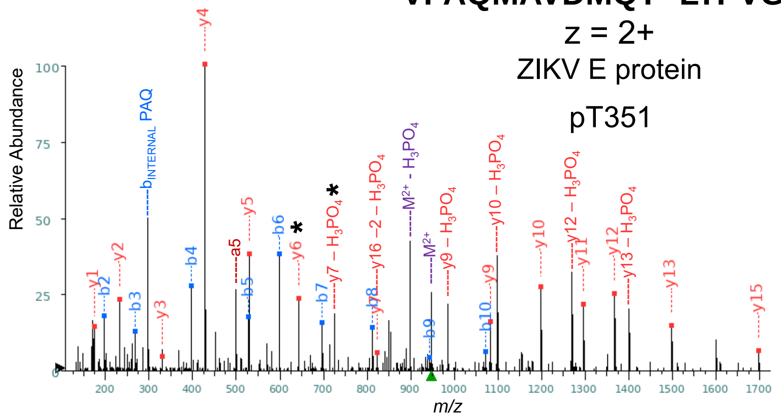


| Seq | # | b: Δ Error | b | y | y: Δ Error | +1 |
|---|---|---|---|---|---|---|
| V | 1 | --- | 100.076 | --- | --- | 17 |
| P | 2 | -0.878 | 197.128 | 1794.828 | --- | 16 |
| A | 3 | -1.288 | 268.166 | 1697.775 | -1.168 | 15 |
| Q | 4 | -0.754 | 396.224 | 1626.738 | --- | 14 |
| M | 5 | -1.006 | 527.265 | 1498.679 | -0.191 | 13 |
| A | 6 | -0.180 | 598.302 | 1367.639 | 1.545 | 12 |
| V | 7 | 0.993 | 697.370 | 1296.602 | 0.880 | 11 |
| D | 8 | 0.669 | 812.397 | 1197.533 | 0.081 | 10 |
| M | 9 | -1.449 | 943.438 | 1082.506 | 0.170 | 9 |
| Q | 10 | 2.214 | 1071.496 | 951.466 | --- | 8 |
| T@ | 11 | --- | 1252.510 | 823.407 | -0.811 | 7 |
| L | 12 | --- | 1365.594 | 642.393 | -0.135 | 6 |
| T | 13 | --- | 1466.642 | 529.309 | -0.706 | 5 |
| P | 14 | --- | 1563.695 | 428.262 | -0.991 | 4 |
| V | 15 | --- | 1662.763 | 331.209 | -1.838 | 3 |
| G | 16 | --- | 1719.785 | 232.140 | -0.941 | 2 |
| R | 17 | --- | --- | 175.119 | -0.494 | 1 |

#### VPAQMAVDM*QTLT@PVGR

z = 2+

ZIKV E protein

pT353

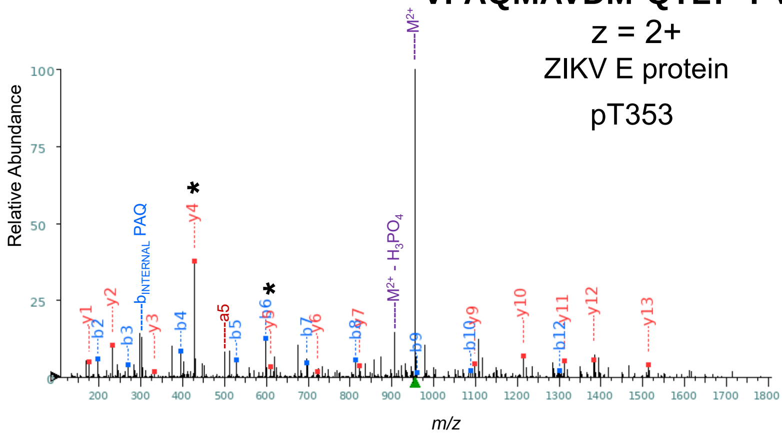


| Seq | # | b: Δ Error | b | y | y: Δ Error | +1 |
|---|---|---|---|---|---|---|
| V | 1 | --- | 100.076 | --- | --- | 17 |
| P | 2 | -0.491 | 197.128 | 1810.823 | --- | 16 |
| A | 3 | -0.605 | 268.166 | 1713.770 | --- | 15 |
| Q | 4 | -0.446 | 396.224 | 1642.733 | --- | 14 |
| M | 5 | -0.312 | 527.265 | 1514.674 | -0.619 | 13 |
| A | 6 | 0.024 | 598.302 | 1383.634 | 0.262 | 12 |
| V | 7 | 4.231 | 697.370 | 1312.597 | -1.116 | 11 |
| D | 8 | 0.218 | 812.397 | 1213.528 | 0.247 | 10 |
| M* | 9 | -0.068 | 959.432 | 1098.501 | 0.686 | 9 |
| Q | 10 | 1.695 | 1087.491 | 951.466 | --- | 8 |
| T | 11 | --- | 1188.539 | 823.407 | 1.042 | 7 |
| L | 12 | -0.763 | 1301.623 | 722.360 | 1.370 | 6 |
| T@ | 13 | --- | 1482.637 | 609.276 | -2.452 | 5 |
| P | 14 | --- | 1579.690 | 428.262 | -0.564 | 4 |
| V | 15 | --- | 1678.758 | 331.209 | -1.285 | 3 |
| G | 16 | --- | 1735.779 | 232.140 | -0.613 | 2 |
| R | 17 | --- | --- | 175.119 | 0.203 | 1 |

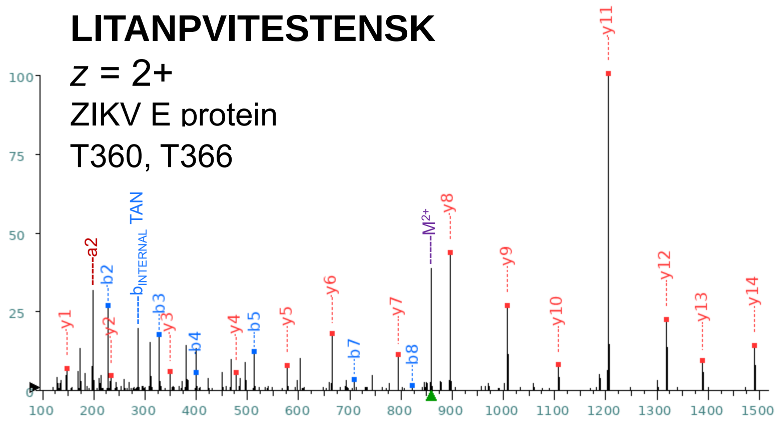


| Seq | # | b: Δ Error | b | y | y: Δ Error | +1 |
|---|---|---|---|---|---|---|
| L | 1 | --- | 114.091 | --- | --- | 16 |
| I | 2 | -1.160 | 227.175 | 1603.817 | --- | 15 |
| T | 3 | -0.927 | 328.223 | 1490.733 | 1.790 | 14 |
| A | 4 | -1.384 | 399.260 | 1389.686 | -0.049 | 13 |
| N | 5 | 0.193 | 513.303 | 1318.649 | 0.044 | 12 |
| P | 6 | --- | 610.356 | 1204.606 | 0.116 | 11 |
| V | 7 | -0.065 | 709.424 | 1107.553 | -1.942 | 10 |
| I | 8 | 0.812 | 822.508 | 1008.484 | -0.142 | 9 |
| T | 9 | --- | 923.556 | 895.400 | -0.889 | 8 |
| E | 10 | --- | 1052.599 | 794.353 | -2.526 | 7 |
| S | 11 | --- | 1139.631 | 665.310 | -0.737 | 6 |
| T | 12 | --- | 1240.678 | 578.278 | -0.346 | 5 |
| E | 13 | --- | 1369.721 | 477.230 | 2.799 | 4 |
| N | 14 | --- | 1483.764 | 348.188 | -2.764 | 3 |
| S | 15 | --- | 1570.796 | 234.145 | 0.015 | 2 |
| K | 16 | --- | --- | 147.113 | -1.738 | 1 |

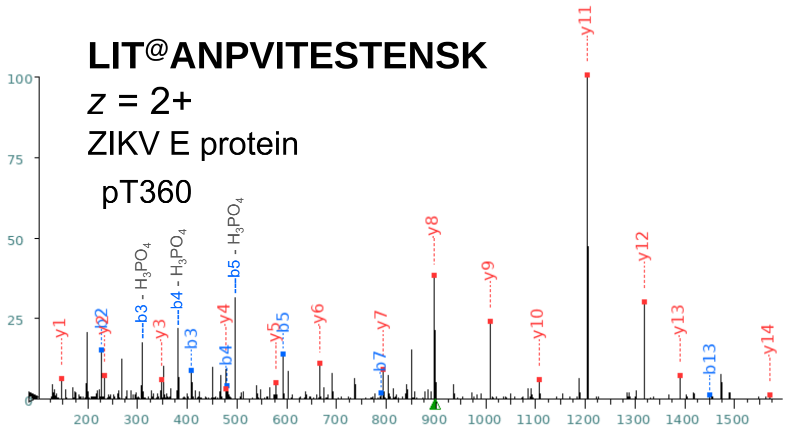


| Seq | # | b: Δ Error | b | y | y: Δ Error | +1 |
|---|---|---|---|---|---|---|
| L | 1 | --- | 114.091 | --- | --- | 16 |
| I | 2 | -1.026 | 227.175 | 1683.784 | --- | 15 |
| T@ | 3 | -0.425 | 408.189 | 1570.700 | -1.035 | 14 |
| A | 4 | 0.521 | 479.227 | 1389.686 | 0.302 | 13 |
| N | 5 | -0.899 | 593.269 | 1318.649 | -1.159 | 12 |
| P | 6 | --- | 690.322 | 1204.606 | -0.695 | 11 |
| V | 7 | 0.069 | 789.391 | 1107.553 | -0.840 | 10 |
| I | 8 | --- | 902.475 | 1008.484 | -1.050 | 9 |
| T | 9 | --- | 1003.522 | 895.400 | -0.957 | 8 |
| E | 10 | --- | 1132.565 | 794.353 | -0.067 | 7 |
| S | 11 | --- | 1219.597 | 665.310 | -1.562 | 6 |
| T | 12 | --- | 1320.645 | 578.278 | 0.499 | 5 |
| E | 13 | 0.750 | 1449.687 | 477.230 | 2.288 | 4 |
| N | 14 | --- | 1563.730 | 348.188 | -0.835 | 3 |
| S | 15 | --- | 1650.762 | 234.145 | -0.441 | 2 |
| K | 16 | --- | --- | 147.113 | -0.805 | 1 |

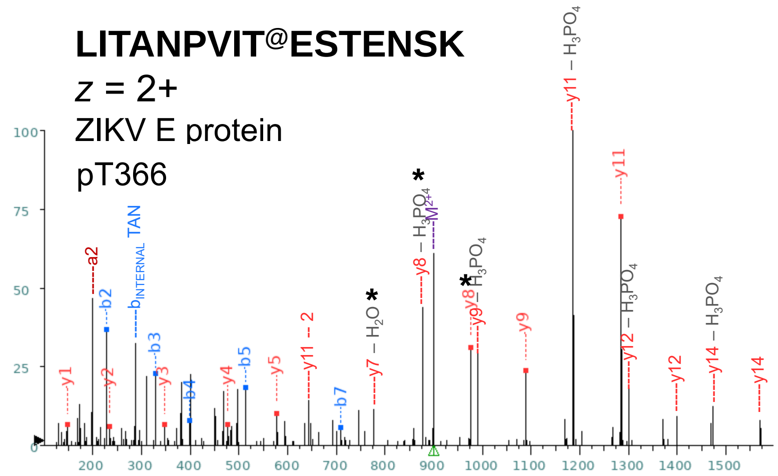


| Seq | # | b: Δ Error | b | y | y: Δ Error | +1 |
|---|---|---|---|---|---|---|
| L | 1 | --- | 114.091 | --- | --- | 16 |
| I | 2 | -1.362 | 227.175 | 1683.784 | --- | 15 |
| T | 3 | -1.577 | 328.223 | 1570.700 | --- | 14 |
| A | 4 | -0.696 | 399.260 | 1469.652 | --- | 13 |
| N | 5 | -2.542 | 513.303 | 1398.615 | --- | 12 |
| P | 6 | --- | 610.356 | 1284.572 | -1.049 | 11 |
| V | 7 | -0.753 | 709.424 | 1187.519 | --- | 10 |
| I | 8 | --- | 822.508 | 1088.451 | -2.058 | 9 |
| T@ | 9 | --- | 1003.522 | 975.367 | -1.527 | 8 |
| E | 10 | --- | 1132.565 | 794.353 | --- | 7 |
| S | 11 | --- | 1219.597 | 665.310 | --- | 6 |
| T | 12 | --- | 1320.645 | 578.278 | -0.240 | 5 |
| E | 13 | --- | 1449.687 | 477.230 | 0.561 | 4 |
| N | 14 | --- | 1563.730 | 348.188 | -1.011 | 3 |
| S | 15 | --- | 1650.762 | 234.145 | -1.810 | 2 |
| K | 16 | --- | --- | 147.113 | -3.605 | 1 |

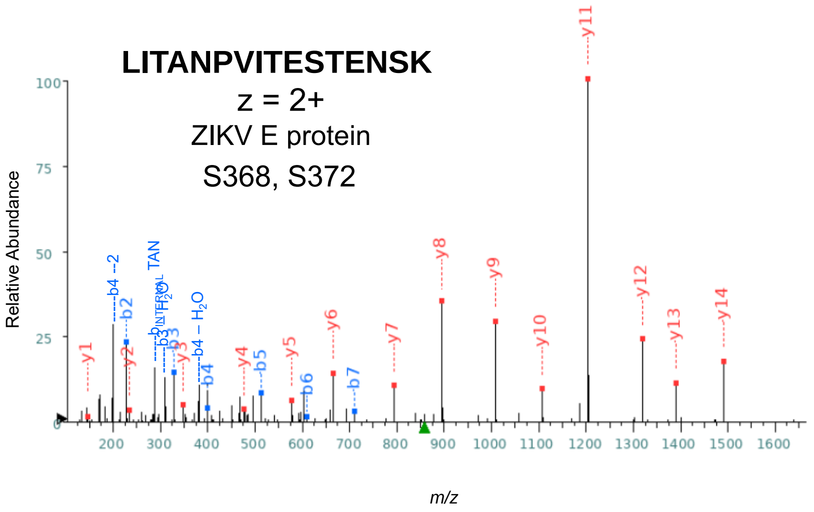


| Seq | # | b: Δ Error | b | y | y: Δ Error | +1 |
|---|---|---|---|---|---|---|
| L | 1 | --- | 114.091 | --- | --- | 16 |
| I | 2 | -0.354 | 227.175 | 1603.817 | --- | 15 |
| T | 3 | -0.741 | 328.223 | 1490.733 | -0.339 | 14 |
| A | 4 | 0.985 | 399.260 | 1389.686 | -1.542 | 13 |
| N | 5 | 0.431 | 513.303 | 1318.649 | 0.044 | 12 |
| P | 6 | -0.586 | 610.356 | 1204.606 | -0.391 | 11 |
| V | 7 | 0.451 | 709.424 | 1107.553 | -0.509 | 10 |
| I | 8 | --- | 822.508 | 1008.484 | 0.100 | 9 |
| T | 9 | --- | 923.556 | 895.400 | -1.025 | 8 |
| E | 10 | --- | 1052.599 | 794.353 | 0.010 | 7 |
| S | 11 | --- | 1139.631 | 665.310 | 0.089 | 6 |
| T | 12 | --- | 1240.678 | 578.278 | 3.032 | 5 |
| E | 13 | --- | 1369.721 | 477.230 | 0.561 | 4 |
| N | 14 | --- | 1483.764 | 348.188 | 0.479 | 3 |
| S | 15 | --- | 1570.796 | 234.145 | -0.767 | 2 |
| K | 16 | --- | --- | 147.113 | -1.738 | 1 |

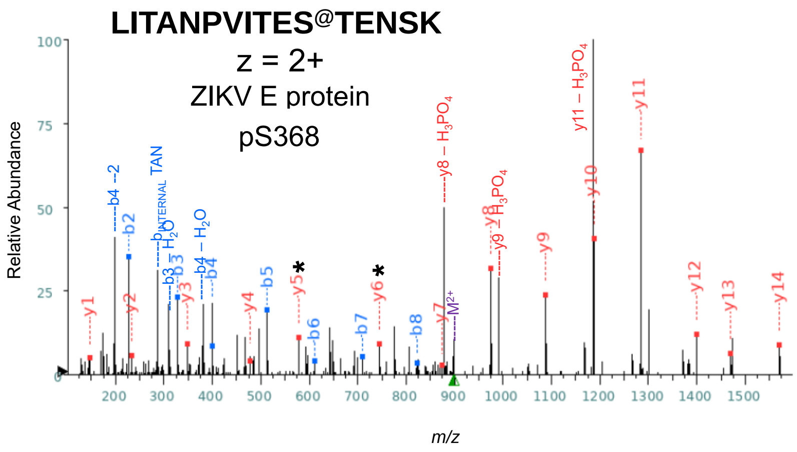


| Seq | # | b: Δ Error | b | y | y: Δ Error | +1 |
|---|---|---|---|---|---|---|
| L | 1 | --- | 114.091 | --- | --- | 16 |
| I | 2 | -1.630 | 227.175 | 1683.784 | --- | 15 |
| T | 3 | -1.484 | 328.223 | 1570.700 | -1.424 | 14 |
| A | 4 | -2.607 | 399.260 | 1469.652 | 0.686 | 13 |
| N | 5 | -1.234 | 513.303 | 1398.615 | -0.410 | 12 |
| P | 6 | -0.686 | 610.356 | 1284.572 | -1.239 | 11 |
| V | 7 | --- | 709.424 | 1187.519 | --- | 10 |
| I | 8 | 1.777 | 822.508 | 1088.451 | -1.385 | 9 |
| T | 9 | --- | 923.556 | 975.367 | -1.339 | 8 |
| E | 10 | --- | 1052.599 | 874.319 | -0.854 | 7 |
| S@ | 11 | --- | 1219.597 | 745.276 | -1.178 | 6 |
| T | 12 | --- | 1320.645 | 578.278 | 0.182 | 5 |
| E | 13 | --- | 1449.687 | 477.230 | 1.393 | 4 |
| N | 14 | --- | 1563.730 | 348.188 | -2.588 | 3 |
| S | 15 | --- | 1650.762 | 234.145 | -2.136 | 2 |
| K | 16 | --- | --- | 147.113 | -0.701 | 1 |

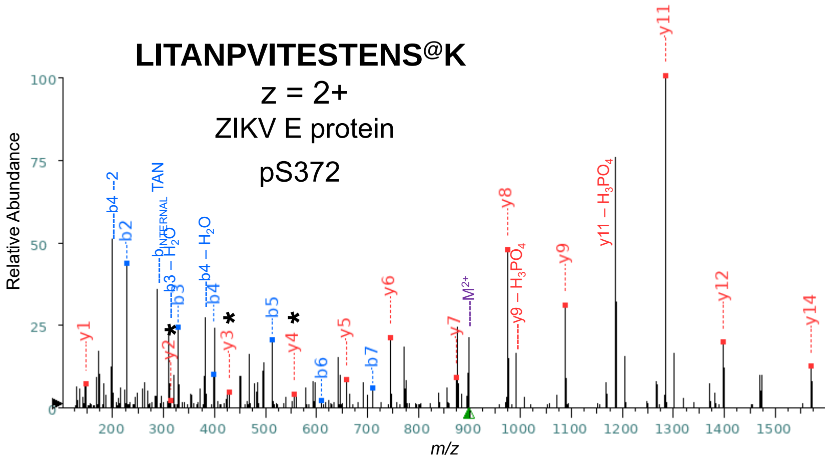


| Seq | # | b: Δ Error | b | y | y: Δ Error | +1 |
|---|---|---|---|---|---|---|
| L | 1 | --- | 114.091 | --- | --- | 16 |
| I | 2 | -0.959 | 227.175 | 1683.784 | --- | 15 |
| T | 3 | -1.670 | 328.223 | 1570.700 | -1.268 | 14 |
| A | 4 | -0.543 | 399.260 | 1469.652 | --- | 13 |
| N | 5 | -1.115 | 513.303 | 1398.615 | -0.323 | 12 |
| P | 6 | 3.114 | 610.356 | 1284.572 | -0.669 | 11 |
| V | 7 | 0.107 | 709.424 | 1187.519 | --- | 10 |
| I | 8 | --- | 822.508 | 1088.451 | -1.834 | 9 |
| T | 9 | --- | 923.556 | 975.367 | -1.214 | 8 |
| E | 10 | --- | 1052.599 | 874.319 | 1.031 | 7 |
| S | 11 | --- | 1139.631 | 745.276 | -1.751 | 6 |
| T | 12 | --- | 1240.678 | 658.244 | 0.219 | 5 |
| E | 13 | --- | 1369.721 | 557.197 | -0.818 | 4 |
| N | 14 | --- | 1483.764 | 428.154 | 0.481 | 3 |
| S@ | 15 | --- | 1650.762 | 314.111 | 5.188 | 2 |
| K | 16 | --- | --- | 147.113 | -0.701 | 1 |

**QSDTQYVC^KR**
$z$ = 3+
ZIKV E protein
K93

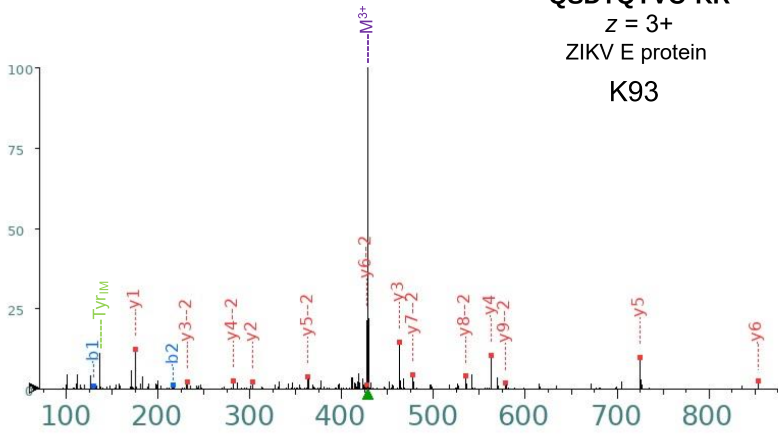


Sequence

QSDTQYVC^KR

Predicted Fragmentation Pattern

+1

| Seq | # | b: Δ Error | b | y | y: Δ Error | +1 |
|---|---|---|---|---|---|---|
| Q | 1 | -1.629 | 129.066 | --- | --- | 10 |
| S | 2 | -1.397 | 216.098 | 1156.542 | --- | 9 |
| D | 3 | --- | 331.125 | 1069.510 | --- | 8 |
| T | 4 | --- | 432.173 | 954.483 | --- | 7 |
| Q | 5 | --- | 560.231 | 853.435 | -1.297 | 6 |
| Y | 6 | --- | 723.294 | 725.376 | -0.959 | 5 |
| V | 7 | --- | 822.363 | 562.313 | -1.284 | 4 |
| C^ | 8 | --- | 982.393 | 463.245 | -1.309 | 3 |
| K | 9 | --- | 1110.488 | 303.214 | -1.870 | 2 |

**QSDTQYVC^K~R**
$z$ = 3+
ZIKV E protein
ubK93

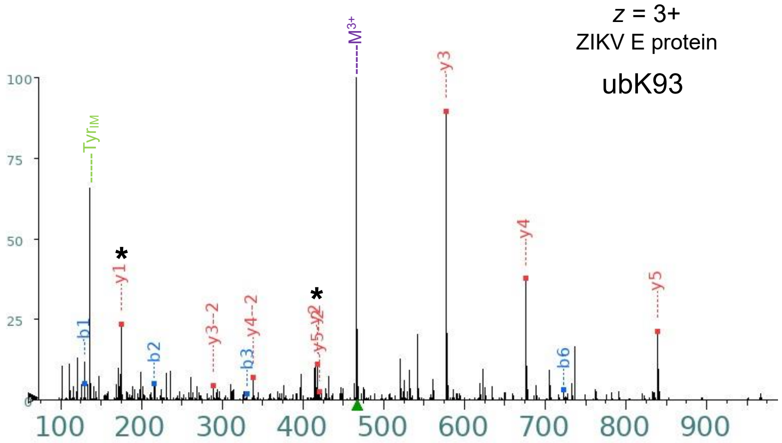


Sequence

QSDTQYVC^K~R

Predicted Fragmentation Pattern

+1

| Seq | # | b: Δ Error | b | y | y: Δ Error | +1 |
|---|---|---|---|---|---|---|
| Q | 1 | -0.565 | 129.066 | --- | --- | 10 |
| S | 2 | -0.479 | 216.098 | 1270.584 | --- | 9 |
| D | 3 | 1.819 | 331.125 | 1183.552 | --- | 8 |
| T | 4 | --- | 432.173 | 1068.525 | --- | 7 |
| Q | 5 | --- | 560.231 | 967.478 | --- | 6 |
| Y | 6 | -1.232 | 723.294 | 839.419 | -0.925 | 5 |
| V | 7 | --- | 822.363 | 676.356 | -1.006 | 4 |
| C^ | 8 | --- | 982.393 | 577.287 | -0.450 | 3 |

**GSLVTCAKFC^SK**

$z$ = 3+

ZIKV E protein

K118

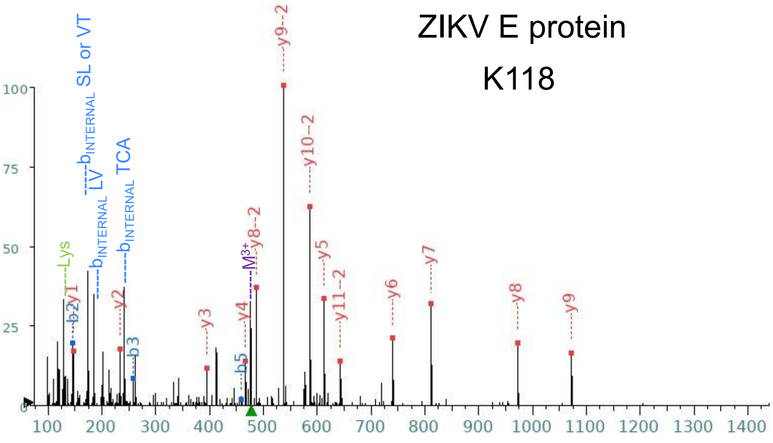


+1

| Seq | # | b: Δ Error | b | y | y: Δ Error | +1 |
|---|---|---|---|---|---|---|
| G | 1 | --- | 58.029 | --- | --- | 13 |
| S | 2 | -0.368 | **145.061** | 1371.676 | --- | 12 |
| L | 3 | -0.693 | **258.145** | 1284.644 | --- | 11 |
| V | 4 | --- | 357.213 | 1171.560 | --- | 10 |
| T | 5 | -0.131 | **458.261** | **1072.491** | 0.154 | 9 |
| C^ | 6 | --- | 618.292 | **971.444** | -1.328 | 8 |
| A | 7 | --- | 689.329 | **811.413** | 0.376 | 7 |
| K | 8 | --- | 817.424 | **740.376** | 0.089 | 6 |
| F | 9 | --- | 964.492 | **612.281** | -0.703 | 5 |
| A | 10 | --- | 1035.529 | **465.213** | -1.857 | 4 |
| C^ | 11 | --- | 1195.560 | **394.175** | -1.019 | 3 |
| S | 12 | --- | 1282.592 | **234.145** | -1.028 | 2 |
| K | 13 | --- | --- | **147.113** | -0.493 | 1 |

+2

| Seq | # | b: Δ Error | b | y | y: Δ Error | +1 |
|---|---|---|---|---|---|---|
| G | 1 | --- | 29.518 | --- | --- | 13 |
| S | 2 | --- | 73.034 | 686.342 | --- | 12 |
| L | 3 | --- | 129.576 | **642.826** | -0.508 | 11 |
| V | 4 | --- | 179.110 | **586.284** | -0.489 | 10 |
| T | 5 | --- | 229.634 | **536.749** | -0.484 | 9 |
| C^ | 6 | --- | 309.649 | **486.226** | -0.838 | 8 |
| A | 7 | --- | 345.168 | 406.210 | --- | 7 |
| K | 8 | --- | 409.215 | 370.692 | --- | 6 |
| F | 9 | --- | 482.750 | 306.644 | --- | 5 |
| A | 10 | --- | 518.268 | 233.110 | --- | 4 |
| C^ | 11 | --- | 598.284 | 197.591 | --- | 3 |
| S | 12 | --- | 641.800 | 117.576 | --- | 2 |
| K | 13 | --- | --- | 74.060 | --- | 1 |

**GSLVTCAK~FC^SK**

$z$ = 3+

ZIKV E protein

ubK118

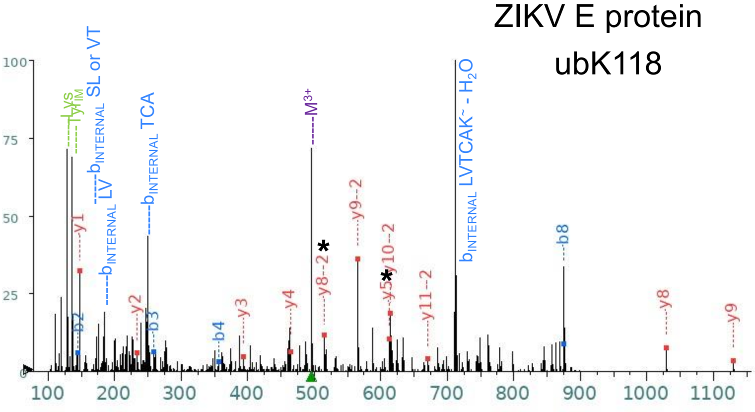


+1

| Seq | # | b: Δ Error | b | y | y: Δ Error | +1 |
|---|---|---|---|---|---|---|
| G | 1 | --- | 58.029 | --- | --- | 13 |
| S | 2 | 0.263 | **145.061** | 1428.697 | --- | 12 |
| L | 3 | -1.047 | **258.145** | 1341.665 | --- | 11 |
| V | 4 | -0.653 | **357.213** | 1228.581 | --- | 10 |
| T | 5 | --- | 458.261 | **1129.513** | 0.488 | 9 |
| C | 6 | --- | 561.270 | **1028.465** | 0.724 | 8 |
| A | 7 | --- | 632.307 | 925.456 | --- | 7 |
| K~ | 8 | 1.903 | **874.445** | 854.419 | --- | 6 |
| F | 9 | --- | 1021.514 | **612.281** | -0.304 | 5 |
| A | 10 | --- | 1092.551 | **465.213** | -0.152 | 4 |
| C^ | 11 | --- | 1252.581 | **394.175** | -3.187 | 3 |
| S | 12 | --- | 1339.613 | **234.145** | 0.276 | 2 |
| K | 13 | --- | --- | **147.113** | -0.182 | 1 |

+2

| Seq | # | b: Δ Error | b | y | y: Δ Error | +1 |
|---|---|---|---|---|---|---|
| G | 1 | --- | 29.518 | --- | --- | 13 |
| S | 2 | --- | 73.034 | 714.852 | --- | 12 |
| L | 3 | --- | 129.576 | **671.336** | -0.562 | 11 |
| V | 4 | --- | 179.110 | **614.794** | -0.053 | 10 |
| T | 5 | --- | 229.634 | **565.260** | -0.225 | 9 |
| C | 6 | --- | 281.139 | **514.736** | -0.060 | 8 |
| A | 7 | --- | 316.657 | 463.232 | --- | 7 |
| K~ | 8 | --- | 437.726 | 427.713 | --- | 6 |
| F | 9 | --- | 511.260 | 306.644 | --- | 5 |
| A | 10 | --- | 546.779 | 233.110 | --- | 4 |
| C^ | 11 | --- | 626.794 | 197.591 | --- | 3 |
| S | 12 | --- | 670.310 | 117.576 | --- | 2 |
| K | 13 | --- | --- | 74.060 | --- | 1 |

**EALVEFKDAHAK**
$z$ = 3+
ZIKV E protein
K246

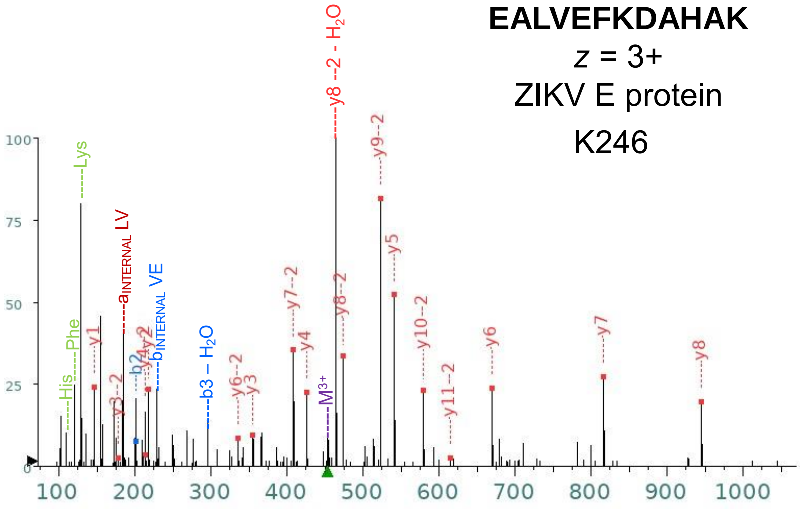


| Seq | # | b: Δ Error | b | y | y: Δ Error | +1 |
|---|---|---|---|---|---|---|
| E | 1 | --- | 130.050 | --- | --- | 12 |
| A | 2 | 1.024 | **201.087** | 1228.668 | --- | 11 |
| L | 3 | --- | 314.171 | 1157.631 | --- | 10 |
| V | 4 | --- | 413.239 | 1044.547 | --- | 9 |
| E | 5 | --- | 542.282 | **945.479** | -0.490 | 8 |
| F | 6 | --- | 689.350 | **816.436** | -2.523 | 7 |
| K | 7 | --- | 817.445 | **669.368** | 0.287 | 6 |
| D | 8 | --- | 932.472 | **541.273** | -1.013 | 5 |
| A | 9 | --- | 1003.509 | **426.246** | -1.224 | 4 |
| H | 10 | --- | 1140.568 | **355.209** | -0.511 | 3 |
| A | 11 | --- | 1211.606 | **218.150** | -0.913 | 2 |
| K | 12 | --- | --- | **147.113** | -0.908 | 1 |

+2

| Seq | # | b: Δ Error | b | y | y: Δ Error | +1 |
|---|---|---|---|---|---|---|
| E | 1 | --- | 65.529 | --- | --- | 12 |
| A | 2 | --- | 101.047 | **614.838** | 2.230 | 11 |
| L | 3 | --- | 157.589 | **579.319** | -0.369 | 10 |
| V | 4 | --- | 207.123 | **522.777** | -0.916 | 9 |
| E | 5 | --- | 271.645 | **473.243** | -0.955 | 8 |
| F | 6 | --- | 345.179 | **408.722** | -1.266 | 7 |
| K | 7 | --- | 409.226 | **335.188** | -0.552 | 6 |
| D | 8 | --- | 466.740 | 271.140 | --- | 5 |
| A | 9 | --- | 502.258 | **213.627** | 1.106 | 4 |
| H | 10 | --- | 570.788 | **178.108** | 0.740 | 3 |
| A | 11 | --- | 606.306 | 109.579 | --- | 2 |
| K | 12 | --- | --- | 74.060 | --- | 1 |

**EALVEFK~DAHAK**
$z$ = 3+
ZIKV E protein
ubK246

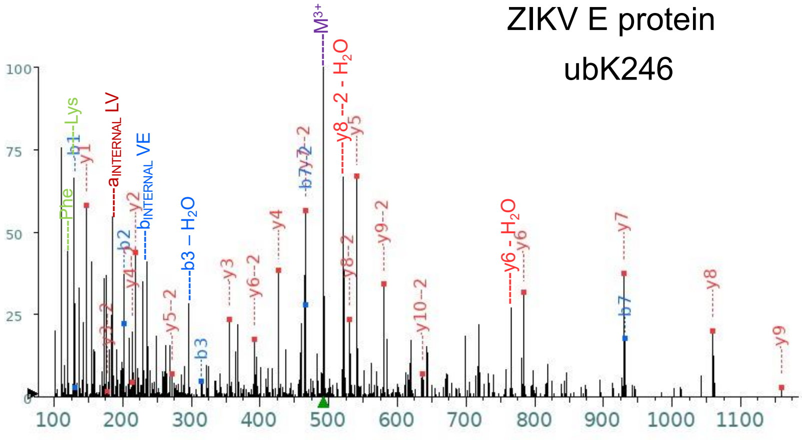


+1

| Seq | # | b: Δ Error | b | y | y: Δ Error | +1 |
|---|---|---|---|---|---|---|
| E | 1 | 1.382 | **130.050** | --- | --- | 12 |
| A | 2 | 0.189 | **201.087** | 1342.711 | --- | 11 |
| L | 3 | -1.250 | **314.171** | 1271.674 | --- | 10 |
| V | 4 | --- | 413.239 | **1158.590** | 0.958 | 9 |
| E | 5 | --- | 542.282 | **1059.522** | 1.215 | 8 |
| F | 6 | --- | 689.350 | **930.479** | 0.061 | 7 |
| K~ | 7 | -2.507 | **931.488** | **783.411** | 1.778 | 6 |
| D | 8 | --- | 1046.515 | **541.273** | 0.679 | 5 |
| A | 9 | --- | 1117.552 | **426.246** | 0.065 | 4 |
| H | 10 | --- | 1254.611 | **355.209** | 0.005 | 3 |
| A | 11 | --- | 1325.648 | **218.150** | -0.003 | 2 |
| K | 12 | --- | --- | **147.113** | 0.129 | 1 |

+2

| Seq | # | b: Δ Error | b | y | y: Δ Error | +1 |
|---|---|---|---|---|---|---|
| E | 1 | --- | 65.529 | --- | --- | 12 |
| A | 2 | --- | 101.047 | 671.859 | --- | 11 |
| L | 3 | --- | 157.589 | **636.341** | 2.382 | 10 |
| V | 4 | --- | 207.123 | **579.799** | 0.052 | 9 |
| E | 5 | --- | 271.645 | **530.265** | 0.223 | 8 |
| F | 6 | --- | 345.179 | **465.743** | 0.178 | 7 |
| K~ | 7 | -5.856 | **466.248** | **392.209** | 0.904 | 6 |
| D | 8 | --- | 523.761 | **271.140** | -0.809 | 5 |
| A | 9 | --- | 559.280 | **213.627** | -2.965 | 4 |
| H | 10 | --- | 627.809 | **178.108** | 1.853 | 3 |
| A | 11 | --- | 663.328 | 109.579 | --- | 2 |
| K | 12 | --- | --- | 74.060 | --- | 1 |

**RQTVVVLGSQEGAVHTALAGALEAEMDGAKGR**

*z* = 4+

ZIKV E protein

K281

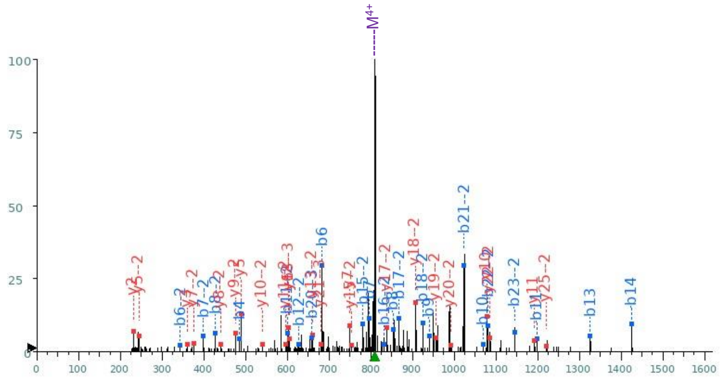


+1

| Seq | # | b: Δ Error | b | y | y: Δ Error | +1 |
|---|---|---|---|---|---|---|
| R | 1 | --- | 157.108 | --- | --- | 32 |
| Q | 2 | --- | 285.167 | 3081.563 | --- | 31 |
| T | 3 | --- | 386.215 | 2953.505 | --- | 30 |
| V | 4 | -1.963 | **485.283** | 2852.457 | --- | 29 |
| V | 5 | --- | 584.351 | 2753.389 | --- | 28 |
| V | 6 | -1.285 | **683.420** | 2654.320 | --- | 27 |
| L | 7 | -2.965 | **796.504** | 2555.252 | --- | 26 |
| G | 8 | -3.315 | **853.525** | 2442.168 | --- | 25 |
| S | 9 | -1.175 | **940.557** | 2385.146 | --- | 24 |
| Q | 10 | -0.962 | **1068.616** | 2298.114 | --- | 23 |
| E | 11 | 0.067 | **1197.659** | 2170.056 | --- | 22 |
| G | 12 | --- | 1254.680 | 2041.013 | --- | 21 |
| A | 13 | -1.309 | **1325.717** | 1983.992 | --- | 20 |
| V | 14 | 0.286 | **1424.786** | 1912.955 | --- | 19 |
| H | 15 | --- | 1561.845 | 1813.886 | --- | 18 |
| T | 16 | --- | 1662.892 | 1676.827 | --- | 17 |
| A | 17 | --- | 1733.929 | 1575.780 | --- | 16 |
| L | 18 | --- | 1847.013 | 1504.742 | --- | 15 |
| A | 19 | --- | 1918.050 | 1391.658 | --- | 14 |
| G | 20 | --- | 1975.072 | 1320.621 | --- | 13 |
| A | 21 | --- | 2046.109 | 1263.600 | --- | 12 |
| L | 22 | --- | 2159.193 | **1192.563** | 2.218 | 11 |
| E | 23 | --- | 2288.236 | **1079.479** | -1.887 | 10 |
| A | 24 | --- | 2359.273 | 950.436 | --- | 9 |
| E | 25 | --- | 2488.315 | 879.399 | --- | 8 |
| M* | 26 | --- | 2635.351 | **750.356** | -1.595 | 7 |
| D | 27 | --- | 2750.378 | **603.321** | -2.086 | 6 |
| G | 28 | --- | 2807.399 | **488.294** | -2.023 | 5 |
| A | 29 | --- | 2878.436 | 431.272 | --- | 4 |
| K | 30 | --- | 3006.531 | **360.235** | -2.534 | 3 |
| G | 31 | --- | 3063.553 | **232.140** | -2.913 | 2 |
| R | 32 | --- | --- | 175.119 | --- | 1 |

+2

| Seq | # | b: Δ Error | b | y | y: Δ Error | +1 |
|---|---|---|---|---|---|---|
| R | 1 | --- | 79.058 | --- | --- | 32 |
| Q | 2 | --- | 143.087 | 1541.285 | --- | 31 |
| T | 3 | --- | 193.611 | 1477.256 | --- | 30 |
| V | 4 | --- | 243.145 | 1426.732 | --- | 29 |
| V | 5 | --- | 292.679 | 1377.198 | --- | 28 |
| V | 6 | -1.748 | **342.214** | 1327.664 | --- | 27 |
| L | 7 | -0.375 | **398.756** | 1278.130 | --- | 26 |
| G | 8 | 0.388 | **427.266** | **1221.588** | 1.797 | 25 |
| S | 9 | --- | 470.782 | 1193.077 | --- | 24 |
| Q | 10 | --- | 534.812 | 1149.561 | --- | 23 |
| E | 11 | 0.005 | **599.333** | **1085.531** | 0.340 | 22 |
| G | 12 | 3.715 | **627.844** | 1021.010 | --- | 21 |
| A | 13 | --- | 663.362 | **992.499** | -0.443 | 20 |
| V | 14 | --- | 712.896 | **956.981** | -2.881 | 19 |
| H | 15 | -0.615 | **781.426** | **907.447** | 0.624 | 18 |
| T | 16 | -0.620 | **831.950** | **838.917** | -2.627 | 17 |
| A | 17 | -3.271 | **867.468** | 788.393 | --- | 16 |
| L | 18 | -1.859 | **924.010** | **752.875** | -3.040 | 15 |
| A | 19 | --- | 959.529 | 696.333 | --- | 14 |
| G | 20 | --- | 988.040 | **660.814** | -1.829 | 13 |
| A | 21 | -1.076 | **1023.558** | 632.304 | --- | 12 |
| L | 22 | 0.243 | **1080.100** | **596.785** | 2.256 | 11 |
| E | 23 | -0.886 | **1144.621** | **540.243** | 0.419 | 10 |
| A | 24 | --- | 1180.140 | **475.722** | 0.466 | 9 |
| E | 25 | --- | 1244.661 | **440.203** | 1.410 | 8 |
| M* | 26 | --- | 1318.179 | **375.682** | -1.041 | 7 |
| D | 27 | --- | 1375.693 | 302.164 | --- | 6 |
| G | 28 | --- | 1404.203 | **244.651** | -1.983 | 5 |
| A | 29 | --- | 1439.722 | 216.140 | --- | 4 |
| K | 30 | --- | 1503.769 | 180.621 | --- | 3 |
| G | 31 | --- | 1532.280 | 116.574 | --- | 2 |
| R | 32 | --- | --- | 88.063 | --- | 1 |

+3

| Seq | # | b: Δ Error | b | y | y: Δ Error | +1 |
|---|---|---|---|---|---|---|
| R | 1 | --- | 53.041 | --- | --- | 32 |
| Q | 2 | --- | 95.727 | 1027.859 | --- | 31 |
| T | 3 | --- | 129.410 | 985.173 | --- | 30 |
| V | 4 | --- | 162.433 | 951.491 | --- | 29 |
| V | 5 | --- | 195.455 | 918.468 | --- | 28 |
| V | 6 | --- | 228.478 | 885.445 | --- | 27 |
| L | 7 | --- | 266.173 | 852.422 | --- | 26 |
| G | 8 | --- | 285.180 | 814.727 | --- | 25 |
| S | 9 | --- | 314.191 | 795.720 | --- | 24 |
| Q | 10 | --- | 356.877 | 766.710 | --- | 23 |
| E | 11 | --- | 399.891 | 724.023 | --- | 22 |
| G | 12 | --- | 418.898 | **681.009** | -1.263 | 21 |
| A | 13 | --- | 442.577 | 662.002 | --- | 20 |
| V | 14 | --- | 475.600 | 638.323 | --- | 19 |
| H | 15 | --- | 521.286 | **605.300** | -0.694 | 18 |
| T | 16 | --- | 554.969 | 559.614 | --- | 17 |
| A | 17 | --- | 578.648 | 525.931 | --- | 16 |
| L | 18 | --- | 616.343 | 502.252 | --- | 15 |
| A | 19 | --- | 640.022 | 464.558 | --- | 14 |
| G | 20 | 0.425 | **659.029** | 440.879 | --- | 13 |
| A | 21 | --- | 682.708 | 421.871 | --- | 12 |
| L | 22 | --- | 720.403 | 398.192 | --- | 11 |
| E | 23 | --- | 763.417 | 360.498 | --- | 10 |
| A | 24 | --- | 787.096 | 317.484 | --- | 9 |
| E | 25 | --- | 830.110 | 293.804 | --- | 8 |
| M* | 26 | --- | 879.122 | 250.790 | --- | 7 |
| D | 27 | --- | 917.464 | 201.778 | --- | 6 |
| G | 28 | --- | 936.471 | 163.436 | --- | 5 |
| A | 29 | --- | 960.150 | 144.429 | --- | 4 |
| K | 30 | --- | 1002.849 | 120.750 | --- | 3 |
| G | 31 | --- | 1021.856 | 78.052 | --- | 2 |
| R | 32 | --- | --- | 59.045 | --- | 1 |

### RQTVVVLGSQEGAVHTALAGALEAEMDGAK~GR

*z* = 4+

ZIKV E protein

ubK281

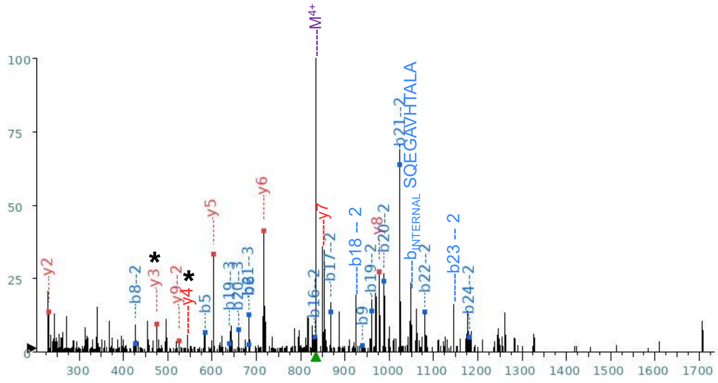


**+1**

| Seq | # | b: Δ Error | b | y | y: Δ Error | +1 |
|---|---|---|---|---|---|---|
| R | 1 | --- | 157.108 | --- | --- | 32 |
| Q | 2 | --- | 285.167 | 3179.611 | --- | 31 |
| T | 3 | --- | 386.215 | 3051.553 | --- | 30 |
| V | 4 | --- | 485.283 | 2950.505 | --- | 29 |
| V | 5 | -2.037 | **584.351** | 2851.437 | --- | 28 |
| V | 6 | -3.071 | **683.420** | 2752.368 | --- | 27 |
| L | 7 | --- | 796.504 | 2653.300 | --- | 26 |
| G | 8 | --- | 853.525 | 2540.216 | --- | 25 |
| S | 9 | 2.394 | **940.557** | 2483.194 | --- | 24 |
| Q | 10 | --- | 1068.616 | 2396.162 | --- | 23 |
| E | 11 | --- | 1197.659 | 2268.104 | --- | 22 |
| G | 12 | --- | 1254.680 | 2139.061 | --- | 21 |
| A | 13 | --- | 1325.717 | 2082.040 | --- | 20 |
| V | 14 | --- | 1424.786 | 2011.003 | --- | 19 |
| H | 15 | --- | 1561.845 | 1911.934 | --- | 18 |
| T | 16 | --- | 1662.892 | 1774.875 | --- | 17 |
| A | 17 | --- | 1733.929 | 1673.828 | --- | 16 |
| L | 18 | --- | 1847.013 | 1602.790 | --- | 15 |
| A | 19 | --- | 1918.050 | 1489.706 | --- | 14 |
| G | 20 | --- | 1975.072 | 1418.669 | --- | 13 |
| A | 21 | --- | 2046.109 | 1361.648 | --- | 12 |
| L | 22 | --- | 2159.193 | 1290.611 | --- | 11 |
| E | 23 | --- | 2288.236 | 1177.527 | --- | 10 |
| A | 24 | --- | 2359.273 | 1048.484 | --- | 9 |
| E | 25 | --- | 2488.315 | **977.447** | -3.003 | 8 |
| M | 26 | --- | 2619.356 | 848.404 | --- | 7 |
| D | 27 | --- | 2734.383 | **717.364** | -2.463 | 6 |
| G | 28 | --- | 2791.404 | **602.337** | -3.598 | 5 |
| A | 29 | --- | 2862.441 | 545.315 | --- | 4 |
| K~ | 30 | --- | 3104.579 | **474.278** | -3.446 | 3 |
| G | 31 | --- | 3161.601 | **232.140** | -3.636 | 2 |
| R | 32 | --- | --- | 175.119 | --- | 1 |

**+2**

| Seq | # | b: Δ Error | b | y | y: Δ Error | +1 |
|---|---|---|---|---|---|---|
| R | 1 | --- | 79.058 | --- | --- | 32 |
| Q | 2 | --- | 143.087 | 1590.309 | --- | 31 |
| T | 3 | --- | 193.611 | 1526.280 | --- | 30 |
| V | 4 | --- | 243.145 | 1475.756 | --- | 29 |
| V | 5 | --- | 292.679 | 1426.222 | --- | 28 |
| V | 6 | --- | 342.214 | 1376.688 | --- | 27 |
| L | 7 | --- | 398.756 | 1327.154 | --- | 26 |
| G | 8 | -3.969 | **427.266** | 1270.612 | --- | 25 |
| S | 9 | --- | 470.782 | 1242.101 | --- | 24 |
| Q | 10 | --- | 534.812 | 1198.585 | --- | 23 |
| E | 11 | --- | 599.333 | 1134.555 | --- | 22 |
| G | 12 | --- | 627.844 | 1070.034 | --- | 21 |
| A | 13 | --- | 663.362 | 1041.523 | --- | 20 |
| V | 14 | --- | 712.896 | 1006.005 | --- | 19 |
| H | 15 | --- | 781.426 | 956.471 | --- | 18 |
| T | 16 | -0.693 | **831.950** | 887.941 | --- | 17 |
| A | 17 | -2.427 | **867.468** | 837.417 | --- | 16 |
| L | 18 | --- | 924.010 | 801.899 | --- | 15 |
| A | 19 | -1.474 | **959.529** | 745.357 | --- | 14 |
| G | 20 | -2.224 | **988.040** | 709.838 | --- | 13 |
| A | 21 | -2.388 | **1023.558** | 681.328 | --- | 12 |
| L | 22 | 2.617 | **1080.100** | 645.809 | --- | 11 |
| E | 23 | --- | 1144.621 | 589.267 | --- | 10 |
| A | 24 | -1.792 | **1180.140** | **524.746** | -4.500 | 9 |
| E | 25 | --- | 1244.661 | 489.227 | --- | 8 |
| M | 26 | --- | 1310.182 | 424.706 | --- | 7 |
| D | 27 | --- | 1367.695 | 359.186 | --- | 6 |
| G | 28 | --- | 1396.206 | 301.672 | --- | 5 |
| A | 29 | --- | 1431.724 | 273.161 | --- | 4 |
| K~ | 30 | --- | 1552.793 | 237.643 | --- | 3 |
| G | 31 | --- | 1581.304 | 116.574 | --- | 2 |
| R | 32 | --- | --- | 88.063 | --- | 1 |

**+3**

| Seq | # | b: Δ Error | b | y | y: Δ Error | +1 |
|---|---|---|---|---|---|---|
| R | 1 | --- | 53.041 | --- | --- | 32 |
| Q | 2 | --- | 95.727 | 1060.542 | --- | 31 |
| T | 3 | --- | 129.410 | 1017.856 | --- | 30 |
| V | 4 | --- | 162.433 | 984.173 | --- | 29 |
| V | 5 | --- | 195.455 | 951.150 | --- | 28 |
| V | 6 | --- | 228.478 | 918.128 | --- | 27 |
| L | 7 | --- | 266.173 | 885.105 | --- | 26 |
| G | 8 | --- | 285.180 | 847.410 | --- | 25 |
| S | 9 | --- | 314.191 | 828.403 | --- | 24 |
| Q | 10 | --- | 356.877 | 799.392 | --- | 23 |
| E | 11 | --- | 399.891 | 756.706 | --- | 22 |
| G | 12 | --- | 418.898 | 713.692 | --- | 21 |
| A | 13 | --- | 442.577 | 694.685 | --- | 20 |
| V | 14 | --- | 475.600 | 671.006 | --- | 19 |
| H | 15 | --- | 521.286 | 637.983 | --- | 18 |
| T | 16 | --- | 554.969 | 592.297 | --- | 17 |
| A | 17 | --- | 578.648 | 558.614 | --- | 16 |
| L | 18 | --- | 616.343 | 534.935 | --- | 15 |
| A | 19 | -0.304 | **640.022** | 497.240 | --- | 14 |
| G | 20 | -4.020 | **659.029** | 473.561 | --- | 13 |
| A | 21 | -2.572 | **682.708** | 454.554 | --- | 12 |
| L | 22 | --- | 720.403 | 430.875 | --- | 11 |
| E | 23 | --- | 763.417 | 393.180 | --- | 10 |
| A | 24 | --- | 787.096 | 350.166 | --- | 9 |
| E | 25 | --- | 830.110 | 326.487 | --- | 8 |
| M | 26 | --- | 873.790 | 283.473 | --- | 7 |
| D | 27 | --- | 912.132 | 239.793 | --- | 6 |
| G | 28 | --- | 931.140 | 201.450 | --- | 5 |
| A | 29 | --- | 954.819 | 182.443 | --- | 4 |
| K~ | 30 | --- | 1035.531 | 158.764 | --- | 3 |
| G | 31 | --- | 1054.538 | 78.052 | --- | 2 |
| R | 32 | --- | --- | 59.045 | --- | 1 |

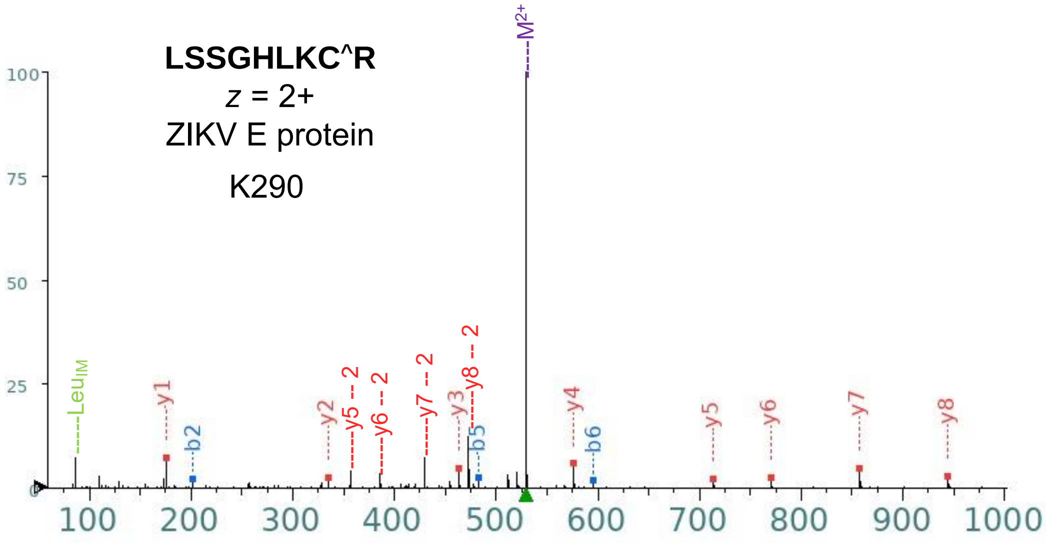
LSSGHLKC^R
z = 2+
ZIKV E protein
K290
M2+
LeuIM
y1
b2
y2
y5 -- 2
y6 -- 2
y7 -- 2
y3
y8 -- 2
b5
y4
b6
y5
y6
y7
y8
100
75
50
25
0
100
200
300
400
500
600
700
800
900
1000

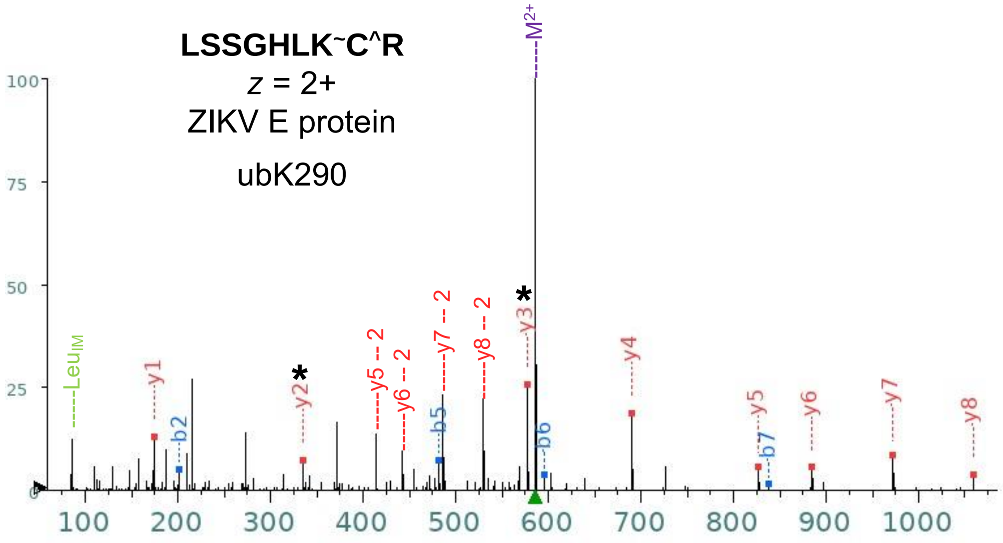
LSSGHLK~C^R
z = 2+
ZIKV E protein
ubK290
M2+
LeuIM
y1
b2
*
y2
y5 -- 2
y6 -- 2
b5
y7 -- 2
y8 -- 2
*
y3
b6
y4
y5
b7
y6
y7
y8
100
75
50
25
0
100
200
300
400
500
600
700
800
900
1000

**CGT@GVFVYNDVEAWR**

$z$ = 2+

ZIKV NS1 protein

pT17

**CGTGVFVY@NDVEAWR**

$z$ = 2+

ZIKV NS1 protein

pY22

| Seq | # | b: Δ Error | b | y | y: Δ Error | +1 |
|---|---|---|---|---|---|---|
| C | 1 | --- | 104.016 | --- | --- | 15 |
| G | 2 | --- | 161.038 | 1692.742 | --- | 14 |
| T | 3 | -5.825 | **262.086** | 1635.720 | --- | 13 |
| G | 4 | -2.138 | **319.107** | 1534.673 | --- | 12 |
| V | 5 | -1.251 | **418.175** | 1477.651 | --- | 11 |
| F | 6 | -3.127 | **565.244** | 1378.583 | --- | 10 |
| V | 7 | -2.192 | **664.312** | 1231.514 | --- | 9 |
| Y@ | 8 | --- | 907.342 | 1132.446 | --- | 8 |
| N | 9 | --- | 1021.385 | **889.416** | 2.131 | 7 |
| D | 10 | --- | 1136.412 | **775.373** | -0.600 | 6 |
| V | 11 | --- | 1235.480 | **660.346** | -1.127 | 5 |
| E | 12 | --- | 1364.523 | **561.278** | -0.576 | 4 |
| A | 13 | --- | 1435.560 | **432.235** | -1.194 | 3 |
| W | 14 | --- | 1621.639 | **361.198** | -1.839 | 2 |
| R | 15 | --- | --- | **175.119** | -1.714 | 1 |

| Seq | # | b: Δ Error | b | y | y: Δ Error | +1 |
|---|---|---|---|---|---|---|
| C | 1 | --- | 104.016 | --- | --- | 15 |
| G | 2 | --- | 161.038 | 1692.742 | --- | 14 |
| T@ | 3 | --- | 342.052 | 1635.720 | --- | 13 |
| G | 4 | --- | 399.073 | **1454.706** | -1.841 | 12 |
| V | 5 | --- | 498.142 | **1397.685** | -2.018 | 11 |
| F | 6 | --- | 645.210 | **1298.616** | -1.190 | 10 |
| V | 7 | --- | 744.279 | **1151.548** | 0.084 | 9 |
| Y | 8 | --- | 907.342 | **1052.480** | -0.784 | 8 |
| N | 9 | --- | 1021.385 | 889.416 | --- | 7 |
| D | 10 | --- | 1136.412 | **775.373** | -0.600 | 6 |
| V | 11 | --- | 1235.480 | **660.346** | -1.127 | 5 |
| E | 12 | --- | 1364.523 | **561.278** | -2.207 | 4 |
| A | 13 | --- | 1435.560 | **432.235** | -2.041 | 3 |
| W | 14 | --- | 1621.639 | **361.198** | -1.078 | 2 |
| R | 15 | --- | --- | **175.119** | -1.801 | 1 |

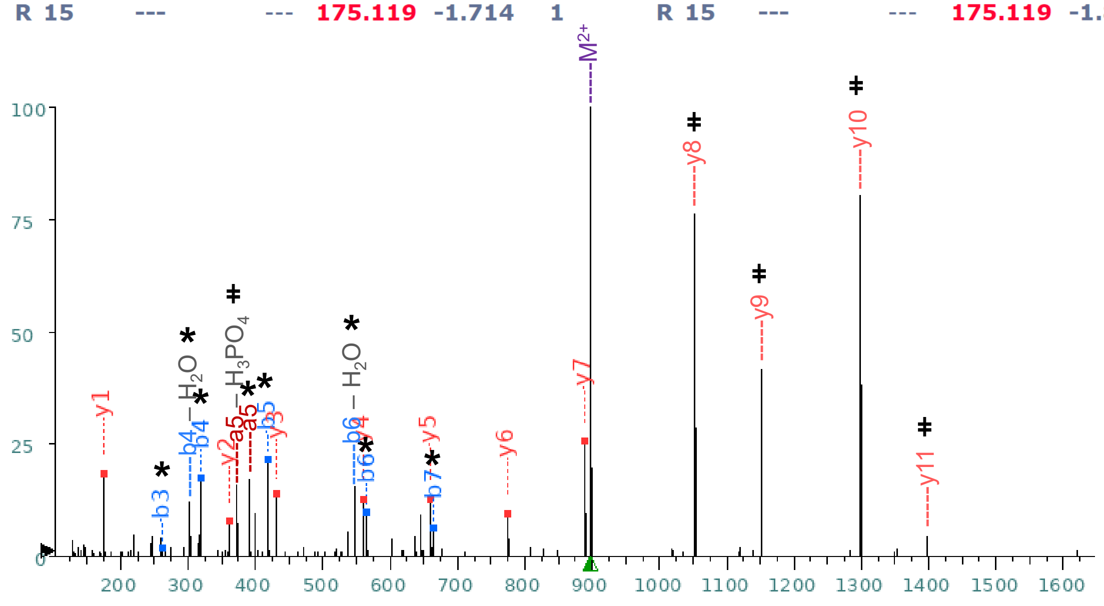

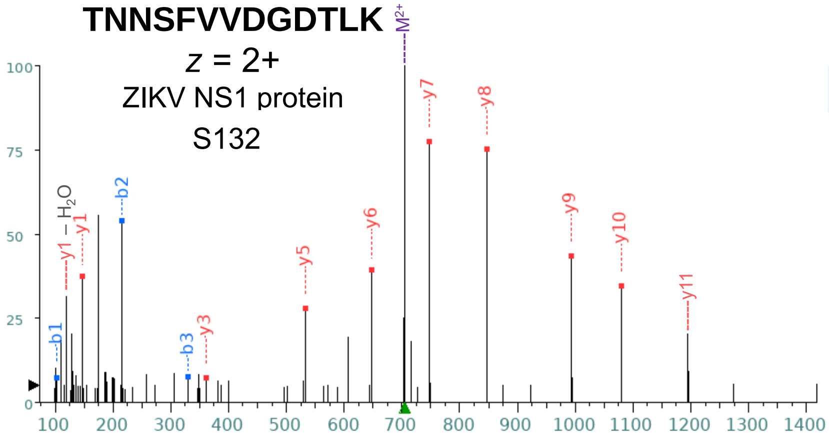


| Seq | # | b: Δ Error | b | y | y: Δ Error | +1 |
|---|---|---|---|---|---|---|
| T | 1 | 2.542 | 102.055 | --- | --- | 13 |
| N | 2 | -0.691 | 216.098 | 1308.643 | --- | 12 |
| N | 3 | -2.407 | 330.141 | 1194.600 | --- | 11 |
| S | 4 | --- | 417.173 | 1080.557 | 0.078 | 10 |
| F | 5 | --- | 564.241 | 993.525 | 2.834 | 9 |
| V | 6 | --- | 663.310 | 846.457 | -2.738 | 8 |
| V | 7 | --- | 762.378 | 747.388 | 0.238 | 7 |
| D | 8 | --- | 877.405 | 648.320 | -2.277 | 6 |
| G | 9 | --- | 934.426 | 533.293 | -5.351 | 5 |
| D | 10 | --- | 1049.453 | 476.271 | --- | 4 |
| T | 11 | --- | 1150.501 | 361.245 | -1.463 | 3 |
| L | 12 | --- | 1263.585 | 260.197 | --- | 2 |
| K | 13 | --- | --- | 147.113 | -2.879 | 1 |

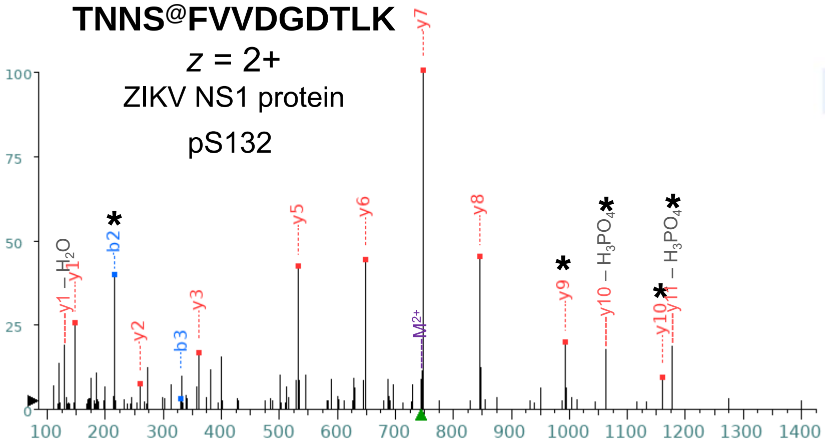


| Seq | # | b: Δ Error | b | y | y: Δ Error | +1 |
|---|---|---|---|---|---|---|
| T | 1 | --- | 102.055 | --- | --- | 13 |
| N | 2 | -1.468 | 216.098 | 1388.609 | --- | 12 |
| N | 3 | -0.189 | 330.141 | 1274.566 | --- | 11 |
| S@ | 4 | --- | 497.139 | 1160.523 | 1.000 | 10 |
| F | 5 | --- | 644.208 | 993.525 | -2.818 | 9 |
| V | 6 | --- | 743.276 | 846.457 | -1.152 | 8 |
| V | 7 | --- | 842.344 | 747.388 | -1.722 | 7 |
| D | 8 | --- | 957.371 | 648.320 | -2.842 | 6 |
| G | 9 | --- | 1014.393 | 533.293 | -2.261 | 5 |
| D | 10 | --- | 1129.420 | 476.271 | --- | 4 |
| T | 11 | --- | 1230.467 | 361.245 | -1.801 | 3 |
| L | 12 | --- | 1343.552 | 260.197 | -2.931 | 2 |
| K | 13 | --- | --- | 147.113 | -1.012 | 1 |

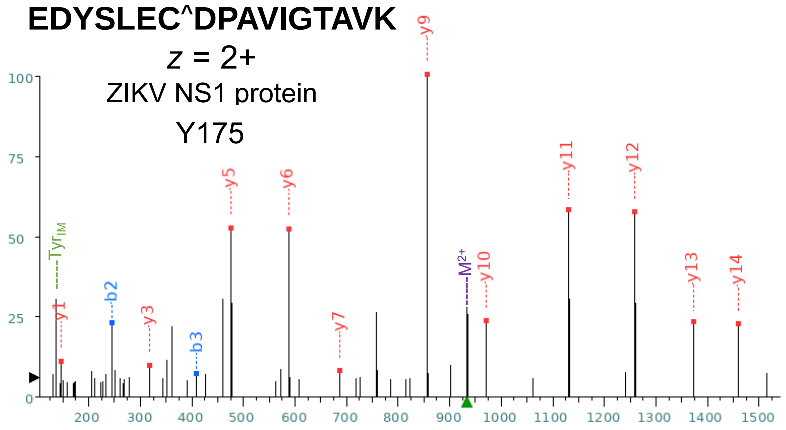


| Seq | # | b: Δ Error | b | y | y: Δ Error | +1 |
|---|---|---|---|---|---|---|
| E | 1 | --- | 130.050 | --- | --- | 17 |
| D | 2 | -3.858 | 245.077 | 1737.836 | --- | 16 |
| Y | 3 | -4.496 | 408.140 | 1622.809 | --- | 15 |
| S | 4 | --- | 495.172 | 1459.746 | -0.343 | 14 |
| L | 5 | --- | 608.256 | 1372.714 | -1.577 | 13 |
| E | 6 | --- | 737.299 | 1259.630 | -1.946 | 12 |
| C^ | 7 | --- | 897.329 | 1130.587 | -1.636 | 11 |
| D | 8 | --- | 1012.356 | 970.557 | 0.178 | 10 |
| P | 9 | --- | 1109.409 | 855.530 | -3.405 | 9 |
| A | 10 | --- | 1180.446 | 758.477 | --- | 8 |
| V | 11 | --- | 1279.515 | 687.440 | -3.390 | 7 |
| I | 12 | --- | 1392.599 | 588.372 | -0.963 | 6 |
| G | 13 | --- | 1449.620 | 475.287 | -1.410 | 5 |
| T | 14 | --- | 1550.668 | 418.266 | --- | 4 |
| A | 15 | --- | 1621.705 | 317.218 | -3.589 | 3 |
| V | 16 | --- | 1720.773 | 246.181 | --- | 2 |
| K | 17 | --- | --- | 147.113 | -4.020 | 1 |

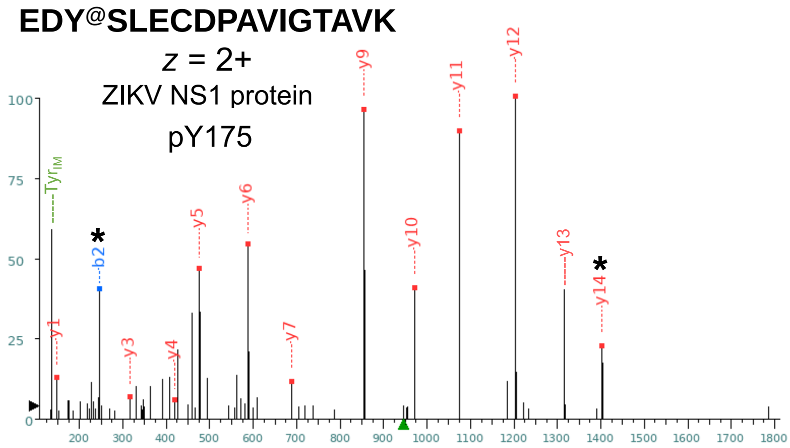


| Seq | # | b: Δ Error | b | y | y: Δ Error | +1 |
|---|---|---|---|---|---|---|
| E | 1 | --- | 130.050 | --- | --- | 17 |
| D | 2 | -0.371 | 245.077 | 1760.781 | --- | 16 |
| Y@ | 3 | --- | 488.106 | 1645.754 | --- | 15 |
| S | 4 | --- | 575.138 | 1402.725 | -0.894 | 14 |
| L | 5 | --- | 688.223 | 1315.693 | --- | 13 |
| E | 6 | --- | 817.265 | 1202.609 | 1.092 | 12 |
| C | 7 | --- | 920.274 | 1073.566 | 0.532 | 11 |
| D | 8 | --- | 1035.301 | 970.557 | 1.310 | 10 |
| P | 9 | --- | 1132.354 | 855.530 | -0.338 | 9 |
| A | 10 | --- | 1203.391 | 758.477 | --- | 8 |
| V | 11 | --- | 1302.460 | 687.440 | 0.606 | 7 |
| I | 12 | --- | 1415.544 | 588.372 | 0.385 | 6 |
| G | 13 | --- | 1472.565 | 475.287 | 0.452 | 5 |
| T | 14 | --- | 1573.613 | 418.266 | 3.383 | 4 |
| A | 15 | --- | 1644.650 | 317.218 | -0.511 | 3 |
| V | 16 | --- | 1743.718 | 246.181 | --- | 2 |
| K | 17 | --- | --- | 147.113 | -2.360 | 1 |

**VREDYSLEC^DPAVIGTAVK**

$z$ = 3+

ZIKV NS1 protein

S176, T186

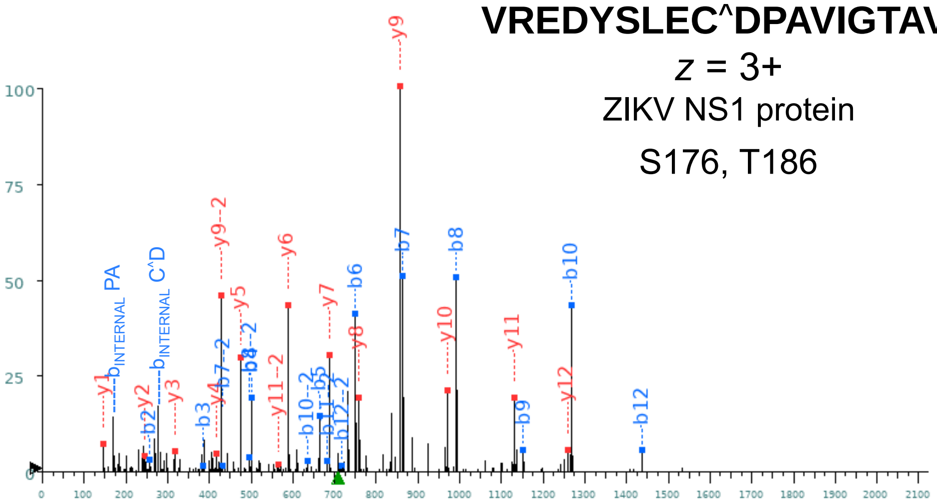


| | | | +1 | | | |
|---|---|---|---|---|---|---|
| Seq | # | b: Δ Error | b | y | y: Δ Error | +1 |
| V | 1 | --- | 100.076 | --- | --- | 19 |
| R | 2 | -1.121 | **256.177** | 2022.980 | --- | 18 |
| E | 3 | -0.642 | **385.219** | 1866.879 | --- | 17 |
| D | 4 | -0.913 | **500.246** | 1737.836 | --- | 16 |
| Y | 5 | -1.156 | **663.310** | 1622.809 | --- | 15 |
| S | 6 | -0.351 | **750.342** | 1459.746 | --- | 14 |
| L | 7 | -0.892 | **863.426** | 1372.714 | --- | 13 |
| E | 8 | -0.521 | **992.468** | **1259.630** | -0.201 | 12 |
| C^ | 9 | -0.297 | **1152.499** | **1130.587** | -0.017 | 11 |
| D | 10 | -0.628 | **1267.526** | **970.557** | -0.262 | 10 |
| P | 11 | --- | 1364.579 | **855.530** | -0.338 | 9 |
| A | 12 | -0.833 | **1435.616** | **758.477** | -0.503 | 8 |
| V | 13 | --- | 1534.684 | **687.440** | -0.815 | 7 |
| I | 14 | --- | 1647.768 | **588.372** | -0.445 | 6 |
| G | 15 | --- | 1704.790 | **475.287** | -0.640 | 5 |
| T | 16 | --- | 1805.837 | **418.266** | -3.330 | 4 |
| A | 17 | --- | 1876.875 | **317.218** | -0.030 | 3 |
| V | 18 | --- | 1975.943 | **246.181** | 0.351 | 2 |
| K | 19 | --- | --- | **147.113** | -0.182 | 1 |

| | | | +2 | | | |
|---|---|---|---|---|---|---|
| Seq | # | b: Δ Error | b | y | y: Δ Error | +1 |
| V | 1 | --- | 50.541 | --- | --- | 19 |
| R | 2 | --- | 128.592 | 1011.994 | --- | 18 |
| E | 3 | --- | 193.113 | 933.943 | --- | 17 |
| D | 4 | --- | 250.627 | 869.422 | --- | 16 |
| Y | 5 | --- | 332.158 | 811.908 | --- | 15 |
| S | 6 | --- | 375.674 | 730.377 | --- | 14 |
| L | 7 | 0.789 | **432.217** | 686.861 | --- | 13 |
| E | 8 | 1.249 | **496.738** | 630.319 | --- | 12 |
| C^ | 9 | --- | 576.753 | **565.797** | 0.026 | 11 |
| D | 10 | 0.180 | **634.267** | 485.782 | --- | 10 |
| P | 11 | -0.569 | **682.793** | **428.269** | -0.424 | 9 |
| A | 12 | -2.668 | **718.312** | 379.742 | --- | 8 |
| V | 13 | --- | 767.846 | 344.224 | --- | 7 |
| I | 14 | --- | 824.388 | 294.689 | --- | 6 |
| G | 15 | --- | 852.899 | 238.147 | --- | 5 |
| T | 16 | --- | 903.422 | 209.637 | --- | 4 |
| A | 17 | --- | 938.941 | 159.113 | --- | 3 |
| V | 18 | --- | 988.475 | 123.594 | --- | 2 |
| K | 19 | --- | --- | 74.060 | --- | 1 |

VREDYS@LEC^DPAVIGTAVK

$z$ = 3+

ZIKV NS1 protein

pS176

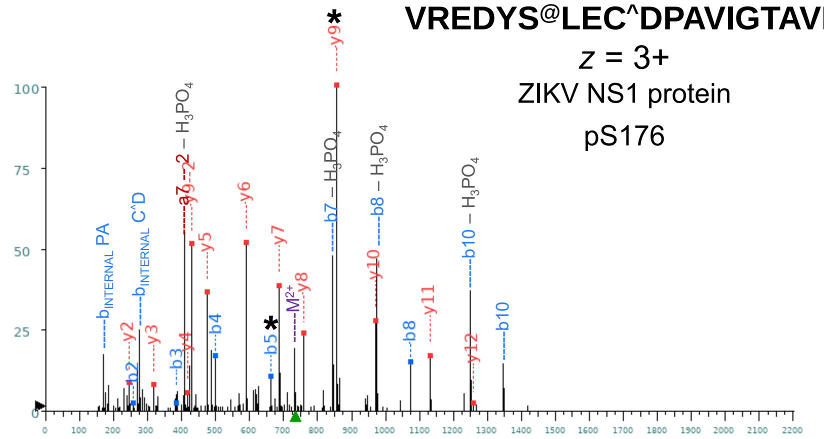


| Seq | # | b: Δ Error | b | y | y: Δ Error | +1 |
|---|---|---|---|---|---|---|
| V | 1 | --- | 100.076 | --- | --- | 19 |
| R | 2 | -1.598 | **256.177** | 2102.946 | --- | 18 |
| E | 3 | -1.830 | **385.219** | 1946.845 | --- | 17 |
| D | 4 | -1.707 | **500.246** | 1817.803 | --- | 16 |
| Y | 5 | -1.800 | **663.310** | 1702.776 | --- | 15 |
| S@ | 6 | --- | 830.308 | 1539.712 | --- | 14 |
| L | 7 | --- | 943.392 | 1372.714 | --- | 13 |
| E | 8 | 0.465 | **1072.435** | **1259.630** | -0.783 | 12 |
| C^ | 9 | --- | 1232.465 | **1130.587** | 1.063 | 11 |
| D | 10 | --- | 1347.492 | **970.557** | -1.268 | 10 |
| P | 11 | --- | 1444.545 | **855.530** | -1.051 | 9 |
| A | 12 | --- | 1515.582 | **758.477** | -0.744 | 8 |
| V | 13 | --- | 1614.651 | **687.440** | -0.993 | 7 |
| I | 14 | --- | 1727.735 | **588.372** | -1.171 | 6 |
| G | 15 | --- | 1784.756 | **475.287** | -1.346 | 5 |
| T | 16 | --- | 1885.804 | **418.266** | 0.756 | 4 |
| A | 17 | --- | 1956.841 | **317.218** | -1.858 | 3 |
| V | 18 | --- | 2055.909 | **246.181** | -2.934 | 2 |
| K | 19 | --- | --- | 147.113 | --- | 1 |

| Seq | # | b: Δ Error | b | y | y: Δ Error | +1 |
|---|---|---|---|---|---|---|
| V | 1 | --- | 50.541 | --- | --- | 19 |
| R | 2 | --- | 128.592 | 1051.977 | --- | 18 |
| E | 3 | --- | 193.113 | 973.926 | --- | 17 |
| D | 4 | --- | 250.627 | 909.405 | --- | 16 |
| Y | 5 | --- | 332.158 | 851.892 | --- | 15 |
| S@ | 6 | --- | 415.658 | 770.360 | --- | 14 |
| L | 7 | --- | 472.200 | 686.861 | --- | 13 |
| E | 8 | --- | 536.721 | 630.319 | --- | 12 |
| C^ | 9 | --- | 616.736 | 565.797 | --- | 11 |
| D | 10 | --- | 674.250 | 485.782 | --- | 10 |
| P | 11 | --- | 722.776 | **428.269** | -0.424 | 9 |
| A | 12 | --- | 758.295 | 379.742 | --- | 8 |
| V | 13 | --- | 807.829 | 344.224 | --- | 7 |
| I | 14 | --- | 864.371 | 294.689 | --- | 6 |
| G | 15 | --- | 892.882 | 238.147 | --- | 5 |
| T | 16 | --- | 943.406 | 209.637 | --- | 4 |
| A | 17 | --- | 978.924 | 159.113 | --- | 3 |
| V | 18 | --- | 1028.458 | 123.594 | --- | 2 |
| K | 19 | --- | --- | 74.060 | --- | 1 |

VREDYSLEC^DPAVIGT@AVK

$z$ = 3+

ZIKV NS1 protein

pS186

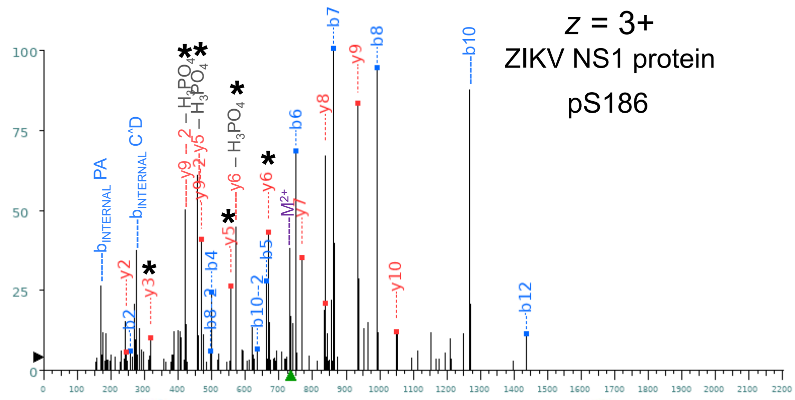


| Seq | # | b: Δ Error | b | y | y: Δ Error | +1 |
|---|---|---|---|---|---|---|
| V | 1 | --- | 100.076 | --- | --- | 19 |
| R | 2 | -4.933 | **256.177** | 2102.946 | --- | 18 |
| E | 3 | --- | 385.219 | 1946.845 | --- | 17 |
| D | 4 | -1.096 | **500.246** | 1817.803 | --- | 16 |
| Y | 5 | 0.592 | **663.310** | 1702.776 | --- | 15 |
| S | 6 | -3.035 | **750.342** | 1539.712 | --- | 14 |
| L | 7 | -2.871 | **863.426** | 1452.680 | --- | 13 |
| E | 8 | -2.427 | **992.468** | 1339.596 | --- | 12 |
| C^ | 9 | --- | 1152.499 | 1210.554 | --- | 11 |
| D | 10 | --- | 1267.526 | **1050.523** | -1.309 | 10 |
| P | 11 | --- | 1364.579 | **935.496** | -3.203 | 9 |
| A | 12 | -0.578 | **1435.616** | **838.443** | -2.010 | 8 |
| V | 13 | --- | 1534.684 | **767.406** | -0.838 | 7 |
| I | 14 | --- | 1647.768 | **668.338** | -1.429 | 6 |
| G | 15 | --- | 1704.790 | **555.254** | 0.622 | 5 |
| T@ | 16 | --- | 1885.804 | 498.232 | --- | 4 |
| A | 17 | --- | 1956.841 | **317.218** | 0.259 | 3 |
| V | 18 | --- | 2055.909 | **246.181** | -3.554 | 2 |
| K | 19 | --- | --- | 147.113 | --- | 1 |

| Seq | # | b: Δ Error | b | y | y: Δ Error | +1 |
|---|---|---|---|---|---|---|
| V | 1 | --- | 50.541 | --- | --- | 19 |
| R | 2 | --- | 128.592 | 1051.977 | --- | 18 |
| E | 3 | --- | 193.113 | 973.926 | --- | 17 |
| D | 4 | --- | 250.627 | 909.405 | --- | 16 |
| Y | 5 | --- | 332.158 | 851.892 | --- | 15 |
| S | 6 | --- | 375.674 | 770.360 | --- | 14 |
| L | 7 | --- | 432.217 | 726.844 | --- | 13 |
| E | 8 | -4.956 | **496.738** | 670.302 | --- | 12 |
| C^ | 9 | --- | 576.753 | 605.781 | --- | 11 |
| D | 10 | 0.180 | **634.267** | 525.765 | --- | 10 |
| P | 11 | --- | 682.793 | **468.252** | -2.301 | 9 |
| A | 12 | --- | 718.312 | 419.725 | --- | 8 |
| V | 13 | --- | 767.846 | 384.207 | --- | 7 |
| I | 14 | --- | 824.388 | 334.673 | --- | 6 |
| G | 15 | --- | 852.899 | 278.131 | --- | 5 |
| T@ | 16 | --- | 943.406 | 249.620 | --- | 4 |
| A | 17 | --- | 978.924 | 159.113 | --- | 3 |
| V | 18 | --- | 1028.458 | 123.594 | --- | 2 |
| K | 19 | --- | --- | 74.060 | --- | 1 |

#### EAVHSDLGYWIESEK

*z* = 3+

ZIKV NS1 protein

Y200

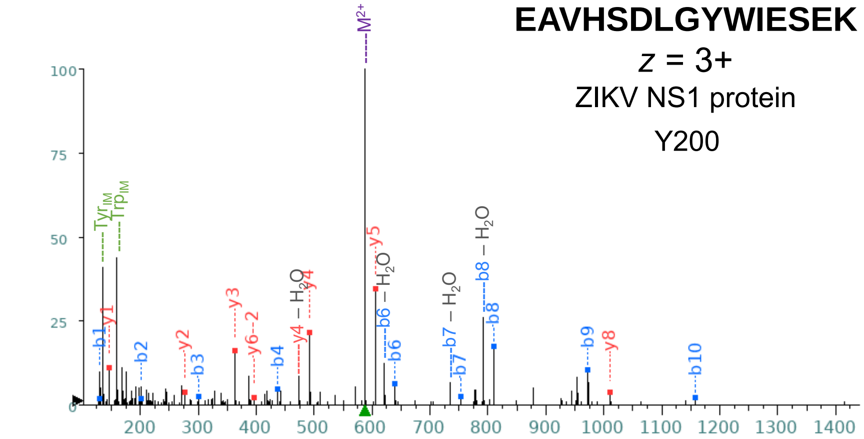


| Seq | # | b: Δ Error | b | y | y: Δ Error | +1 |
|---|---|---|---|---|---|---|
| E | 1 | -1.082 | **130.050** | --- | --- | 15 |
| A | 2 | -1.860 | **201.087** | 1633.786 | --- | 14 |
| V | 3 | -3.156 | **300.155** | 1562.749 | --- | 13 |
| H | 4 | 0.107 | **437.214** | 1463.680 | --- | 12 |
| S | 5 | --- | 524.246 | 1326.621 | --- | 11 |
| D | 6 | -4.146 | **639.273** | 1239.589 | --- | 10 |
| L | 7 | 0.509 | **752.357** | 1124.562 | --- | 9 |
| G | 8 | -1.764 | **809.379** | **1011.478** | 2.677 | 8 |
| Y | 9 | -0.186 | **972.442** | 954.457 | --- | 7 |
| W | 10 | -0.813 | **1158.521** | 791.393 | --- | 6 |
| I | 11 | --- | 1271.606 | **605.314** | -0.901 | 5 |
| E | 12 | --- | 1400.648 | **492.230** | -0.947 | 4 |
| S | 13 | --- | 1487.680 | **363.187** | -1.141 | 3 |
| E | 14 | --- | 1616.723 | **276.155** | -3.986 | 2 |
| K | 15 | --- | --- | **147.113** | -1.219 | 1 |

+2

| Seq | # | b: Δ Error | b | y | y: Δ Error | +1 |
|---|---|---|---|---|---|---|
| E | 1 | --- | 65.529 | --- | --- | 15 |
| A | 2 | --- | 101.047 | 817.396 | --- | 14 |
| V | 3 | --- | 150.581 | 781.878 | --- | 13 |
| H | 4 | --- | 219.111 | 732.344 | --- | 12 |
| S | 5 | --- | 262.627 | 663.814 | --- | 11 |
| D | 6 | --- | 320.140 | 620.298 | --- | 10 |
| L | 7 | --- | 376.682 | 562.785 | --- | 9 |
| G | 8 | --- | 405.193 | 506.243 | --- | 8 |
| Y | 9 | --- | 486.725 | 477.732 | --- | 7 |
| W | 10 | --- | 579.764 | **396.200** | 0.800 | 6 |
| I | 11 | --- | 636.306 | 303.161 | --- | 5 |
| E | 12 | --- | 700.828 | 246.619 | --- | 4 |
| S | 13 | --- | 744.344 | 182.097 | --- | 3 |
| E | 14 | --- | 808.865 | 138.581 | --- | 2 |
| K | 15 | --- | --- | 74.060 | --- | 1 |

#### EAVHSDLGY@WIESEK

*z* = 3+

ZIKV NS1 protein

pY200

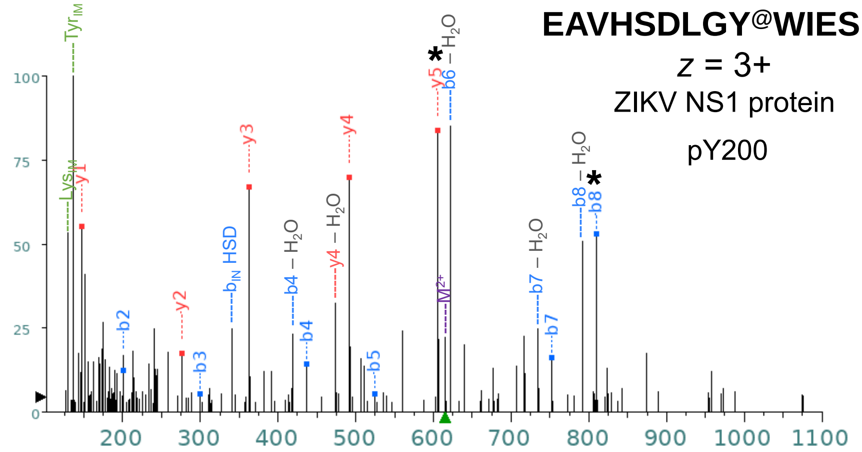


+1

| Seq | # | b: Δ Error | b | y | y: Δ Error | +1 |
|---|---|---|---|---|---|---|
| E | 1 | --- | 130.050 | --- | --- | 15 |
| A | 2 | -1.556 | **201.087** | 1713.752 | --- | 14 |
| V | 3 | -3.563 | **300.155** | 1642.715 | --- | 13 |
| H | 4 | -0.870 | **437.214** | 1543.646 | --- | 12 |
| S | 5 | -1.279 | **524.246** | 1406.588 | --- | 11 |
| D | 6 | --- | 639.273 | 1319.556 | --- | 10 |
| L | 7 | 3.592 | **752.357** | 1204.529 | --- | 9 |
| G | 8 | -0.105 | **809.379** | 1091.445 | --- | 8 |
| Y@ | 9 | --- | 1052.408 | 1034.423 | --- | 7 |
| W | 10 | --- | 1238.488 | 791.393 | --- | 6 |
| I | 11 | --- | 1351.572 | **605.314** | -1.708 | 5 |
| E | 12 | --- | 1480.614 | **492.230** | 0.045 | 4 |
| S | 13 | --- | 1567.646 | **363.187** | -1.981 | 3 |
| E | 14 | --- | 1696.689 | **276.155** | -1.555 | 2 |
| K | 15 | --- | --- | **147.113** | -0.908 | 1 |

+2

| Seq | # | b: Δ Error | b | y | y: Δ Error | +1 |
|---|---|---|---|---|---|---|
| E | 1 | --- | 65.529 | --- | --- | 15 |
| A | 2 | --- | 101.047 | 857.380 | --- | 14 |
| V | 3 | --- | 150.581 | 821.861 | --- | 13 |
| H | 4 | --- | 219.111 | 772.327 | --- | 12 |
| S | 5 | --- | 262.627 | 703.797 | --- | 11 |
| D | 6 | --- | 320.140 | 660.281 | --- | 10 |
| L | 7 | --- | 376.682 | 602.768 | --- | 9 |
| G | 8 | --- | 405.193 | 546.226 | --- | 8 |
| Y@ | 9 | --- | 526.708 | 517.715 | --- | 7 |
| W | 10 | --- | 619.748 | 396.200 | --- | 6 |
| I | 11 | --- | 676.290 | 303.161 | --- | 5 |
| E | 12 | --- | 740.811 | 246.619 | --- | 4 |
| S | 13 | --- | 784.327 | 182.097 | --- | 3 |
| E | 14 | --- | 848.848 | 138.581 | --- | 2 |
| K | 15 | --- | --- | 74.060 | --- | 1 |

**SHTLWTDGIEESDLIIPK**

$z$ = 3+

ZIKV NS1 protein

T233, S239

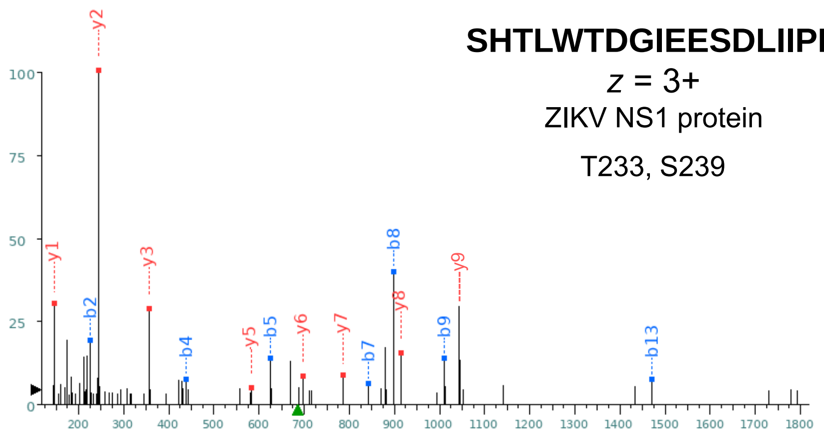


| Seq | # | b: Δ Error | b | y | y: Δ Error | +1 |
|---|---|---|---|---|---|---|
| S | 1 | --- | 88.039 | --- | --- | 18 |
| H | 2 | --- | 225.098 | 1967.012 | --- | 17 |
| T | 3 | -0.531 | **326.146** | 1829.953 | --- | 16 |
| L | 4 | -3.633 | **439.230** | 1728.905 | --- | 15 |
| W | 5 | 1.746 | **625.309** | 1615.821 | --- | 14 |
| T | 6 | --- | 726.357 | 1429.742 | --- | 13 |
| D | 7 | --- | 841.384 | 1328.694 | --- | 12 |
| G | 8 | 1.402 | **898.405** | 1213.667 | --- | 11 |
| I | 9 | 2.071 | **1011.489** | 1156.646 | --- | 10 |
| E | 10 | --- | 1140.532 | **1043.562** | -0.144 | 9 |
| E | 11 | --- | 1269.575 | **914.519** | -0.108 | 8 |
| S | 12 | --- | 1356.607 | **785.477** | 0.717 | 7 |
| D | 13 | --- | 1471.634 | 698.445 | --- | 6 |
| L | 14 | --- | 1584.718 | 583.418 | --- | 5 |
| I | 15 | --- | 1697.802 | 470.334 | --- | 4 |
| I | 16 | --- | 1810.886 | **357.250** | 0.431 | 3 |
| P | 17 | --- | 1907.939 | **244.166** | 0.332 | 2 |
| K | 18 | --- | --- | **147.113** | -0.701 | 1 |

+2

| Seq | # | b: Δ Error | b | y | y: Δ Error | +1 |
|---|---|---|---|---|---|---|
| S | 1 | --- | 44.523 | --- | --- | 18 |
| H | 2 | --- | 113.053 | 984.010 | --- | 17 |
| T | 3 | --- | 163.577 | 915.480 | --- | 16 |
| L | 4 | --- | 220.119 | 864.956 | --- | 15 |
| W | 5 | --- | 313.158 | 808.414 | --- | 14 |
| T | 6 | --- | 363.682 | 715.375 | --- | 13 |
| D | 7 | --- | 421.196 | 664.851 | --- | 12 |
| G | 8 | --- | 449.706 | 607.337 | --- | 11 |
| I | 9 | --- | 506.248 | 578.827 | --- | 10 |
| E | 10 | --- | 570.770 | 522.285 | --- | 9 |
| E | 11 | --- | 635.291 | 457.763 | --- | 8 |
| S | 12 | --- | 678.807 | 393.242 | --- | 7 |
| D | 13 | --- | 736.320 | 349.726 | --- | 6 |
| L | 14 | --- | 792.862 | 292.213 | --- | 5 |
| I | 15 | --- | 849.404 | 235.670 | --- | 4 |
| I | 16 | --- | 905.947 | 179.128 | --- | 3 |
| P | 17 | --- | 954.473 | 122.586 | --- | 2 |
| K | 18 | --- | --- | 74.060 | --- | 1 |

**SHTLWT@DGIEESDLIIPK**

$z$ = 3+

ZIKV NS1 protein

pT233

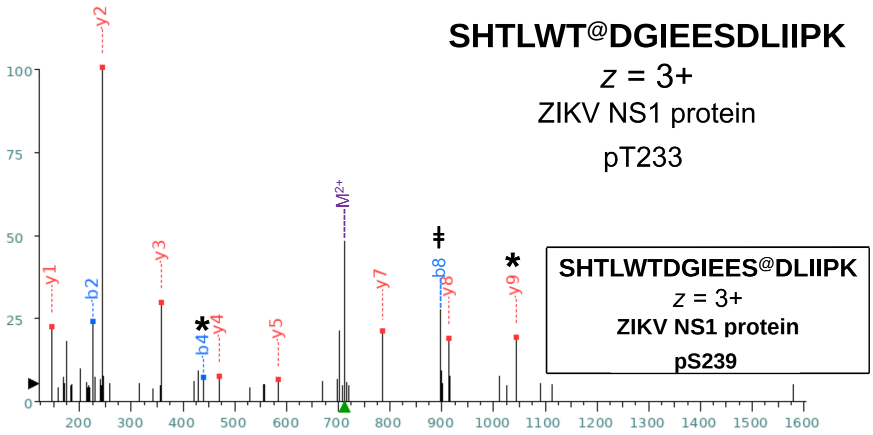


+1

| Seq | # | b: Δ Error | b | y | y: Δ Error | +1 |
|---|---|---|---|---|---|---|
| S | 1 | --- | 88.039 | --- | --- | 18 |
| H | 2 | -1.946 | **225.098** | 2046.978 | --- | 17 |
| T | 3 | --- | 326.146 | 1909.919 | --- | 16 |
| L | 4 | -3.424 | **439.230** | 1808.872 | --- | 15 |
| W | 5 | --- | 625.309 | 1695.788 | --- | 14 |
| T@ | 6 | --- | 806.323 | 1509.708 | --- | 13 |
| D | 7 | --- | 921.350 | 1328.694 | --- | 12 |
| G | 8 | --- | 978.372 | 1213.667 | --- | 11 |
| I | 9 | --- | 1091.456 | 1156.646 | --- | 10 |
| E | 10 | --- | 1220.498 | **1043.562** | -1.314 | 9 |
| E | 11 | --- | 1349.541 | **914.519** | 0.493 | 8 |
| S | 12 | --- | 1436.573 | **785.477** | -1.536 | 7 |
| D | 13 | --- | 1551.600 | 698.445 | --- | 6 |
| L | 14 | --- | 1664.684 | **583.418** | -3.409 | 5 |
| I | 15 | --- | 1777.768 | **470.334** | -1.010 | 4 |
| I | 16 | --- | 1890.852 | **357.250** | -1.363 | 3 |
| P | 17 | --- | 1987.905 | **244.166** | -2.355 | 2 |
| K | 18 | --- | --- | **147.113** | -3.398 | 1 |

+2

| Seq | # | b: Δ Error | b | y | y: Δ Error | +1 |
|---|---|---|---|---|---|---|
| S | 1 | --- | 44.523 | --- | --- | 18 |
| H | 2 | --- | 113.053 | 1023.993 | --- | 17 |
| T | 3 | --- | 163.577 | 955.463 | --- | 16 |
| L | 4 | --- | 220.119 | 904.940 | --- | 15 |
| W | 5 | --- | 313.158 | 848.397 | --- | 14 |
| T@ | 6 | --- | 403.665 | 755.358 | --- | 13 |
| D | 7 | --- | 461.179 | 664.851 | --- | 12 |
| G | 8 | --- | 489.689 | 607.337 | --- | 11 |
| I | 9 | --- | 546.232 | 578.827 | --- | 10 |
| E | 10 | --- | 610.753 | 522.285 | --- | 9 |
| E | 11 | --- | 675.274 | 457.763 | --- | 8 |
| S | 12 | --- | 718.790 | 393.242 | --- | 7 |
| D | 13 | --- | 776.304 | 349.726 | --- | 6 |
| L | 14 | --- | 832.846 | 292.213 | --- | 5 |
| I | 15 | --- | 889.388 | 235.670 | --- | 4 |
| I | 16 | --- | 945.930 | 179.128 | --- | 3 |
| P | 17 | --- | 994.456 | 122.586 | --- | 2 |
| K | 18 | --- | --- | 74.060 | --- | 1 |

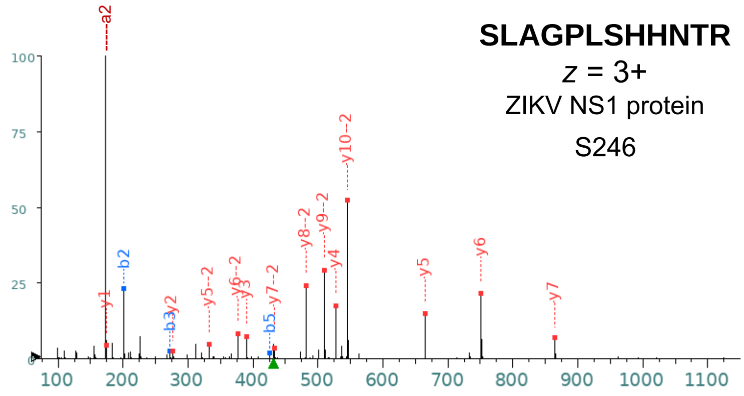


+1

| Seq | # | b: Δ Error | b | y | y: Δ Error | +1 |
|---|---|---|---|---|---|---|
| S | 1 | --- | 88.039 | --- | --- | 12 |
| L | 2 | -1.902 | **201.123** | 1202.639 | --- | 11 |
| A | 3 | -1.982 | **272.160** | 1089.555 | --- | 10 |
| G | 4 | --- | 329.182 | 1018.518 | --- | 9 |
| P | 5 | -3.936 | **426.235** | 961.496 | --- | 8 |
| L | 6 | --- | 539.319 | **864.443** | -1.969 | 7 |
| S | 7 | --- | 626.351 | **751.359** | -1.672 | 6 |
| H | 8 | --- | 763.410 | **664.327** | -2.281 | 5 |
| H | 9 | --- | 900.469 | **527.268** | -1.808 | 4 |
| N | 10 | --- | 1014.512 | **390.210** | -2.410 | 3 |
| T | 11 | --- | 1115.559 | **276.167** | -2.781 | 2 |
| R | 12 | --- | --- | **175.119** | -2.847 | 1 |

+2

| Seq | # | b: Δ Error | b | y | y: Δ Error | +1 |
|---|---|---|---|---|---|---|
| S | 1 | --- | 44.523 | --- | --- | 12 |
| L | 2 | --- | 101.065 | 601.823 | --- | 11 |
| A | 3 | --- | 136.584 | **545.281** | -1.976 | 10 |
| G | 4 | --- | 165.095 | **509.762** | -2.110 | 9 |
| P | 5 | --- | 213.621 | **481.252** | -1.812 | 8 |
| L | 6 | --- | 270.163 | **432.725** | 1.333 | 7 |
| S | 7 | --- | 313.679 | **376.183** | -1.687 | 6 |
| H | 8 | --- | 382.208 | **332.667** | -2.389 | 5 |
| H | 9 | --- | 450.738 | 264.138 | --- | 4 |
| N | 10 | --- | 507.759 | 195.608 | --- | 3 |
| T | 11 | --- | 558.283 | 138.587 | --- | 2 |
| R | 12 | --- | --- | 88.063 | --- | 1 |

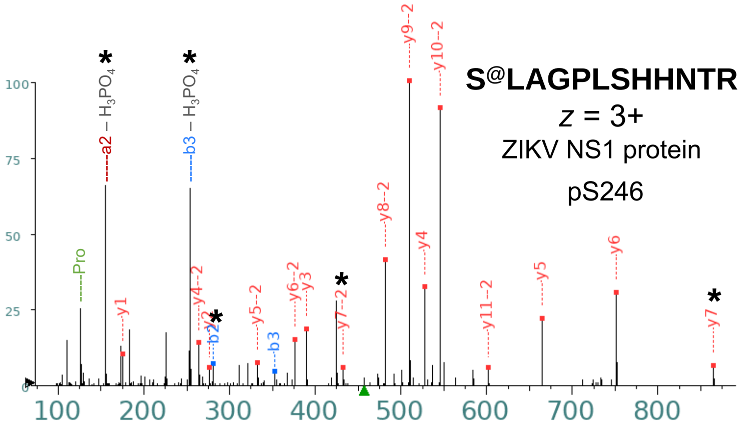


+1

| Seq | # | b: Δ Error | b | y | y: Δ Error | +1 |
|---|---|---|---|---|---|---|
| S@ | 1 | --- | 168.006 | --- | --- | 12 |
| L | 2 | -2.850 | **281.090** | 1202.639 | --- | 11 |
| A | 3 | -2.981 | **352.127** | 1089.555 | --- | 10 |
| G | 4 | --- | 409.148 | 1018.518 | --- | 9 |
| P | 5 | --- | 506.201 | 961.496 | --- | 8 |
| L | 6 | --- | 619.285 | **864.443** | 0.220 | 7 |
| S | 7 | --- | 706.317 | **751.359** | -2.566 | 6 |
| H | 8 | --- | 843.376 | **664.327** | -3.108 | 5 |
| H | 9 | --- | 980.435 | **527.268** | -1.924 | 4 |
| N | 10 | --- | 1094.478 | **390.210** | -3.974 | 3 |
| T | 11 | --- | 1195.526 | **276.167** | -1.897 | 2 |
| R | 12 | --- | --- | **175.119** | -2.063 | 1 |

+2

| Seq | # | b: Δ Error | b | y | y: Δ Error | +1 |
|---|---|---|---|---|---|---|
| S@ | 1 | --- | 84.506 | --- | --- | 12 |
| L | 2 | --- | 141.048 | **601.823** | -3.074 | 11 |
| A | 3 | --- | 176.567 | **545.281** | -2.424 | 10 |
| G | 4 | --- | 205.078 | **509.762** | -2.649 | 9 |
| P | 5 | --- | 253.604 | **481.252** | -2.827 | 8 |
| L | 6 | --- | 310.146 | **432.725** | -3.604 | 7 |
| S | 7 | --- | 353.662 | **376.183** | -2.985 | 6 |
| H | 8 | --- | 422.192 | **332.667** | -4.407 | 5 |
| H | 9 | --- | 490.721 | **264.138** | -2.406 | 4 |
| N | 10 | --- | 547.743 | 195.608 | --- | 3 |
| T | 11 | --- | 598.266 | 138.587 | --- | 2 |
| R | 12 | --- | --- | 88.063 | --- | 1 |

**SM*VTAAAGGGGM*DEK**

$z$ = 3+

ZIKV NS1 protein

S348

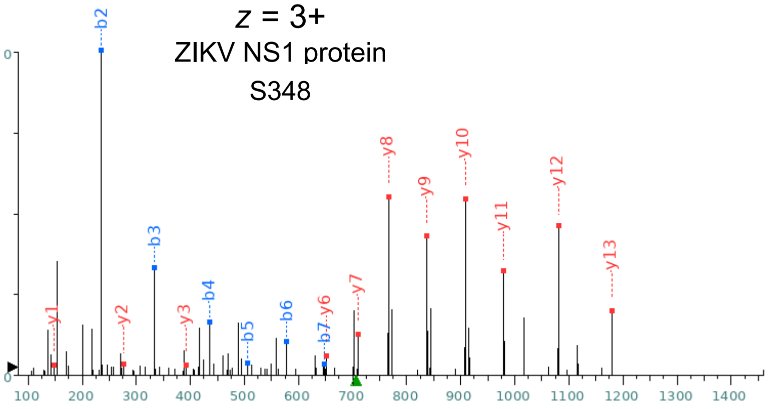


| Seq | # | b: Δ Error | b | y | y: Δ Error | +1 |
|---|---|---|---|---|---|---|
| S | 1 | --- | 88.039 | --- | --- | 15 |
| M* | 2 | -1.933 | **235.075** | 1326.566 | --- | 14 |
| V | 3 | -2.436 | **334.143** | **1179.531** | 0.498 | 13 |
| T | 4 | -1.543 | **435.191** | **1080.463** | -0.874 | 12 |
| A | 5 | -1.094 | **506.228** | **979.415** | -1.328 | 11 |
| A | 6 | 1.465 | **577.265** | **908.378** | -1.696 | 10 |
| A | 7 | -1.904 | **648.302** | **837.341** | -1.908 | 9 |
| G | 8 | --- | 705.324 | **766.304** | -1.840 | 8 |
| G | 9 | --- | 762.345 | **709.282** | 0.995 | 7 |
| G | 10 | --- | 819.367 | **652.261** | -3.722 | 6 |
| G | 11 | --- | 876.388 | 595.239 | --- | 5 |
| M* | 12 | --- | 1023.423 | 538.218 | --- | 4 |
| D | 13 | --- | 1138.450 | **391.182** | 0.473 | 3 |
| E | 14 | --- | 1267.493 | **276.155** | -0.339 | 2 |
| K | 15 | --- | --- | **147.113** | -1.116 | 1 |

**S@M*VTAAAGGGGM*DEK**

$z$ = 3+

ZIKV NS1 protein

pS348

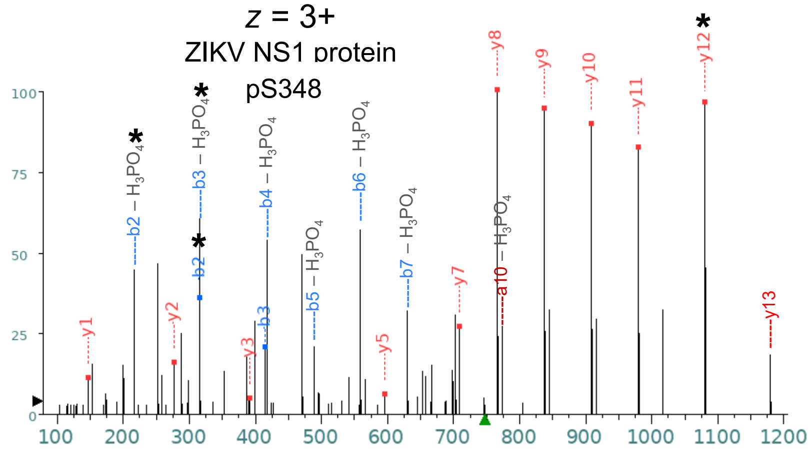


| Seq | # | b: Δ Error | b | y | y: Δ Error | +1 |
|---|---|---|---|---|---|---|
| S@ | 1 | --- | 168.006 | --- | --- | 15 |
| M* | 2 | 0.038 | **315.041** | 1326.566 | --- | 14 |
| V | 3 | 0.266 | **414.109** | 1179.531 | --- | 13 |
| T | 4 | --- | 515.157 | **1080.463** | -1.778 | 12 |
| A | 5 | --- | 586.194 | **979.415** | -1.141 | 11 |
| A | 6 | --- | 657.231 | **908.378** | -0.823 | 10 |
| A | 7 | --- | 728.268 | **837.341** | -1.033 | 9 |
| G | 8 | --- | 785.290 | **766.304** | -1.043 | 8 |
| G | 9 | --- | 842.311 | **709.282** | 0.909 | 7 |
| G | 10 | --- | 899.333 | 652.261 | --- | 6 |
| G | 11 | --- | 956.354 | **595.239** | -4.523 | 5 |
| M* | 12 | --- | 1103.390 | 538.218 | --- | 4 |
| D | 13 | --- | 1218.417 | **391.182** | -4.130 | 3 |
| E | 14 | --- | 1347.459 | **276.155** | 0.766 | 2 |
| K | 15 | --- | --- | **147.113** | -0.805 | 1 |

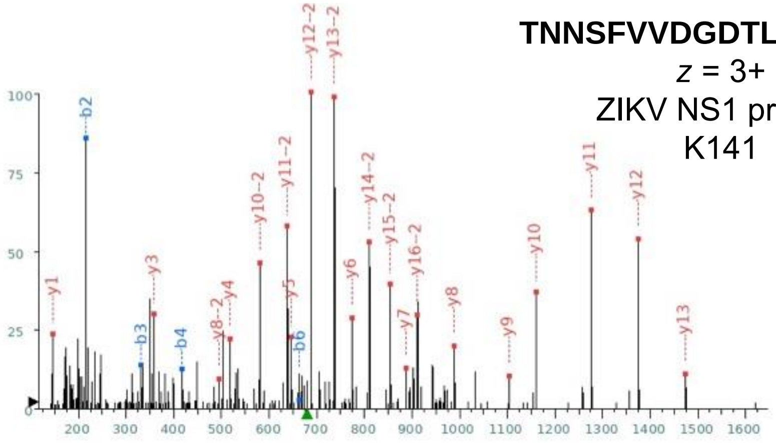


**TNNSFVVDGDTLKEC^PLK**

*z* = 3+

ZIKV NS1 protein

K141

| +1 | | | | | | |
|---|---|---|---|---|---|---|
| Seq | # | b: Δ Error | b | y | y: Δ Error | +1 |
| T | 1 | --- | 102.055 | --- | --- | 18 |
| N | 2 | -0.550 | **216.098** | 1935.948 | --- | 17 |
| N | 3 | 0.181 | **330.141** | 1821.905 | --- | 16 |
| S | 4 | -4.137 | **417.173** | 1707.862 | --- | 15 |
| F | 5 | --- | 564.241 | 1620.830 | --- | 14 |
| V | 6 | 3.721 | **663.310** | **1473.762** | 1.631 | 13 |
| V | 7 | --- | 762.378 | **1374.693** | -2.119 | 12 |
| D | 8 | --- | 877.405 | **1275.625** | -1.284 | 11 |
| G | 9 | --- | 934.426 | **1160.598** | 0.032 | 10 |
| D | 10 | --- | 1049.453 | **1103.577** | 2.448 | 9 |
| T | 11 | --- | 1150.501 | **988.550** | -1.995 | 8 |
| L | 12 | --- | 1263.585 | **887.502** | 2.810 | 7 |
| K | 13 | --- | 1391.680 | **774.418** | -0.302 | 6 |
| E | 14 | --- | 1520.723 | **646.323** | 3.592 | 5 |
| C^ | 15 | --- | 1680.753 | **517.280** | 0.222 | 4 |
| P | 16 | --- | 1777.806 | **357.250** | -1.448 | 3 |
| L | 17 | --- | 1890.890 | 260.197 | --- | 2 |
| K | 18 | --- | --- | **147.113** | -0.493 | 1 |

| +2 | | | | | | |
|---|---|---|---|---|---|---|
| Seq | # | b: Δ Error | b | y | y: Δ Error | +1 |
| T | 1 | --- | 51.531 | --- | --- | 18 |
| N | 2 | --- | 108.553 | 968.478 | --- | 17 |
| N | 3 | --- | 165.574 | **911.456** | -1.701 | 16 |
| S | 4 | --- | 209.090 | **854.435** | -0.839 | 15 |
| F | 5 | --- | 282.624 | **810.919** | -1.006 | 14 |
| V | 6 | --- | 332.158 | **737.385** | -1.235 | 13 |
| V | 7 | --- | 381.693 | **687.850** | -1.018 | 12 |
| D | 8 | --- | 439.206 | **638.316** | -0.385 | 11 |
| G | 9 | --- | 467.717 | **580.803** | -1.293 | 10 |
| D | 10 | --- | 525.230 | 552.292 | --- | 9 |
| T | 11 | --- | 575.754 | **494.778** | -0.093 | 8 |
| L | 12 | --- | 632.296 | 444.255 | --- | 7 |
| K | 13 | --- | 696.344 | 387.713 | --- | 6 |
| E | 14 | --- | 760.865 | 323.665 | --- | 5 |
| C^ | 15 | --- | 840.880 | 259.144 | --- | 4 |
| P | 16 | --- | 889.407 | 179.128 | --- | 3 |
| L | 17 | --- | 945.949 | 130.602 | --- | 2 |
| K | 18 | --- | --- | 74.060 | --- | 1 |

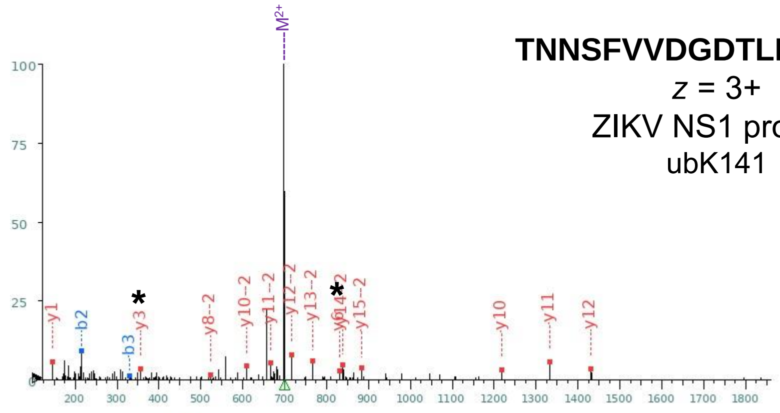


**TNNSFVVDGDTLK~ECPLK**

*z* = 3+

ZIKV NS1 protein

ubK141

| +1 | | | | | | |
|---|---|---|---|---|---|---|
| Seq | # | b: Δ Error | b | y | y: Δ Error | +1 |
| T | 1 | --- | 102.055 | --- | --- | 18 |
| N | 2 | -1.115 | **216.098** | 1992.970 | --- | 17 |
| N | 3 | 2.400 | **330.141** | 1878.927 | --- | 16 |
| S | 4 | --- | 417.173 | 1764.884 | --- | 15 |
| F | 5 | --- | 564.241 | 1677.852 | --- | 14 |
| V | 6 | --- | 663.310 | 1530.783 | --- | 13 |
| V | 7 | --- | 762.378 | **1431.715** | 1.731 | 12 |
| D | 8 | --- | 877.405 | **1332.646** | -0.572 | 11 |
| G | 9 | --- | 934.426 | **1217.619** | -1.357 | 10 |
| D | 10 | --- | 1049.453 | 1160.598 | --- | 9 |
| T | 11 | --- | 1150.501 | 1045.571 | --- | 8 |
| L | 12 | --- | 1263.585 | 944.523 | --- | 7 |
| K~ | 13 | --- | 1505.723 | **831.439** | -0.404 | 6 |
| E | 14 | --- | 1634.766 | 589.301 | --- | 5 |
| C | 15 | --- | 1737.775 | 460.259 | --- | 4 |
| P | 16 | --- | 1834.828 | **357.250** | -0.338 | 3 |
| L | 17 | --- | 1947.912 | 260.197 | --- | 2 |
| K | 18 | --- | --- | **147.113** | -0.390 | 1 |

| +2 | | | | | | |
|---|---|---|---|---|---|---|
| Seq | # | b: Δ Error | b | y | y: Δ Error | +1 |
| T | 1 | --- | 51.531 | --- | --- | 18 |
| N | 2 | --- | 108.553 | 996.988 | --- | 17 |
| N | 3 | --- | 165.574 | 939.967 | --- | 16 |
| S | 4 | --- | 209.090 | **882.945** | 0.583 | 15 |
| F | 5 | --- | 282.624 | **839.429** | -2.704 | 14 |
| V | 6 | --- | 332.158 | **765.895** | -3.327 | 13 |
| V | 7 | --- | 381.693 | **716.361** | -0.707 | 12 |
| D | 8 | --- | 439.206 | **666.827** | -2.733 | 11 |
| G | 9 | --- | 467.717 | **609.313** | 0.387 | 10 |
| D | 10 | --- | 525.230 | 580.803 | --- | 9 |
| T | 11 | --- | 575.754 | **523.289** | 3.314 | 8 |
| L | 12 | --- | 632.296 | 472.765 | --- | 7 |
| K~ | 13 | --- | 753.365 | 416.223 | --- | 6 |
| E | 14 | --- | 817.886 | 295.154 | --- | 5 |
| C | 15 | --- | 869.391 | 230.633 | --- | 4 |
| P | 16 | --- | 917.917 | 179.128 | --- | 3 |
| L | 17 | --- | 974.459 | 130.602 | --- | 2 |
| K | 18 | --- | --- | 74.060 | --- | 1 |

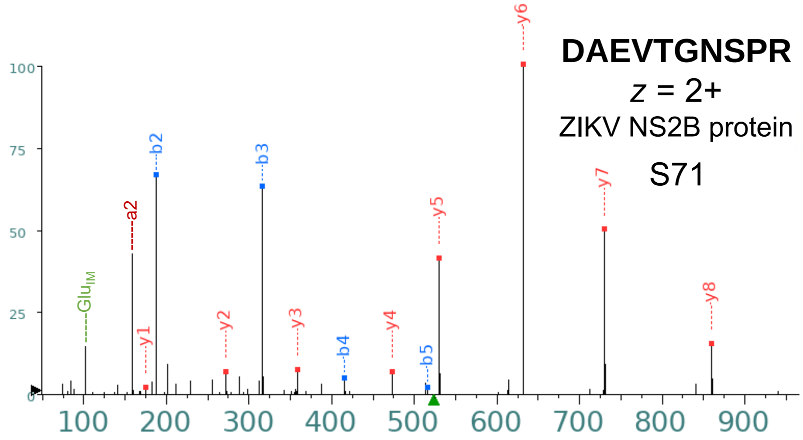


| Seq | # | b: Δ Error | b | y | y: Δ Error | +1 |
|---|---|---|---|---|---|---|
| D | 1 | --- | 116.034 | --- | --- | 10 |
| A | 2 | -0.479 | **187.071** | 930.464 | --- | 9 |
| E | 3 | -1.074 | **316.114** | **859.427** | 0.423 | 8 |
| V | 4 | -3.081 | **415.182** | **730.384** | -0.601 | 7 |
| T | 5 | -2.793 | **516.230** | **631.316** | -0.706 | 6 |
| G | 6 | --- | 573.251 | **530.268** | -0.246 | 5 |
| N | 7 | --- | 687.294 | **473.247** | -1.416 | 4 |
| S | 8 | --- | 774.326 | **359.204** | -1.216 | 3 |
| P | 9 | --- | 871.379 | **272.172** | -0.875 | 2 |
| R | 10 | --- | --- | **175.119** | 0.726 | 1 |

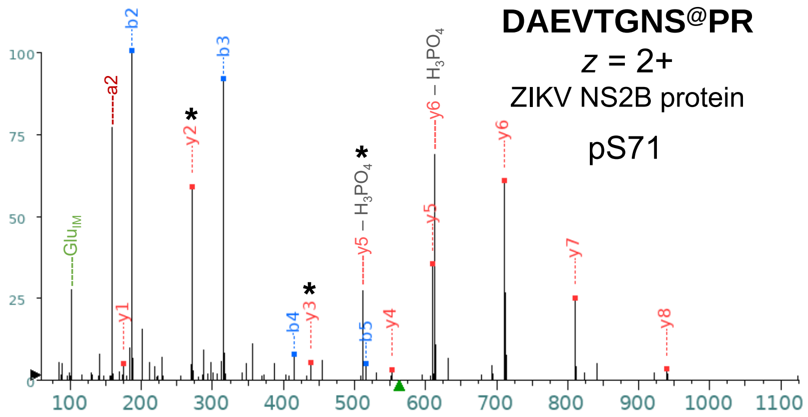


| Seq | # | b: Δ Error | b | y | y: Δ Error | +1 |
|---|---|---|---|---|---|---|
| D | 1 | --- | 116.034 | --- | --- | 10 |
| A | 2 | 0.174 | **187.071** | 1010.430 | --- | 9 |
| E | 3 | -0.302 | **316.114** | **939.393** | -0.611 | 8 |
| V | 4 | 0.668 | **415.182** | **810.351** | -0.343 | 7 |
| T | 5 | -2.674 | **516.230** | **711.282** | -0.057 | 6 |
| G | 6 | --- | 573.251 | **610.234** | -0.950 | 5 |
| N | 7 | --- | 687.294 | **553.213** | 0.128 | 4 |
| S@ | 8 | --- | 854.293 | **439.170** | 0.693 | 3 |
| P | 9 | --- | 951.346 | **272.172** | -0.202 | 2 |
| R | 10 | --- | --- | **175.119** | 0.029 | 1 |

**YWNSSTATSLC^NIFR**

*z* = 2+

ZIKV NS4B protein

S217, S218, T219

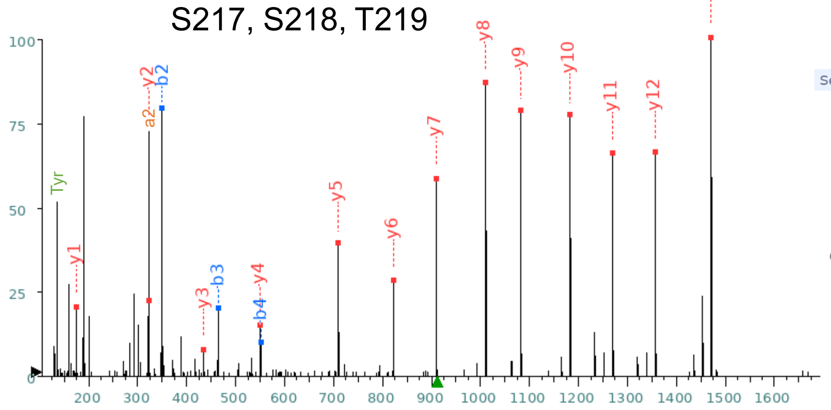


**Predicted Fragmentation Pattern**

| Seq | # | b: Δ Error | b | y | y: Δ Error | +1 |
|---|---|---|---|---|---|---|
| Y | 1 | --- | 164.071 | --- | --- | 15 |
| W | 2 | -2.919 | **350.150** | 1656.780 | --- | 14 |
| N | 3 | -3.231 | **464.193** | **1470.701** | -2.319 | 13 |
| S | 4 | 2.289 | **551.225** | **1356.658** | -3.264 | 12 |
| S | 5 | --- | 638.257 | **1269.626** | -2.394 | 11 |
| T | 6 | --- | 739.305 | **1182.594** | -3.563 | 10 |
| A | 7 | --- | 810.342 | **1081.546** | -2.702 | 9 |
| T | 8 | --- | 911.389 | **1010.509** | -2.525 | 8 |
| S | 9 | --- | 998.421 | **909.461** | -3.533 | 7 |
| L | 10 | --- | 1111.505 | **822.429** | -1.847 | 6 |
| C^ | 11 | --- | 1271.536 | **709.345** | -1.857 | 5 |
| N | 12 | --- | 1385.579 | **549.314** | -0.271 | 4 |
| I | 13 | --- | 1498.663 | **435.271** | -1.699 | 3 |
| F | 14 | --- | 1645.732 | **322.187** | -0.059 | 2 |
| R | 15 | --- | --- | **175.119** | -2.150 | 1 |

**YWNS@STATSLCNIFR**

*z* = 2+

ZIKV NS4B protein

pS217, pS218, pT219

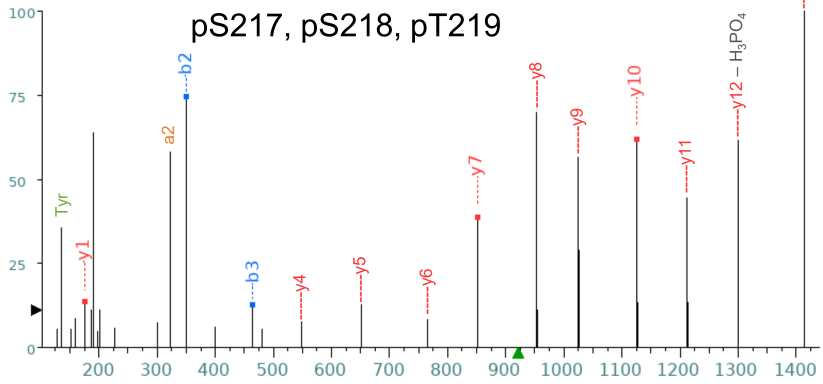


| Seq | # | b: Δ Error | b | y | y: Δ Error | +1 |
|---|---|---|---|---|---|---|
| Y | 1 | --- | 164.071 | --- | --- | 15 |
| W | 2 | -4.314 | **350.150** | 1679.725 | --- | 14 |
| N | 3 | -9.345 | **464.193** | 1493.645 | --- | 13 |
| S@ | 4 | --- | 551.225 | 1379.602 | --- | 12 |
| S | 5 | --- | 718.223 | 1212.6041 | --- | 11 |
| T | 6 | --- | 819.271 | **1125.572** | -0.834 | 10 |
| A | 7 | --- | 890.308 | 1024.524 | --- | 9 |
| T | 8 | --- | 991.356 | 953.487 | --- | 8 |
| S | 9 | --- | 1078.388 | **852.440** | -2.075 | 7 |
| L | 10 | --- | 1191.472 | 765.408 | --- | 6 |
| C | 11 | --- | 1294.481 | 652.324 | --- | 5 |
| N | 12 | --- | 1408.524 | 549.314 | --- | 4 |
| I | 13 | --- | 1521.608 | 435.271 | --- | 3 |
| F | 14 | --- | 1668.676 | 322.187 | --- | 2 |
| R | 15 | --- | --- | **175.119** | -3.805 | 1 |

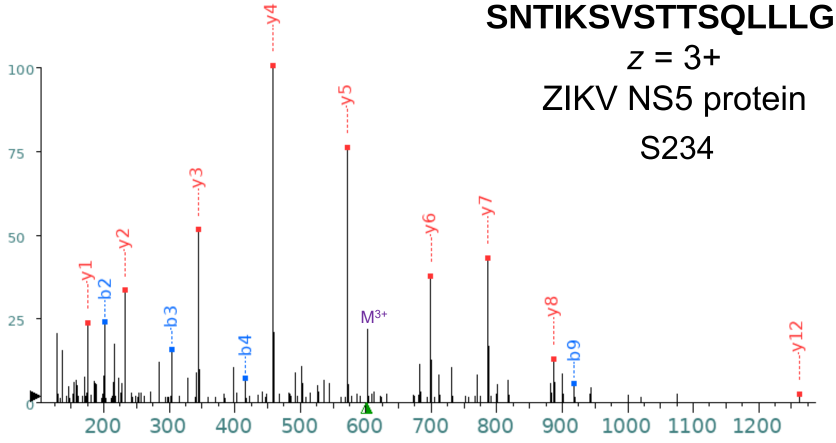


**SNTIKSVSTTSQLLLGR**

$z$ = 3+

ZIKV NS5 protein

S234

+1

| Seq | # | b: Δ Error | b | y | y: Δ Error | +1 |
|---|---|---|---|---|---|---|
| S | 1 | --- | 88.039 | --- | --- | 17 |
| N | 2 | -4.315 | **202.082** | 1717.981 | --- | 16 |
| T | 3 | -6.031 | **303.130** | 1603.938 | --- | 15 |
| I | 4 | -10.743 | **416.214** | 1502.890 | --- | 14 |
| K | 5 | --- | 544.309 | 1389.806 | --- | 13 |
| S | 6 | --- | 631.341 | **1261.711** | -1.727 | 12 |
| V | 7 | --- | 730.409 | 1174.679 | --- | 11 |
| S | 8 | --- | 817.441 | 1075.611 | --- | 10 |
| T | 9 | -4.598 | **918.489** | 988.579 | --- | 9 |
| T | 10 | --- | 1019.537 | **887.531** | -1.729 | 8 |
| S | 11 | --- | 1106.569 | **786.483** | -3.258 | 7 |
| Q | 12 | --- | 1234.627 | **699.451** | -5.692 | 6 |
| L | 13 | --- | 1347.711 | **571.393** | -5.500 | 5 |
| L | 14 | --- | 1460.795 | **458.309** | -5.219 | 4 |
| L | 15 | --- | 1573.880 | **345.224** | -5.814 | 3 |
| G | 16 | --- | 1630.901 | **232.140** | -6.265 | 2 |
| R | 17 | --- | --- | **175.119** | -3.544 | 1 |

+2

| Seq | # | b: Δ Error | b | y | y: Δ Error | +1 |
|---|---|---|---|---|---|---|
| S | 1 | --- | 44.523 | --- | --- | 17 |
| N | 2 | --- | 101.545 | 859.494 | --- | 16 |
| T | 3 | --- | 152.069 | 802.473 | --- | 15 |
| I | 4 | --- | 208.611 | 751.949 | --- | 14 |
| K | 5 | --- | 272.658 | 695.407 | --- | 13 |
| S | 6 | --- | 316.174 | 631.359 | --- | 12 |
| V | 7 | --- | 365.708 | 587.843 | --- | 11 |
| S | 8 | --- | 409.224 | 538.309 | --- | 10 |
| T | 9 | --- | 459.748 | 494.793 | --- | 9 |
| T | 10 | --- | 510.272 | 444.269 | --- | 8 |
| S | 11 | --- | 553.788 | 393.745 | --- | 7 |
| Q | 12 | --- | 617.817 | 350.229 | --- | 6 |
| L | 13 | --- | 674.359 | 286.200 | --- | 5 |
| L | 14 | --- | 730.901 | 229.658 | --- | 4 |
| L | 15 | --- | 787.443 | 173.116 | --- | 3 |
| G | 16 | --- | 815.954 | 116.574 | --- | 2 |
| R | 17 | --- | --- | 88.063 | --- | 1 |

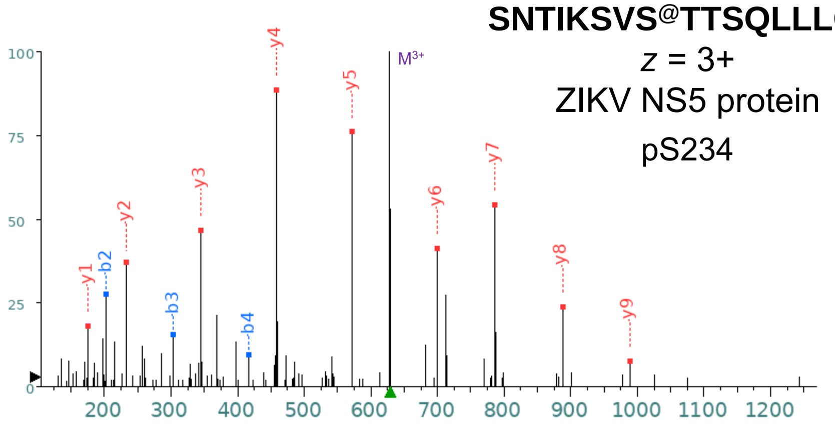


**SNTIKSVS@TTSQLLLGR**

$z$ = 3+

ZIKV NS5 protein

pS234

+1

| Seq | # | b: Δ Error | b | y | y: Δ Error | +1 |
|---|---|---|---|---|---|---|
| S | 1 | --- | 88.039 | --- | --- | 17 |
| N | 2 | -1.068 | **202.082** | 1797.947 | --- | 16 |
| T | 3 | 1.419 | **303.130** | 1683.904 | --- | 15 |
| I | 4 | -3.044 | **416.214** | 1582.856 | --- | 14 |
| K | 5 | --- | 544.309 | 1469.772 | --- | 13 |
| S | 6 | --- | 631.341 | 1341.677 | --- | 12 |
| V | 7 | --- | 730.409 | 1254.645 | --- | 11 |
| S@ | 8 | --- | 897.408 | 1155.577 | --- | 10 |
| T | 9 | --- | 998.455 | **988.579** | 4.241 | 9 |
| T | 10 | --- | 1099.503 | **887.531** | 1.159 | 8 |
| S | 11 | --- | 1186.535 | **786.483** | 2.097 | 7 |
| Q | 12 | --- | 1314.594 | **699.451** | -2.550 | 6 |
| L | 13 | --- | 1427.678 | **571.393** | -1.228 | 5 |
| L | 14 | --- | 1540.762 | **458.309** | -1.690 | 4 |
| L | 15 | --- | 1653.846 | **345.224** | -1.659 | 3 |
| G | 16 | --- | 1710.867 | **232.140** | -2.190 | 2 |
| R | 17 | --- | --- | **175.119** | 1.336 | 1 |

+2

| Seq | # | b: Δ Error | b | y | y: Δ Error | +1 |
|---|---|---|---|---|---|---|
| S | 1 | --- | 44.523 | --- | --- | 17 |
| N | 2 | --- | 101.545 | 899.477 | --- | 16 |
| T | 3 | --- | 152.069 | 842.456 | --- | 15 |
| I | 4 | --- | 208.611 | 791.932 | --- | 14 |
| K | 5 | --- | 272.658 | 735.390 | --- | 13 |
| S | 6 | --- | 316.174 | 671.342 | --- | 12 |
| V | 7 | --- | 365.708 | 627.826 | --- | 11 |
| S@ | 8 | --- | 449.208 | 578.292 | --- | 10 |
| T | 9 | --- | 499.731 | 494.793 | --- | 9 |
| T | 10 | --- | 550.255 | 444.269 | --- | 8 |
| S | 11 | --- | 593.771 | 393.745 | --- | 7 |
| Q | 12 | --- | 657.800 | 350.229 | --- | 6 |
| L | 13 | --- | 714.343 | 286.200 | --- | 5 |
| L | 14 | --- | 770.885 | 229.658 | --- | 4 |
| L | 15 | --- | 827.427 | 173.116 | --- | 3 |
| G | 16 | --- | 855.937 | 116.574 | --- | 2 |
| R | 17 | --- | --- | 88.063 | --- | 1 |

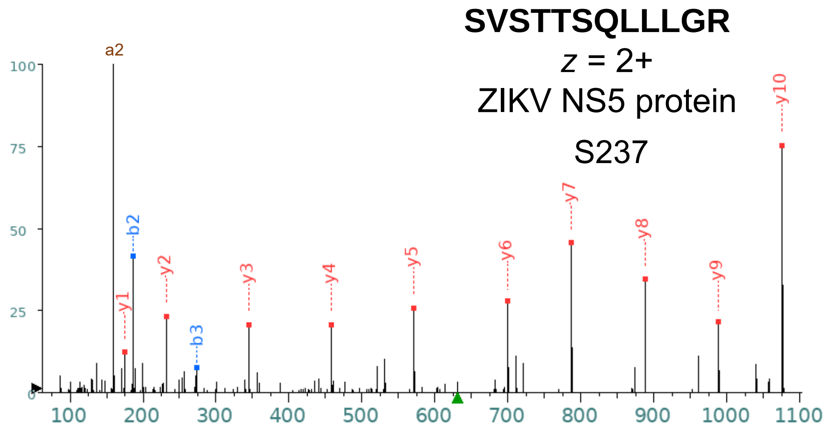


| Seq | # | b: Δ Error | b | y | y: Δ Error | +1 |
|---|---|---|---|---|---|---|
| S | 1 | --- | 88.039 | --- | --- | 12 |
| V | 2 | -0.035 | 187.108 | 1174.679 | --- | 11 |
| S | 3 | -0.359 | 274.140 | 1075.611 | -0.344 | 10 |
| T | 4 | --- | 375.187 | 988.579 | 1.216 | 9 |
| T | 5 | --- | 476.235 | 887.531 | -0.147 | 8 |
| S | 6 | --- | 563.267 | 786.483 | 1.088 | 7 |
| Q | 7 | --- | 691.326 | 699.451 | -0.980 | 6 |
| L | 8 | --- | 804.410 | 571.393 | 0.802 | 5 |
| L | 9 | --- | 917.494 | 458.309 | -1.423 | 4 |
| L | 10 | --- | 1030.578 | 345.224 | -2.189 | 3 |
| G | 11 | --- | 1087.599 | 232.140 | -0.941 | 2 |
| R | 12 | --- | --- | 175.119 | 1.336 | 1 |

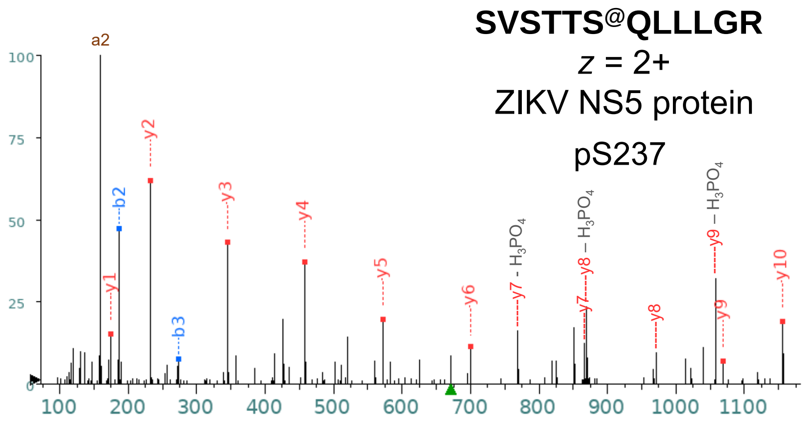


| Seq | # | b: Δ Error | b | y | y: Δ Error | +1 |
|---|---|---|---|---|---|---|
| S | 1 | --- | 88.039 | --- | --- | 12 |
| V | 2 | -0.850 | 187.108 | 1254.645 | --- | 11 |
| S | 3 | 0.087 | 274.140 | 1155.577 | 1.351 | 10 |
| T | 4 | --- | 375.187 | 1068.545 | -3.294 | 9 |
| T@ | 5 | --- | 556.201 | 967.497 | --- | 8 |
| S | 6 | --- | 643.233 | 786.483 | --- | 7 |
| Q | 7 | --- | 771.292 | 699.451 | -0.892 | 6 |
| L | 8 | --- | 884.376 | 571.393 | 1.122 | 5 |
| L | 9 | --- | 997.460 | 458.309 | -1.090 | 4 |
| L | 10 | --- | 1110.544 | 345.224 | -1.040 | 3 |
| G | 11 | --- | 1167.566 | 232.140 | -0.744 | 2 |
| R | 12 | --- | --- | 175.119 | 0.900 | 1 |

#### **SEHAETWFFDENHPYR**

*z* = 3+

ZIKV NS5 protein

T292

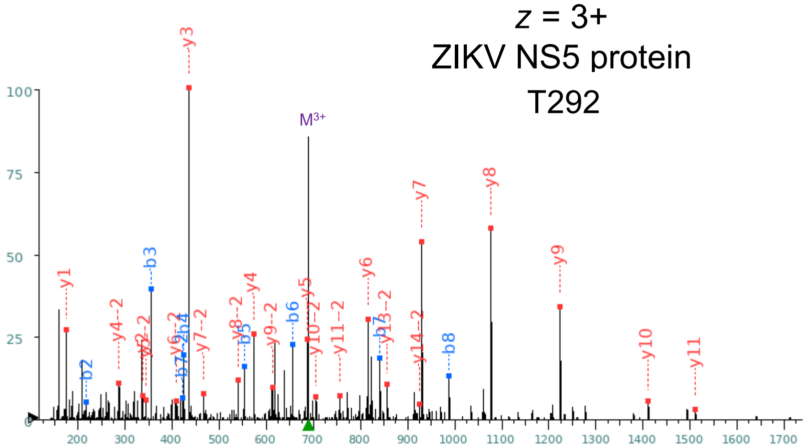


+1

| Seq | # | b: Δ Error | b | y | y: Δ Error | +1 |
|---|---|---|---|---|---|---|
| S | 1 | --- | 88.039 | --- | --- | 16 |
| E | 2 | -3.250 | 217.082 | 1977.851 | --- | 15 |
| H | 3 | -2.847 | 354.141 | 1848.809 | --- | 14 |
| A | 4 | -2.167 | 425.178 | 1711.750 | --- | 13 |
| E | 5 | -6.271 | 554.221 | 1640.713 | --- | 12 |
| T | 6 | -1.779 | 655.268 | 1511.670 | -1.098 | 11 |
| W | 7 | -1.782 | 841.348 | 1410.623 | 1.989 | 10 |
| F | 8 | -1.510 | 988.416 | 1224.543 | -1.125 | 9 |
| F | 9 | --- | 1135.484 | 1077.475 | -2.021 | 8 |
| D | 10 | --- | 1250.511 | 930.406 | -2.938 | 7 |
| E | 11 | --- | 1379.554 | 815.379 | -2.796 | 6 |
| N | 12 | --- | 1493.597 | 686.337 | -1.646 | 5 |
| H | 13 | --- | 1630.656 | 572.294 | 0.407 | 4 |
| P | 14 | --- | 1727.708 | 435.235 | -1.609 | 3 |
| Y | 15 | --- | 1890.772 | 338.182 | 3.250 | 2 |
| R | 16 | --- | --- | 175.119 | -1.888 | 1 |

+2

| Seq | # | b: Δ Error | b | y | y: Δ Error | +1 |
|---|---|---|---|---|---|---|
| S | 1 | --- | 44.523 | --- | --- | 16 |
| E | 2 | --- | 109.045 | 989.429 | --- | 15 |
| H | 3 | --- | 177.574 | 924.908 | -1.305 | 14 |
| A | 4 | --- | 213.093 | 856.379 | -0.725 | 13 |
| E | 5 | --- | 277.614 | 820.860 | --- | 12 |
| T | 6 | --- | 328.138 | 756.339 | -2.922 | 11 |
| W | 7 | 4.364 | 421.177 | 705.815 | -2.043 | 10 |
| F | 8 | --- | 494.712 | 612.775 | -0.587 | 9 |
| F | 9 | --- | 568.246 | 539.241 | -1.069 | 8 |
| D | 10 | --- | 625.759 | 465.707 | -1.376 | 7 |
| E | 11 | --- | 690.281 | 408.193 | -3.706 | 6 |
| N | 12 | --- | 747.302 | 343.672 | -2.018 | 5 |
| H | 13 | --- | 815.831 | 286.651 | -2.917 | 4 |
| P | 14 | --- | 864.358 | 218.121 | --- | 3 |
| Y | 15 | --- | 945.890 | 169.595 | --- | 2 |
| R | 16 | --- | --- | 88.063 | --- | 1 |

#### **SEHAET@WFFDENHPYR**

*z* = 3+

ZIKV NS5 protein

pT292

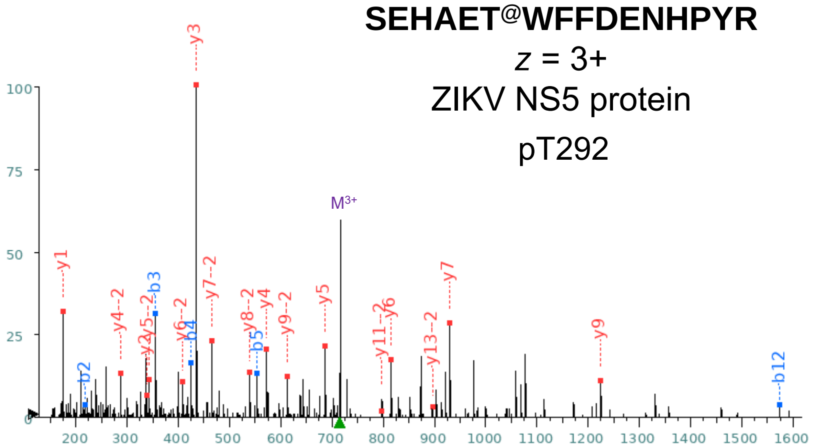


+1

| Seq | # | b: Δ Error | b | y | y: Δ Error | +1 |
|---|---|---|---|---|---|---|
| S | 1 | --- | 88.039 | --- | --- | 16 |
| E | 2 | -2.125 | 217.082 | 2057.818 | --- | 15 |
| H | 3 | -8.880 | 354.141 | 1928.775 | --- | 14 |
| A | 4 | -1.664 | 425.178 | 1791.716 | --- | 13 |
| E | 5 | -2.967 | 554.221 | 1720.679 | --- | 12 |
| T@ | 6 | --- | 735.235 | 1591.637 | --- | 11 |
| W | 7 | --- | 921.314 | 1410.623 | --- | 10 |
| F | 8 | --- | 1068.382 | 1224.543 | 2.264 | 9 |
| F | 9 | --- | 1215.451 | 1077.475 | --- | 8 |
| D | 10 | --- | 1330.478 | 930.406 | -1.954 | 7 |
| E | 11 | --- | 1459.520 | 815.379 | -5.641 | 6 |
| N | 12 | -1.486 | 1573.563 | 686.337 | 2.978 | 5 |
| H | 13 | --- | 1710.622 | 572.294 | 0.193 | 4 |
| P | 14 | --- | 1807.675 | 435.235 | -2.170 | 3 |
| Y | 15 | --- | 1970.738 | 338.182 | -2.616 | 2 |
| R | 16 | --- | --- | 175.119 | -1.888 | 1 |

+2

| Seq | # | b: Δ Error | b | y | y: Δ Error | +1 |
|---|---|---|---|---|---|---|
| S | 1 | --- | 44.523 | --- | --- | 16 |
| E | 2 | --- | 109.045 | 1029.413 | --- | 15 |
| H | 3 | --- | 177.574 | 964.891 | --- | 14 |
| A | 4 | --- | 213.093 | 896.362 | 5.083 | 13 |
| E | 5 | --- | 277.614 | 860.843 | --- | 12 |
| T@ | 6 | --- | 368.121 | 796.322 | -4.705 | 11 |
| W | 7 | --- | 461.161 | 705.815 | --- | 10 |
| F | 8 | --- | 534.695 | 612.775 | -0.786 | 9 |
| F | 9 | --- | 608.229 | 539.241 | -3.673 | 8 |
| D | 10 | --- | 665.742 | 465.707 | -2.490 | 7 |
| E | 11 | --- | 730.264 | 408.193 | -2.285 | 6 |
| N | 12 | --- | 787.285 | 343.672 | -1.751 | 5 |
| H | 13 | --- | 855.815 | 286.651 | -3.023 | 4 |
| P | 14 | --- | 904.341 | 218.121 | --- | 3 |
| Y | 15 | --- | 985.873 | 169.595 | --- | 2 |
| R | 16 | --- | --- | 88.063 | --- | 1 |

#### M*AVSGDDCVVKPIDDR

*z* = 2+

ZIKV NS5 protein

S663

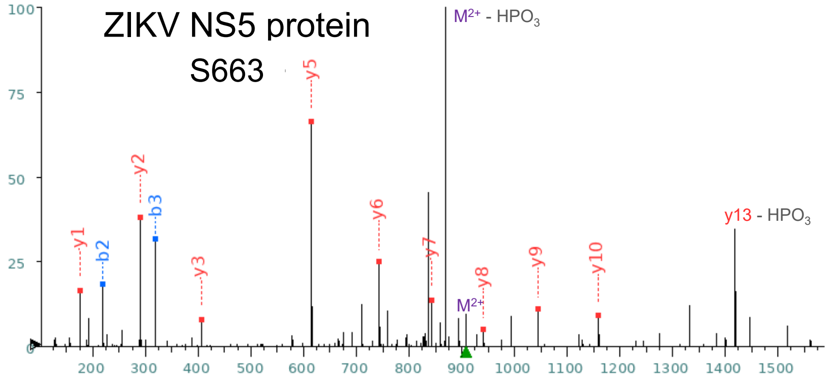


**Predicted Fragmentation Pattern**

| Seq | # | b: Δ Error | b | y | y: Δ Error | +1 |
|---|---|---|---|---|---|---|
| M* | 1 | --- | 148.043 | --- | --- | 16 |
| A | 2 | -4.113 | **219.080** | 1668.730 | --- | 15 |
| V | 3 | -4.826 | **318.148** | 1597.693 | --- | 14 |
| S@ | 4 | --- | 485.147 | 1498.624 | --- | 13 |
| G | 5 | --- | 542.168 | 1331.626 | --- | 12 |
| D | 6 | --- | 657.195 | 1274.605 | --- | 11 |
| D | 7 | --- | 772.222 | **1159.578** | -2.798 | 10 |
| C | 8 | --- | 875.231 | **1044.551** | -1.619 | 9 |
| V | 9 | --- | 974.300 | **941.541** | -2.543 | 8 |
| V | 10 | --- | 1073.368 | **842.473** | -3.140 | 7 |
| K | 11 | --- | 1201.463 | **743.405** | -4.142 | 6 |
| P | 12 | --- | 1298.516 | **615.310** | -4.322 | 5 |
| I | 13 | --- | 1411.600 | 518.257 | --- | 4 |
| D | 14 | --- | 1526.627 | **405.173** | 4.928 | 3 |
| D | 15 | --- | 1641.654 | **290.146** | -3.861 | 2 |
| R | 16 | --- | --- | **175.119** | -3.805 | 1 |

#### M*AVS@GDDCVVKPIDDR

*z* = 2+

ZIKV NS5 protein

pS663

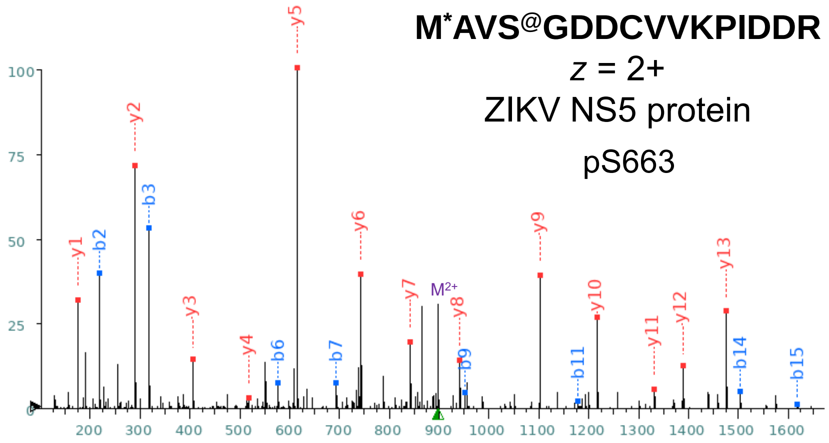


**Predicted Fragmentation Pattern**

| Seq | # | b: Δ Error | b | y | y: Δ Error | +1 |
|---|---|---|---|---|---|---|
| M* | 1 | --- | 148.043 | --- | --- | 16 |
| A | 2 | -2.790 | **219.080** | 1645.785 | --- | 15 |
| V | 3 | -3.291 | **318.148** | 1574.748 | --- | 14 |
| S | 4 | --- | 405.180 | **1475.679** | -2.674 | 13 |
| G | 5 | --- | 462.202 | **1388.647** | -0.348 | 12 |
| D | 6 | -2.538 | **577.229** | **1331.626** | -0.928 | 11 |
| D | 7 | -0.656 | **692.256** | **1216.599** | -3.051 | 10 |
| C^ | 8 | --- | 852.286 | **1101.572** | -2.403 | 9 |
| V | 9 | -0.800 | **951.355** | **941.541** | -1.441 | 8 |
| V | 10 | --- | 1050.423 | **842.473** | -3.937 | 7 |
| K | 11 | -2.861 | **1178.518** | **743.405** | -3.075 | 6 |
| P | 12 | --- | 1275.571 | **615.310** | -2.735 | 5 |
| I | 13 | --- | 1388.655 | **518.257** | -4.251 | 4 |
| D | 14 | -0.100 | **1503.682** | **405.173** | 5.154 | 3 |
| D | 15 | -2.334 | **1618.709** | **290.146** | -2.283 | 2 |
| R | 16 | --- | --- | **175.119** | -2.150 | 1 |
