## Supplementary figures and images for "A global map of the Zika virus phosphoproteome reveals host-driven regulation of viral budding and cytopathogenicity"

### Supplementary Information 4

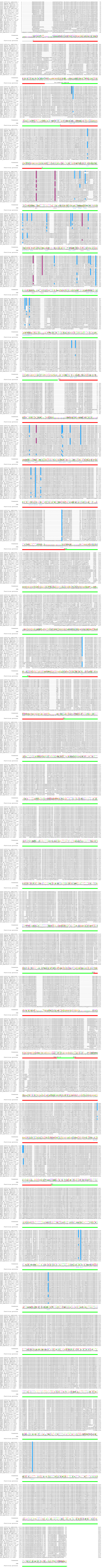
